# Compartmentalisation proteomics revealed endolysosomal protein network changes in a goat model of atrial fibrillation

**DOI:** 10.1101/2023.03.10.532119

**Authors:** Thamali Ayagama, Philip D Charles, Samuel J Bose, Barry Boland, David A Priestman, Daniel Aston, Georgina Berridge, Roman Fisher, Adam P Cribbs, Qianqian Song, Gary R Mirams, Lisa Heather, Antony Galione, Neil Herring, Ulrich Schotten, Rebecca A Capel, Frances M Platt, Frances M Platt, Holger Krame, Sander Verheule, Rebecca AB Burton

## Abstract

Endolysosomes (EL) are known for their role in regulating both intracellular trafficking and proteostasis. EL help facilitate elimination of damaged membrane and cytosolic proteins, protein aggregates, membranous organelles and also play an important role in calcium signalling. Despite the importance of EL, their specific role in cardiovascular disease is not well understood. In particular, it’s unclear how EL contribute to atrial pathology over longer time frames. To shed light on this question, we conducted a comprehensive analysis that involved proteomics, transcriptomics, integrated analysis, electron tomography, western blotting, and enzyme assays. To identify the role of EL in atrial fibrillation (AF), we applied a recently published organelle protein isolation method. We used this method to study biopsies from AF goat model and analyse the EL-specific proteins and pathways involved in this condition. Our results revealed the upregulation of the AMPK pathway and the expression of EL-specific proteins that were not found in whole tissue lysates (TL), including GAA, DYNLRB1, CLTB, SIRT3, CCT2, and muscle-specific HSPB2. We also observed structural anomalies, such as autophago-vacuole formation, irregularly shaped mitochondria, and glycogen deposition, which provide insights into the EL’s contribution to AF and related pathways and molecular mechanisms. Overall, our findings suggest that EL play an important role in the development of AF over longer time frames, and provide a more detailed understanding of the underlying molecular processes involved.

## Introduction

Atrial fibrillation (AF) is the most common sustained cardiac arrhythmia, accounting for around 14% of all strokes in the UK, and linked to a significantly high risk of developing ischaemic stroke (Workman *et al*., 2008; Samuthpongtorn *et al*., 2021). The prevalence of AF in the general population is 2%, although AF is age-dependent and this figure rises to 3.7-4.2% in ages 60-70 and 10-17% in the over 80s (Zoni-Berisso *et al*., 2014). AF is a progressive disease, progressing from paroxysmal to persistent forms, with self-sustaining progression driven by AF-triggered cardiac remodelling (Wijffels *et al*., 1995; Botto *et al*., 2003) and potentially progression of comorbidities associated with AF (Heijman *et al*., 2021). Atrial remodelling in AF can be the result of structural (Frustaci *et al*., 1997; Ausma *et al*., 2000) or electrical (Lee, 2013) changes, and organelle dysfunction is also observed in AF progression (Pool *et al*., 2021).

Regional and cell-type specific quantitative proteomics studies have enabled significant progress to be made in the understanding of the proteomic and transcriptomic contribution to AF pathology (Doll *et al*., 2017). However, the contribution of organelle remodelling in AF has not been studied extensively, including the potential contribution of changes in acidic organelles (Kim *et al*., 2021) such as lysosomes and endolysosomes, which play key roles in cellular energy metabolism (Halcrow *et al*., 2021) and the trafficking of cellular components (Bose *et al*., 2022; Halcrow *et al*., 2022). Lysosomal changes have long been linked to cardiac disease (Wildenthal & Decker, 1980), and changes in acidic organelles may be linked to underlying alterations in AF molecular pathways (Papathanasiou *et al*., 2021). Lysosomes may be linked to AF progression, for example via changes in autophagy (Yuan *et al*., 2018). Indeed, several studies have recently identified autophagy as a potential mechanism underlying cardiac remodelling in AF progression (Garcia *et al*., 2012; Yuan *et al*., 2018), and autophagy has been shown to be increased in AF patients (Yuan *et al*., 2018). However, little is known of how lysosomal proteins may be altered in AF. Techniques that enable proteomic characterisation at the level of individual organelles (Au *et al*., 2007; Ayagama *et al*., 2021), therefore, have the potential to highlight such changes. We have previously developed a modified density gradient method to isolate endolysosomal proteins, which increases the identification of endo-membrane proteins that are trafficked to acidic organelles (Ayagama *et al*., 2021), we applied this endolysosomal isolation method to conduct a proteomic study in a large animal AF model. We combined transcriptomics and proteomics to obtain the mRNA-protein correlation. In addition, we carried out high resolution electron microscopy imaging to confirm structural changes that have previously been reported in goat AF studies (Ausma *et al*., 1997; Ausma *et al*., 2001) and we specifically look at acidic organelles at the cellular level.

The endolysosomal (EL) system consists of a series of membranous vesicles, composed of early endosomes, recycling endosomes, late endosomes and lysosomes. Auto phagosomes are responsible for delivering the intracellular contents to lysosomes to complete autophagy. The endocytic pathway consists of acidification of the endosomes, maturation of endosomes to lysosomes accompanied by vesicle trafficking, protein sorting and targeted degradation of mostly sorted cargo. The two opposing sorting systems that are operating in these processes include the endosomal sorting complex required for transport (ESCRT, supports targeted degradation) and the retromer (supports retrograde retrieval of cargo). The EL system is emerging as a central player in a host of neurodegenerative diseases (Hu *et al*., 2015) and its relevance in other diseases including AF is now being explored.

The present study was conducted using the AF goat as the animal model. The goat model is an ideal substitute for human AF as it has better tolerance than most contemporary animal models for AF, and the goat model is more comparable physiologically to humans compared to other models (Schüttler *et al*., 2020). The goat is a suitable model for conducting long-term cardiac pacing to develop sustained AF (Wijffels *et al*., 1995), and has been successfully utilised for studying electrical, contractile and structural remodelling in sustained AF pathology (Schotten *et al*., 2003; Verheule *et al*., 2010; Verheule *et al*., 2013) Maesen *et al*., 2013). During prolonged pacing, the AF goat model has been shown to undergo structural remodelling through endomysial fibrosis (Verheule *et al*., 2013). Similar to humans, the AF goat model demonstrates the development of electrical conduction disturbances that give rise to complex activation patterns and endocardial-epicardial dissociation, providing a suitable substrate for the development of atrial arrhythmia (Eckstein *et al*., 2011; Verheule *et al*., 2013). Studies from Wijffels *et al*. (1999) and Eijsbouts *et al*. (2006) highlight the relevance of the AF goat model’s suitability for antiarrhythmic drug targeted studies.

Our integrated approach combines transcriptomics and proteomics to provide a comprehensive comparative analysis of omics data that includes post-translational data. Dysregulated proteins were identified by performing label-free quantitative mass spectrometric analysis of the AF goat peptides compared to the sham goat models. After identifying dysregulated proteins, molecular pathways were used to understand how these dysregulations potentially affected the lysosomes and acidic organelles. Molecular pathways were analysed using Cytoscape 3.7.2 with STRING, Kyoto Encyclopaedia of Genes and Genomes (KeGG), Gene Ontology (GO) and Reactome pathway annotations to predict the relevant pathways. Furthermore, using GO, an over-representation study was performed. The most significantly regulated proteins identified in the AF goat model were compared against the human proteome, and the highest represented biological processes and cellular components that were altered in the AF goat model were identified based on comparison against human data.

## Results

### Identifying differential protein expression by quantitative proteomic analysis

A density gradient approach (Ayagama *et al*., 2021) (see Star Methods) was used to isolate fractions corresponding to whole tissue lysate (TL), mitochondria (Mito) and endolysosomal lysate (EL) from sham (N=3) and AF (N=3) left atrial tissue biopsies from goat hearts. Tissue samples were then prepared as described in Star methods for liquid chromatography-tandem mass spectrometry (LC-MS). The differential protein-expression levels between AF and sham groups were identified by quantitatively analysing the mass spectrometric data of TL and EL using the Perseus software platform (Tyanova *et al*., 2016) (version 1.6.15.0) (Figure 1A and B). The protein intensity values of each biological replicate were separated into protein groups (AF and sham) and imported into Perseus. These data matrixes were filtered by removing proteins with more than two missing values and used for quantitative analysis.

**Figure 1:**
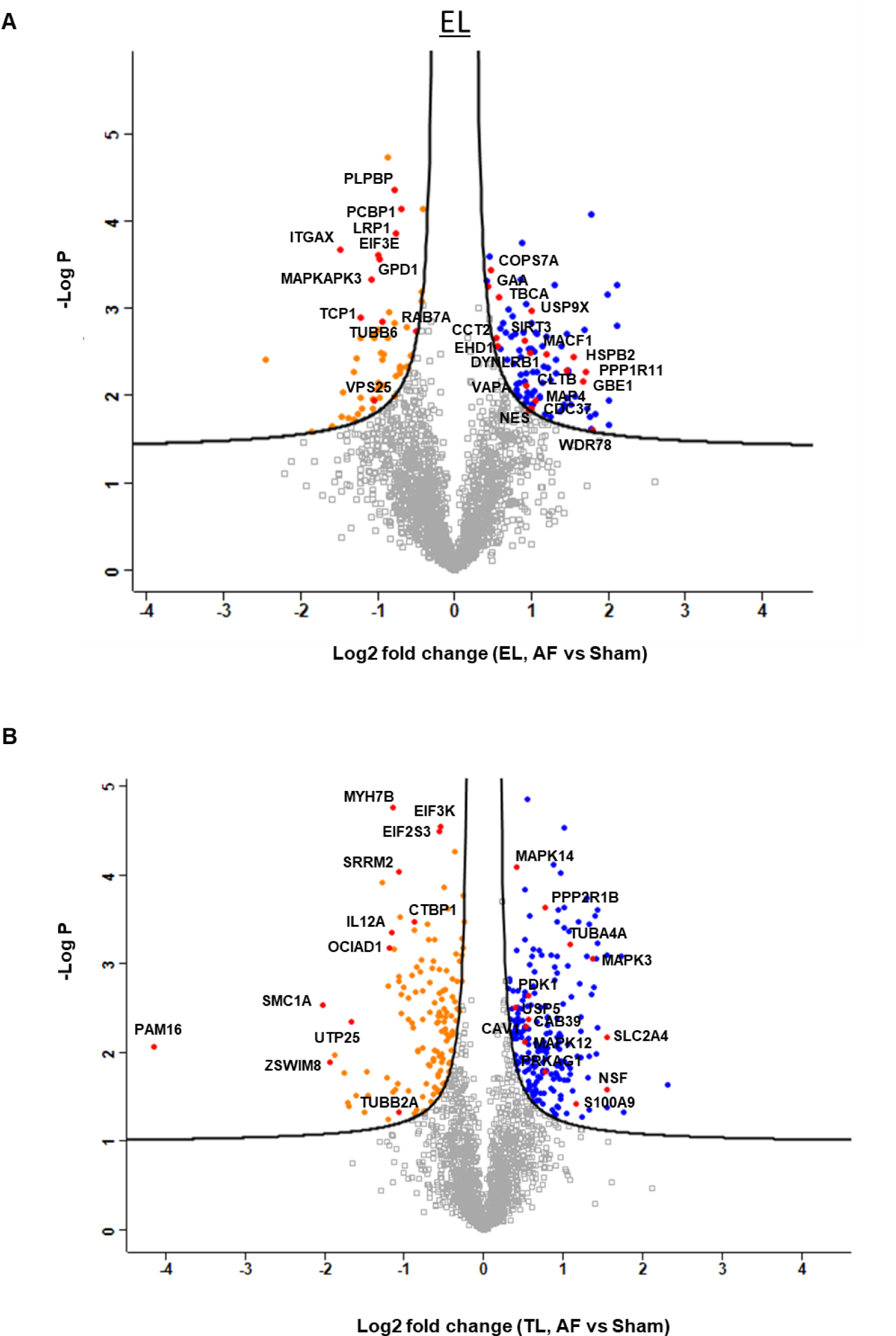
**A** and **B.** Volcano plot analysis of Endolysosome fraction (EL) and Tissue Lysate (TL) of the AF vs sham goat models showing significantly upregulated (Blue) and downregulated (Orange) proteins in the AF (*p*=0.05 and FDR =0.05). The –log2 transformed *p* values is plotted against the differentially regulated levels of Proteins in AF and sham.

After the filtration, TL and EL samples each remained with 2,104 proteins. Data were log-transformed (log2) and normalized via Z score. The missing data points of the matrix were imputed based on normal distribution. Volcano plots were generated for each TL and EL sample by applying a two-way Student’s t-test to identify the significant differences in protein regulations between AF and sham conditions (Figures 1A and 1B). The regulation levels were detected using a permutation-based false-discovery rate (FDR) of 5% with 250 randomizations at S0 = 0.1 and 99% confidence level.

Violin plots (Figures 2A) were created using the kernel density estimation indicating the underlying distribution of the protein intensities between and within sham and AF group biological replicates by samples, colour-coded with a gradient of purple colour, with higher protein intensity values presented as darker than the samples with comparatively lower intensities. Subsequently, Pearson coefficient correlation plots were created to observe the actual correlation between the groups (Figures 2B). Here the correlation ρ represents the interaction between two variables, or the AF and sham biological replicates, on a continuous scale of 1 to −1, where 1, depicts positive correlation, 0 depicts no correlation and −1 depicts negative correlation (Dytham, 2011). Heat maps (Figure 2C) were created using Euclidian distance and K-mean clustering of the normalised protein intensities obtained from the quantified protein data matrix. A total of 2104 proteins in EL was observed in the three (EL) protein clusters. The red and blue colour codes represent differential protein intensity levels and intensity whisker plots of each biological replicate were created to visualize the protein intensity distribution (Figure 2D). Principal Component Analysis (PCA) (Figure 2E) enabled observation of the vector distribution between and within the sample groups. In the PCA plots (Figure 2E), EL component 1 showed a 46.1% deviation respectively between AF (purple symbols) and sham (green symbols) groups. However, an exceptional segregation of 23.8% was observed in EL first biological replicates of sham groups, demonstrating that the variability between the differential experiment groups is more influential than the variability observed within the biological replicates. The most significantly regulated proteins of the EL fraction (Supplementary Table 1) were further analysed in STRING network to identify endolysosomal proteins with their respective structural or functional entities (Figure 3 and Supplementary Table 1).

**Figure 2:**
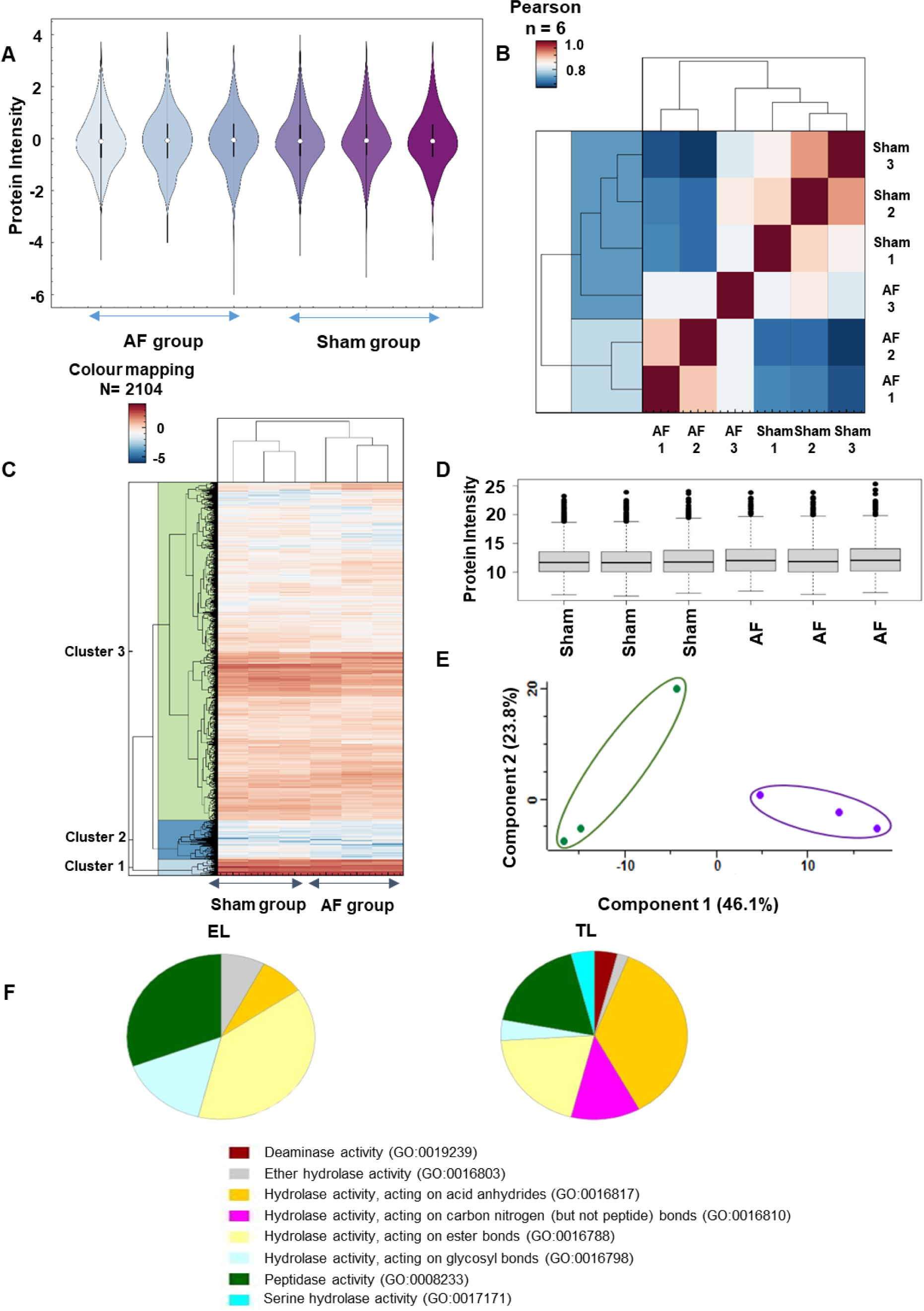
**A** Violin plot shows protein intensity levels in the triplicated AF vs sham EL samples. **B** Pearson co-efficient correlation plot explains the direct correlation between the sample protein intensities (1= highest and 0= lowest). **C** Heat map for the AF and sham EL Samples demonstrating the z scored intensities of the differentially expressed proteins after unsupervised hierarchical clustering. **D** Whisker plot of the EL AF vs sham protein intensity distribution shows the median, interquartile and the minimum to maximum outlier distribution. **E** Principal Component Analysis (PCA) demonstrating the spatial resolution among the averaged vectors of the AF vs sham goat EL samples, and component one and two variations presents respectively 46.1% and 23.8%. **F** Gene ontology panther pathway analysis for EL and TL fractions. The molecular function of the endo-lysosome fraction (EL) showed 25% of catalytic activity, compared to the tissue lysate (TL) which showed 26.3% of catalytic activity. The catalytic hydrolase activity was further analysed, and compared to TL (18%), EL fraction showed higher peptidase activity (30.8%), and hydrolases acting on ester bonds (TL=20%, EL= 38.5%)

**Figure 3:**
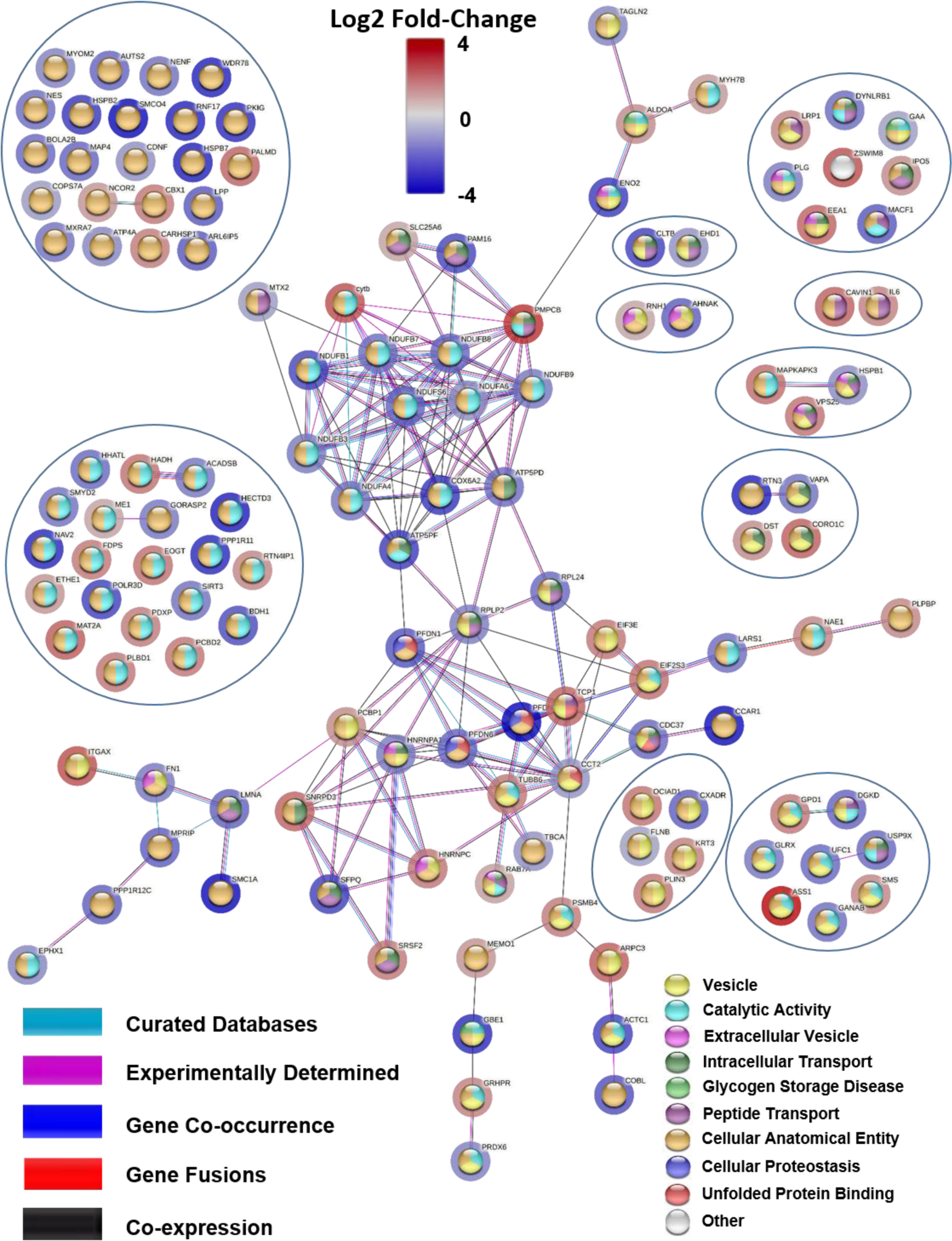
Endolysosomal proteins displaying network. Edges based on curated databases, experiments, gene co-occurrences, gene fusions, co-expressions and the nodes are coloured according to their functional enrichments. The halos around the nodes display the significant log2 fold-change.

### *The overrepresentation of Capra hircus* (goat) protein regulation enrichment comparison to total human proteome and to the C. *porcellus* (guinea pig)

The significantly up and down regulated protein/gene names with the respective log2 fold-change values were then uploaded to the Panther–Gene Ontology (GO), and an overrepresentation test was conducted against the reference *H. sapiens* proteome (Taxonomy ID: 9606) (Supplementary Figure 1A and B). This overrepresentation test was conducted on GO terms, Cellular Component (GOCC) and Biological Process (GOBP). The highest enriched GOCC term included muscle filaments, myofibrils proteasome complex, sarcoplasmic reticulum lumen, endocytic vesicles, and vesicle lumen (Supplementary Figure 1A). In contrast, the highest enriched GOBP term included the energy metabolism and vesicle-mediated trafficking pathways (Supplementary Figure 1B).

We performed a Venn analysis to understand the total protein yield of C. *hircus* TL and EL compared to the C. *porcellus* protein list that was published previously using this density-gradient based method from Ayagama *et al* 2021 (Supplementary Figure 1C). These comparisons showed a good overlap between goat and guinea pig data from these separate investigations, with 44.2% shared proteins for TL, and 28.8% shared proteins for EL samples. Due to the higher availability of annotation data for *H. sapiens*, the UniProt KB identifiers of the most significantly regulated *C. hircus* proteins identified from the TL volcano plot of the AF compared to the sham samples (Figure 1B) were converted to the *H. sapiens* identifiers (Ayagama *et al*., 2021).

### mRNA transcript analysis using next generation sequencing

#### Distance Matrix Heatmap

The distances between samples were analysed using hierarchical clustering to provide an overview of the similarities and dissimilarities between the samples. Sample distances were displayed using a range of shaded colour bar from dark blue to white, 0 being highly similar and displayed as dark blue and 200 being highly dissimilar represented in white colour (Supplementary Figure 2A).

#### Heatmap

The topmost regulated genes were plotted using hierarchical clustering to determine whether the samples cluster together according to the AF or sham conditions. All the AF and sham goat samples of LA tissue were clustered together, (Supplementary Figure 2B

#### PCA

PCA was performed to analyse the sample distribution and to assess the reliable correlation between the sample triplicates between AF and sham model groups by reduced data dimension for a simpler interpretation. As indicated in Supplementary Figure 2C, vector deviation of 67.3% was observed at component 1 (PC1) between AF (purple symbols) and sham/control groups (green symbols). An exceptional 11.8% segregation was observed in component 2 (PC2), or the biological replicates within the groups. Since, PC1 value was higher than PC2, it can be concluded that the variance between the two conditions is higher than the variance within a group (Supplementary Figure 2C).

#### Whisker Plot

The transcript intensity distribution between AF and sham goat model samples were observed using whisker plot (Supplementary Figure 2D)

The whole tissue lysates (same samples as used in proteomics and Western blotting) were used to isolate mRNA, genomic expression was quantified, and the differential expression plotted as a volcano plot (Figure 4A). These differentially expressed genes were fed into Reactome pathway analysis to study the pathway regulations. We observed a significant upregulation of 235 genes, while 297 genes were significantly down regulated (Supplementary File 1 Table 1).

**Figure 4:**
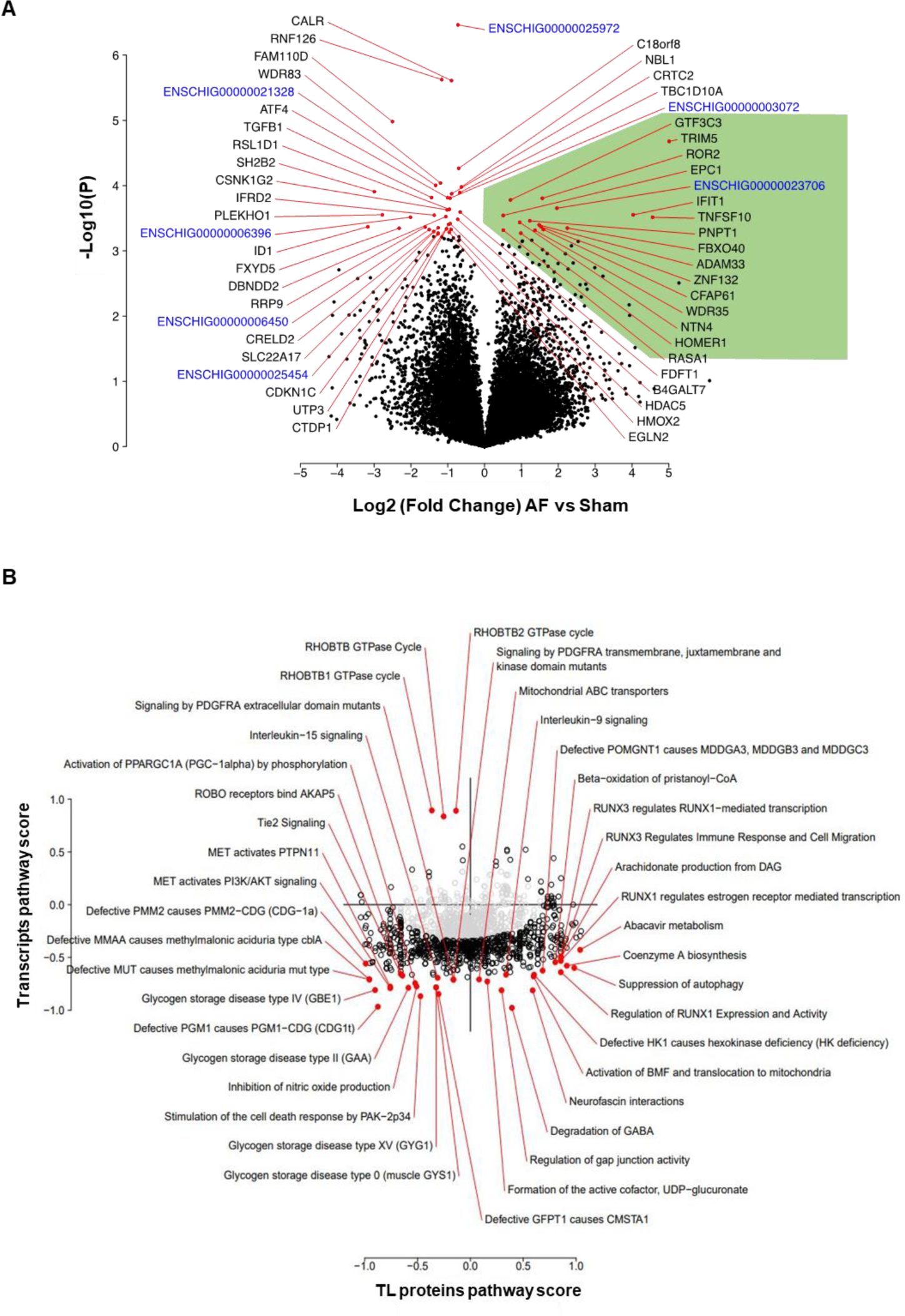
**A** Volcano plot of the most significantly regulated genes identified from the transcriptomics analysis**. B Integrated Analysis;** TL protein pathways vs mRNA transcript pathway scores. The pathways falling well under the horizontal line are up/downregulated more strongly in the proteomics than the transcriptomics vice versa for pathways falling above the line. grey = nonsignificant pathways, black= significant pathways red= highest-significant pathways (FDR=0.01) and the black horizontal line indicates y=x.

### Integrated Analysis of Transcriptomics and Proteomics

The integrated omics analysis highlighted several regulated pathways that were categorized under three confidence levels. These are colour coded in grey, black and red which represents non-confident, confident and highly confident pathways, respectively in Figure 4B and 5B. In our analysis and discussion, we do not consider non-confident pathways. Furthermore, the significance of these pathways were determined with a pathway score (PS) to show their enrichment levels in both transcriptomics and proteomics analysis. Interestingly, our integrated analysis highlighted RHOBTB GTPase, RHOBTB1 and RHOBTB2 GTPase cycles to be significantly enriched in transcriptomics. Whilst the integrated analysis did not highlight a significant enrichment in proteomics. The pathways, mitochondrial ABC transporters, interleukin (IL) 9 signalling, defective POMGNT1 causes MDDGA3, MDDGB3 and MDDGC3, beta-oxidation of pristanoyl-CoA, RUNX3 regulates RUNX1-mediated transcription, RUNX3 Regulates Immune Response and Cell Migration, arachidonate production from DAG, RUNX1 regulates estrogen receptor mediated transcription, abacavir metabolism, coenzyme A biosynthesis, suppression of autophagy, regulation of RUNX1 expression and activity, defective HK1 causes hexokinase deficiency (HK deficiency), activation of BMF and translocation to mitochondria, neurofascin interactions, degradation of GABA, and formation of the active co-factor, UDP-glucuronate are significantly downregulated in proteomics compared to transcriptomics. Some of the downregulated pathways in both the integrated TL proteomics and transcriptomics were, defective PMM2 causes PMM2-CDG (CDG-1a), defective MMAA causes methylmalonic aciduria type cblA, defective MUT causes methyl malonic aciduria mut type, glycogen storage disease types II, IV, XV and 0, defective PGM1 cause of PGM1-CDGII, MET activation of PI3K/AKT signalling, MET activation of PTPNII, inhibition of NO production, and stimulation of the cell death response by PAK-2P34 (labelled in red circles, Figure 4B and Supplementary file 2 Table 1).

**Figure 5:**
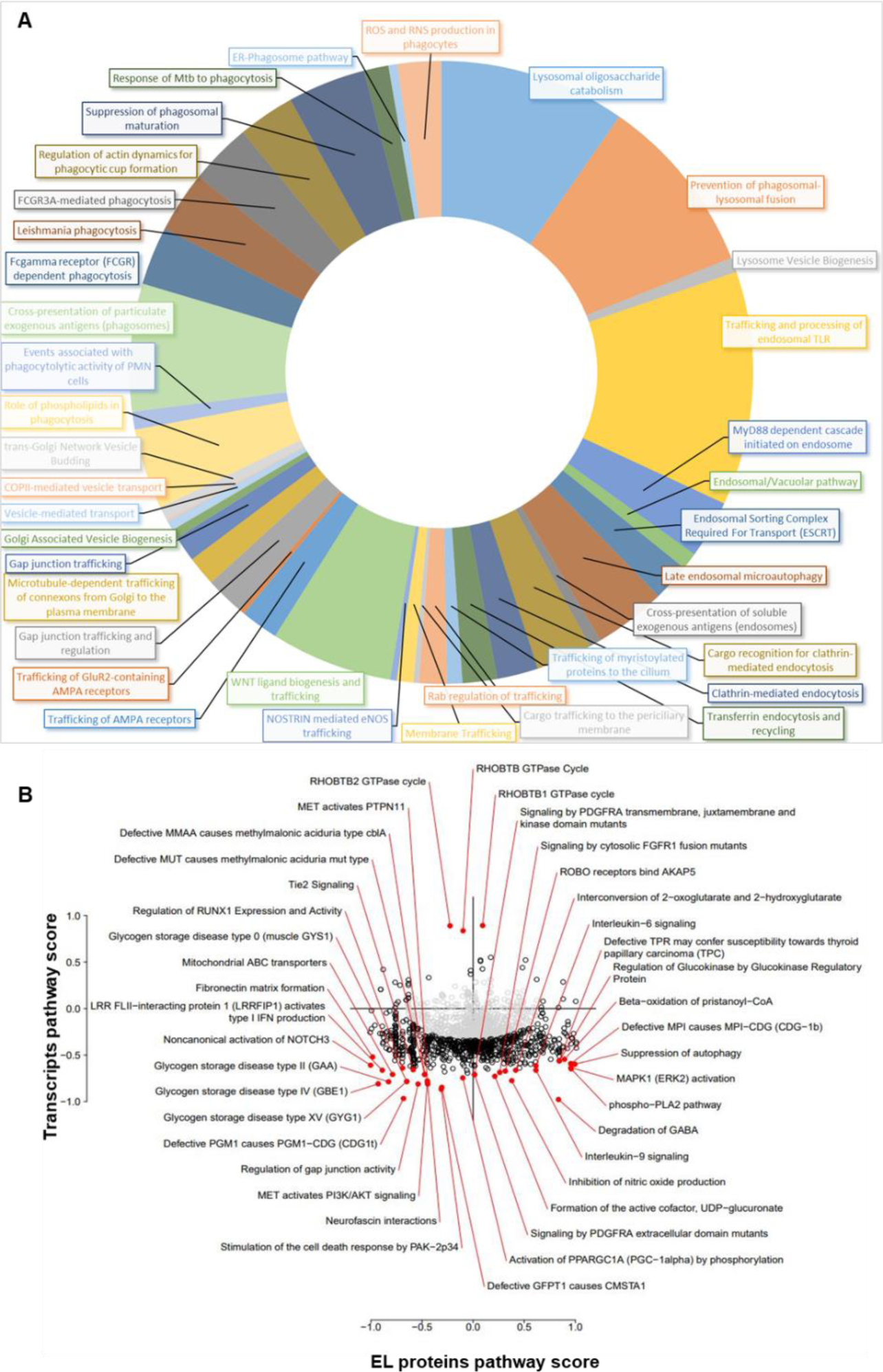
**A** Pathways representing the acidic organelles, vesicles and trafficking network from integrated analysis are represented using a donut chart against the enrichment scores. Data used to create this plot can be found in Supplementary File 2 Table 1. **B** Integrated analysis of EL protein pathways vs mRNA transcript pathway scores. Pathways represented in black and red are significantly altered (based on null hypothsis of no change) in either the transcriptomic data, proteomic data, or both. Pathways in the lower left and upper right (i.e. close to y=x) are regulated in the same direction in both modalities. Pathways in the upper left and lower right (i.e. close to y=-x) are regulated in the opposite direction in the two modalities. Grey = nonsignificant pathways; Black = significantly altered pathways (FDR<0.01); Red = top 40 most significant pathways.

Like the TL fraction, the integrated analysis of the most significantly up and down regulated EL proteins and genes of transcriptomics analysis showed RHOBTB and RHOBTB2 GTPase cycles up regulated. From the integrated analysis, the pathways such as, signaling by PDGFRA transmembrane, juxtamembrane and kinase domain mutants, signaling by cytosolic FGFR1 fusion mutants, ROBO receptors bind AKAP5, interconversion of 2-oxoglutarate and 2-hydroxyglutarate, IL-6 signaling, defective TPR may confer susceptibility towards thyroid papillary carcinoma (TPC), regulation of Glucokinase by Glucokinase Regulatory Protein, beta-oxidation of pristanoyl-CoA, defective MPI causes MPI-CDG (CDG-1b), suppression of autophagy, MAPK1 (ERK2) activation, phosphor-PLA2 pathway, degradation of GABA, IL-9 signalling, inhibition of NO production, formation of the active cofactor, UDP-glucuronate, and signaling by PDGFRA extracellular domain mutants were downregulated in proteomics compared to transcriptomics. Activation of PPARGC1A (PGC-1alpha) by phosphorylation, defective GFPT1 causes CMSTA1, stimulation of the cell death response by PAK-2P34, MET activation of PI3K/AKT signalling, glycogen storage disorders such as type II, IV,XV and 0, non-canonical activation of NOTCH3, defective PGM1 cause of PGM1-CDGII, neurofascin interactions, LRR FLII-interacting protein 1 (LRRFIP1) activates type I IFN production, fibronectin matrix formation, mitochondrial ABC transporters, regulation of RUNX1 expression and activity, Tie2 signaling, defective MMAA causes methyl malonic aciduria type cblA, defective MUT causes methyl malonic aciduria mut type and MET activates PTPN11 were the significantly down regulated pathways in, both proteomics and transcriptomics. The regulation of GAP junction activity pathway was downregulated in TL proteomics compared to transcriptomics, (Figure 4B, Figure 5 and Supplementary File 2 Table 1).

To identify and understand the overall representation of the pathways related to the endolysosomes, acidic organelles, and vesicle trafficking, the enrichment scores from the listed significant pathways of integrated proteomics and transcriptomics analysis were separated (Figure 5A and Supplementary File 2 Table 1). Furthermore, the integrated analysis of the most significantly up or downregulated EL, TL proteins and genes highlighted by the proteomic and transcriptomic analysis showed pathways related to the following as being confidently up regulated (Black in colour circled; Figure 4B and 5B and Supplementary file 2 table 1); membrane trafficking, vesicle mediated transport, TCA cycle and respiratory pathway, chaperone mediated protein folding, ER to Golgi anterograde transport, ER-Phagosome pathway, lipid metabolism, COPI mediated anterograde transport, while anabolic pathways such as rRNA processing, and cell cycle were among the down regulated pathways with confidence (the most significant list of pathways are summarised in Figure 4B, and 5B and the complete list of the pathways are shown in the supplementary file 2 table 1).

### Confirmation of selected proteins by Western blotting

The Ras-related protein Rab-11A (Rab11A) is expressed ubiquitously (Khvotchev *et al*., 2003; Lapierre *et al*., 2003) and plays important roles in intracellular transport. Rab11A was previously found to be significantly upregulated in the AF goat proteomics model. Western blotting was conducted on the same (N=3) biological samples retrieved from sham and AF goat models to evaluate this previous finding (Supplementary Figure 3A and B). GAPDH was used as the control protein, and intensities of Rab11A bands were normalized using the GAPDH band intensity. The normalized data are presented as mean±SD. The normalized band intensity for Rab11A was at 0.61±0.20 for the control group and 0.90±0.20 for the AF group. The Rab11A upregulation in AF was analysed by conducting a one-way t-test to analyse for upregulation. Although our data did not support a significant upregulation in Rab11A in the AF group, the *p*-value of 0.07 suggests a trend towards Rab11A upregulation in the AF group when compared to sham controls (N=3) (Supplementary Figure 3A and B). This difference between our data and data published previously by Lapierre *et al*. (2003) is likely the result of the lower power (N=3) of our data.

### Lysosome hydrolase activity assays

To check for impairment of autophagy, we conducted biochemical assays to investigate whether lysosomal enzyme activity is changed in goat AF. Lysosomal enzyme activity assays were performed on the three most common lysosomal enzymes: β galactosidase, β-hexosaminidase type A, and β hexosaminidase type B (Supplementary Figure 3C, 3D and 3E). The enzymatic activities were analysed using a two-way t-test and no significant difference was detected in any of the 3 lysosomal enzymes studied.

Altered lysosomal enzyme activities are observed in lysosomal storage disorders (LSDs) (Platt *et al*., 2018), and LSD patients show severe cardiac phenotypes in some LSDs (Nair *et al*., 2019). However, we did not observe such significant changes in the AF goat model, indicating little or no impairment in autophagy and this result is reinforced by the western blotting data on LC3I proteins (Supplementary Figure 4A).

### Structural insights using Electron Microscopy (EM)

We qualitatively analysed EM images from goat AF and sham control tissue and observed that the sarcomeres were regularly distributed throughout the cytoplasm including rows of uniformly sized mitochondria between them (Figure 6 A and C panels); similar to observations reported in (Ausma *et al*., 2003). In AF tissue, we observed increased myolysis (Figure 6 B and D), with areas depleted in sarcomeres and smaller, irregular mitochondria observed (Figure 6 D). Endolysosomes and autophagic vacuoles and autophagic-lysosomes are more commonly seen in AF tissue (Figure 6 E and more zoomed in detail in Figure 6 F). Since glycogen is associated with lysosomes or lysosome-like organelles (Geddes & Stratton, 1977), we quantitatively analysed the amount of glycogen accumulation in our AF and sham tissue samples, observing increased glycogen levels in AF samples (disease mean glycogen counts 378.7±42.6/μm^2 vs. control 190.5±24.9/μm^2, N=3, *p*=0.0002, student t-test; Supplementary Figure S5).

**Figure 6.**
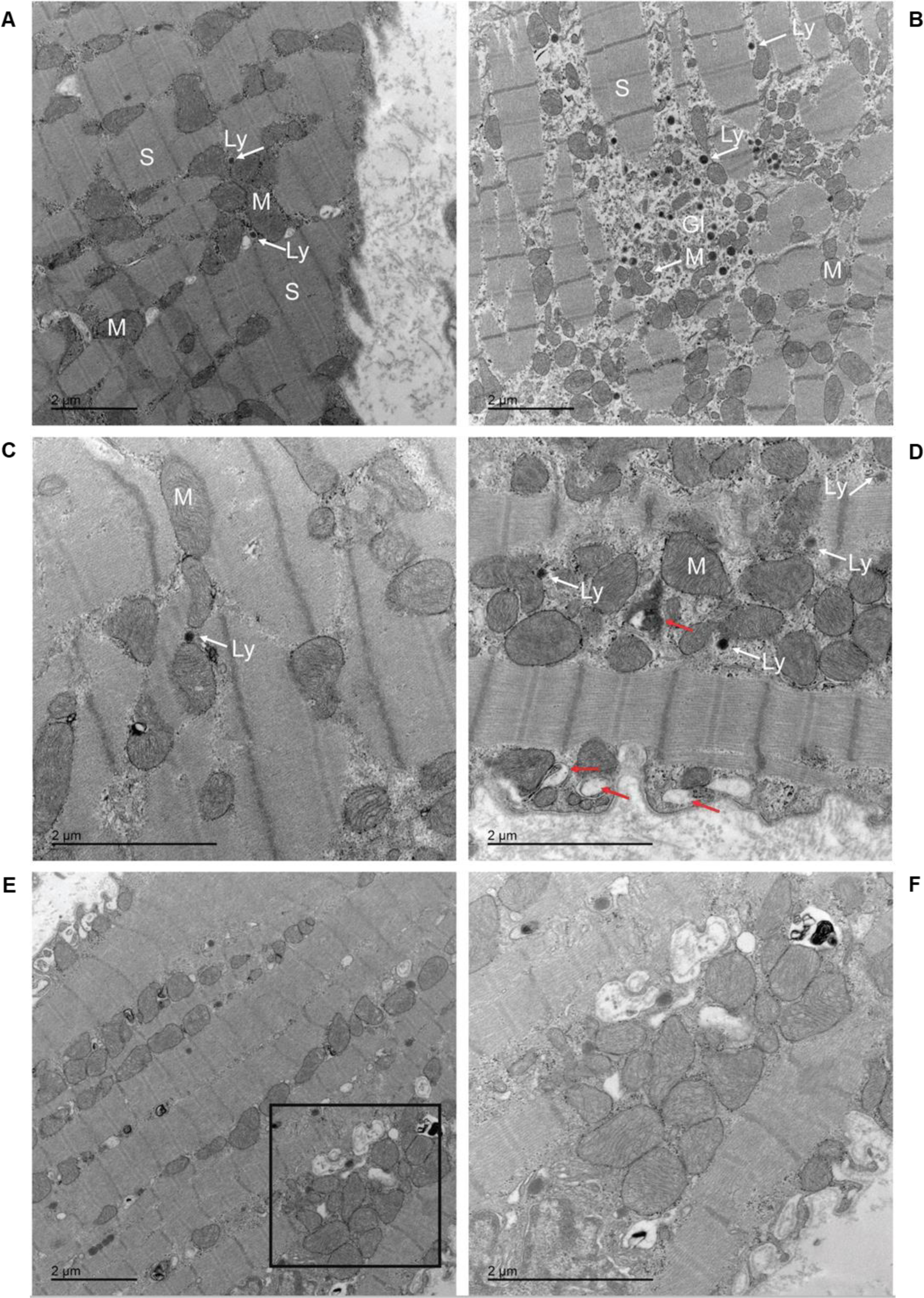
Electron microscopy images of goat left atrial myocardium tissue samples in sinus rhythm (sham, Panel A and C) and after prolonged sustained atrial fibrillation (Panel B, D, E and F). Panel A shows sarcomeres (S) regularly spaced and surrounded by mitochondria. Glycogen (Gl), lysosomes (Ly) and numerous irregular shaped mitochondria observed in atrial fibrillation sample (B). We observe many irregular shaped mitochondria dispersed in myolytic areas, autophagic vacuoles (red arrows) and lysosomes (Ly) in Panel D. Electron micrographs highlighting increased number of endocytic vesicles and vacuoles including endosomes, autophagosomes, lysosomes in atrial fibrillation samples (Panel E and F). Panel F is a higher resolution image of the marked square area in Panel E.

## DISCUSSION

Previous studies investigating mechanisms of AF using the goat model have shown structural, electrical, contractile and molecular changes during AF (some examples include Ausma *et al*. (2003), Wijffels *et al*. (1995) Allessie *et al*. (2002), van Hunnik *et al*. (2018) and Neuberger *et al*. (2006)) Proteomics studies have been performed in many cardiac diseases including AF (Liu *et al*., 2020) and reviewed by (De Souza & Camm, 2012) and have highlighted the need for more multi-omics research to investigate possible implicated molecular pathways in the development of AF. In this study we have focussed on changes in endolysosome-related proteins as another factor with a slower time course of development, and their involvement in this disease. Our endolysosomal organelle proteomics approach contributes new insight into regulation and differential functionalities related to these pathways and molecular mechanisms observed in this large animal model of AF. Here, we provide data related to pathway dysregulations in endolysosomal proteins previously not explored. Analysis of such proteins in separated tissue lysate (TL) and endolysosome fractions (EL) increased our ability to uncover endolysosome-specific proteins that were not identified in the TL, such as GAA, CLTB, DYNLRB1, SIRT3, CCT2, and muscle specific HSPB2 (Sugiyama *et al*., 2000) (See Supplementary File 3 Table 1 for the list of proteins). Combining an integrative multi-omics approach in this study helps us highlight interrelationships of the biomolecules and their functions in this disease and deriving insights into the data we have collected.

Lysosome number and dysfunction has been characterised in several cardiac conditions (Chi *et al*., 2020), including atrial septal defects, and AF is a common complication in these patients (Kottmeier & Wheat, 1967). Further, degenerative changes, including accumulation of lysosomes, have been found to correlate with atrial cellular electrophysiological changes (Mary-Rabine *et al*., 1983). Whilst AF is a common complication in atrial septal defect patients, no published studies have investigated the role of the endolysosomes or their corresponding interactive effects on other organelles in AF itself.

The major pathway regulations identified in this study were identified to be influenced from the AMPK signalling pathway (Supplementary Figure 6 and 7), selective autophagy (aggrephagy and proteasome pathway) (Supplementary Figure 8 and 9), NADPH oxidase pathway (Supplementary Figure 10), OXPHOS (Supplementary Figure 11), gap junction assembly and degradation, ESCRT, protein processing and folding pathway, vesicle-mediated transport, and lysosome vesicle biogenesis (Supplementary Figure 9, 10, 12 and 13). Most of the proteins and interpreted pathways identified in our study are increasingly being recognised in cardiac pathology (Papathanasiou *et al*., 2021; Sygitowicz *et al*., 2021), such as gap junctional remodelling (van der Velden *et al*., 2000), mitochondrial-bioenergetics and proteostasis (Papathanasiou *et al*., 2021), NADPH oxidase (Kim *et al*., 2005), and metabolic stress caused by oxidative stress (Manna & Jain, 2015).

Comparative analysis of LC3-I and II levels found that in AF tissue, levels of LC3-I were almost completely depleted, while no detection of the lipidated autophagic vacuole (AV)-associated LC3-II was observed. This suggests that high levels of autophagic flux may be occurring in the AF tissue, where the efficient clearance of AVs by lysosomes, outpaces the supply of LC3-I. We further tested by Western blotting the levels of p-p70 (p-P70:Total p70) to provide insights into mTOR activity. This was measured using a p-P70(Thr 389):Total p70 ratio, where Thr389 is an mTOR specific epitope. The p-p70 was unchanged, which suggests the heightened autophagy is not mTOR-dependent.

The contemporary transcriptomic and proteome of studied samples may vary substantially in terms of overall correlation between transcript and associated protein. In static cell contexts with minimal temporal dynamics, overall correlation can be quite strong (Lundberg *et al*., 2010), but in cells undergoing dynamic response to stimuli or stress, correlation can vary from strongly correlated to almost uncorrelated on a gene by gene basis (Buccitelli & Selbach, 2020) . To integrate across proteomic and transcriptomic results, we adopted the method of (Cox & Mann, 2012) Cox and Mann to compare changes in terms of pathway up or down regulation relative to a global median across all pathways with identified members.

This approach synthesises pathway changes across all genes/proteins and therefore allowed for more meaningful comparisons of the implication of changes in the transcriptome versus proteome. The approach revealed that, while there was a general downregulation of pathways the endolysosome as scored by mRNA levels, this drop was not consistently reflected by changes in the same pathways as scored by proteins. This is partially reflective of the time lag between response at the mRNA level versus response at the protein level, determined by regulation factors, in particular protein translation and turnover rates that vary from protein to protein.

Bioinformatics analysis of our proteomics results suggests that AMPK upregulation is triggered through an ATP depletion path. V-ATPase proton transporters utilise ATP to conduct protons which acidify the lysosomes or endolysosomes (Godbey, 2014), which is pivotal for the lysosomal enzymes to be functionally activated. For example, dysfunctional V-ATPases lead to neurodegenerative disorders (Song *et al*., 2020) caused by poor substrate digestion in the lysosomes. The increased ATP consumption in tachyarrhythmias (Nesheiwat *et al*., 2022), such as AF, is expected to create an energy demand in the atria (Rennison *et al*., 2021). As the cell’s primary energy source, ATP drives active and coupled membrane ion transport (HASSELBACH, 1964) to maintain cellular ion homeostasis (Grant, 2009).

### Exploring the functional endolysosomal network proteins in AF

The endolysosomal String network (Figure 3) consists of 134 nodes and 171 edges with an average node degree of 2.55. The functional enrichments in the network were determined using the GO database. 25 proteins from the biological process of intracellular transport and 24 proteins related to peptide transport were identified in the EL network. Six proteins were associated with unfolded protein binding, and 58 proteins were identified for catalytic activity. Furthermore, 48 proteins were components of vesicles, and 11 were components of extracellular vesicles. 118 proteins were part of the anatomical entity of the cell, and 3 proteins were from cellular proteostasis. Notably, 2/8 proteins that regulate the lysosomal storage disorder glycogen storage disease were highlighted in the EL network of AF and 2 more (PYGB and AGL) were significantly upregulated in TL (Figure 3, Supplementary Table 1).

#### (i) Endosomal sorting complex required for transport (ESCRT)

Our analysis suggests lower expression levels of proteins involved in the ESCRT pathway in the AF diseased condition (down-regulation of ESCRT-II complex and its cargo). Proteins such as Ras-related protein Rab-7a (RAB7A), and Vacuolar protein-sorting-associated protein 25 (VPS25), were downregulated by −0.50 and −1.04 log2 fold-change respectively (Supplementary Table 1). RAB7A, a recruiting protein for the tethering molecules (Gutierrez *et al*., 2004) (Vanlandingham & Ceresa, 2009), plays an essential role in the late endosomes to multivesicular bodies (MVB) transition in the ESCRT pathway. Rab-7 is a common modulator/participant in endocytosis and autophagy (Hyttinen *et al*., 2013). Moreover, RAB7A participates in the lysosome biogenesis through autophago-lysosome formation (Press *et al*., 1998). Vacuolar protein-sorting-associated protein 25 (VPS25) is a leading regulatory component of the ESCRT complex-II that sorts the endosomal cargo proteins to the MVB (Im *et al*., 2009) (Figure 3 and Supplementary Table 1).

Lysosomal alpha-glucosidase (GAA) was log2 0.44-fold upregulated in the EL fraction of AF. GAA functions as an enzyme that degrades glycogen in the lysosome (Zirin *et al*., 2013; Adeva-Andany *et al*., 2016; Roig-Zamboni *et al*., 2017) (Figure 3 and Table 1). Studies have indicated that a substantially high recombinant human GAA level is required to minimize the abnormal glycogen storage in the skeletal and cardiac muscles (Van den Hout *et al*., 2001; Zhu *et al*., 2004) (Figure 3 and Table 1).

1,4-alpha-glucan-branching enzyme (GBE1) is an enzyme that functions as a vital component in glycogen biosynthesis. GBE1 was upregulated by log2 1.67-fold in AF goat model compared to that of sham model (Figure 3 and Supplementary Table 1).To increase the glycogen molecule solubility, GBE1 generates α-1,6-glucosidic branches from α-1,4-linked glucose chains (Froese *et al*., 2015).

#### (ii) Lysosome motility and motor-protein based transport

Lysosome motility is mediated by motor proteins (Bandyopadhyay *et al*., 2014) such as kinesin and dynein and microtubules (Levi *et al*., 2006; Cabukusta & Neefjes, 2018). Multiple biological processes, such as degradation of macro molecules (de Duve, 2005), cellular homeostasis such as autophagy, signal transduction and metabolic signalling for ATP energy and amino acids, intracellular organelle signalling (Capel *et al*., 2015; Aston *et al*., 2017; Wong *et al*., 2019), require lysosomes to move and position themselves throughout the cytoplasm (Pu *et al*., 2016). We identified several motor proteins that are associated with lysosome motility that were upregulated, such as tubulins MACF1 up by log2 1.19-fold, Microtubule-associated protein (MAP4) up by log2 1.05-fold, Tubulin-specific chaperone A (TBCA) up by log2 0.58-fold and we also identified TUBB6 (tubulin beta 6 class V) in EL proteomics fraction and it was found to be downregulated log2 −0.95-fold Clathrin light chain B (CLTB) was upregulated by log2 1.5-fold in the AF condition. CLTB, along with Vesicle-associated membrane proteins (VAMPs) and adaptor protein complex 1 (AP1) play an essential role in lysosome membrane biogenesis (Saftig & Klumperman, 2009) (Figure 3 and Supplementary Table 1). Dynein light chain roadblock-type 1 (DYNLRB1) was upregulated by log2 0.98-fold. DYNLRB1 is an essential protein for general dynein-mediated transport and has been shown to be vital for sensory neuron survival (Terenzio *et al*., 2020). Furthermore, DYNLRB1 links dynein with adaptor proteins to regulate dynein and cargo for ideal cellular vesicle transport (Vaughan & Vallee, 1995). Simultaneously, Cytoplasmic Dynein 1 acts as a motor protein for the intracellular motility of organelles and retrograde motility of vesicles along the microtubules (Ansar *et al*., 2019) (Figure 3 and Supplementary Table 1). WDR78, a Dynein-f associated motor protein required for the axonemal localization (Zhang *et al*., 2019) was upregulated by log2 1.8-fold. Nestin (NES, upregulated by log2 0.96-fold in AF goat) is an intermediate filament (IFs) involved in vesicle-based communication, vesicle interaction and trafficking (Lasič *et al*., 2020). Similarly, vesicle-associated membrane protein-associated protein A (VAPA), a protein involved in vesicle trafficking, was upregulated by log2 0.93-fold in the AF goat model (Weir *et al*., 2001) (Figure 3 and Supplementary Table 1) and is a major ER-lysosome anchor protein (Zhao *et al*., 2018). Meng Lu et al. showed that abolished VAPA-mediated anchoring compromises ER remodelling and significantly increases the speed of lysosome motility (Lu *et al*., 2020). Eps15 homology (EH) domain-containing protein 1 (EHD1) was upregulated by log2 0.6-fold in the EL fraction. EHDs play critical roles in endosome-based membrane protein targeting (Gudmundsson *et al*., 2010) . EHD1 is a retrograde trafficking mediator and a regulator of the cluster of differentiation 44 protein (CD44) that participates in endocytic recycling and lysosomal degradation (Liu *et al*., 2022) (Zhang *et al*., 2012). Gudmundsson *et al* (2010) found modulation of EHD expression during myocardial infarction, which suggests that these proteins may play important roles in regulating membrane excitability (Gudmundsson *et al*., 2010) (Figure 3 and Supplementary Table 1).

### Intermediate protein networks between the whole cell and endolysosomes

#### Proteasome and aggresome pathway

The proteasome pathway is part of the selective autophagy process (Kraft et al., 2010) and several dysregulated proteins in our EL disease fraction were from this pathway (Supplementary Figure 8). Protein degradation through ubiquitination is termed as proteasome or aggresome pathway. Aggregated and misfolded proteins are identified by chaperone proteins and tagged by ubiquitination before protein degradation. Proteins related to both aggresome and proteasome formation were found to be up-or downregulated in our AF samples, including; E3 ubiquitin-protein ligase PPP1R11 (PPP1R11) up by log2 1.7, tubulin α chain (TUBA4A) up by log2 1.09, ubiquitin carboxyl-terminal hydrolase 5 (USP5) up by log2 0.39, probable ubiquitin carboxyl-terminal hydrolase FAF-X (USP9X) up by log2 1.00-fold, tubulin β chain (TUBB2A) down by log2 −1.07-fold (Supplementary Table 1 and Supplementary File 1 Table 2). PPP1R11 and USP5 are ubiquitination triggering proteins, and USP9X is both a deubiquitinase, that prevents a protein from the removal of conjugated ubiquitin and a ubiquitin precursor processor (Dupont *et al*., 2009); (Zhang *et al*., 2018) (Supplementary Table 1). The upregulated aggresome formation process that we observed in AF goat model is a well-known factor for the changes in lysosome distribution and its motility (Zaarur *et al*., 2014). Furthermore, Chaperonin containing TPC1 subunit 2 (CCT2) functions independently of ubiquitin and the TRiC complex to facilitate autophagic clearance of solid protein aggregates (Zhang & Klionsky, 2022). We detect modestly upregulated levels log2 (0.55) of CCT2 in the EL fraction. In addition to CCT2, we also observe upregulated levels log2 (1.00) of CDC37 and log2 (0.48) COPS7A in the EL fraction. Hsp90-CDC37 complex appear to participate in upstream of autophagy activation for the control of protein quality (Calderwool, Murshid *et al* 2009), and COPS7A is a protein from the COP9 signalosome complex (CSN) that mediates de-neddylation to regulate the ubiquitin conjugation pathway and is presumed to participate in structural remodelling that plays a crucial part in the developing stages of AF (Liu *et al*., 2020). .

#### (iv) AMPK upregulation

AMPK pathway is a central regulator of cellular metabolisms that is activated mainly by reduced adenosine triphosphate (ATP) levels in the cell (Supplementary Figure 6 and 7) and cascades a series of downstream chemical reactions in the cell that reprogram metabolism, autophagy (Li & Chen, 2019), cell polarity and growth (Shackelford & Shaw, 2009) As a result, catabolic pathways are upregulated while inhibiting the anabolic pathways to reduce the cellular ATP consumption level (Mihaylova & Shaw, 2011).

Our Integrated proteomic and transcriptomic data showed a significant upregulation in AMP-activated protein kinase (AMPK) pathway related proteins in AF samples. 5’-AMP-activated protein kinase subunit gamma-1 (PRKAG1/AMPK) log2 0.8-fold, calcium-binding protein 39 (CAB39/MO25) log2 0.54 -fold, Serine/threonine-protein phosphatase 2A regulatory subunit A beta isoform (PPP2R1B/PP2A) log2 0.77-fold and Protein-serine/threonine kinase (PDK1) log2 0.56-fold from AMPK pathway were observed to be upregulated in AF samples (Figure 1B, Supplementary figure 6, 7 and Supplementary File 1, Table 2). These observed changes suggest a link between AMPK regulatory pathway protein upregulation and AF (Chen *et al*., 2011; Garcia *et al*., 2012; Czegledi *et al*., 2019).

The identified AMPK pathway proteins are from AMPK activation through the ATP pathway. CAB39/MO25 is part of the liver kinase B1 (LKB1) and STE-related adaptor protein STRAD protein complex (Zeqiraj *et al*., 2009), which binds and activates STK11/LKB1. AMPK protein activity is controlled by LKB1, acting as a key upstream regulator for AMPK phosphorylation. PPP2R1B assembles the catalytic subunits of AMPK, and signals from insulin to PKB/AKT1 are transduced by PDK1 through activated phosphorylation. This downstream signalling cascade targets cell survival, glucose and amino acid uptake and glucose storage (Pullen *et al*., 1998). Therefore, processes such as upregulation of ATP production, activation of glucose intake, inhibition of the cell proliferation and growth, autophagy activation, cytoskeletal remodelling, DNA damage response, and apoptosis (Mihaylova & Shaw, 2011) are regulated by AMPK. The increased expression of the CAB39, AMPK, PP2A and PDK1 proteins provides further support for the upregulation of the AMPK signalling pathway in AF. Of interest is the finding by Christ et al. 2004 (Christ *et al*., 2004), related to molecular mechanism of electrical modelling, where they showed an increased activity of PP2A (protein phosphatase 2A) results in hypophosphorylation of the calcium channed *I_CaL_* (see also Review (Schotten *et al*., 2011).

#### (v) Ras-homologous guanosine triphosphatases (Rho GTPases) activation of Nicotinamide adenine dinucleotide phosphate oxidase (NADPH oxidase)

We observed increased levels of proteins from the MAP Kinase signalling network, which represent the RHO-GTPase activation of NADPH Oxidase. These regulated proteins are: Mitogen-activated protein kinase 3 (MAPK3) up by log2 1.37-fold; Mitogen-activated protein kinase 12 (MAPK12) up by log2 0.51-fold; Mitogen-activated protein kinase 14 (MAPK14) up by log2 0.42-fold; and Protein S100-A9 (S100A9) up by log2 1.17-fold (a complete list of the most significantly regulated TL proteins identified in the AF goat model are provided in Supplementary File 1 Table 2). MAPK proteins are reported to conduct signalling in mitochondria, Golgi, endoplasmic reticulum (ER) and endosomes (Mor & Philips, 2006). Rho family small GTPase proteins activate NADPH Oxidase (NOX), which activates the leading cell stress-response signalling network MAPK (see integrated omics Figure 4B, 5B, and Supplementary Figure 7 and 10). NOX signalling is a cellular stress-responsive mechanism in the cardiovascular system (Jiang *et al*., 2011) (Schröder *et al*., 2017), causing the production of superoxide, a reactive oxygen species, during cellular stresses initiated by biological, physical or chemical triggers. Cytochrome b-245 heavy chain (CYBB) or NADPH Oxidase 2 is a membrane-bound enzyme that generates superoxide (Upregulated by log2 2.06-fold change in our transcriptomics data, a complete list of the most significantly regulated genes identified in the AF goat model mRNA analysis are provided in Supplementary File 1 Table 1). The Yoo S et al. 2020 study showed that the oxidative injury by the CYBB/NOX2 caused an electrical remodelling by upregulating the constitutively active acetylcholine dependent potassium current (IKACh) in the canine model (Yoo *et al*., 2020). Significantly higher NOX-induced superoxide levels have been observed in AF patients, and unlike NOX2, the superoxides produced by dysfunctional NOS had a lesser contribution to electrophysiological remodelling and oxidative injury in the atria of AF patients. (Kim *et al*., 2005) and these superoxides further worsen the disease condition (Dudley *et al*., 2005; Antoniades *et al*., 2012; Youn *et al*., 2013; Mighiu *et al*., 2021).

MAPK3 is a signal transducing protein that regulates transcription, translation (Sha *et al*., 2017), and cell cycle-related functions such as the arrangement of the cytoskeleton and cell-cell adhesion during the cell survival state (Pullikuth & Catling, 2007; Shi *et al*., 2012). Furthermore, MAPK/ERK participate in lysosomal dynamics (Takemasu *et al*., 2019) and endosomal recycling (He & Kogut, 2003). MAPK12 is another signal transducing protein that, is triggered by extracellular stress stimuli, such as pro cytokines (Enslen *et al*., 1998) (Supplementary Figure 10).

MAPK14 is also stimulated by inflammatory triggers (Xu & Derynck, 2010) that conduct the cellular protein turnover for degradation through proteasomes (Qi *et al*., 2007), and S100A9 is a calcium- and zinc-binding protein that plays a prominent role in the inflammatory and immune response (Simard *et al*., 2010); The observed protein changes in this pathway, indicate increased cellular stress status in our AF goat model (Supplementary Figure 10) highlighting the relevance of these findings relating to the emerging role of NOX pathways in AF (Youn *et al*., 2013).

### Identifying differential mRNA expression by quantitative transcriptomic analysis

We performed transcriptomics as a confirmatory and supplementary method to our proteomics screen in our goat AF model (Figure 4A). The transcriptome approach has allowed for the detection of unbiased molecular changes in AF (Steenman, 2020). There is evidence for inter-related pathways such as oxidative stress, inflammation, thrombogenesis and fibrosis (Steenman, 2020) and more recently autophagy (Çubukçuoğlu Deniz *et al*., 2021). Our transcriptomic analysis identified upregulation of major ion channel Potassium/sodium hyperpolarization-activated cyclic nucleotide-gated channel 1 (HCN1) by log2 2.80-fold (Supplementary File 1, Table 1). HCN1 gain of function promotes AF (Fenske *et al*., 2013; Rivolta *et al*., 2020). Furthermore, Glycogen debranching enzyme (AGL) was upregulated by log2 2.85-fold change (Supplementary File 1, Table 1).

Reactome analysis (www.reactome.org) was performed on the differentially expressed genes (the most significantly regulated proteins and genes are presented in a hierarchical visualization of pathways using space filling graphs in Supplementary Files 4, 5 and 6); KEGG pathways of proteomics as well as transcriptomics and integrated Reactome pathway analysis highlighted AMPK signalling pathways as significantly upregulated (Supplementary Figure 6, Supplementary File 1 Table 1 and a complete list of pathways from the integrated EL, TL proteomics and transcriptomics analysis is in, Supplementary File 2 Table 1, and the most significantly highlighted pathways of EL fraction and TL from Reactome analysis are shown in Supplementary File 2 Table 2 and 3), and Ribosome biogenesis as significantly down regulated (Supplementary Figure 14). AMPK (upregulated): These include the following 8 genes SLC2A4, PFKFB2, PRKAA2, STRADB, CAB39L, PRKAG3, PPP2R3A and CCND1. Ribosome biogenesis (downregulated): These include the following 8 genes TRMT112, TBL3, GNL3, GAR1, FBL, NOP56, POP1 and NOB1 (Supplementary File 1 Table 1).

Reactome allows us to overlay our quantitative expression data to visualise the extent of change and progression in affected pathways. In the mRNA analysis we found 231 out of 380 identifiers and 885 pathways were identified by at least one identifier (Supplementary File 6). Supplementary File 2 Table 4 shows the 25 most relevant pathways sorted according to the p-value.

In summary, the analysis highlights rRNA processing (down regulations), major pathway of rRNA processing in the nucleolus and cytosol (down regulations), unfolded protein response (UPR, down regulations), metabolism of RNA (down regulations), and interferon signalling (up regulations). AMPK, as well as being the master regulator of energy homeostasis, is also a physiological suppressor of UPR (Leclerc *et al*., 2013).

There is growing evidence linking AF to metabolic stress and inflammation (Gutierrez & Van Wagoner, 2015; Su *et al*., 2022). In the cell, ribosomes control translation of proteins and their activity accounts for most of the cells energy consumption (Turi *et al*., 2019). Ribosome biogenesis depends on the nutritional and energy status of the cells and is vulnerable to internal and external stress stimuli. Impaired ribosome biogenesis (e.g. in aged tissue) may be protective or a compensatory mechanism (Jung *et al*., 2015). AMPK is a central regulator of energy homeostasis (Kim *et al*., 2016) playing a key role in monitoring cellular energy metabolism and there is growing evidence around the importance of AMPK in the heart (Su *et al*., 2022); (Dyck & Lopaschuk, 2006) (Arad *et al*., 2007). Recently Cao et al (Cao *et al*., 2017) showed that γ2-AMPK translocate into the nucleus to suppress pre-rRNA transcription and ribosome biosynthesis during stress, this reduces ER stress and cell death. We identified PRKAG3 (5’-AMP-activated protein kinase subunit gamma-3) among our upregulated genes (upregulated by log2 2.31fold, Supplementary File 1, Table 1). This seems particularly interesting in light of the findings by Cao et al (2017) where activation of γ2-AMPK suppresses ribosome biogenesis and protects against myocardial ischemia/reperfusion injury. Our transcriptomics data suggests an adaptation process to the stress conditions created during AF. AMPK signalling is increased, and this consequently would be expected to affect ribosomal RNA transcription, as reflected in the genes that are upregulated (Supplementary File 1 Table 1) Su et al (2022) highlight a critical role played by AMPK signalling and the resultant alterations in electrophysiological function and structural remodelling in the atria. Additionally, Su et al (2022) highlight loss of AMPK affecting the expression of mRNA transcripts encoding gap junction proteins and ion channels in the atrium. Here, our goat model aligns with the notion of an upregulation of AMPK to downregulate energy-consuming processes like ribosome biogenesis.

### Glycogen Accumulation and Lysosomal GAA upregulation

Histological studies in healthy/normal goat by Embi *et al*. (2014) showed left atrial appendage glycogen levels always exceeded right atrial appendage levels. The density and location of glycogen was also distinct and suggested these differences in glycogen were a potential contributory mechanism for the initiation and maintenance of AF, particularly the greater propensity for developing an AF substrate in the left versus the right atrium (Embi *et al*., 2014). Studies by Zhang et al in pacing-induced AF (dog model), showed AF promoted glycogen deposition (Zhang *et al*., 2015). In our proteomic data, significantly regulated proteins in the LA of AF goat included GLUT4/ SLC2A4, a protein that transports glucose inside the cell (upregulated by log2 1.55) (Supplementary File 1 Table 2 and Supplementary File 2 Table 4). The GLUT4 vesicle translocation to the plasma membrane (Supplementary Figure 7 and 12) is conducted by the tethering and docking proteins Caveolin (CAV1) and Vesicle-fusing ATPase N-ethylmaleimide-sensitive fusion protein (NSF) (Kanzaki & Pessin, 2003). We observe a significant upregulation of NSF by log2 1.55 (Figure 1B and Supplementary File 1 Table 2) and CAV1 by a fold-change of log2 0.57 (Supplementary Figure 12 and Supplementary File 1 Table 2). Moreover, lysosomal-α-glucosidase (GAA), the glycogen degrading enzyme in the lysosomes (Zirin *et al*., 2013); (Adeva-Andany *et al*., 2016), was upregulated by a log2 fold-change of 0.44 in the EL fraction of the AF goat model (Figure 3 and Supplementary Table 1). We chose to quantitatively asses the levels of Glycogenin 1 (GYG1) because in eukaryotes, Glycogenin 1 enzyme initiates glycogen biogenesis by producing an oligosaccharide primer that functions as a substrate for glycogen synthesis in bulk (Chaikuad *et al*., 2011). Western blotting was performed on the enzyme GYG1 and a significant (*p*=0.007) upregulation of GYG1 was observed in the AF samples (Supplementary Figure 4B and 4C), providing evidence for increased glycogen synthesis in the AF goat model (Embi *et al*., 2014). Here we present specific protein changes that impact on glycogen levels in the cells (Supplementary Figure 4B and 4C). Reassuringly, the evidence presented from our whole tissue omics analysis (Figure 4B, Figure 5B and Supplementary Figure 7) is in keeping with existing structural data (canine (Zhang *et al*., 2015) and goat AF model (Ausma *et al*., 1997)) linking glycogen accumulation and fibrosis as factors in the persistent forms of AF (Ausma *et al*., 1997).

### Increased autophagic flux in AF goat model

Western blotting was performed on sham and AF groups to identify effects of AMPK upregulation in autophagy (Supplementary Figure 4A). For the upregulation of autophagy flux LC3I, and for impaired autophagy flux, LC3II protein markers were blotted. A depletion of LC3I was detected by the absence of LC3I protein in AF compared to the sham goat model, suggesting the upregulation of autophagy flux. Moreover, the LC3II protein was absent in both sham and AF goat models suggesting the absence of impaired autophagic flux instead pointing towards the presence of an overactive flux. This result aligns with a recent study by (Yuan *et al*., 2018). Network analysis shown in Supplementary Figure 7, which presents a close network interaction of AMPK with lysosomal and vesicle localized proteins Ras-related GTP-binding protein A (RRAGA), 1,4-alpha-glucan-branching enzyme (GBE1), Glycogen debranching enzyme (AGL), Phosphoglucomutase-1 (PGM1), and Glycogen phosphorylase, brain form (PYGB).

Our integrated omics analysis comparing EL proteomics and transcriptomics highlights important changes occurring in EL proteins involved in this chronic goat AF model. We see a downregulation in suppression of autophagy, MAPK1 activation, inhibition of nitric oxide (NO) production, mitochondrial electron transport chain deregulation, changes in inflammation status (Interleukins), glycogen disease-like pathologies, changes in gap junction activity, stimulation of cell death response by PAK-2p34, upregulation of RhoBTB proteins that are involved in vesicle trafficking processes and retrograde transport from endosomes to the Golgi apparatus (Figure 4B integrated omics). Our observations are relevant and fit with many observations published over the years including the findings that human atrial samples from patients with AF have increased immune cell infiltration compared to those from patients without AF (Bonilla *et al*., 2012). Additionally, NO produced by endothelial NO synthase (eNOS) plays a role in the regulation of cell growth, apoptosis, and tissue perfusion (Dimmeler *et al*., 1997) and our findings of apoptosis and downregulation of NOS corroborates early findings (Feng *et al*., 2002). ER stress and oxidative stress have been highly implicated in many cardiac pathologies (Binder *et al*., 2022) (Dhalla *et al*., 2000) including the pathogenesis of AF (Li *et al*., 2010). Recent studies highlight the importance of PAK2. In Pak2 cardiac deleted mice under stress or overload, there is a defective ER response, cardiac dysfunction, and profound cell death (Binder *et al*., 2019).

Out of the 2104 proteins, 340 proteins in TL and 148 in EL were significantly changed in AF. We validated Rab11 (Ras-related protein 11) using Western blotting. The LC3I protein was absent in AF, suggesting an increased autophagic flux. The TL fraction, highlighted mitochondrial oxidative-phosphorylation (OXPHOS) and AMPK pathway protein upregulation, indicating a potential increased ATP energy demand in AF. The EL proteins GAA, Rab7a, CLTB, VPS25 and CCT2 were significantly upregulated. The upregulation of protein processing suggests increased vesicular trafficking, potentially related to increased metabolic energy demands. Here we use an endolysosomal purification protocol to study atrial specific protein changes in AF and noted changes in EL proteins. We believe this approach allows us to study differences in protein expression more specifically related to endolysosomes and globally in health and disease.

## Conclusion

Studies have shown that prominent differences between paroxysmal AF and sinus rhythm patients relate to changes in expression of proteins involved in metabolic processes (Chiang *et al*., 2015; Ghezelbash *et al*., 2015). Ozcan et al. provided evidence into the role of atrial metabolism for AF substrate evolution, findings that are relevant when considering alternative therapeutic approaches to prevent AF progression (Ozcan *et al*., 2015). Our EL organelle omics approach helps us describe a disease setting where there is increased cellular stress, increased vesicle trafficking, changes in ATP demands, accumulation of glycogen, inflammation and stimulation of cell death. Our findings uncover new insights linking endolysosomal proteins, ER stress response, RNA biogenesis, and cell apoptosis pathways which may be triggered by failure of protective ER stress response. The current pharmacological therapies for AF are not sufficiently effective to control disease progression and new molecular insights can fuel the development of novel therapeutic strategies. Endolysosomes in the heart provide a significant contribution to basal calcium transient amplitude and beta-adrenergic responses in both atrial (Collins *et al*., 2011) and ventricular (Capel *et al*., 2015) myocytes. Clearly, the multi-functional role of the endolysosomes appears to be at play in this disease setting and these results highlight the need for further investigation into the role of endolysosomal pathways in cellular dysfunction and apoptosis in AF. In summary, our endolysosomal proteomics and integrated omics analysis pave the way for future studies focused on identifying suitable EL targets, drug discoveries and biomarker identification. The novel information provided present a promising option for exploring new pathways for the treatment of AF.

## Limitations

The molecular pathways applied on the protein regulations found in this project are interpretations of published literature and databases and provide a fundamental understanding of the disease pathways. Therefore, an establishment of the protein regulation as a complete pathway alteration needs further and extensive investigation. We have used a large animal goat AF model in which chronic AF was maintained by pacing for 6 months. The data we present is a snapshot of genes and proteins relevant at that point in time of the disease. The multifactorial and heterogeneous nature of AF as a disease (Xi & Cheng, 2015; Nattel & Dobrev, 2016; Schüttler *et al*., 2020) poses limitations in the assessment of the occurrence and changes in gene and protein expression especially during the process of disease progression from paroxysmal to chronic AF.

## Competing Interests

None to declare

## Author contributions

RABB conceived and designed the study; TA, GB, RF and HK performed LC-MS/MS; TA and BB performed western blots; TA and DP performed enzyme assays; RABB, TA, RAC, SJB and FP contributed intellectually to the project and RABB, TA, and SJB drafted the paper. TA performed statistical analysis of proteomics and pathways analysis. APC and TA performed statistical analysis, on transcriptomics. PDC performed integrated analysis of proteomics versus transcriptomics. SJB performed the species comparison study. RAC, DA, QS and DM analysed EM data and created EM figures. TA created the figures related to proteomics. TA and APC created figures related to transcriptomics. PDC and TA created integrated proteomics and transcriptomics analysis figures. GM created code for analysing

EM images for glycogen and QS manually analysed EM images. BB performed Western blotting studies on LC3 and p-p70. All authors contributed to writing of the paper.

## STAR Methods

## EXPERIMENTAL MODEL AND SUBJECT DETAILS

### Animals

AF was induced and maintained in female goats (C. hircus) for 6 months (AF goat model was created as conducted in (van Hunnik *et al*., 2018), followed by an open chest sacrifice experiment (N=4 AF and N=4 sham controls) (The goat study was carried out in accordance with the principles of the Basel declaration and regulations of European directive 2010/63/EU, and the local ethical board for animal experimentation of the Maastricht University approved the protocol).

## METHOD DETAILS

### Tissue homogenization

Frozen left atrial tissue biopsies of AF and sham goat were thoroughly cleaned using Phosphate buffered solution (PBS) and weighed. A minimum of 100 mg tissue is weighted in order to perform proteomics. Each atrium biopsy sample was cut using sterile scalpels and gently homogenized using a 7 mL Dounce homogenizer in Lysosome isolation buffer (LIB) [Containing 1:500 protease inhibitor cocktail (PIC) and phosphatase inhibitor (PHI) (Bio vision), (PhosSTOP Roche)]. Preparations were further homogenized in 1 mL Dounce homogeniser and transferred to chilled 1.5 mL ultracentrifugation tubes (Beckmann coulter). Sample preparations were mixed at a ratio of 1:1.5 Lysosome enrichment buffer [(LEB) (Biovision, containing 1:500 PIC)] to homogenate by inverting tubes, and were stored on ice for 5 min until the centrifugation (Ayagama *et al*., 2021).

### Tissue Lysate (TL)

Samples were centrifuged at 13,000 g x 2 min at 4 °C (TLX Beckmann Coulter Ultra Centrifuge) and the supernatant or the TL, was collected (Ayagama et al., 2021).

### Endo-lysosome Fraction (EL)

The collected supernatant was retained and repeated for a further centrifugation step at 29,000 g x 30 min at 15 °C (500 µL of under-laid 2.5 M sucrose with over-laid 500 µL Percoll). The supernatant above the sucrose and Percoll intermediate was collected for further fractionation. Firstly, ultracentrifuge tubes were underlaid with 2.5 M sucrose and overlaid with a series of Percoll dilutions (1.11 g/mL – 1.04 g/mL in ddH2O). The ultracentrifuge tubes were centrifuged at 67,000 g x 30min at 4 °C. The fraction at 1.04 g/mL was collected and labelled as the endolysosomal fraction (EL) (Ayagama et al., 2021). N = 3 biological replicates for each AF and sham conditions were used in proteomic analysis.

### Liquid chromatography-tandem mass spectrometry analysis

Samples were reduced using 5 µL of 200 mM dithiothreitol (30 min at room temperature) and alkylated with 20 µL of 200 mM iodoacetamide (30 min at room temperature), followed by methanol-chloroform precipitation (supplementary data). The Pelleted protein was re-suspended in 6 M urea in 400 mM Tris-HCl, pH 7.8. Urea was then diluted to 1 M with 400 mM Tris-HCl at pH 7.8, and the proteins were digested in trypsin at a ratio of 1:50 (overnight at 37C°). Once the trypsin digestion is completed, the samples are processed at the Target Discovery Institute, Oxford.

Samples were then acidified to a final concentration of 1% formic acid, and the samples were desalted on Sola HRP SPE cartridges (Thermo Fisher Scientific) and dried down using a SpeedVaccum centrifuge. Dried down protein Samples were further desalted online (PepMAP C18, 300 µm x 5 mm, 5 µm particle, Thermo Fisher Scientific) for 1 min (flow rate of 20 µL/min and separated on an EASY-Spray column) (PepMAP C18, 75 µm x 500 mm, 2 µm particle, ES803, Thermo Fisher Scientific) over 60 min using a gradient of 2–35% acetonitrile in 5% DMSO/ 0.1% formic acid at 250 nL/min. Separation and analysis were performed on a Dionex Ultimate 3000 RSLC system coupled to an Orbitrap Fusion Lumos platform (both Thermo Fisher Scientific) using standard parameters (Universal Method (Davis et al., 2017).

MS scans were acquired at a resolution of 120,000 between 400 and 1,500 m/z. An AGC target of 4.0E5 and MS/MS spectra detection was carried out using rapid scan mode in the linear ion trap at a 35% collision energy after collision-induced dissociation fragmentation (CIDF). An AGC target of 4.0E3 for up to 250 ms, employing a maximal duty cycle of 3 s, prioritising the most intense ions and injecting ions for all available parallelisable time. Selected precursor masses were excluded for 30 s (Davis et al., 2017).

Mass spectrometry data were analysed quantitatively with the Progenesis QI software platform (WatersTM Cooperation, www.nonlinear.com) (version 4.2), and database searches were carried out against the UniProt C. hircus database (UP000291000). Automatic processing was selected. All runs in the experiment were adjusted to the function suitability, and runs were aligned automatically. The peak picking was selected between 10 and 75 min. The group runs option was set to conditions, and relative quantitation using Hi-N was selected. Finally, proteins were grouped.

### Quantification and statistical analysis of mass spectrometry data

Quantitative analysis for significant differences of protein regulation between the AF and sham conditions of TL and EL samples and data visualization were performed using the Perseus software platform (Tyanova et al., 2016) (version 1.6.15.0). Using protein intensity values of biological replicates, the protein groups were created and uploaded as a data matrix in Perseus with the respective protein abundances as main columns. The data matrix was reduced by filtering based on categorical columns to remove proteins where more than two intensity values were absent from six biological replicates, and remaining data with no more than 2 missing values were quantitatively analysed. A total of 2,104 proteins in TL and EL remained after filtering. Groups of biological replicates for TL and EL fractions were defined in categorical annotation rows. Data were log transformed (log2) and normalised via Z score.

Missing data points were imputed based on normal distribution and visualized as normalised intensity histograms (per biological replicate) with imputed values. Principal component analysis (PCA) was performed on 100% valid values. A volcano plot was generated based on normalised intensities applying two-way Student’s t-test to probe for significant difference of protein regulation between AF and sham conditions of each TL and EL samples. A permutation-based false-discovery rate (FDR) was determined with 250 randomizations and S0 = 0.1 (default). The quantified proteins were accounted with 99% confidence level at 5% FDR.

### Principal Component Analysis (PCA)

Principal component analysis is a reduced data dimension-interpretation of the protein groups’ distribution between the sample groups and it is one of the most popular multivariate statistical techniques. A PCA plot can provide a window to a large data set by identifying the common vectors, therefore summarizing the variation (Yacine & Dacheng, 2017). As indicated in Figure 1E the respective component 1 and component 2 vector deviations were observed between AF (purple symbols) and sham (green symbols) EL groups demonstrating that the variability is driven by the differential experiment groups rather than the variability within the sample group.

### Violin plot analysis

The violin plots were produced using the InstantClue omics tool (Nolte et al., 2018). The protein intensities used were transformed to log2, normalised by z scoring, and AF vs sham groups were colour-coded; a gradient of purple for the distribution of protein intensities (Figure 2A). This Violin plot represents a kernel density estimation of the underlying protein intensity distribution (The kernel estimated density distribution is the nonparametric representation of the probability density function of a random variable (www.mathworks.com). The quantified protein intensity matrix used for the violin plot is created in Perseus 1.6.15.0 (Tyanova et al., 2016).

### Heat map analysis

The heat or clustering maps were produced in InstantClue omics tool using Euclidian distance and K-mean clustering of the normalised protein intensities. The intensity values are obtained from the quantified matrix created in Perseus 1.6.15.0. A total of 148 proteins in EL were categorized into protein clusters according to the protein intensities (EL = 3 clusters). The regulation level variations are displayed using colour codes (red and blue), where red represents the up regulation of the protein clusters and blue represents the down regulation of these clusters (Figure 2C).

### Statistics for Network analysis

The string network edges of EL and TL were created based on the interaction evidence, which are experiments, gene fusions, data bases, co-occurrences and co-expressions. The minimum required interaction score was set to default or medium confidence (0.4), and only the query proteins were used and external interactors were excluded.

### Sample Preparation for Transcriptomics

Frozen AF and sham LA goat tissue were collected without the RNase contamination, and samples were thawed using an RNAlater stabilisation solution (AM7020, Invitrogen). Samples were then homogenised using bead disruption, and in a 1mL of Trizol (15596026, Thermo Fischer), approximately 50 – 100 mg of tissue were solubilised. The supernatant was collected from lysates after incubating them in RT for 5minutes and centrifuging at 12,000 x g for 5 min at 4°C. Furthermore, it was followed with a re-centrifugation at 12,000 g at 4°C for 15 min, after vigorously shaking with 200 µL chloroform. The upper aqueous phase was separated, 1:1 volume ice-cold isopropanol was added, and gently mixed. This procedure was followed by centrifugation at 12,000 g for 30 min at 4°C, and the collected RNA pellet was washed with 75% ethyl alcohol (10048291, Thermo Fisher) and dried.

### Qualification and Quantification of mRNA

The isolated RNA was screened for contaminants and degradation levels using 1% agarose, and a NanoPhotometer® spectrophotometer (IMPLEN, CA, USA) was used to check the purity (RNA Nano 6000 Assay Kit of the Bio-analyser 2100 system (Agilent Technologies, CA, USA) was used to detect the RNA integrity and assess quantitation.

Sequence libraries were prepared using the NEBNext® Ultra TM RNA Illumina® (NEB, USA) Library Prep Kit. Furthermore, using a PE Cluster Kit cBot-HS all the samples were clustered, and sequenced using an Illumina.

Fastp was used to remove the poly-N, and adapter reads from the raw data and process. The reads Q score over 50% was considered low quality, and Q score at 20 and 30 were identified as clean. Genome web browser National Centre for Biotechnology Information/European Molecular Biology Laboratories-European Bioinformatics Institute (NCBI/Ensembl-EBI) was used as the reference genome with HISAT2 programme (daehwankimlab.github.io/hisat2/manual/).

### Integrated Analysis of Proteomics and transcriptomics statistics

The proteomic data from Progenesis and the transcript data were analysed in R v4.1.2. Both proteomic intensities and transcript counts were normalised by Variance Stabilizing Normalisation (VSN)(Huber *et al*., 2002) (Supplementary figure 8 and 9). Due to the poor annotation status of Capra hircus gene, protein and pathway IDs, the data were ’humanised’; human protein-coding homolog gene IDs were assigned to matchable Goat IDs using the getLDS function in the biomaRt package (Durinck *et al*., 2009). In total, 27552 IDs across the proteomic and transcriptomic datasets were mapped to 18658 human IDs. Further analysis was performed on the human IDs.

### Removal of sample Outliers

The outliers of AF and sham samples were identified by conducting principal component analysis on the individual and shared proteomics and transcriptomics data, then the outliers were removed to increase the confidence of the study (Please refer to Supplementary file 3, a PCA plot showing the outliers of combined proteomics and transcriptomics data is in the Supplementary File 3 Figure 1).

For each AF vs Control comparison, t-statistics were calculated for the AF versus control comparison for each gene. For all human Reactome pathways containing genes found in the dataset, we calculated rank-based 2D enrichment MANOVA p-values after (Cox & Mann, 2012) for all pathways with at least 2 genes found on both sides of each paired comparison, over the three proteomics datasets and the transcriptomics dataset. Missing values for genes that had non-missing value at least one in any proteomics dataset were treated as ranked last; conversely, genes with no non-missing values except in the transcriptomics dataset were treated as missing (and excluded from ranking) in the proteomics datasets. To correct for multiple testing, pathway p-values were converted to q-values using the fdr tool package (Strimmer, 2008). We set a threshold of 1% FDR for significance.

### Protein quantitation

Sample fractions EL or TL were mixed at a ratio of 1:1 with radio-immunoprecipitation (RIPA) buffer (Thermo Scientific). Protein concentrations of EL fractions and TL were determined using the Bicinchoninic acid assay (BCA Protein Assay Kit, Thermo Scientific). Bovine serum albumin was used as a protein standard, and serial dilutions were prepared from the initial stock concentration of 2 mg/mL to prepare a standard curve. To ensure accuracy and reproducibility, protein assays were performed in triplicate. Absorbance values were measured at 562 nm. Protein concentrations were calculated by linear regression analysis (Ayagama *et al*., 2021).

### SDS/PAGE gel preparation and Western blotting

Sample fractions EL and TL were solubilised, and proteins denatured using SDS/PAGE loading buffer (bio rad) and 2-mercaptoethanol (Sigma-Aldrich). Proteins were separated by gel electrophoresis (NW04120BOX, NuPAGE 4%–12% Bis-Tris protein gels, 20X MES buffer). The gel was transferred to nitrocellulose membrane (NC) (Bio-Rad) for protein transfer (X-cell-II blot module, Thermo Fisher Scientific). NC membrane was incubated in 5% skimmed milk. The primary antibodies anti-GYG1 (1:1000, sc-271109, Santa Cruz), anti-Rab11 (1:1000, ab65200, abcam) were incubated. Goat anti-rabbit antibody (1:2,500, Dako P0448) was used as the secondary antibody to detect the protein markers. The secondary antibodies were detected via chemiluminescence using Westar Supernova (XLS3, 0020, Cyanogen) and the protein bands were visualized in a ChemiDoc XRS + imager (Bio-rad with image Lab software) (Ayagama *et al*., 2021).

### Lysosomal hydrolase activity assays

To fluorometrically measure the lysosome enzyme levels, artificial sugar substrates containing the fluorophore 4-methylumbelliferone (4-MU) were used. For measuring β-hexosaminidase activity, 3 mM 4-MU N-acetyl-β-D-glucosaminide (Sigma Aldrich) in 200 mM sodium citrate buffer, pH 4.5 and 0.1% Triton X-100 was used as substrate. For β-galactosidase activity, 1 mM 4-MU β-D-galactopyranoside (Sigma Aldrich) in 200 mM sodium acetate buffer, pH 4.3, 100 mM NaCl, and 0.1% Triton X-100 was used as substrate. The reaction was stopped by adding chilled 0.5 M Na2CO3, and the released fluorescent 4-MU was measured in a Clariostar OPTIMA plate reader (BMG Labtech, Ortenberg, Germany) with an excitation at 360 nm and emission at 460 nm. A standard curve for free 4-MU was used to calculate the enzyme activity. Results were calculated as total Units of enzyme activity (nmol/hr) and normalised with respect to protein content (Ayagama *et al*., 2021).

### Sample Preparation for Electron Microscopy (EM)

Left atrial samples (from n = 3 sham animals and 3 AF animals) were prepared by chemical fixation. Approximately 1 mm^3^ pieces of tissue from the left atria were rapidly dissected and fixed in Karnovsky fixative (paraformaldehyde 4%, glutaraldehyde 5%, cacodylate buffer 80 mM, pH 7.4, (Karnovsky & Karnovsky, 1965) and were embedded in Spurr’s resin (Spurr, 1969) . Sections were cut to approximately 70-80 nm thickness (Reichart Ultracut) then post-stained with 2% aqueous uranyl acetate and Reynolds lead citrate for contrast. Images were obtained using a transmission electron microscope (Thermo Fisher Tecnai T12 TEM, operated at 120kV, using a Gatan OneView camera). Analysis and measurements were made using Gatan DigitalMicrograph and ImageJ software.

Raw EM images were analysed using ImageJ (Version 2.3.0/1.53s). The area of interest was manually selected using the box tool, avoiding cellular organelles such as Golgi, lysosome, mitochondria etc. 5-7 boxes per image were selected in order to cover most of the cytosolic space. Box sizes in pixels were generated by ImageJ. Glycogen quantity was evaluated manually using the multi-point tool, one point represents one glycogen. The overall number of points was recorded as an output by ImageJ. The results are presented as glycogen counts per μm^2.

## Acknowledgments

RABB is funded by a Sir Henry Dale Wellcome Trust and Royal Society Fellowship (109371/Z/15/Z) and TA acknowledges support from The Returning Carers’ Fund, Medical Sciences Division, and Global Challenges Research Fund, University of Oxford. RABB is a Senior Research Fellow of at Linacre College. SJB is funded by the British Heart Foundation. RAC is funded by the Wellcome Trust and Royal Society. FMP is a Wellcome Trust Investigator in Science and a Royal Society Wolfson merit award holder. DAP was funded by the Mizutani Foundation. AG is a Wellcome Trust Senior Investigator and a Principal Investigator of the British Heart Foundation Centre of Research Excellence at the University of Oxford. NH is a British Heart Foundation Senior Clinical Research Fellow (FS/SCRF/20/32005). We would like to thank the Parrington and Tammaro Groups, Department of Pharmacology, University of Oxford for access to scientific equipment. We thank Dr Errin Johnson and Raman Dhaliwal from the Dunn School of Pathology for Electron Microscopy technical support. The goat model work was supported by the Netherlands Heart Foundation (CVON2014-09, RACE V Reappraisal of Atrial Fibrillation: Interaction between hypercoagulability, Electrical remodeling, and Vascular Destabilisation in the Progression of AF, and Grant number 01-002-2022-0118, EmbRACE: Electro-Molecular Basis and the theRapeutic management of Atrial Cardiomyopathy, fibrillation and associated outcomEs), the European Union (CATCH ME: Characterizing Atrial fibrillation by Translating its Causes into Health Modifiers in the Elderly, grant number 633196; MAESTRIA: Machine Learning Artificial Intelligence Early Detection Stroke Atrial Fibrillation, grant number 965286).

---

**Supplementary Table 1:** The most significantly regulated EL fraction proteins of the AF goat model

| Protein ID | Protein name | Gene | Fold change |
| --- | --- | --- | --- |
| A0A452F9W2_CAPHI | Prefoldin subunit 4 | PFDN4 | 2.12 |
| A0A452DN27_CAPHI | Single-pass membrane and coiled-coil domain-containing protein 4 | SMCO4 | 2.01 |
| A0A452DY79_CAPHI | Structural maintenance of chromosomes protein | SMC1A | 2.01 |
| A0A452DL17_CAPHI | SAP domain-containing protein | CCAR1 | 1.99 |
| A0A452G2Z7_CAPHI | E3 ubiquitin-protein ligase HECTD3 | HECTD3 | 1.83 |
| A0A452DPX9_CAPHI | WD_REPEATS_REGION domain-containing protein | WDR78 | 1.79 |
| A0A452EJX2_CAPHI | Reticulon | RTN3 | 1.78 |
| A0A452G662_CAPHI | SHSP domain-containing protein | HSPB7 | 1.77 |
| A0A452G8P6_CAPHI | RING finger protein 17 | RNF17 | 1.75 |
| A0A452DM61_CAPHI | Calponin-homology (CH) domain-containing protein | NAV2 | 1.72 |
| A0A452F797_CAPHI | E3 ubiquitin-protein ligase PPP1R11 | PPP1R11 | 1.71 |
| A0A452FPZ5_CAPHI | cAMP-dependent protein kinase inhibitor | PKIG | 1.68 |
| A0A452G133_CAPHI | 1,4-alpha-glucan branching enzyme | GBE1 | 1.67 |
| A0A452F8B4_CAPHI | Cytochrome c oxidase subunit | COX6A2 | 1.56 |
| A0A452G4I2_CAPHI | SHSP domain-containing protein | HSPB2 | 1.55 |
| A0A452F5D8_CAPHI | 2-phospho-D-glycerate hydro-lyase | ENO2 | 1.51 |
| A0A452FLK7_CAPHI | ATP synthase-coupling factor 6, mitochondrial | ATP5PF | 1.47 |
| A0A452DQP7_CAPHI | Actin, alpha cardiac muscle 1 | ACTC1 | 1.47 |
| A0A452EBL3_CAPHI | DNA-directed RNA polymerase III subunit RPC4 | POLR3D | 1.46 |
| A0A452F3N4_CAPHI | Clathrin light chain | CLTB | 1.45 |
| A0A452FRY3_CAPHI | Splicing factor, proline- and glutamine-rich | SFPQ | 1.43 |
| A0A452DRC3_CAPHI | Mitochondrial import inner membrane translocase subunit TIM16 | PAM16 | 1.39 |
| A0A452ERZ9_CAPHI | Complex I-MNLL | NDUFB1 | 1.37 |
| A0A452G711_CAPHI | D-beta-hydroxybutyrate dehydrogenase, mitochondrial | BDH1 | 1.32 |
| A0A452FQ01_CAPHI | Protein cordon-bleu | COBL | 1.25 |
| A0A452E6Y1_CAPHI | Prefoldin subunit 1 | PFDN1 | 1.22 |
| A0A452FJJ3_CAPHI | PRA1 family protein | ARL6IP5 | 1.20 |
| A0A452DY71_CAPHI | Microtubule-actin cross-linking factor 1, isoforms 1/2/3/5 | MACF1 | 1.19 |
| A0A452FIK7_CAPHI | Lipoma-preferred partner | LPP | 1.18 |
| A0A452F973_CAPHI | Cathelicidin-2 | CATHL2 | 1.16 |
| A0A452EW34_CAPHI | Myosin phosphatase Rho-interacting protein | MPRIP | 1.14 |
| A0A452EQJ4_CAPHI | PDZ domain-containing protein | AHNAK | 1.14 |
| A0A452FP52_CAPHI | Coxsackievirus and adenovirus receptor | CXADR | 1.11 |
| A0A452F158_CAPHI | Protein phosphatase 1 regulatory subunit | PPP1R12C | 1.10 |
| A0A452FDC1_CAPHI | NADH dehydrogenase [ubiquinone] iron-sulfur protein 6, mitochondrial | NDUFS6 | 1.08 |
| A0A452FQ33_CAPHI | Prelamin-A/C | LMNA | 1.07 |
| A0A452G3Z1_CAPHI | Diacylglycerol kinase | DGKD | 1.06 |
| A0A452EIY1_CAPHI | Microtubule-associated protein | MAP4 | 1.05 |
| A0A452FHF8_CAPHI | Prefoldin subunit 6 | PFDN6 | 1.04 |
| A0A452EW73_CAPHI | Autism susceptibility gene 2 protein | AUTS2 | 1.04 |
| A0A452EKC1_CAPHI | NADH dehydrogenase [ubiquinone] 1 beta subcomplex subunit 8, mitochondrial | NDUFB8 | 1.02 |
| A0A452FL98_CAPHI | Ubiquitinyl hydrolase 1 | USP9X | 1.00 |
| A0A452DMC5_CAPHI | Hsp90 chaperone protein kinase-targeting subunit | CDC37 | 1.00 |
| A0A452FYK7_CAPHI | BolA-like protein 2 | BOLA2B | 1.00 |
| A0A452E9A8_CAPHI | TRASH domain-containing protein | RPL24 | 1.00 |
| A0A452FDZ6_CAPHI | Ubiquitin-fold modifier-conjugating enzyme 1 | UFC1 | 1.00 |
| A0A452FSE0_CAPHI | Protein-cysteine N-palmitoyltransferase HHAT-like protein | HHATL | 1.00 |
| A0A452F2G4_CAPHI | Dynein light chain roadblock | DYNLRB1 | 0.98 |
| A0A452G757_CAPHI | IF rod domain-containing protein | NES | 0.96 |
| A0A452G5E3_CAPHI | ATP synthase subunit d, mitochondrial | ATP5PD | 0.96 |
| A0A452EGT0_CAPHI | Complex I-B18 | NDUFB7 | 0.95 |
| A0A452FP74_CAPHI | Gal_mutarotas_2 domain-containing protein | GANAB | 0.94 |
| A0A452ENN3_CAPHI | Leucyl-tRNA synthetase | LARS1 | 0.93 |
| A0A452DKF1_CAPHI | Vesicle-associated membrane protein-associated protein A | VAPA | 0.93 |
| A0A452F4D3_CAPHI | Cystatin domain-containing protein | LOC102186806 | 0.92 |
| A0A452EPW1_CAPHI | NAD-dependent protein deacetylase | SIRT3 | 0.91 |
| A0A452G0B4_CAPHI | Cytochrome b5 heme-binding domain-containing protein | NENF | 0.91 |
| A0A452F760_CAPHI | N-lysine methyltransferase SMYD2 | SMYD2 | 0.90 |
| A0A452DZF9_CAPHI | Matrix-remodeling-associated protein 7 | MXRA7 | 0.87 |
| A0A452FYY0_CAPHI | Short/branched chain specific acyl-CoA dehydrogenase | ACADSB | 0.87 |
| A5JSS3_CAPHI | NADH dehydrogenase (Ubiquinone) 1 alpha subcomplex 4 | NDUFA4 | 0.86 |
| A0A452EKQ0_CAPHI | GRASP55_65 domain-containing protein | GORASP2 | 0.86 |
| A0A452FF94_CAPHI | Glutaredoxin domain-containing protein | GLRX | 0.85 |
| A0A452FEX6_CAPHI | Complex I-B22 | NDUFB9 | 0.85 |
| A0A452FJK2_CAPHI | Fibronectin | FN1 | 0.84 |
| PLMN_CAPHI | Plasminogen (Fragment) | PLG | 0.81 |
| A0A452FQM9_CAPHI | Complex I-B12 | NDUFB3 | 0.81 |
| A0A452G1G3_CAPHI | Helix-destabilizing protein | HNRNPA1 | 0.81 |
| A0A452F1F7_CAPHI | 60S acidic ribosomal protein P2 | RPLP2 | 0.77 |
| A0A452EUX2_CAPHI | Peroxiredoxin-6 | PRDX6 | 0.76 |
| A0A452E9Y7_CAPHI | Myomesin-2 | MYOM2 | 0.74 |
| A0A452GB99_CAPHI | Transgelin | TAGLN2 | 0.71 |
| A0A452F2I2_CAPHI | Epoxide hydrolase | EPHX1 | 0.68 |
| A0A452FAW7_CAPHI | Sodium/potassium-transporting ATPase subunit alpha | ATP4A | 0.67 |
| A0A452ESU7_CAPHI | Heat shock 27 kDa protein | HSPB1 | 0.63 |
| A0A452F8A8_CAPHI | Cerebral dopamine neurotrophic factor | CDNF | 0.60 |
| A0A452EI29_CAPHI | NADH dehydrogenase [ubiquinone] 1 alpha subcomplex subunit 6 | NDUFA6 | 0.60 |
| A0A452FI28_CAPHI | Tubulin-specific chaperone A | TBCA | 0.58 |
| A0A452FFI1_CAPHI | EH domain-containing protein 1 | EHD1 | 0.57 |
| A0A452G4D6_CAPHI | CCT-beta | CCT2 | 0.55 |
| A0A452EF68_CAPHI | PCI domain-containing protein | COPS7A | 0.48 |
| A0A452DXZ3_CAPHI | Metaxin-2 | MTX2 | 0.46 |
| A0A452EV92_CAPHI | P-type domain-containing protein | GAA | 0.44 |
| A0A452ETD8_CAPHI | Filamin-B | FLNB | 0.41 |
| A0A452ERC8_CAPHI | Ribonuclease inhibitor | RNH1 | -0.41 |
| A0A452EB34_CAPHI | ADP/ATP translocase 3 | SLC25A6 | -0.43 |
| A0A452FQE3_CAPHI | Malic enzyme | ME1 | -0.43 |
| A0A452DMR1_CAPHI | Ras-related protein Rab-7a | RAB7A | -0.50 |
| A0A452F3X0_CAPHI | Myosin-7B | MYH7B | -0.57 |
| A0A452DSU6_CAPHI | Mediator of ErbB2-driven cell motility 1 | MEMO1 | -0.61 |
| A0A452EI23_CAPHI | Lactamase_B domain-containing protein | ETHE1 | -0.62 |
| A0A452E1G5_CAPHI | Nuclear receptor corepressor 2 | NCOR2 | -0.62 |
| A0A452DVL8_CAPHI | Poly(rC)-binding protein 1 | PCBP1 | -0.70 |
| A0A452G6A1_CAPHI | Dystonin | DST | -0.70 |
| A0A452F3M1_CAPHI | Importin N-terminal domain-containing protein | IPO5 | -0.70 |
| A0A452EH63_CAPHI | Fructose-bisphosphate aldolase | ALDOA | -0.72 |
| A0A452DZ25_CAPHI | NEDD8-activating enzyme E1 regulatory subunit | NAE1 | -0.76 |
| A0A452FYK1_CAPHI | Exosome complex protein LRP1 | LRP1 | -0.76 |
| A0A452FU89_CAPHI | Pyridoxal phosphate homeostasis protein | PLPBP | -0.78 |
| A0A452DU00_CAPHI | PABS domain-containing protein | SMS | -0.79 |
| A0A452E7Y7_CAPHI | PKS_ER domain-containing protein | RTN4IP1 | -0.79 |
| IL6_CAPHI | Interleukin-6 | IL6 | -0.86 |
| A0A452FN18_CAPHI | IF rod domain-containing protein | KRT3 | -0.86 |
| A0A452FAD3_CAPHI | Glyoxylate reductase/hydroxypyruvate reductase | GRHPR | -0.87 |
| A0A452EQN7_CAPHI | CSD_1 domain-containing protein | CARHSP1 | -0.92 |
| A0A452EKB8_CAPHI | Pyridoxal phosphate phosphatase | PDXP | -0.92 |
| A0A452FAZ1_CAPHI | Proteasome subunit beta | PSMB4 | -0.94 |
| A0A452G1A9_CAPHI | Tubulin beta chain | TUBB6 | -0.95 |
| A0A452EXY9_CAPHI | Serine/arginine-rich splicing factor 2 | SRSF2 | -0.95 |
| A0A452FD14_CAPHI | Glycerol-3-phosphate dehydrogenase [NAD(+)] | GPD1 | -0.97 |
| A0A452FCU2_CAPHI | Farnesyl pyrophosphate synthase | FDPS | -0.98 |
| A0A452E698_CAPHI | Chromobox protein homolog 1 | CBX1 | -0.98 |
| A0A452FV62_CAPHI | 4a-hydroxytetrahydrobiopterin dehydratase | PCBD2 | -0.98 |
| A0A452DRD5_CAPHI | Eukaryotic translation initiation factor 3 subunit E | EIF3E | -0.99 |
| A0A452FVJ7_CAPHI | Phospholipase B-like | PLBD1 | -0.99 |
| A0A452FUU5_CAPHI | Perilipin | PLIN3 | -1.00 |
| A0A452FP83_CAPHI | RRM domain-containing protein | HNRNPC | -1.00 |
| A0A452ES90_CAPHI | ESCRT-II complex subunit VPS25 | VPS25 | -1.04 |
| A0A452FY19_CAPHI | OCIA domain-containing protein | OCIAD1 | -1.04 |
| A0A452EHJ6_CAPHI | Hydroxyacyl-coenzyme A dehydrogenase, mitochondrial | HADH | -1.05 |
| A0A452EJR1_CAPHI | EGF domain-specific O-linked N-acetylglucosamine transferase | EOGT | -1.06 |
| A0A452DQ55_CAPHI | Protein-synthesizing GTPase | EIF2S3 | -1.07 |
| A0A452EIV8_CAPHI | Non-specific serine/threonine protein kinase | MAPKAPK3 | -1.08 |
| A0A452DKT5_CAPHI | Small nuclear ribonucleoprotein Sm D3 | SNRPD3 | -1.21 |
| A0A452FIX8_CAPHI | Early endosome antigen 1 | EEA1 | -1.21 |
| A0A452EVX4_CAPHI | CCT-alpha | TCP1 | -1.23 |
| A0A452G5M5_CAPHI | Palmdelphin | PALMD | -1.23 |
| A0A452FLJ3_CAPHI | Caveolae-associated protein 1 | CAVIN1 | -1.26 |
| A0A452FNA0_CAPHI | Coronin | CORO1C | -1.28 |
| A0A452G7L4_CAPHI | S-adenosylmethionine synthase | MAT2A | -1.31 |
| A0A452FVB0_CAPHI | Actin-related protein 2/3 complex subunit 3 | ARPC3 | -1.32 |
| A0A452FX49_CAPHI | Hemoglobin subunit beta-C | HBBC | -1.38 |
| Q8WF85_CAPHI | Cytochrome b | cytb | -1.46 |
| A0A452DJR4_CAPHI | SWIM-type domain-containing protein | ZSWIM8 | -1.48 |
| A0A452DS40_CAPHI | VWFA domain-containing protein | ITGAX | -1.48 |
| A0A452F341_CAPHI | Beta-MPP | PMPCB | -1.86 |
| A0A452F1Z3_CAPHI | Argininosuccinate synthase | ASS1 | -2.45 |

**Supplementary Figure 1.**
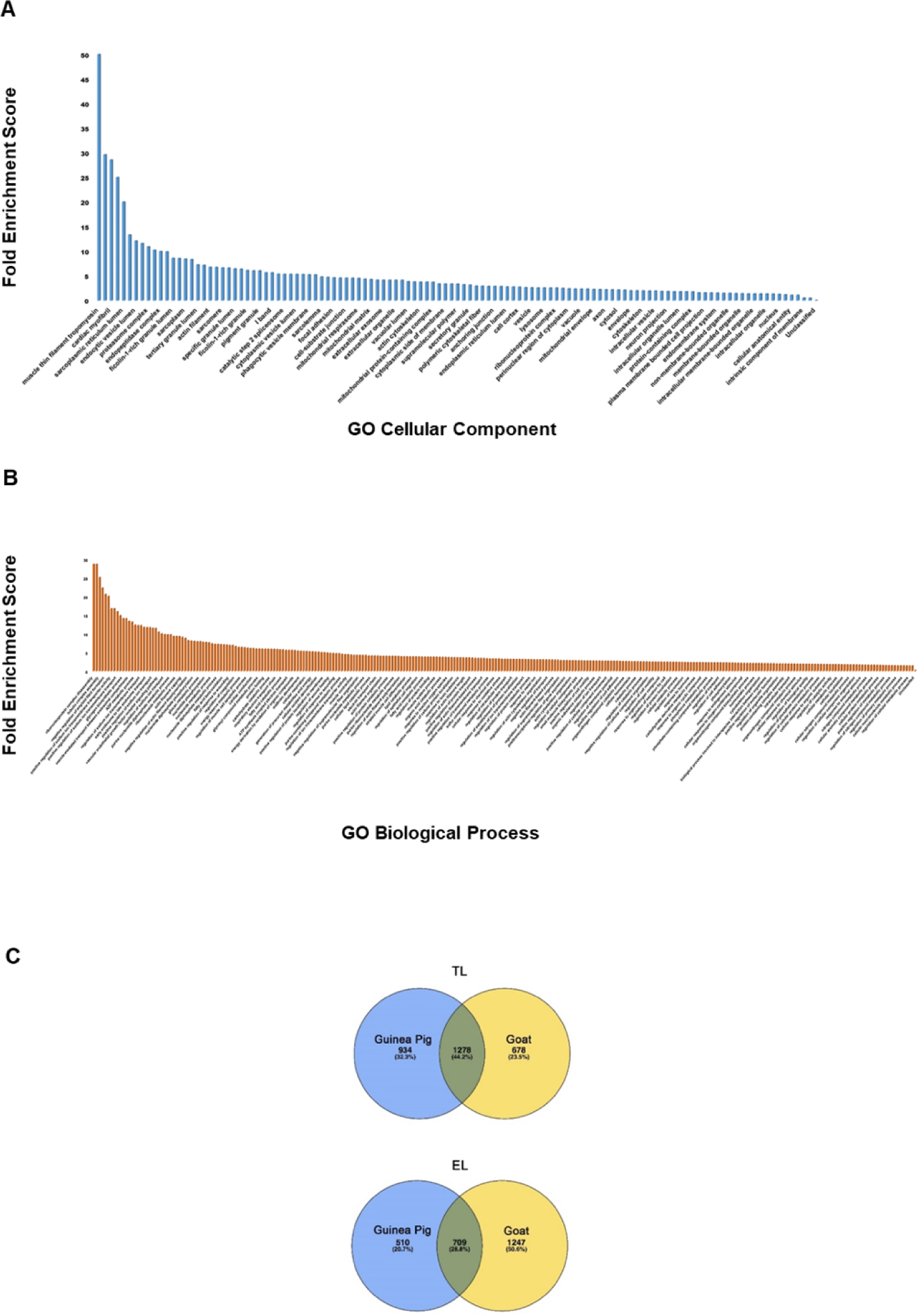
(relating to figure 1 and 3) **A and B** The enrichment of regulated proteins in AF goat model compared to the total human proteome. The over representation test was performed using the converted C. *hircus* protein identifiers to H. *sapiens*. The higher fold enrichment scores were plotted against the Gene Ontology functional and anatomical parameters, Cellular component and Biological process. **C** Species comparison between C. *porcellus* and C. *hircus* TL and EL fractions against the protein yield discovered from Ayagama *et al* 2021 study to understand the protein/gene recovery.

**Supplementary Figure 2.**
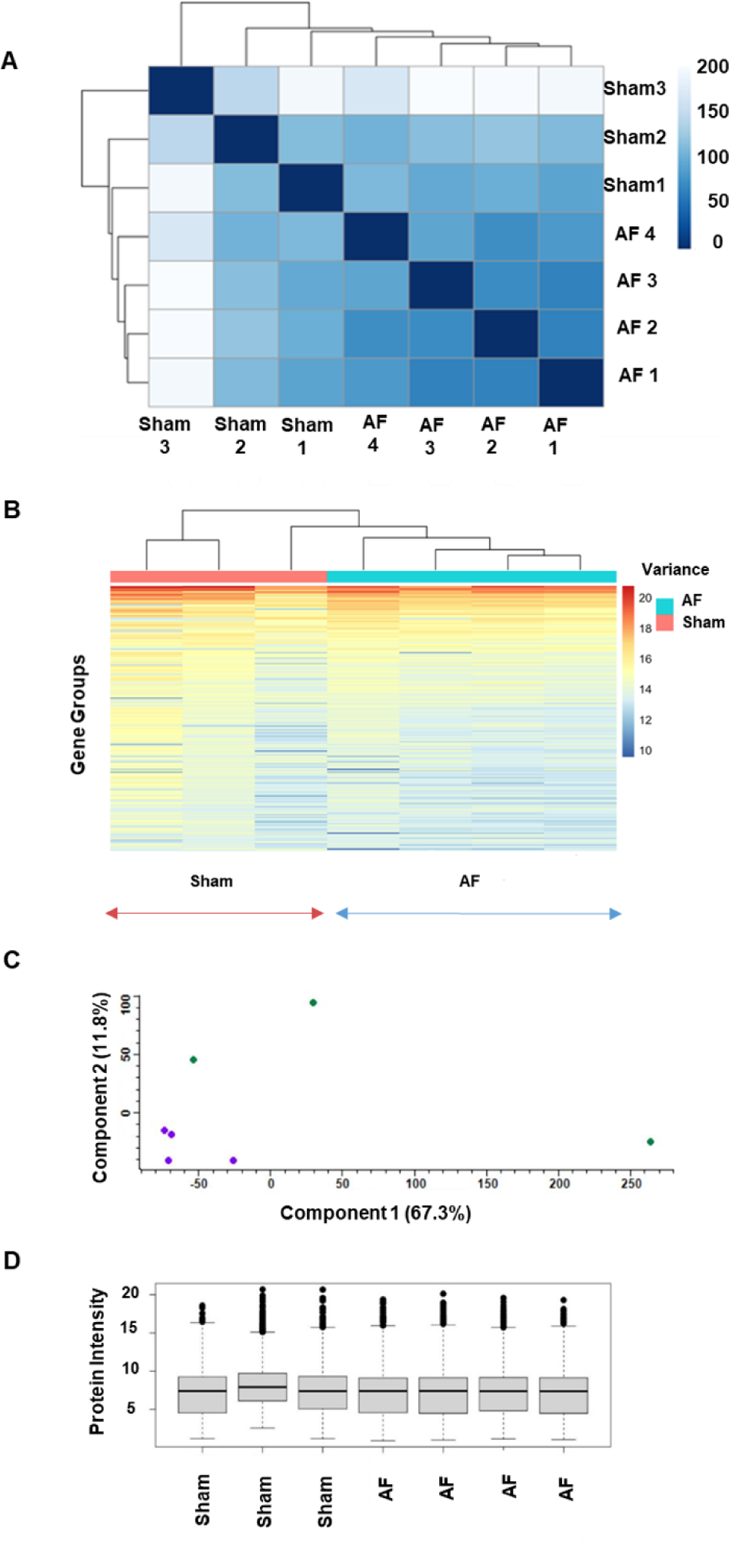
(relating to figure 4A): Quality control analysis for transcriptomics. **A** Distance matrix heatmap to determine the distance between the AF and sham samples according to their gene groups. **B** A heat map of the transcripts obtained from all the samples after analysed for hierarchical clustering. **C** PCA plot to verify the sample distribution between AF (purple circles) and sham (green circles) groups. Component 1 showed a deviation of 67.3% between the groups and component 2 showed a 11.8% of deviation between the samples of sham group biological replicates. **D** Whisker plot showing the transcript distribution between the AF and sham biological replicates.

**Supplementary Figure 3.**
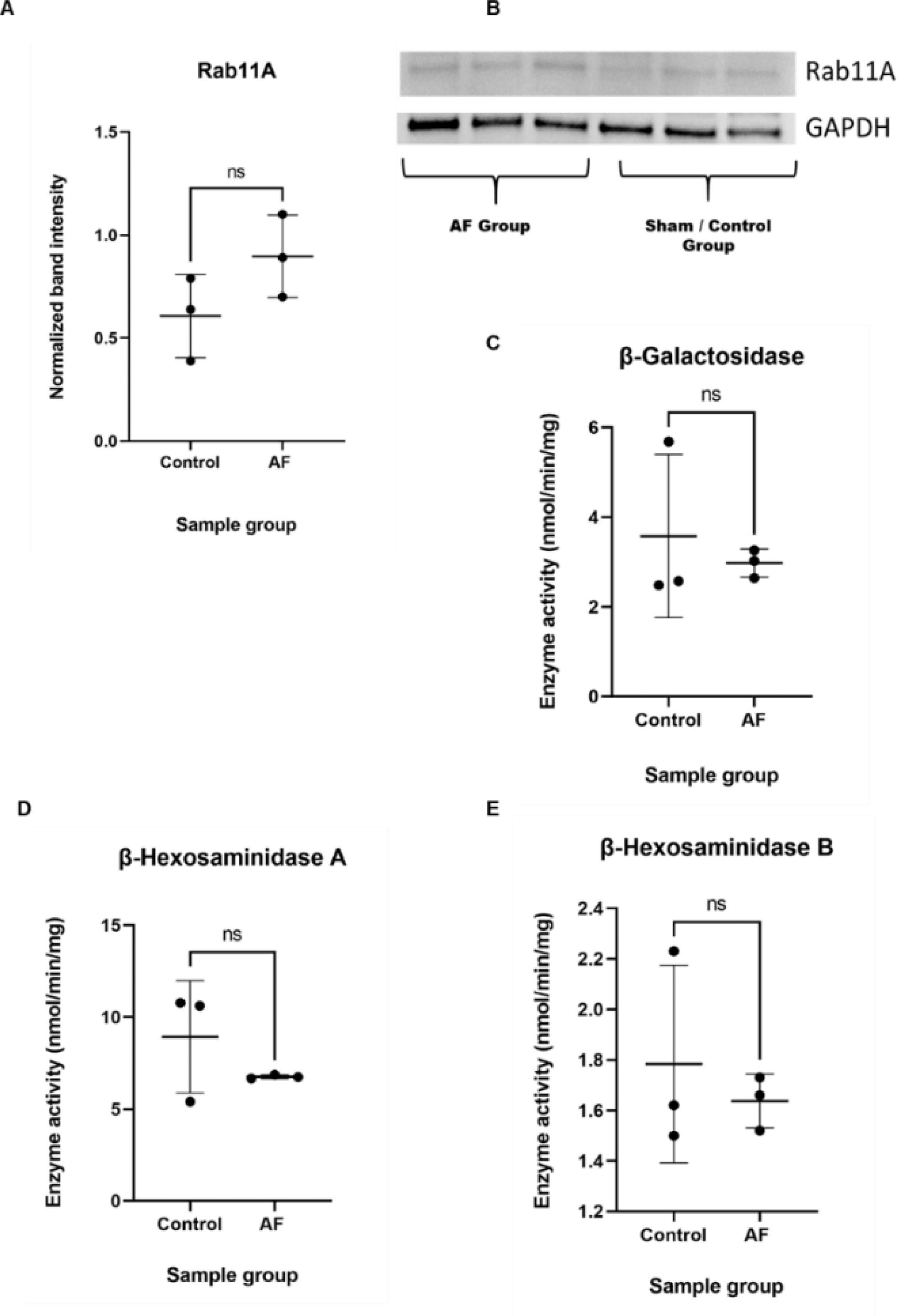
(relating to figure 1, 2 and): Western Blotting and Lysosomal Enzymatic Assays. **(A)** Western blots were performed on N=3 biological samples of each sham and AF group. Rab11A (24-25 kDa) and GAPDH control (37 kDa) were detected. **(B)** The normalised band intensities of Rab11A data are presented as mean±SD. The normalised band intensity value for the control group was 0.61±0.20, and for the AF group at 0.90±0.20. After performing a one-way t-test, the normalised protein band intensity showed a trend towards a significant upregulation in the AF group (*p* = 0.07). **(C)** β galactosidase activity is displayed as mean±SD. The AF group showed an enzymatic activity of 2.64, 3.02 and 3.26 nmol/min/mg, respectively, and the sham group showed 2.48, 2.57 and 5.68 nmol/min/mg. The mean enzymatic activity was 2.57±0.31 for the AF group and 3.57±1.8 for the control group. After performing a Two-way t-test, the β galactosidase activity of the AF group showed no significant regulation (*p*=0.60). **(D)** β hexosaminidase type-A activity of the AF group (N=3) showed 6.66, 6.74 and 6.87 nmol/min/mg, respectively, and the sham group showed 5.4, 10.61 and 10.78 nmol/min/mg activity. The normalised mean enzymatic activity was at 8.93±3.1 for the control group and 6.75±0.11 for the AF group, and β hexosaminidase type-A activity of the AF group showed no significant regulation (*p*=0.28). **(E)** AF group showed (N=3) 1.52, 1.66 and 1.73 nmol/min/mg of β hexosaminidase type B activity, and the sham group (N=3) showed 1.5, 1.62 and 2.23 nmol/min/mg of β hexosaminidase type B activity, The mean enzymatic activity was at 1.8±0.39 for the control group and 1.6±0.11 for AF group. No significant regulation was observed in β hexosaminidase type B activity (*p*=0.56).

**Supplementary Figure 4.**
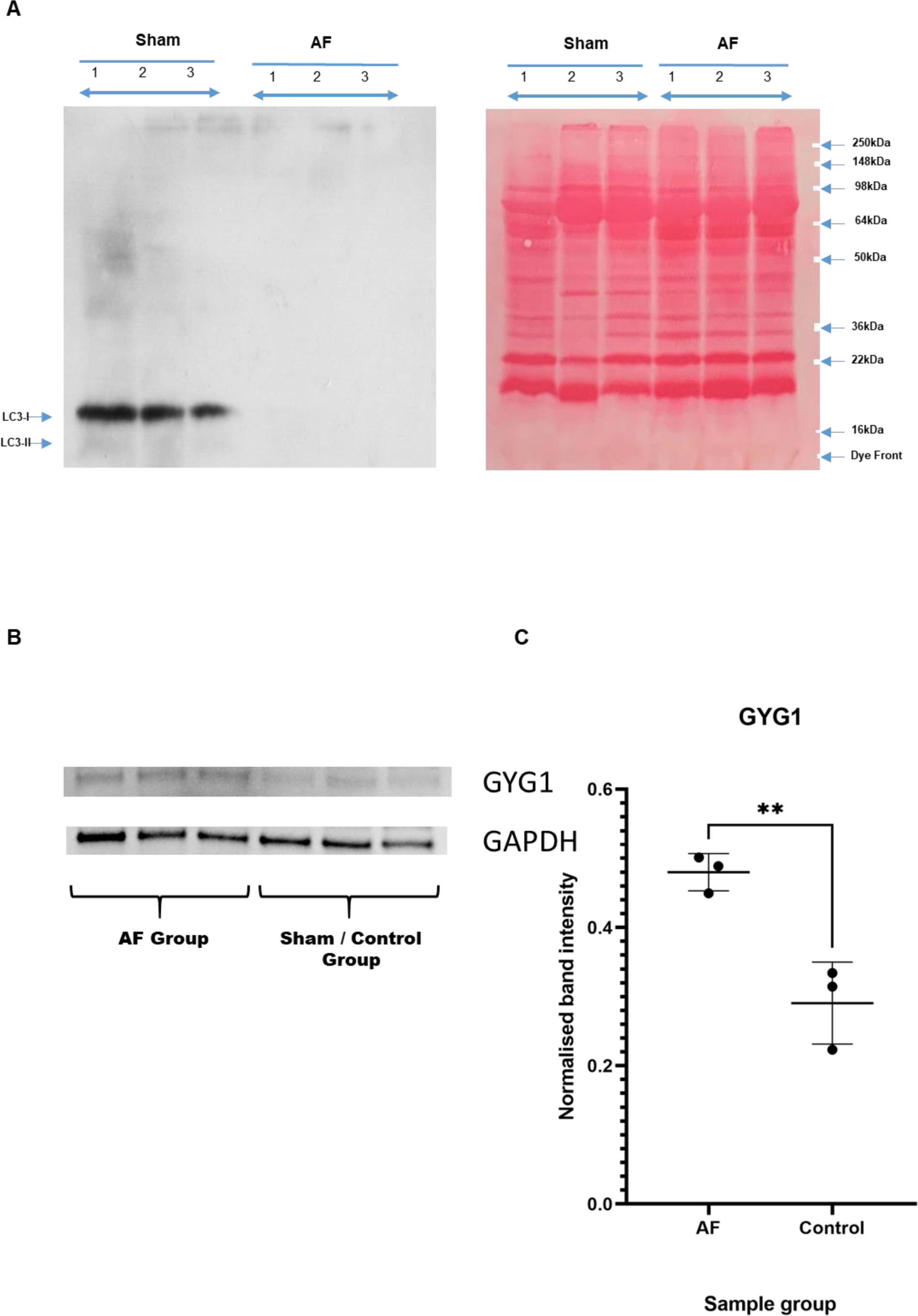
(relating to figure 1, 2 and 3) **A** The absence of the LC3-I and LC3-II proteins showed an overactive autophagic flux in the LA tissue of the AF goat model, and a Ponceau blot of the LC3-I and LC3-II membrane was prepared to observe the equal protein loading. **B and C** Western Blotting of Glycogenin-1 and LC3-I and II. (A &B) western blots displayed a significant upregulation of GYG-1 protein in the LA tissue of the AF goat model (*p* = *0.007*).

**Supplementary Figure 5.**
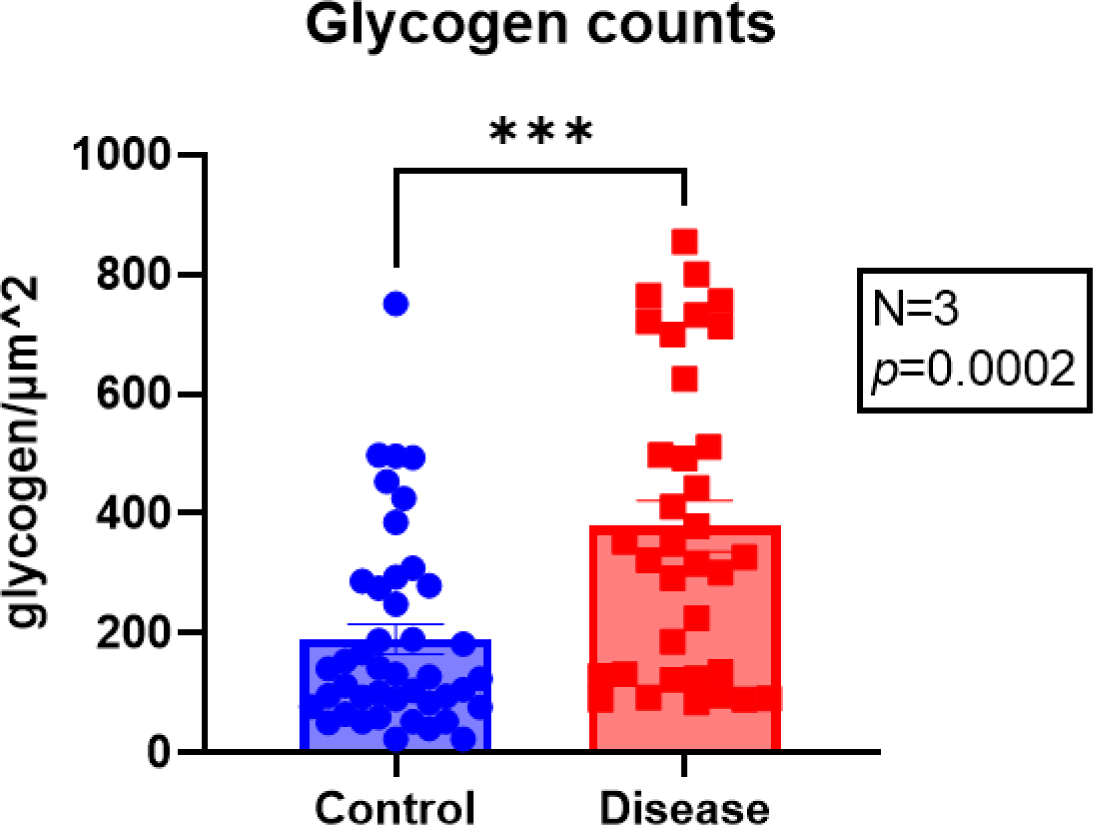
(relating to Figure 6) Electron microscopy image analysis to assess changes in glycogen in LA AF vs. Sham control samples. Increased glycogen levels was observed in AF samples (disease mean glycogen counts 378.7±42.6/μm^2 vs. control 190.5±24.9/μm^2, N=3, *p*=0.0002, student t-test).

**Supplementary Figure 6.**
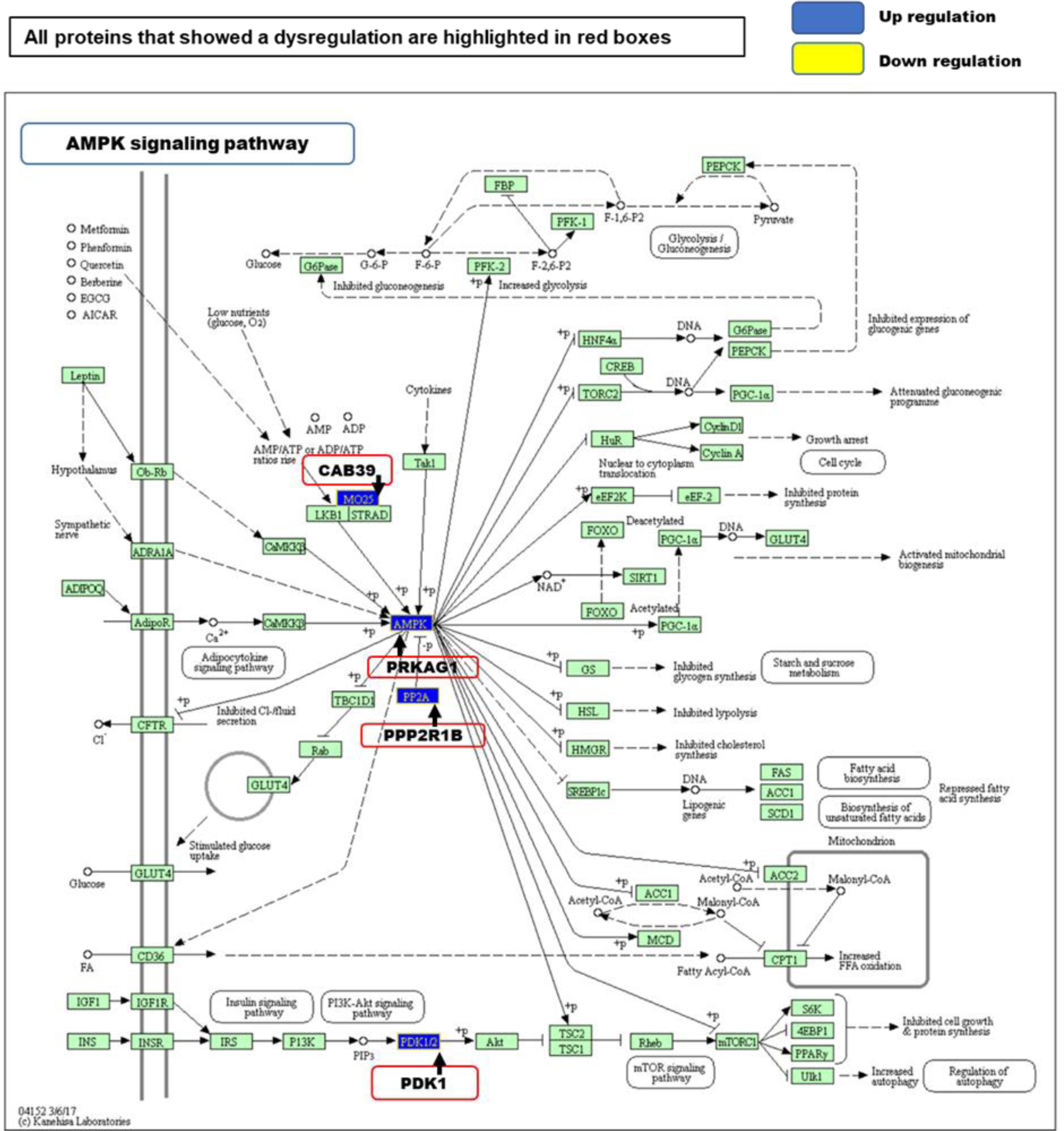
(relating to figure 1B) The level of upregulation of the proteins observed are respectively: CAB39/MO25 log2 0.54-fold, PRKAG1/AMPK log2 0.77-fold, PPP2R1B/PP2A log2 0.77-fold and PDK1 log2 0.56-fold. (Arrow = Protein name, protein complex name or protein function. Red bordered boxes= regulated proteins from the query data set. +P = phosphorylating events. Blue = up regulation (log2 fold-change) and Yellow= down regulation (log2 fold-change)). Pathway analysed using KeGG database.

**Supplementary Figure 7.**
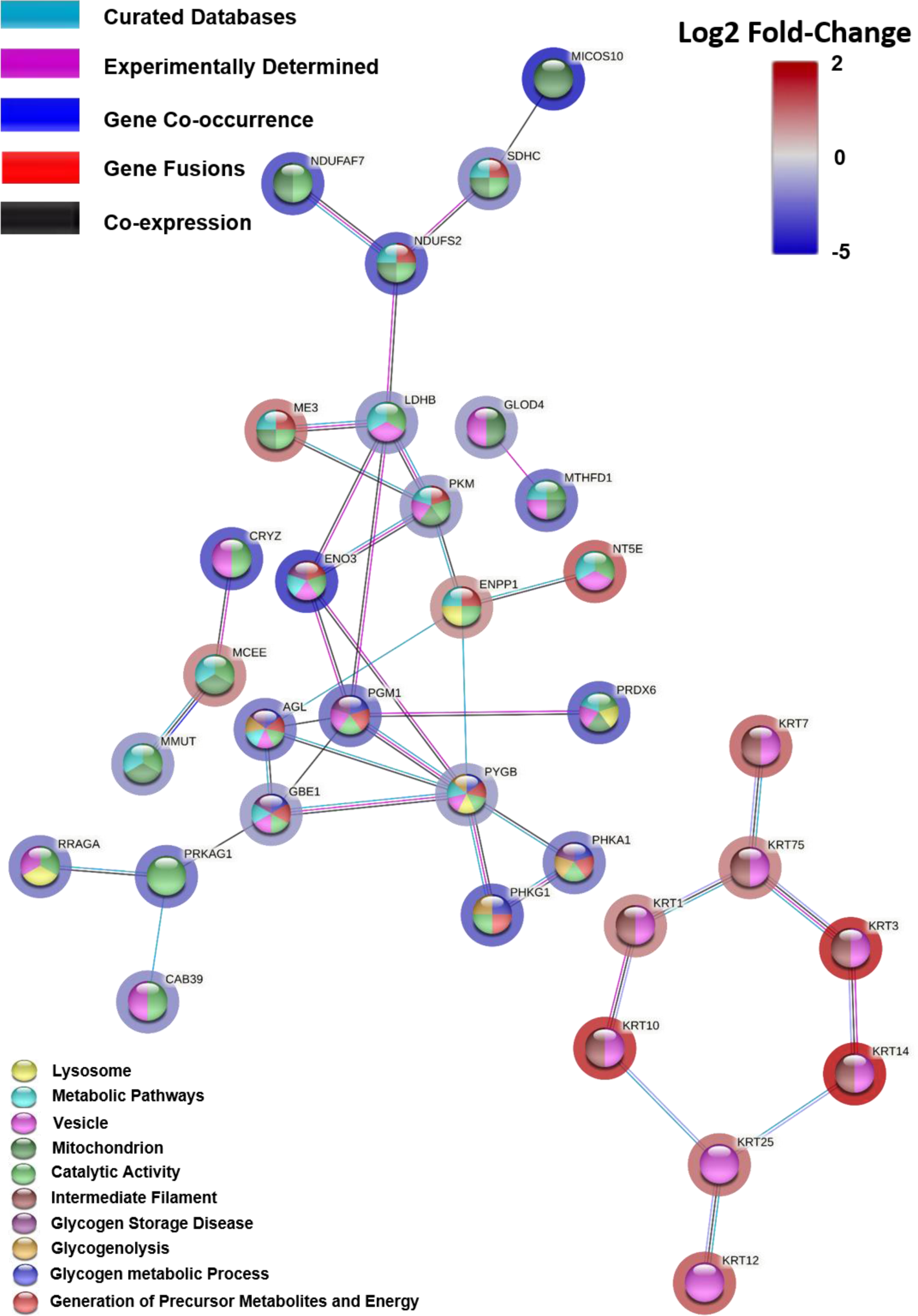
(relating to figure 1B): TL Network Cluster 5. The most significant proteins that have a functional role of regulatory and catalytic activity are clustered together Edges based on curated databases, experiments, gene co-occurrences, gene fusions, co-expressions and the nodes are coloured according to their functional enrichments. The halo around the nodes display the significant log2 fold-change.

**Supplementary Figure 8.**
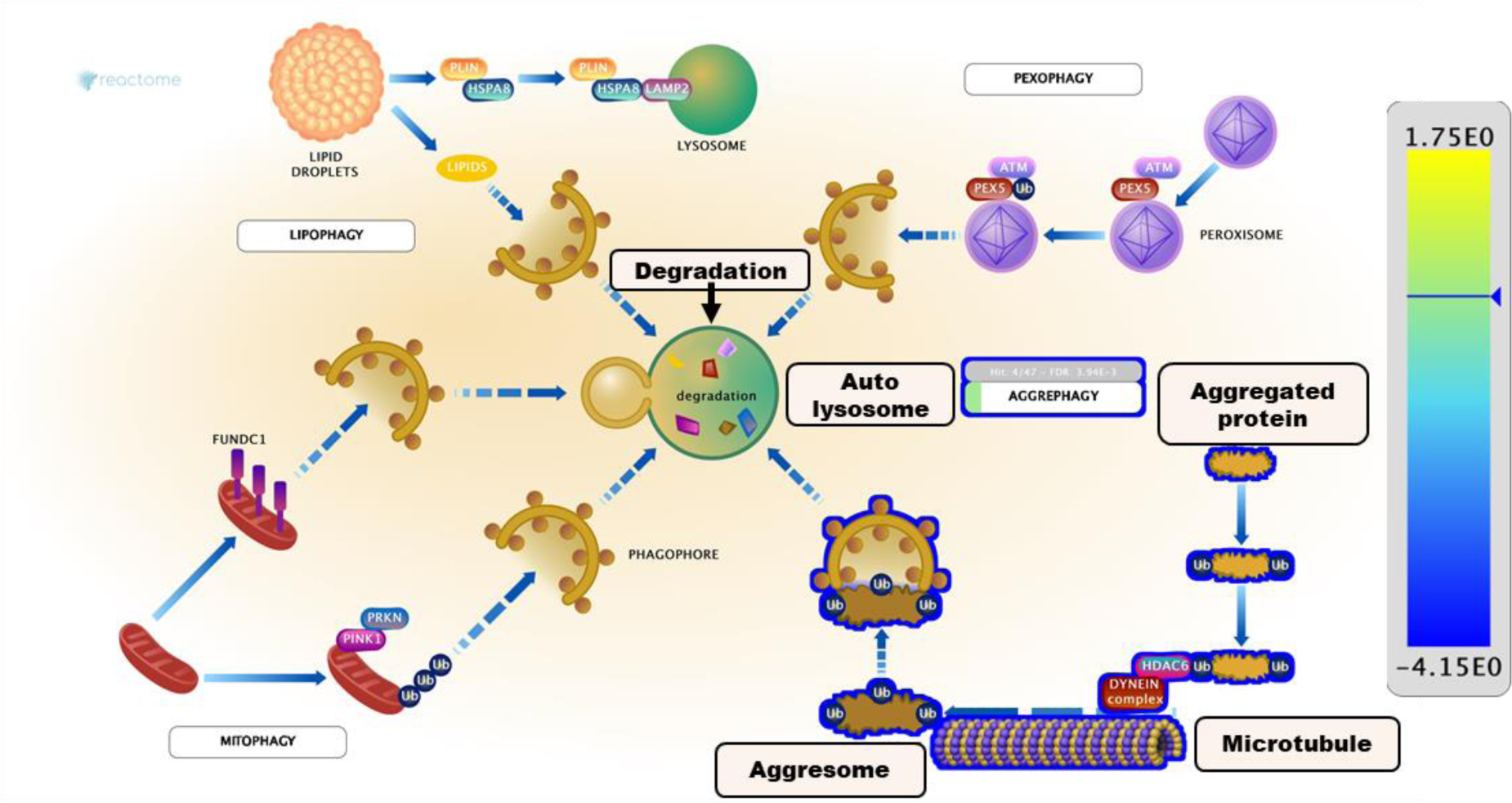
(relating to figure 1) Aggresome formation. The majority of the misfolded protein degradation was detected by then over expressed ubiquitin levels in the left atrial cells of the AF goat model. (Colour code= yellow to blue gradient; yellow represent highest up regulation while blue represent highest down regulation. Grey box above the aggrephagy label indicates the pathway entities detected in the query dataset, the total number of entities in the pathway, and the FDR corrected probability score. The protein complexes with complete or part-coloured representation showed the total molecules present from the query data set to complete the complex. Arrow = Protein, protein complex name or protein function). Figure created using www.reactome.com pathway analysis.

**Supplementary Figure 9.**
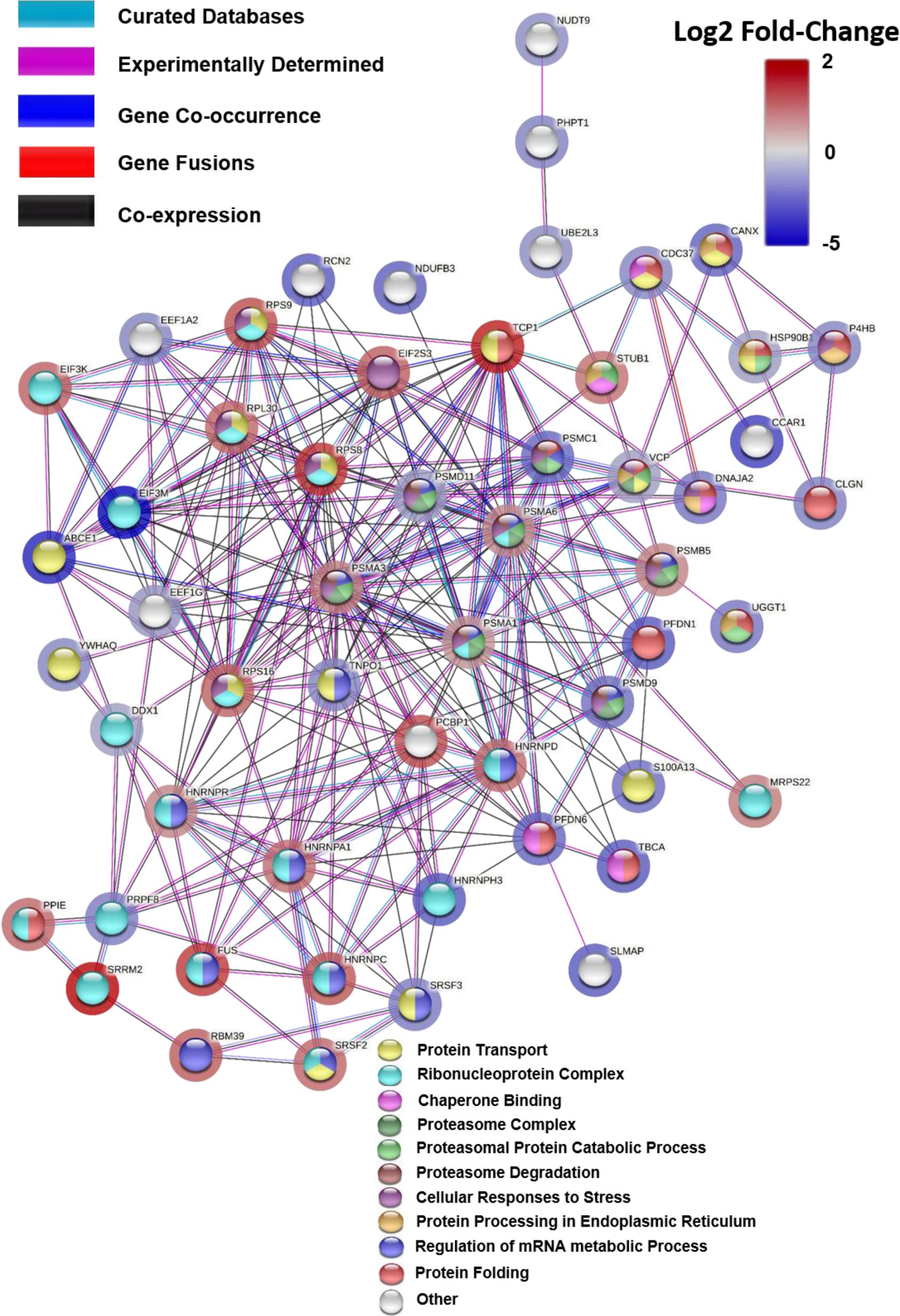
(relating to figure 1B): TL Network cluster 3. The most significant proteins that were regulated as part of stress induced stimuli are clustered together and Edges based on curated databases, experiments, gene co-occurrences, gene fusions, co-expressions and the nodes are coloured according to their functional enrichments. The halo around the nodes display the significant log2 fold-change.

**Supplementary Figure 10.**
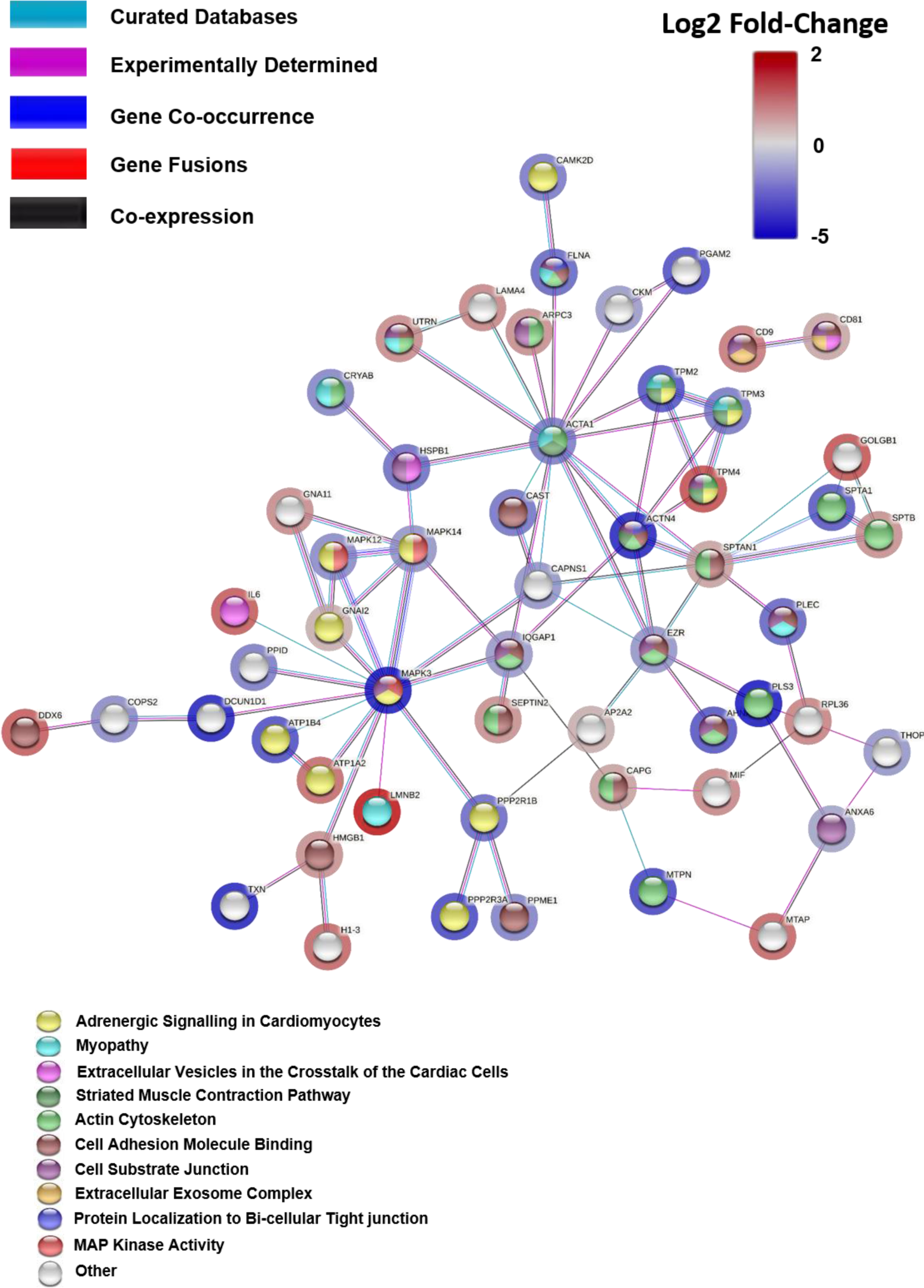
(relating to figure 1B): TL Network cluster 1. The most significant proteins that represented dysfunctional atrial conduction are clustered together and Edges based on curated databases, experiments, gene co-occurrences, gene fusions, co-expressions and the nodes are coloured according to their functional enrichments. The halo around the nodes display the significant log2 fold-change.

**Supplementary Figure 11.**
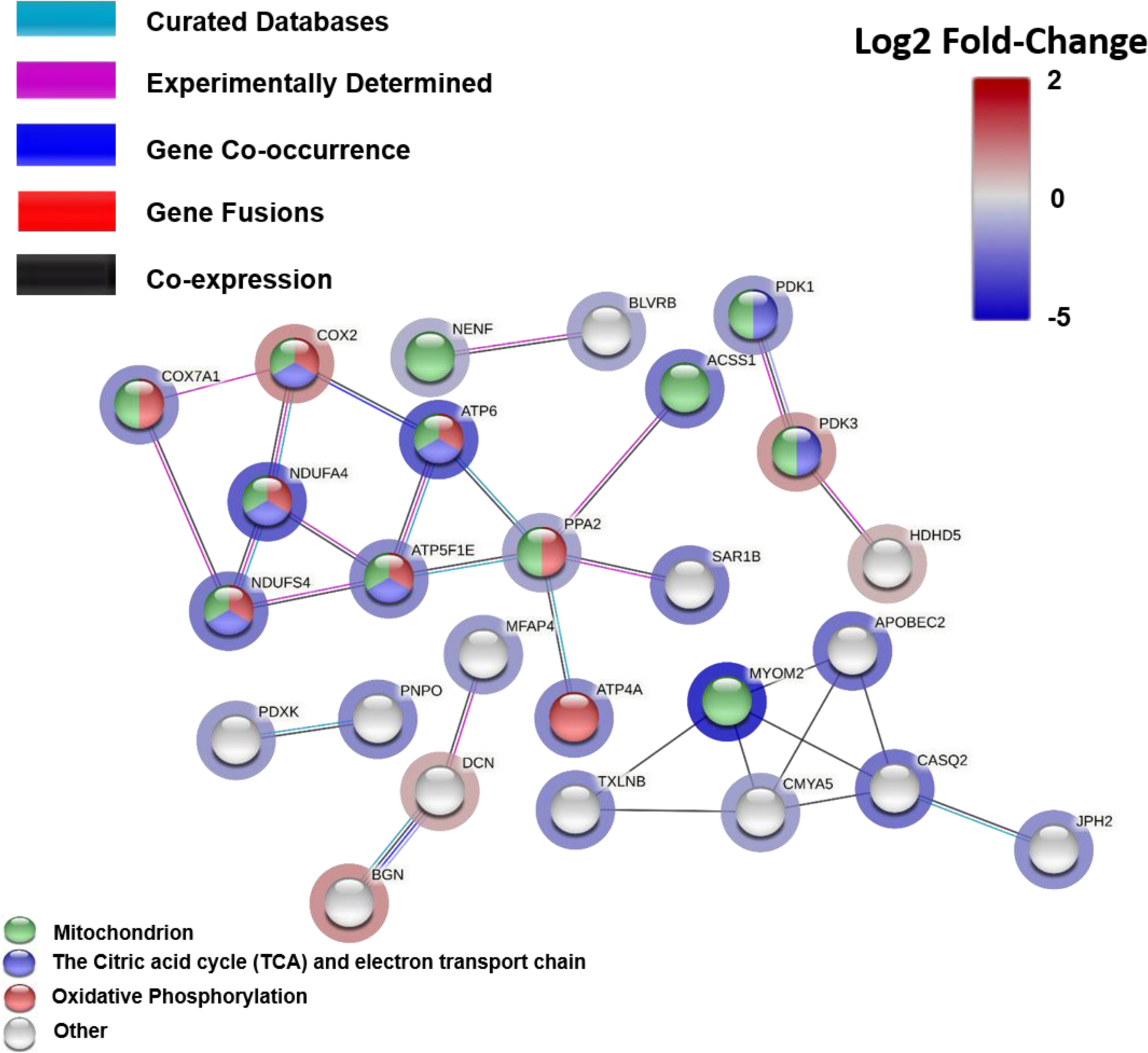
(relating to figure 1B): TL Network Cluster 4. The most significant proteins that belong to the metabolism of mitochondria are clustered together and Edges based on curated databases, experiments, gene co-occurrences, gene fusions, co-expressions and the nodes are coloured according to their functional enrichments. The halo around the nodes display the significant log2 fold-change.

**Supplementary Figure 12.**
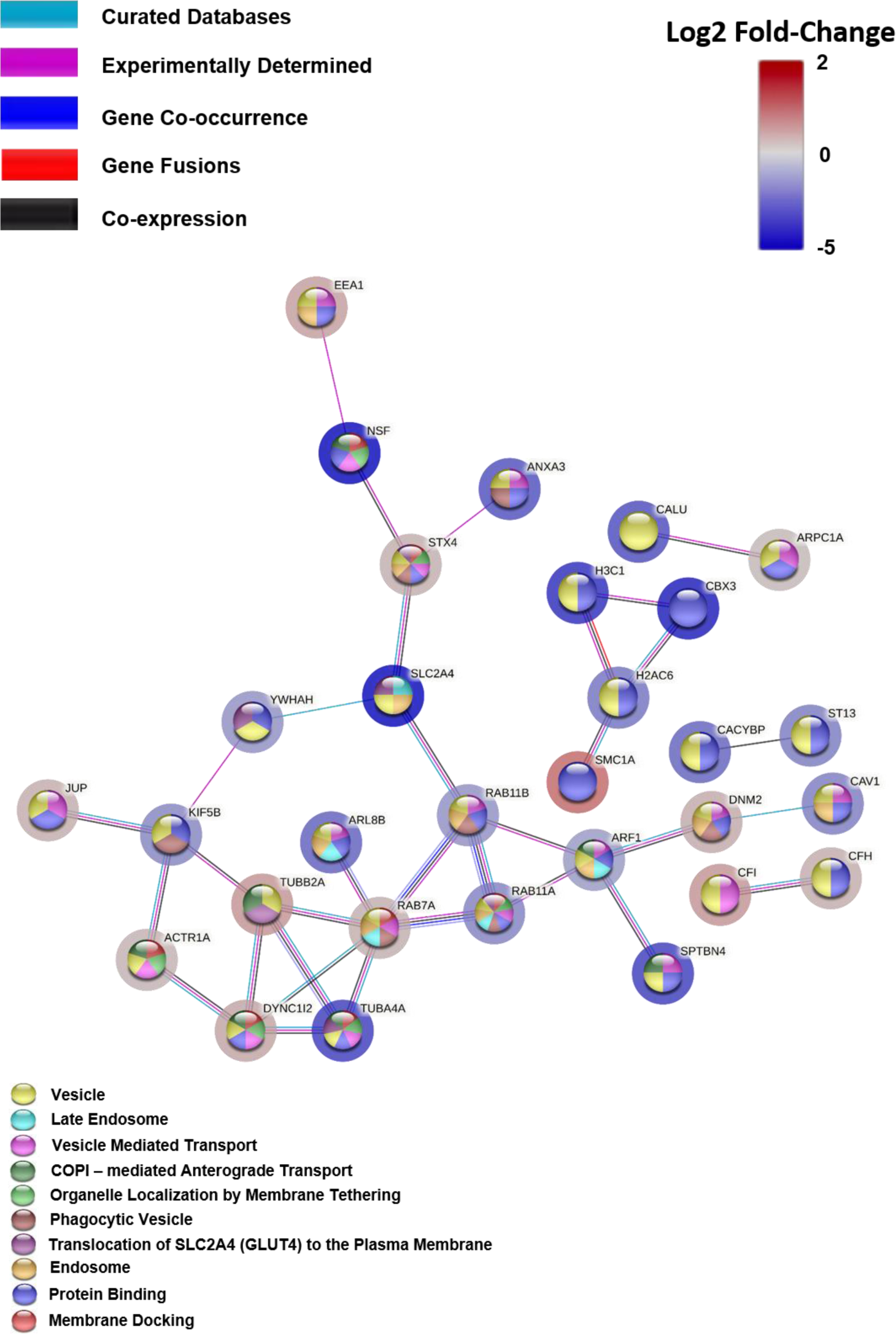
(relating to figure 1B): TL Network Cluster 6. The most significant proteins that have a functional role of trafficking and transport are clustered together and the Edges based on curated databases, experiments, gene co-occurrences, gene fusions, co-expressions and the nodes are coloured according to their functional enrichments. The halo around the nodes display the significant log2 fold-change.

**Supplementary Figure 13.**
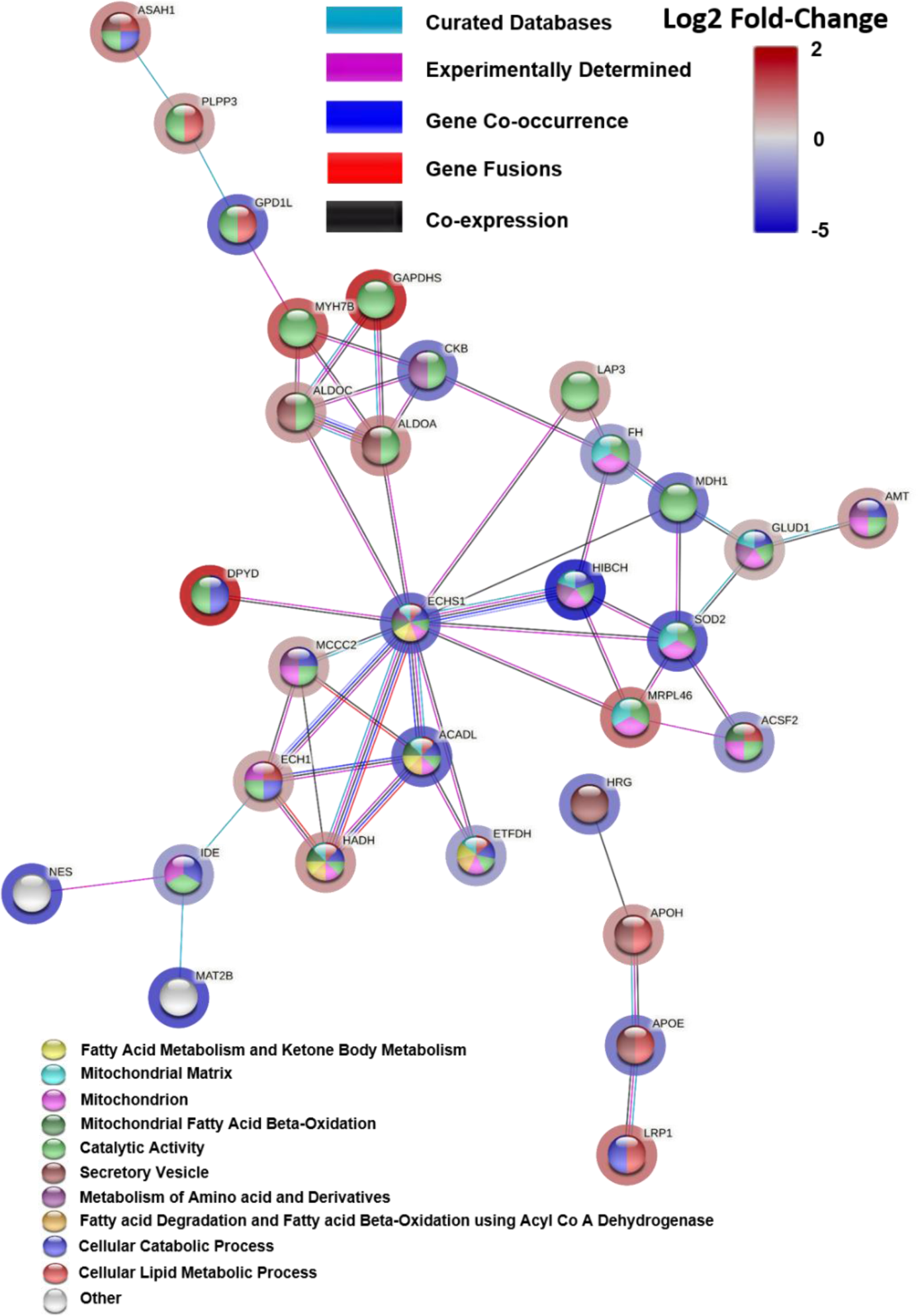
(relating to figure 1B): TL Network cluster 2. The most significant proteins that represented cellular metabolisms are clustered together and Edges based on curated databases, experiments, gene co-occurrences, gene fusions, co-expressions and the nodes are coloured according to their functional enrichments. The halo around the nodes display the significant log2 fold-change.

**Supplementary Figure 14.**
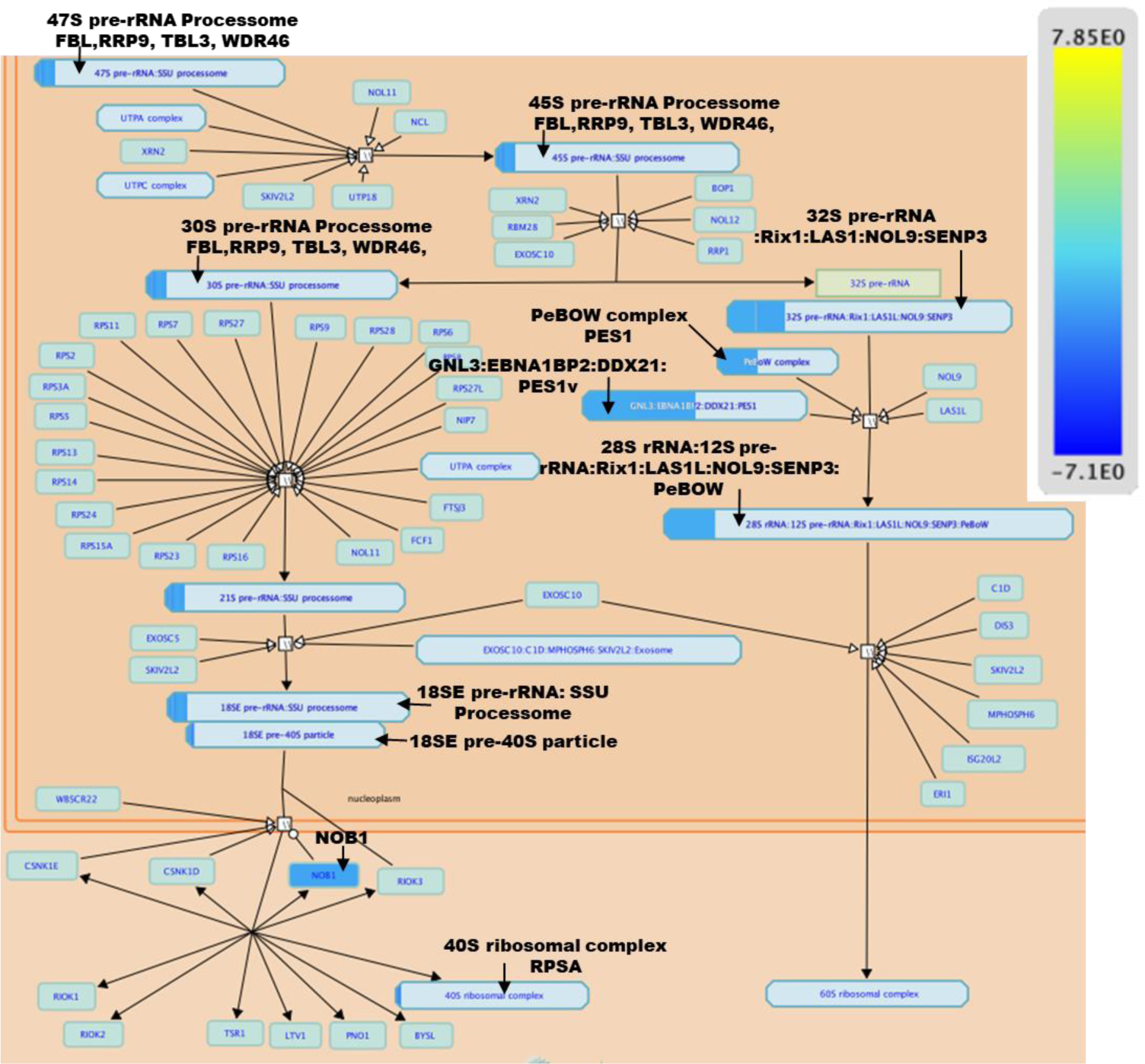
(relating to figure 1B) Transcriptomics analysis showing the Down regulation of the rRNA processing pathway in the nucleolus and cytosol of AF goat model. (Arrow = Gene name, protein complex name or protein function. Yellow = up regulation (log2 fold-change) and Blue= down regulation (log2 fold-change)). Pathway analysed using Reactome database.

