## Supplementary material for "Compartmentalisation proteomics revealed endolysosomal protein network changes in a goat model of atrial fibrillation": Supplementary File 1.docx

### Supplementary File 1 contains;

### Table 1: The complete list of the most significantly regulated genes of the LA of AF goat model obtained from Transcriptomics analysis

### Table 2: The complete list of the most significantly regulated proteins of the TL, LA of AF goat model obtained from Proteomics analysis

| Gene Name | Fold change (Log2) |
| --- | --- |
| SAMD9 | 5.27 |
| TRIM5 | 5.01 |
| GBP2 | 4.22 |
| RNF213 | 4.08 |
| IFIT1 | 4.03 |
| DDX58 | 3.94 |
| MX2 | 3.93 |
| RGS5 | 3.92 |
| BST2 | 3.62 |
| HLA-E | 3.27 |
| GBP1 | 3.26 |
| COL6A6 | 3.17 |
| CHRDL1 | 2.99 |
| TRANK1 | 2.86 |
| AGL | 2.85 |
| ZC3HAV1 | 2.81 |
| HCN1 | 2.80 |
| KLHL31 | 2.80 |
| PPFIBP1 | 2.75 |
| TRIM5 | 2.74 |
| SDPR | 2.74 |
| FRAS1 | 2.72 |
| GPR22 | 2.71 |
| MLKL | 2.68 |
| GADL1 | 2.62 |
| CMPK2 | 2.59 |
| FAM13B | 2.57 |
| FNDC1 | 2.56 |
| SLC25A30 | 2.49 |
| PARP12 | 2.47 |
| CACNA2D1 | 2.45 |
| TRIM56 | 2.45 |
| BCL2L14 | 2.42 |
| KIAA1033 | 2.41 |
| HES4 | 2.40 |
| HNRNPH2 | 2.39 |
| MB21D1 | 2.37 |
| MORC3 | 2.36 |
| LAMC2 | 2.36 |
| LPL | 2.35 |
| GREB1L | 2.34 |
| LZTS1 | 2.33 |
| TRIM5 | 2.32 |
| PRKAG3 | 2.31 |
| ZNFX1 | 2.30 |
| IFI44L | 2.28 |
| NUP43 | 2.28 |
| WNT5A | 2.27 |
| ZNF106 | 2.26 |
| GRB7 | 2.26 |
| ALG13 | 2.26 |
| FBXO40 | 2.25 |
| ITGB6 | 2.22 |
| SYNPO2 | 2.20 |
| PDP1 | 2.20 |
| PRKAA2 | 2.19 |
| VCPIP1 | 2.17 |
| TNFRSF19 | 2.13 |
| HERC6 | 2.12 |
| SLC2A4 | 2.10 |
| CYBB | 2.06 |
| PAQR3 | 2.06 |
| TIFA | 2.06 |
| CMTR2 | 2.05 |
| TANC1 | 2.05 |
| ZNF382 | 2.03 |
| SLC25A42 | 2.01 |
| CCNG2 | 1.97 |
| ITIH5 | 1.97 |
| MYO5A | 1.95 |
| C1orf109 | 1.93 |
| CFAP54 | 1.93 |
| DENND6A | 1.91 |
| SLC16A7 | 1.91 |
| NEBL | 1.90 |
| COL15A1 | 1.89 |
| BBS7 | 1.89 |
| DAXX | 1.88 |
| MEGF6 | 1.87 |
| MYO1B | 1.84 |
| ZNF292 | 1.83 |
| AGBL2 | 1.81 |
| B3GALT2 | 1.81 |
| NRK | 1.79 |
| EXO5 | 1.78 |
| THSD7B | 1.77 |
| ACSS1 | 1.77 |
| ZNF84 | 1.76 |
| TCAIM | 1.74 |
| SEMA4F | 1.72 |
| EIF2AK2 | 1.72 |
| TECRL | 1.72 |
| ENOSF1 | 1.71 |
| SCARA5 | 1.70 |
| ZFPM2 | 1.69 |
| PPP2R3A | 1.69 |
| TRDN | 1.62 |
| ANKRD33B | 1.62 |
| MOB3C | 1.61 |
| SYNE1 | 1.60 |
| C21orf91 | 1.59 |
| CFAP61 | 1.59 |
| DNAJC13 | 1.59 |
| PHACTR4 | 1.59 |
| DTX3L | 1.59 |
| ROR2 | 1.57 |
| FAM45A | 1.55 |
| SLC1A4 | 1.54 |
| ZNF132 | 1.53 |
| MGAT5 | 1.52 |
| SNX24 | 1.51 |
| INPP4B | 1.51 |
| ENTPD2 | 1.50 |
| HDAC9 | 1.49 |
| ADAM33 | 1.48 |
| CERCAM | 1.47 |
| PRRG4 | 1.47 |
| GBP6 | 1.47 |
| CABLES1 | 1.47 |
| FGF7 | 1.46 |
| FAM214A | 1.45 |
| SESN1 | 1.45 |
| GPM6B | 1.44 |
| GANC | 1.44 |
| TRIM2 | 1.44 |
| DST | 1.43 |
| PPIP5K1 | 1.43 |
| TXNDC16 | 1.41 |
| PFKFB2 | 1.41 |
| PIWIL2 | 1.41 |
| ARL5A | 1.40 |
| FAM161A | 1.39 |
| RAPGEF2 | 1.37 |
| EHBP1 | 1.37 |
| BVES | 1.37 |
| WDR35 | 1.37 |
| SH3TC2 | 1.36 |
| PAQR9 | 1.36 |
| NABP1 | 1.33 |
| TRIM26 | 1.33 |
| ARMC2 | 1.33 |
| ASB14 | 1.33 |
| PCYOX1 | 1.32 |
| PLSCR1 | 1.32 |
| COL5A3 | 1.32 |
| METTL24 | 1.31 |
| CAB39L | 1.30 |
| NAP1L3 | 1.30 |
| STK39 | 1.30 |
| KNTC1 | 1.30 |
| GMCL1 | 1.30 |
| KIF21A | 1.29 |
| DMXL2 | 1.29 |
| SLC35D1 | 1.29 |
| CPED1 | 1.28 |
| ATP1B4 | 1.28 |
| CCND1 | 1.28 |
| MPV17 | 1.26 |
| ADAT2 | 1.25 |
| LRRC72 | 1.24 |
| PNPT1 | 1.23 |
| GAS1 | 1.23 |
| KIAA1109 | 1.21 |
| PLP1 | 1.20 |
| PKIA | 1.18 |
| SYCP3 | 1.17 |
| PDK3 | 1.17 |
| HEATR5B | 1.16 |
| NET1 | 1.16 |
| IDH3A | 1.15 |
| F13A1 | 1.15 |
| AHCYL2 | 1.15 |
| RPGRIP1L | 1.15 |
| MGME1 | 1.14 |
| WDR19 | 1.14 |
| TRIM24 | 1.13 |
| CTTNBP2 | 1.12 |
| CA5B | 1.11 |
| TTC3 | 1.11 |
| CHN1 | 1.11 |
| ZNF808 | 1.11 |
| DNAJC28 | 1.10 |
| TNRC6A | 1.10 |
| SLC35F1 | 1.09 |
| AASS | 1.07 |
| WNK2 | 1.07 |
| FUCA2 | 1.07 |
| TSPAN12 | 1.07 |
| EGFLAM | 1.05 |
| TRIM23 | 1.05 |
| TET2 | 1.05 |
| WWP1 | 1.04 |
| PHF11 | 1.03 |
| PBX1 | 1.02 |
| MAT2B | 1.02 |
| AGPAT5 | 1.02 |
| C2CD5 | 1.01 |
| ATG4A | 0.99 |
| HOMER1 | 0.98 |
| LAMA2 | 0.98 |
| NUBPL | 0.96 |
| NTN4 | 0.95 |
| ST5 | 0.95 |
| TGFBI | 0.95 |
| TUBGCP5 | 0.94 |
| STRADB | 0.94 |
| SFMBT2 | 0.93 |
| LETMD1 | 0.93 |
| XRN2 | 0.93 |
| DCAF17 | 0.93 |
| RALGPS2 | 0.93 |
| CDKN2AIP | 0.92 |
| B2M | 0.92 |
| PMS2 | 0.92 |
| MTFR1L | 0.91 |
| C10orf2 | 0.90 |
| ZMYM4 | 0.90 |
| DZANK1 | 0.90 |
| MRPS35 | 0.88 |
| PRKCE | 0.86 |
| DYNC2H1 | 0.86 |
| DNM1L | 0.84 |
| PLEKHA5 | 0.83 |
| PNRC1 | 0.82 |
| DDX11 | 0.81 |
| IKBKAP | 0.79 |
| APBB3 | 0.77 |
| BBS12 | 0.75 |
| MYO9A | 0.73 |
| GTF3C3 | 0.70 |
| FAM13A | 0.66 |
| SLC30A4 | 0.66 |
| AP5S1 | 0.64 |
| RASA1 | 0.51 |
| EPC1 | 0.50 |
| BCCIP | -0.58 |
| TWISTNB | -0.59 |
| NCL | -0.59 |
| SLC48A1 | -0.60 |
| PDAP1 | -0.63 |
| NBL1 | -0.63 |
| PDIA3 | -0.63 |
| PRPF38A | -0.64 |
| TOR1A | -0.64 |
| EIF3I | -0.65 |
| MAPK3 | -0.65 |
| TRADD | -0.65 |
| KDM1A | -0.66 |
| UNC45A | -0.66 |
| MBD2 | -0.66 |
| PSMB2 | -0.66 |
| TTC27 | -0.66 |
| FDFT1 | -0.66 |
| TBC1D10A | -0.68 |
| PAK1IP1 | -0.68 |
| TPCN2 | -0.68 |
| TEX10 | -0.68 |
| NCLN | -0.69 |
| POLR2E | -0.70 |
| C18orf8 | -0.70 |
| PITPNA | -0.71 |
| NT5C3B | -0.71 |
| TMED9 | -0.71 |
| SDCCAG3 | -0.73 |
| TMEM9 | -0.73 |
| B4GALT7 | -0.73 |
| ALG3 | -0.73 |
| HN1 | -0.73 |
| ORAOV1 | -0.74 |
| PBX2 | -0.74 |
| NADK | -0.75 |
| HGS | -0.76 |
| DEXI | -0.77 |
| HSPA5 | -0.78 |
| NOL6 | -0.78 |
| B9D1 | -0.78 |
| DUSP12 | -0.79 |
| SSRP1 | -0.79 |
| UBTD1 | -0.79 |
| CDK5RAP2 | -0.79 |
| ADRM1 | -0.79 |
| ARRB1 | -0.79 |
| ZNF622 | -0.80 |
| METTL1 | -0.80 |
| CTU1 | -0.80 |
| MARK4 | -0.80 |
| VIMP | -0.82 |
| PDGFA | -0.82 |
| SEC23B | -0.82 |
| UBE2J2 | -0.82 |
| VARS | -0.83 |
| PSMD8 | -0.83 |
| B3GAT3 | -0.83 |
| RING1 | -0.83 |
| RABL6 | -0.83 |
| PLEKHJ1 | -0.84 |
| DYNLT1 | -0.85 |
| PGS1 | -0.85 |
| NGDN | -0.85 |
| CYB561D2 | -0.85 |
| ACOT7 | -0.86 |
| PRKCSH | -0.86 |
| HSF1 | -0.86 |
| ABHD12 | -0.86 |
| ZNF706 | -0.86 |
| TOLLIP | -0.87 |
| INO80E | -0.87 |
| HS1BP3 | -0.87 |
| CEBPG | -0.87 |
| TFPT | -0.89 |
| SEC61B | -0.89 |
| CRTC2 | -0.89 |
| CALR | -0.90 |
| GNB2 | -0.90 |
| ZPR1 | -0.91 |
| PELO | -0.91 |
| RASIP1 | -0.91 |
| HMOX2 | -0.91 |
| TSEN34 | -0.92 |
| GARS | -0.92 |
| COLGALT1 | -0.93 |
| HDAC5 | -0.94 |
| SNRPB2 | -0.94 |
| VIM | -0.94 |
| SDE2 | -0.95 |
| GNL3 | -0.95 |
| LSP1 | -0.95 |
| RHOC | -0.96 |
| RSL1D1 | -0.96 |
| SRM | -0.96 |
| RPL29 | -0.96 |
| EGLN2 | -0.96 |
| DDX49 | -0.97 |
| CHMP4B | -0.97 |
| REXO4 | -0.97 |
| SEC61A1 | -0.97 |
| UTP3 | -0.97 |
| GAR1 | -0.98 |
| MYDGF | -0.98 |
| GRINA | -0.98 |
| YPEL3 | -0.98 |
| TRMU | -0.99 |
| WDR5 | -0.99 |
| NOP9 | -0.99 |
| FAM212B | -0.99 |
| ABHD14A | -1.00 |
| TXNDC11 | -1.00 |
| ATF4 | -1.00 |
| RUVBL1 | -1.00 |
| NR1H2 | -1.01 |
| BOP1 | -1.02 |
| CSNK1G2 | -1.02 |
| BRIX1 | -1.03 |
| BEAN1 | -1.03 |
| RBM42 | -1.03 |
| SERP1 | -1.03 |
| CTDP1 | -1.03 |
| GPSM1 | -1.04 |
| KCNK7 | -1.04 |
| TRMT61A | -1.04 |
| EVA1C | -1.05 |
| DBNDD2 | -1.05 |
| WDR4 | -1.05 |
| EIF4A1 | -1.05 |
| EXT1 | -1.06 |
| S100A11 | -1.07 |
| RBM15B | -1.07 |
| GRWD1 | -1.07 |
| PPP1R14B | -1.07 |
| TXN | -1.07 |
| TNIP2 | -1.08 |
| SH3TC1 | -1.08 |
| SIGIRR | -1.09 |
| MANF | -1.09 |
| RAC1 | -1.09 |
| GMPPB | -1.10 |
| EBNA1BP2 | -1.10 |
| ODC1 | -1.10 |
| RALY | -1.10 |
| NOC2L | -1.10 |
| HM13 | -1.11 |
| PFKL | -1.11 |
| TMEM208 | -1.12 |
| SSBP4 | -1.12 |
| RPL29 | -1.13 |
| TRMT112 | -1.13 |
| AATF | -1.13 |
| CCM2 | -1.13 |
| METTL6 | -1.13 |
| ARPC4 | -1.13 |
| TPRA1 | -1.14 |
| MECP2 | -1.14 |
| CCDC71 | -1.14 |
| PRELID1 | -1.14 |
| PLCD1 | -1.15 |
| LMNA | -1.15 |
| PPP4C | -1.15 |
| NOP56 | -1.15 |
| GATSL3 | -1.15 |
| PAGR1 | -1.16 |
| RNF126 | -1.16 |
| TBL3 | -1.17 |
| SELO | -1.17 |
| RCN3 | -1.18 |
| WDR83 | -1.20 |
| POP1 | -1.20 |
| PES1 | -1.20 |
| ABHD1 | -1.20 |
| TNFRSF1A | -1.20 |
| C15orf39 | -1.21 |
| PTRH2 | -1.21 |
| GPATCH4 | -1.22 |
| LRFN3 | -1.23 |
| TALDO1 | -1.24 |
| SNRPB | -1.25 |
| ESAM | -1.25 |
| RRAD | -1.25 |
| SLC22A17 | -1.26 |
| CDKN1C | -1.27 |
| TUBB2A | -1.27 |
| ZDHHC12 | -1.27 |
| ARHGAP22 | -1.27 |
| NOB1 | -1.27 |
| ATOH8 | -1.27 |
| CDC42EP1 | -1.27 |
| SLC27A4 | -1.28 |
| FAM89A | -1.28 |
| EXOSC5 | -1.29 |
| NLE1 | -1.30 |
| SDF2L1 | -1.30 |
| SNCAIP | -1.31 |
| KCTD15 | -1.31 |
| LCAT | -1.31 |
| CARS | -1.32 |
| RRP1 | -1.33 |
| SEMA6B | -1.33 |
| SPIDR | -1.34 |
| RHOG | -1.34 |
| ARRDC5 | -1.34 |
| RENBP | -1.35 |
| PXDC1 | -1.35 |
| JOSD2 | -1.35 |
| WDR46 | -1.36 |
| IFRD2 | -1.37 |
| RRS1 | -1.37 |
| ALPL | -1.37 |
| NPDC1 | -1.38 |
| CRELD2 | -1.38 |
| SMIM5 | -1.38 |
| FKBP11 | -1.41 |
| CNPY3 | -1.41 |
| GAS2L1 | -1.41 |
| UCHL1 | -1.41 |
| SH3BGRL3 | -1.42 |
| GATA2 | -1.43 |
| ECE1 | -1.44 |
| TGFB1 | -1.44 |
| ISYNA1 | -1.47 |
| NINJ2 | -1.48 |
| LIPE | -1.48 |
| MISP3 | -1.50 |
| DDIT3 | -1.50 |
| ZNF444 | -1.51 |
| TESC | -1.51 |
| FBL | -1.52 |
| RGS19 | -1.52 |
| RHOD | -1.53 |
| SLC3A2 | -1.53 |
| FHL3 | -1.53 |
| ITPKA | -1.55 |
| ARPC1B | -1.57 |
| WFIKKN1 | -1.57 |
| CARHSP1 | -1.58 |
| RRP9 | -1.61 |
| SH2D3C | -1.65 |
| ASGR1 | -1.67 |
| ZNF619 | -1.67 |
| ZNF593 | -1.69 |
| ID3 | -1.70 |
| CLU | -1.70 |
| EXOSC6 | -1.71 |
| N4BP3 | -1.72 |
| C1QTNF5 | -1.72 |
| SURF6 | -1.73 |
| RAMP2 | -1.75 |
| DOK3 | -1.76 |
| GGT5 | -1.81 |
| APRT | -1.81 |
| C11orf96 | -1.82 |
| RPL13 | -1.83 |
| CLIC1 | -1.83 |
| CLEC14A | -1.85 |
| CEBPA | -1.85 |
| PGF | -1.87 |
| ITIH4 | -1.87 |
| ENG | -1.88 |
| NT5C | -1.91 |
| IMPA2 | -1.93 |
| EVA1B | -1.93 |
| SLC16A5 | -1.94 |
| SLC35E4 | -1.94 |
| SERPINB6 | -1.95 |
| RPSA | -1.95 |
| DUSP26 | -1.98 |
| IER5L | -2.10 |
| MT1G | -2.16 |
| ZNF467 | -2.17 |
| TOR4A | -2.19 |
| IL21R | -2.22 |
| FXYD5 | -2.32 |
| CDC42EP5 | -2.40 |
| EGFL7 | -2.48 |
| FAM110D | -2.50 |
| IRX3 | -2.54 |
| ETV2 | -2.55 |
| GDPD3 | -2.76 |
| PLEKHO1 | -2.78 |
| CD300LB | -2.88 |
| HOXB2 | -2.89 |
| HES7 | -2.93 |
| RTN1 | -2.94 |
| SH2B2 | -2.99 |
| FAM167A | -3.02 |
| C1QTNF4 | -3.02 |
| MAT1A | -3.10 |
| ID1 | -3.17 |
| EXOC3L2 | -3.25 |
| PLVAP | -3.37 |
| FABP4 | -3.38 |
| SOX18 | -3.39 |
| IQCG | -4.09 |
| DLK1 | -4.32 |

Table 1: The most significantly regulated gene list obtained from transcriptomics analysis.

| Protein ID | Protein Name | Gene | Fold change |
| --- | --- | --- | --- |
| A0A452EB00_CAPHI | Eukaryotic translation initiation factor 3 subunit M | EIF3M | 1.75 |
| A0A452E9Y7_CAPHI | Myomesin-2 | MYOM2 | 1.73 |
| A0A452E134_CAPHI | Protein preY, mitochondrial | PYURF | 1.55 |
| A0A452FEB3_CAPHI | Glucose transporter type 4, insulin-responsive | SLC2A4 | 1.55 |
| A0A452EEQ2_CAPHI | Vesicle-fusing ATPase | NSF | 1.55 |
| A0A452FTN3_CAPHI | 3-hydroxyisobutyryl-CoA hydrolase, mitochondrial | HIBCH | 1.43 |
| A0A452F5Y9_CAPHI | Plastin-3 | PLS3 | 1.43 |
| A0A452F8U9_CAPHI | ATP-binding cassette sub-family E member 1 | ABCE1 | 1.43 |
| A0A452DTB9_CAPHI | MARVEL domain-containing protein | SYPL1 | 1.42 |
| A0A452FZL6_CAPHI | Amine oxidase | MAOB | 1.41 |
| A0A452FSW2_CAPHI | MICOS complex subunit MIC10 | MICOS10 | 1.40 |
| A0A452EU65_CAPHI | Mitogen-activated protein kinase | MAPK3 | 1.37 |
| A0A452E268_CAPHI | Chromobox protein homolog 3 | CBX3 | 1.36 |
| A0A452F5R6_CAPHI | Calcium-transporting ATPase | ATP2A3 | 1.33 |
| A0A452DV61_CAPHI | Huntingtin-interacting protein K | HYPK | 1.32 |
| A0A452ED15_CAPHI | DCN1-like protein | DCUN1D1 | 1.32 |
| A0A452EYY0_CAPHI | Tuftelin-interacting protein 11 | TFIP11 | 1.32 |
| A0A452E7A9_CAPHI | Alpha-actinin-4 | ACTN4 | 1.30 |
| A0A452EGP3_CAPHI | Thioredoxin | TXN | 1.29 |
| D2KMK0_CAPHI | ATP synthase subunit a | ATP6 | 1.24 |
| A0A452FB24_CAPHI | 2-phospho-D-glycerate hydro-lyase | ENO3 | 1.23 |
| A0A452DRQ7_CAPHI | Histone H3 | H3C1 | 1.22 |
| A5JSS3_CAPHI | NADH dehydrogenase (Ubiquinone) 1 alpha subcomplex 4 | NDUFA4 | 1.22 |
| A0A452DL17_CAPHI | SAP domain-containing protein | CCAR1 | 1.21 |
| A0A452EWZ3_CAPHI | SERPIN domain-containing protein | LOC102174166 | 1.19 |
| A0A452FYK7_CAPHI | BolA-like protein 2 | BOLA2B | 1.18 |
| A0A452EXE8_CAPHI | Calgranulin-B | S100A9 | 1.17 |
| A0A452GAJ0_CAPHI | Methionine adenosyltransferase 2 subunit beta | MAT2B | 1.10 |
| A0A452EAC5_CAPHI | Pentatricopeptide repeat domain-containing protein 3, mitochondrial | PTCD3 | 1.09 |
| A0A452DNE3_CAPHI | Tubulin alpha chain | TUBA4A | 1.09 |
| A0A452EPS0_CAPHI | Spectrin beta chain | SPTBN4 | 1.08 |
| A0A452DYZ1_CAPHI | Peptidylprolyl isomerase | LOC108634682 | 1.07 |
| HBA1_CAPHI | Hemoglobin subunit alpha-1 | HBA1 | 1.06 |
| A0A452FZM5_CAPHI | Protein arginine methyltransferase NDUFAF7 | NDUFAF7 | 1.06 |
| A0A452EZS5_CAPHI | PKS_ER domain-containing protein | CRYZ | 1.06 |
| A0A452G757_CAPHI | IF rod domain-containing protein | NES | 1.05 |
| A0A452F5N4_CAPHI | ANK_REP_REGION domain-containing protein | MTPN | 1.05 |
| A0A452DZF9_CAPHI | Matrix-remodeling-associated protein 7 | MXRA7 | 1.04 |
| A0A452E1L0_CAPHI | Serine/threonine-protein phosphatase 2A regulatory subunit B'' subunit alpha | PPP2R3A | 1.02 |
| A0A452E294_CAPHI | Superoxide dismutase | SOD2 | 1.01 |
| A0A452G6Y9_CAPHI | CMP/dCMP-type deaminase domain-containing protein | APOBEC2 | 1.01 |
| A0A452FXF1_CAPHI | Complex I-49kD | NDUFS2 | 1.01 |
| A0A452ERD0_CAPHI | Peptidase M20 domain-containing protein 2 | PM20D2 | 1.01 |
| A0A452EKU0_CAPHI | Calsequestrin | CASQ2 | 1.00 |
| A0A452DR91_CAPHI | Phosphoglycerate mutase | PGAM2 | 0.99 |
| A0A452G7L1_CAPHI | Heterogeneous nuclear ribonucleoprotein H3 | HNRNPH3 | 0.97 |
| A0A452G3A0_CAPHI | Calumenin | CALU | 0.97 |
| A0A452E6Y1_CAPHI | Prefoldin subunit 1 | PFDN1 | 0.97 |
| A0A452EE06_CAPHI | Cytochrome P450 2D14 | LOC102169002 | 0.97 |
| A0A452EW53_CAPHI | Reticulocalbin-2 | RCN2 | 0.97 |
| A0A452FC01_CAPHI | Myosin regulatory light chain 2 | MYL2 | 0.96 |
| A0A452FRP1_CAPHI | Protein-tyrosine-phosphatase | EPM2A | 0.96 |
| A0A452DMV1_CAPHI | Sodium/potassium-transporting ATPase subunit beta | ATP1B4 | 0.96 |
| A0A452E102_CAPHI | FHA domain-containing protein | SLMAP | 0.95 |
| A0A452FI28_CAPHI | Tubulin-specific chaperone A | TBCA | 0.95 |
| A0A452E8X0_CAPHI | Annexin | ANXA3 | 0.94 |
| A0A452EN00_CAPHI | Tropomyosin beta chain | TPM2 | 0.94 |
| A0A452EQ10_CAPHI | PDZ domain-containing protein | AHNAK | 0.93 |
| A0A452EUX2_CAPHI | Peroxiredoxin-6 | PRDX6 | 0.93 |
| A0A452FKE9_CAPHI | Acetyl-coenzyme A synthetase | ACSS1 | 0.93 |
| C6ZP52_CAPHI | II alpha globin | HBAII | 0.93 |
| A0A452F4D3_CAPHI | Cystatin domain-containing protein | LOC102186806 | 0.92 |
| A0A452G117_CAPHI | NADH dehydrogenase [ubiquinone] iron-sulfur protein 4, mitochondrial | NDUFS4 | 0.92 |
| A0A452FGA5_CAPHI | Phosphorylase kinase | PHKG1 | 0.92 |
| A0A452G6M6_CAPHI | Cytochrome b-c1 complex subunit 9 | LOC102187154 | 0.92 |
| A0A452F1S1_CAPHI | Protein-arginine deiminase | PADI2 | 0.92 |
| A0A452FQM9_CAPHI | Complex I-B12 | NDUFB3 | 0.91 |
| A0A452E834_CAPHI | Spectrin alpha chain, erythrocytic 1 | SPTA1 | 0.90 |
| A0A452ECL8_CAPHI | Long-chain specific acyl-CoA dehydrogenase, mitochondrial | ACADL | 0.90 |
| A0A452DM15_CAPHI | Calnexin | CANX | 0.90 |
| A0A452E8U3_CAPHI | Glycerol-3-phosphate dehydrogenase [NAD(+)] | GPD1L | 0.88 |
| A0A452ECE5_CAPHI | Endoplasmic reticulum resident protein 29 | ERP29 | 0.88 |
| A0A452FNY5_CAPHI | GTP-binding protein SAR1b | SAR1B | 0.87 |
| A0A452DLQ3_CAPHI | Enoyl-CoA hydratase, mitochondrial | ECHS1 | 0.85 |
| A0A452FHF8_CAPHI | Prefoldin subunit 6 | PFDN6 | 0.85 |
| A0A452F929_CAPHI | Sulfotransferase | LOC102183750 | 0.84 |
| A0A452EY56_CAPHI | AAA domain-containing protein | PSMC1 | 0.84 |
| A0A452EAA3_CAPHI | ADP-ribosylation factor-like protein 8B | ARL8B | 0.81 |
| A0A452DRF4_CAPHI | Catechol O-methyltransferase domain-containing protein 1 | COMTD1 | 0.80 |
| A0A452DNI3_CAPHI | C-1-tetrahydrofolate synthase, cytoplasmic | MTHFD1 | 0.80 |
| A0A452FMX3_CAPHI | Plectin | PLEC | 0.80 |
| A0A452FJL5_CAPHI | Calcyclin-binding protein | CACYBP | 0.79 |
| A0A452DSQ1_CAPHI | S_100 domain-containing protein | S100A13 | 0.79 |
| A0A452ETQ5_CAPHI | SERPIN domain-containing protein | SERPINB9 | 0.78 |
| A0A452F6I3_CAPHI | Phosphoglucomutase-1 | PGM1 | 0.78 |
| A0A452E7C9_CAPHI | Filamin-A | FLNA | 0.78 |
| A0A452FC43_CAPHI | Serine/threonine-protein phosphatase 2A 65 kDa regulatory subunit A beta isoform | PPP2R1B | 0.77 |
| A0A452EQW7_CAPHI | Malate dehydrogenase | MDH1 | 0.77 |
| A0A452FEA0_CAPHI | 5'-AMP-activated protein kinase subunit gamma-1 | PRKAG1 | 0.77 |
| A0A452EBN6_CAPHI | RNA helicase | EIF4A2 | 0.77 |
| A0A452F2L8_CAPHI | Creatine kinase | CKB | 0.76 |
| A0A452FD51_CAPHI | 26S proteasome non-ATPase regulatory subunit 9 | PSMD9 | 0.75 |
| A0A452DR09_CAPHI | Pyridoxal 5'-phosphate synthase | PNPO | 0.75 |
| A0A452EPN2_CAPHI | Myopalladin | MYPN | 0.75 |
| A0A452FTG4_CAPHI | ATP synthase subunit epsilon, mitochondrial | ATP5F1E | 0.74 |
| A0A452FAW7_CAPHI | Sodium/potassium-transporting ATPase subunit alpha | ATP4A | 0.74 |
| A0A452G9A6_CAPHI | Beta-taxilin | TXLNB | 0.73 |
| A0A452ESU7_CAPHI | Heat shock 27 kDa protein | HSPB1 | 0.73 |
| A0A452FTT1_CAPHI | DnaJ homolog subfamily A member 2 | DNAJA2 | 0.72 |
| A0A452FL25_CAPHI | Apolipoprotein E | APOE | 0.72 |
| A0A452F043_CAPHI | UDP-glucose:glycoprotein glucosyltransferase 1 | UGGT1 | 0.72 |
| A0A452EN08_CAPHI | Cytochrome c oxidase subunit 7A1, mitochondrial | COX7A1 | 0.71 |
| A0A452FWP3_CAPHI | 4-alpha-glucanotransferase | AGL | 0.69 |
| A0A452G2Y4_CAPHI | Histidine-rich glycoprotein | HRG | 0.69 |
| A0A452EH52_CAPHI | Sulfotransferase | SULT1C4 | 0.69 |
| A0A452G3E6_CAPHI | Junctophilin | JPH2 | 0.69 |
| A0A452GB29_CAPHI | Calpain inhibitor | CAST | 0.68 |
| A0A452FL74_CAPHI | Phosphorylase b kinase regulatory subunit | PHKA1 | 0.68 |
| A0A452EVP3_CAPHI | SGNH_hydro domain-containing protein | IAH1 | 0.67 |
| A0A452DQZ4_CAPHI | Ras-related GTP-binding protein | RRAGA | 0.66 |
| A0A452DR58_CAPHI | Histone H2A | H2AC6 | 0.66 |
| A0A452DXY8_CAPHI | PDZ and LIM domain protein 5 | PDLIM5 | 0.65 |
| A0A452E1R3_CAPHI | Calcium/calmodulin-dependent protein kinase | CAMK2D | 0.65 |
| A0A452FRI2_CAPHI | Serum paraoxonase/arylesterase 2 | PON2 | 0.64 |
| A0A452F0S3_CAPHI | MPN domain-containing protein | PRPF8 | 0.64 |
| A0A452DUC4_CAPHI | Nucleosome assembly protein 1-like 1 | LOC102176008 | 0.64 |
| A0A452F2I2_CAPHI | Epoxide hydrolase | EPHX1 | 0.64 |
| A0A452FM48_CAPHI | Phosphomannomutase | PMM2 | 0.63 |
| A0A452F0W7_CAPHI | Tropomyosin alpha-3 chain | TPM3 | 0.63 |
| A0A452E0R7_CAPHI | Serine and arginine rich splicing factor 3 | SRSF3 | 0.63 |
| A0A452G1R9_CAPHI | Nucleobindin-2 | NUCB2 | 0.63 |
| A0A452EPL0_CAPHI | Calmegin | CLGN | 0.63 |
| A0A452F973_CAPHI | Cathelicidin-2 | CATHL2 | 0.63 |
| A0A452EFE3_CAPHI | Actin, alpha skeletal muscle | ACTA1 | 0.62 |
| A0A452E5T3_CAPHI | Peptidyl-prolyl cis-trans isomerase D | PPID | 0.62 |
| A0A452G4A1_CAPHI | Flavodoxin_2 domain-containing protein | NQO1 | 0.61 |
| A0A452DLG8_CAPHI | 14 kDa phosphohistidine phosphatase | PHPT1 | 0.61 |
| A0A452G9I1_CAPHI | Cytosolic 5'-nucleotidase 1A | NT5C1A | 0.60 |
| A0A452FKQ2_CAPHI | Protein NDRG4 | NDRG4 | 0.59 |
| A0A452F1P8_CAPHI | TPR_REGION domain-containing protein | ST13 | 0.58 |
| A0A452EHZ0_CAPHI | Protein phosphatase methylesterase 1 | PPME1 | 0.58 |
| A0A452G7V5_CAPHI | Protein disulfide-isomerase | P4HB | 0.58 |
| A0A452EWA5_CAPHI | Elongation factor 1-alpha | EEF1A2 | 0.57 |
| A0A452F8Y4_CAPHI | Caveolin | CAV1 | 0.57 |
| A0A452GB70_CAPHI | Thioredoxin domain-containing protein | ERP44 | 0.57 |
| A0A452DXH7_CAPHI | Pyridoxal kinase | PDXK | 0.57 |
| A0A452DMC5_CAPHI | Hsp90 chaperone protein kinase-targeting subunit | CDC37 | 0.57 |
| A0A452EB53_CAPHI | Protein-serine/threonine kinase | PDK1 | 0.56 |
| A0A452EMT6_CAPHI | 2'-deoxynucleoside 5'-phosphate N-hydrolase 1 | DNPH1 | 0.56 |
| A0A452FZU7_CAPHI | Fibrinogen C-terminal domain-containing protein | MFAP4 | 0.56 |
| A0A452DYA9_CAPHI | Succinate dehydrogenase cytochrome b560 subunit, mitochondrial | SDHC | 0.56 |
| A0A452FTN8_CAPHI | Importin N-terminal domain-containing protein | TNPO1 | 0.56 |
| A0A452ECX7_CAPHI | 14-3-3 protein theta | YWHAQ | 0.55 |
| A0A452EGV3_CAPHI | Tryptophanyl-tRNA synthetase | WARS2 | 0.55 |
| A0A452DVS7_CAPHI | Neuropilin | NRP1 | 0.54 |
| A0A452EJN6_CAPHI | Alpha(B)-crystallin | CRYAB | 0.54 |
| A0A452FY20_CAPHI | Calcium-binding protein 39 | CAB39 | 0.54 |
| A0A452EBK8_CAPHI | Ras-related protein Rab-11A | RAB11A | 0.53 |
| A0A452DLH9_CAPHI | UBC core domain-containing protein | UBE2L3 | 0.53 |
| A0A452E2L2_CAPHI | HIT domain-containing protein | HINT2 | 0.52 |
| A0A452FIK7_CAPHI | Lipoma-preferred partner | LPP | 0.51 |
| A0A452G8K6_CAPHI | Cardiomyopathy-associated protein 5 | CMYA5 | 0.51 |
| A0A452FDJ0_CAPHI | Nudix hydrolase domain-containing protein | NUDT9 | 0.51 |
| A0A452F874_CAPHI | Kinesin-like protein | KIF5B | 0.51 |
| A0A452FJG6_CAPHI | COP9 signalosome complex subunit 2 | COPS2 | 0.51 |
| A0A452EVI4_CAPHI | Mitogen-activated protein kinase | MAPK12 | 0.51 |
| A0A452G3X4_CAPHI | Rhodanese domain-containing protein | TSTD1 | 0.51 |
| A0A452G8P5_CAPHI | Medium-chain acyl-CoA ligase ACSF2, mitochondrial | ACSF2 | 0.49 |
| A0A452DW40_CAPHI | Inorganic diphosphatase | PPA2 | 0.49 |
| A0A452G9H8_CAPHI | Microsomal prostaglandin E synthase 2 | PTGES2 | 0.47 |
| A0A452DWB5_CAPHI | Inorganic diphosphatase | PPA2 | 0.47 |
| A0A452E480_CAPHI | L-lactate dehydrogenase | LDHB | 0.47 |
| A0A452FS81_CAPHI | Methylmalonyl-CoA isomerase | MMUT | 0.45 |
| A0A452EUI1_CAPHI | 5-formyltetrahydrofolate cyclo-ligase | LOC102172393 | 0.44 |
| A0A452EU44_CAPHI | 14_3_3 domain-containing protein | YWHAH | 0.44 |
| A0A452FK08_CAPHI | NAD(P)-bd_dom domain-containing protein | BLVRB | 0.43 |
| A0A452EWE8_CAPHI | Calcium-activated neutral proteinase small subunit | CAPNS1 | 0.43 |
| A0A452FNL4_CAPHI | Ras-related protein Rab-11B | RAB11B | 0.43 |
| A0A452DS03_CAPHI | ADP-ribosylation factor 4 | LOC108633910 | 0.43 |
| A0A452F2C6_CAPHI | FERM domain-containing protein | EZR | 0.42 |
| A0A452DPJ7_CAPHI | Fumarate hydratase, mitochondrial | FH | 0.42 |
| A0A452EWG8_CAPHI | Insulin-degrading enzyme | IDE | 0.42 |
| A0A452DM03_CAPHI | Mitogen-activated protein kinase | MAPK14 | 0.42 |
| A0A452E9Y1_CAPHI | Pyr_redox_2 domain-containing protein | SQOR | 0.42 |
| A0A452FNW0_CAPHI | Elongation factor 1-gamma | EEF1G | 0.42 |
| A0A452EIW8_CAPHI | Peptidase_M3 domain-containing protein | THOP1 | 0.41 |
| A0A452ET82_CAPHI | Pyruvate kinase | PKM | 0.41 |
| A0A452EU77_CAPHI | Pyruvate kinase | PKM | 0.41 |
| A0A452DYV8_CAPHI | Ig-like domain-containing protein | AZGP1 | 0.41 |
| A0A452G133_CAPHI | 1,4-alpha-glucan branching enzyme | GBE1 | 0.40 |
| A0A452FRH5_CAPHI | Electron transfer flavoprotein-ubiquinone oxidoreductase | ETFDH | 0.40 |
| A0A452DZ71_CAPHI | Glyoxalase domain-containing protein 4 | GLOD4 | 0.40 |
| A0A452EC09_CAPHI | Ubiquitin carboxyl-terminal hydrolase | USP5 | 0.39 |
| A0A452E2E2_CAPHI | Ras GTPase-activating-like protein IQGAP1 | IQGAP1 | 0.39 |
| A0A452FZ73_CAPHI | Creatine kinase | CKM | 0.39 |
| A0A452G5W4_CAPHI | Alpha-1,4 glucan phosphorylase | PYGB | 0.36 |
| A0A452E633_CAPHI | ADP-ribosylation factor 1 | ARF1 | 0.36 |
| A0A452DW82_CAPHI | PCI domain-containing protein | PSMD11 | 0.36 |
| A0A452E1G9_CAPHI | HATPase_c domain-containing protein | HSP90B1 | 0.35 |
| A0A452E6Q6_CAPHI | ATP-dependent RNA helicase DDX1 | DDX1 | 0.33 |
| A0A452G0B4_CAPHI | Cytochrome b5 heme-binding domain-containing protein | NENF | 0.32 |
| A0A452F4B6_CAPHI | 15S Mg(2+)-ATPase p97 subunit | VCP | 0.32 |
| A0A452EVG2_CAPHI | Annexin | ANXA6 | 0.31 |
| A0A452E1P4_CAPHI | Guanine nucleotide-binding protein G(i) subunit alpha-2 | GNAI2 | -0.25 |
| A0A452DLE1_CAPHI | Glutamate dehydrogenase (NAD(P)(+)) | GLUD1 | -0.26 |
| A0A452G2Q0_CAPHI | AP-2 complex subunit alpha | AP2A2 | -0.27 |
| A0A452FB11_CAPHI | Proteasome subunit alpha type | PSMA1 | -0.27 |
| A0A452G4G0_CAPHI | Protein kinase domain-containing protein | NRBP1 | -0.28 |
| A0A452EHP3_CAPHI | Actin-related protein 2/3 complex subunit | ARPC1A | -0.31 |
| B9VR86_CAPHI | Proteasome subunit alpha type | PSMA6 | -0.31 |
| A0A452EJE6_CAPHI | Haloacid dehalogenase-like hydrolase domain-containing 5 | HDHD5 | -0.33 |
| A0A452FWA5_CAPHI | Heterogeneous nuclear ribonucleoprotein R | HNRNPR | -0.34 |
| A0A452E0U0_CAPHI | Tetraspanin | CD81 | -0.34 |
| A0A452E813_CAPHI | Proteasome subunit beta | PSMB5 | -0.34 |
| A0A452FSB8_CAPHI | Methylcrotonoyl-CoA carboxylase beta chain, mitochondrial | MCCC2 | -0.36 |
| A0A452EWR7_CAPHI | Proteasome subunit alpha type | PSMA3 | -0.37 |
| A0A452FZ57_CAPHI | Ectonucleotide pyrophosphatase/phosphodiesterase family member 1 | ENPP1 | -0.39 |
| A0A452EHE9_CAPHI | Spectrin alpha chain, non-erythrocytic 1 | SPTAN1 | -0.39 |
| A0A452EHD5_CAPHI | Spectrin alpha chain, non-erythrocytic 1 | SPTAN1 | -0.39 |
| A0A452G3I1_CAPHI | Delta(3,5)-Delta(2,4)-dienoyl-CoA isomerase, mitochondrial | ECH1 | -0.39 |
| A0A452FGC9_CAPHI | Macrophage-capping protein | CAPG | -0.39 |
| A0A452ELK2_CAPHI | Alpha-centractin | ACTR1A | -0.40 |
| A0A452GAM6_CAPHI | 28S ribosomal protein S22, mitochondrial | MRPS22 | -0.41 |
| A0A452DVV1_CAPHI | Cytosol aminopeptidase | LAP3 | -0.41 |
| A0A452F311_CAPHI | Decorin | DCN | -0.42 |
| A0A452FPG6_CAPHI | acidPPc domain-containing protein | PLPP3 | -0.43 |
| A0A452E8X2_CAPHI | ADP-ribose glycohydrolase ARH3 | ADPRS | -0.44 |
| A0A452F0W6_CAPHI | Aminomethyltransferase | AMT | -0.44 |
| A0A452E6Z8_CAPHI | E3 ubiquitin-protein ligase CHIP | STUB1 | -0.46 |
| A0A452FD94_CAPHI | VOC domain-containing protein | MCEE | -0.46 |
| A0A452G1G3_CAPHI | Helix-destabilizing protein | HNRNPA1 | -0.46 |
| A0A452DZ70_CAPHI | Fructose-bisphosphate aldolase | ALDOC | -0.47 |
| A0A452DMR1_CAPHI | Ras-related protein Rab-7a | RAB7A | -0.47 |
| A0A452FMV1_CAPHI | Heterogeneous nuclear ribonucleoprotein D0 | HNRNPD | -0.48 |
| A0A452EI11_CAPHI | t-SNARE coiled-coil homology domain-containing protein | STX4 | -0.48 |
| A0A452GA47_CAPHI | Cytokeratin-1 | KRT1 | -0.49 |
| A0A452DLZ9_CAPHI | Heme-binding protein 1 | HEBP1 | -0.49 |
| A0A452EXY9_CAPHI | Serine/arginine-rich splicing factor 2 | SRSF2 | -0.50 |
| A0A452EML6_CAPHI | IF rod domain-containing protein | KRT75 | -0.50 |
| A0A452EXP6_CAPHI | 60S ribosomal protein L30 | RPL30 | -0.51 |
| A0A452ECN5_CAPHI | Peptidyl-prolyl cis-trans isomerase E | PPIE | -0.51 |
| A0A452G6Z9_CAPHI | Junction plakoglobin | JUP | -0.52 |
| A0A452FFD7_CAPHI | Complement factor H | CFH | -0.52 |
| A0A452DNF8_CAPHI | 40S ribosomal protein S15a | LOC102173911 | -0.53 |
| A0A452F211_CAPHI | (3R)-3-hydroxyacyl-CoA dehydrogenase | HSD17B8 | -0.53 |
| A0A452E729_CAPHI | Spectrin beta chain | SPTB | -0.53 |
| A0A452EHJ6_CAPHI | Hydroxyacyl-coenzyme A dehydrogenase, mitochondrial | HADH | -0.54 |
| A0A452FWU9_CAPHI | 40S ribosomal protein S16 | RPS16 | -0.54 |
| A0A452F540_CAPHI | High mobility group protein B1 | HMGB1 | -0.54 |
| A0A452FS00_CAPHI | Eukaryotic translation initiation factor 3 subunit K | EIF3K | -0.55 |
| A0A452FZ04_CAPHI | Apolipoprotein H | APOH | -0.55 |
| A0A452G8J5_CAPHI | Septin-2 | SEPTIN2 | -0.55 |
| A0A452G5F8_CAPHI | Dynamin GTPase | DNM2 | -0.55 |
| A0A452FVB0_CAPHI | Actin-related protein 2/3 complex subunit 3 | ARPC3 | -0.56 |
| TRFL_CAPHI | Lactotransferrin | LTF | -0.56 |
| A0A452DQ55_CAPHI | Protein-synthesizing GTPase | EIF2S3 | -0.57 |
| A0A452EQN7_CAPHI | CSD_1 domain-containing protein | CARHSP1 | -0.57 |
| A0A452G0J5_CAPHI | RNA-binding protein 39 | RBM39 | -0.57 |
| A0A452EKF4_CAPHI | Malic enzyme | ME3 | -0.58 |
| A0A452F7G3_CAPHI | Mammalian ependymin-related protein 1 | EPDR1 | -0.58 |
| A0A452EAS1_CAPHI | Guanine nucleotide-binding protein subunit alpha-11 | GNA11 | -0.58 |
| A0A452F9B9_CAPHI | Post-proline cleaving enzyme | PREP | -0.59 |
| A0A452DQU9_CAPHI | Utrophin | UTRN | -0.59 |
| A0A452FP83_CAPHI | RRM domain-containing protein | HNRNPC | -0.60 |
| A0A452FQK5_CAPHI | Macrophage migration inhibitory factor | MIF | -0.62 |
| A0A452FAG1_CAPHI | Laminin subunit alpha-4 | LAMA4 | -0.62 |
| A0A452E6M1_CAPHI | Protein-serine/threonine kinase | PDK3 | -0.62 |
| A0A452EH63_CAPHI | Fructose-bisphosphate aldolase | ALDOA | -0.63 |
| A0A452E6R4_CAPHI | Hydrolase_4 domain-containing protein | MGLL | -0.63 |
| A0A452EQ71_CAPHI | Keratin, type I cytoskeletal 25 | KRT25 | -0.64 |
| A0A452G7N6_CAPHI | Protein S100 | S100A16 | -0.64 |
| A0A452FX42_CAPHI | 40S ribosomal protein S9 | RPS9 | -0.64 |
| A0A452DVL8_CAPHI | Poly(rC)-binding protein 1 | PCBP1 | -0.64 |
| A0A452E9N0_CAPHI | PHB domain-containing protein | STOM | -0.65 |
| A0A452EMZ1_CAPHI | 60S ribosomal protein L36 | RPL36 | -0.66 |
| A0A452ERH0_CAPHI | GST class-pi | LOC102184041 | -0.67 |
| A0A452E0H9_CAPHI | Acid ceramidase | ASAH1 | -0.68 |
| A0A452DR46_CAPHI | 60S ribosomal protein L23 | LOC102169182 | -0.68 |
| A0A452G2P0_CAPHI | WD_REPEATS_REGION domain-containing protein | DYNC1I2 | -0.69 |
| A0A452FER9_CAPHI | IF rod domain-containing protein | KRT7 | -0.70 |
| D2KMJ8_CAPHI | Cytochrome c oxidase subunit 2 | MT-CO2 | -0.71 |
| A0A452DTM3_CAPHI | Biglycan | BGN | -0.72 |
| A0A452E4U4_CAPHI | Ecto-5'-nucleotidase | NT5E | -0.75 |
| A0A452F3I2_CAPHI | IF rod domain-containing protein | KRT12 | -0.76 |
| A0A452EG01_CAPHI | KAT8 regulatory NSL complex subunit 3 | KANSL3 | -0.77 |
| A0A452DQG8_CAPHI | RNA-binding protein FUS | FUS | -0.77 |
| A0A452DYP4_CAPHI | Tetraspanin | CD9 | -0.77 |
| A0A452FDR2_CAPHI | 40S ribosomal protein S8 | RPS8 | -0.79 |
| A0A452G5M5_CAPHI | Palmdelphin | PALMD | -0.80 |
| A0A452G6U6_CAPHI | EF-hand domain-containing protein | TESC | -0.83 |
| A0A452F6V7_CAPHI | Adipsin | CFD | -0.83 |
| A0A452FIX8_CAPHI | Early endosome antigen 1 | EEA1 | -0.85 |
| A0A452DR44_CAPHI | A_deaminase domain-containing protein | ADAL | -0.85 |
| CASA1_CAPHI | Alpha-S1-casein | CSN1S1 | -0.86 |
| A0A452G220_CAPHI | C-terminal-binding protein 1 | CTBP1 | -0.87 |
| A0A452FYK1_CAPHI | Prolow-density lipoprotein receptor-related protein 1 | LRP1 | -0.87 |
| CASA2_CAPHI | Alpha-S2-casein | CSN1S2 | -0.87 |
| A0A452ET13_CAPHI | 39S ribosomal protein L46, mitochondrial | MRPL46 | -0.89 |
| A0A452G969_CAPHI | Sodium/potassium-transporting ATPase subunit alpha | ATP1A2 | -0.91 |
| A0A452EVX4_CAPHI | CCT-alpha | TCP1 | -0.91 |
| A0A452DYE7_CAPHI | Protein kinase domain-containing protein | TNK1 | -0.94 |
| A0A452EN12_CAPHI | S-methyl-5'-thioadenosine phosphorylase | MTAP | -0.94 |
| A0A452E5I0_CAPHI | H15 domain-containing protein | H1-3 | -0.95 |
| A0A452G9D8_CAPHI | WD repeat-containing protein 81 | WDR81 | -1.00 |
| IL6_CAPHI | Interleukin-6 | IL6 | -1.04 |
| A0A452DTU9_CAPHI | Complement factor I | CFI | -1.04 |
| A0A452G600_CAPHI | RNA helicase | DDX6 | -1.04 |
| A0A452G0P6_CAPHI | IF rod domain-containing protein | KRT10 | -1.05 |
| A0A452G3J7_CAPHI | Tubulin beta chain | TUBB2A | -1.07 |
| A0A452ECG5_CAPHI | IF rod domain-containing protein | LOC102176997 | -1.07 |
| A0A452E7D3_CAPHI | CTD domain-containing protein | SRRM2 | -1.07 |
| A0A452DNV7_CAPHI | Acidic leucine-rich nuclear phosphoprotein 32 family member E | ANP32E | -1.08 |
| A0A452GBN0_CAPHI | Golgin subfamily B member 1 | GOLGB1 | -1.11 |
| A0A452FN18_CAPHI | IF rod domain-containing protein | KRT3 | -1.13 |
| A0A452E0A4_CAPHI | Tropomyosin alpha-4 chain | TPM4 | -1.14 |
| A0A452F3X0_CAPHI | Myosin-7B | MYH7B | -1.15 |
| IL12A_CAPHI | Interleukin-12 subunit alpha | IL12A | -1.15 |
| A0A452FY19_CAPHI | OCIA domain-containing protein | OCIAD1 | -1.19 |
| A0A452G2U2_CAPHI | IF rod domain-containing protein | LOC108635997 | -1.20 |
| A0A452EW11_CAPHI | Complement component C7 | C7 | -1.21 |
| A0A452FTY4_CAPHI | Serum amyloid A protein | LOC102168979 | -1.22 |
| A0A452EME0_CAPHI | IF rod domain-containing protein | KRT14 | -1.27 |
| A0A452FLB4_CAPHI | Arginase | ARG1 | -1.46 |
| A0A452FJC1_CAPHI | Coronin | CORO1A | -1.47 |
| A0A452E0Z2_CAPHI | Glyceraldehyde-3-phosphate dehydrogenase | GAPDHS | -1.61 |
| A0A452FJP9_CAPHI | U3 small nucleolar RNA-associated protein 25 homolog | UTP25 | -1.66 |
| A0A452EUG4_CAPHI | Dihydropyrimidine dehydrogenase [NADP(+)] | DPYD | -1.70 |
| A0A452EGA4_CAPHI | Lamin-B2 | LMNB2 | -1.71 |
| A0A452FX49_CAPHI | Hemoglobin subunit beta-C | HBBC | -1.76 |
| A0A452E2E4_CAPHI | TCTP domain-containing protein | LOC102168398 | -1.88 |
| A0A452DJR4_CAPHI | SWIM-type domain-containing protein | ZSWIM8 | -1.94 |
| A0A452DY79_CAPHI | Structural maintenance of chromosomes protein | SMC1A | -2.02 |
| A0A452DRC3_CAPHI | Mitochondrial import inner membrane translocase subunit TIM16 | PAM16 | -4.15 |

Table 2: The most significantly regulated protein list obtained from TL fraction of proteomics analysis.
