## Supplementary material for "Compartmentalisation proteomics revealed endolysosomal protein network changes in a goat model of atrial fibrillation": Supplementary File 2.docx

### Supplementary File 2 contains;

### 1. Table 1: The complete list of pathways obtained from integrated omics analysis of EL, TL proteomics and transcriptomics.

### 2. Table 2: The complete list of most significantly regulated Reactome pathways identified from the EL proteomics.

### 3. Table 3: The complete list of most significantly regulated Reactome pathways identified from the TL proteomics.

### 4. Table 4: The complete list of most significantly regulated Reactome pathways identified from the transcriptomics.

| Pathway ID | Pathway Name | EL enrichment score | TL enrichment score | Transcripts enrichment score | EL vs TL qvalue | EL vs Transcripts qvalue | TL vs Transcripts qvalue |
| --- | --- | --- | --- | --- | --- | --- | --- |
| R-HSA-1280215 | Cytokine Signaling in Immune system | 0.11 | -0.06 | -0.26 | 0.74 | 0.10 | 0.13 |
| R-HSA-162582 | Signal Transduction | -0.02 | -0.01 | -0.13 | 0.74 | 0.36 | 0.38 |
| R-HSA-166016 | Toll Like Receptor 4 (TLR4) Cascade | -0.05 | -0.14 | -0.37 | 0.74 | 0.00 | 0.00 |
| R-HSA-166058 | MyD88:MAL(TIRAP) cascade initiated on plasma membrane | 0.00 | -0.18 | -0.39 | 0.74 | 0.00 | 0.00 |
| R-HSA-166166 | MyD88-independent TLR4 cascade | 0.05 | -0.25 | -0.42 | 0.71 | 0.00 | 0.00 |
| R-HSA-166520 | Signaling by NTRKs | -0.03 | -0.13 | -0.36 | 0.74 | 0.00 | 0.00 |
| R-HSA-167044 | Signalling to RAS | -0.34 | -0.83 | -0.43 | 0.00 | 0.00 | 0.00 |
| R-HSA-168138 | Toll Like Receptor 9 (TLR9) Cascade | 0.17 | -0.13 | -0.41 | 0.73 | 0.00 | 0.00 |
| R-HSA-168142 | Toll Like Receptor 10 (TLR10) Cascade | 0.08 | -0.23 | -0.41 | 0.72 | 0.00 | 0.00 |
| R-HSA-168164 | Toll Like Receptor 3 (TLR3) Cascade | 0.05 | -0.25 | -0.42 | 0.71 | 0.00 | 0.00 |
| R-HSA-168176 | Toll Like Receptor 5 (TLR5) Cascade | 0.08 | -0.23 | -0.41 | 0.72 | 0.00 | 0.00 |
| R-HSA-168179 | Toll Like Receptor TLR1:TLR2 Cascade | 0.00 | -0.18 | -0.36 | 0.74 | 0.00 | 0.00 |
| R-HSA-168181 | Toll Like Receptor 7/8 (TLR7/8) Cascade | 0.08 | -0.23 | -0.42 | 0.72 | 0.00 | 0.00 |
| R-HSA-168188 | Toll Like Receptor TLR6:TLR2 Cascade | 0.00 | -0.18 | -0.39 | 0.74 | 0.00 | 0.00 |
| R-HSA-168249 | Innate Immune System | 0.10 | 0.01 | -0.26 | 0.74 | 0.11 | 0.12 |
| R-HSA-168256 | Immune System | 0.09 | 0.00 | -0.29 | 0.74 | 0.06 | 0.07 |
| R-HSA-168898 | Toll-like Receptor Cascades | 0.02 | -0.14 | -0.34 | 0.74 | 0.01 | 0.01 |
| R-HSA-171007 | p38MAPK events | -0.31 | -0.85 | -0.40 | 0.00 | 0.00 | 0.00 |
| R-HSA-181438 | Toll Like Receptor 2 (TLR2) Cascade | 0.00 | -0.18 | -0.36 | 0.74 | 0.00 | 0.00 |
| R-HSA-187037 | Signaling by NTRK1 (TRKA) | -0.04 | -0.17 | -0.34 | 0.74 | 0.01 | 0.01 |
| R-HSA-187687 | Signalling to ERKs | -0.19 | -0.46 | -0.44 | 0.52 | 0.00 | 0.00 |
| R-HSA-194138 | Signaling by VEGF | -0.13 | -0.49 | -0.41 | 0.45 | 0.00 | 0.00 |
| R-HSA-2262752 | Cellular responses to stress | 0.00 | 0.00 | -0.18 | 0.74 | 0.30 | 0.32 |
| R-HSA-2559580 | Oxidative Stress Induced Senescence | -0.02 | -0.45 | -0.06 | 0.53 | 0.38 | 0.22 |
| R-HSA-2559583 | Cellular Senescence | -0.09 | -0.08 | -0.15 | 0.74 | 0.34 | 0.36 |
| R-HSA-4420097 | VEGFA-VEGFR2 Pathway | -0.19 | -0.47 | -0.43 | 0.50 | 0.00 | 0.00 |
| R-HSA-448424 | Interleukin-17 signaling | 0.14 | -0.20 | -0.39 | 0.71 | 0.00 | 0.00 |
| R-HSA-449147 | Signaling by Interleukins | 0.14 | -0.02 | -0.22 | 0.74 | 0.19 | 0.23 |
| R-HSA-450294 | MAP kinase activation | 0.14 | -0.20 | -0.44 | 0.71 | 0.00 | 0.00 |
| R-HSA-450302 | activated TAK1 mediates p38 MAPK activation | -0.25 | -0.29 | -0.44 | 0.69 | 0.00 | 0.00 |
| R-HSA-8953897 | Cellular responses to stimuli | 0.00 | 0.00 | -0.18 | 0.74 | 0.31 | 0.32 |
| R-HSA-9006934 | Signaling by Receptor Tyrosine Kinases | -0.06 | -0.09 | -0.33 | 0.74 | 0.01 | 0.02 |
| R-HSA-937061 | TRIF(TICAM1)-mediated TLR4 signaling | 0.05 | -0.25 | -0.42 | 0.71 | 0.00 | 0.00 |
| R-HSA-975138 | TRAF6 mediated induction of NFkB and MAP kinases upon TLR7/8 or 9 activation | 0.08 | -0.23 | -0.42 | 0.72 | 0.00 | 0.00 |
| R-HSA-975155 | MyD88 dependent cascade initiated on endosome | 0.08 | -0.23 | -0.42 | 0.72 | 0.00 | 0.00 |
| R-HSA-975871 | MyD88 cascade initiated on plasma membrane | 0.08 | -0.23 | -0.41 | 0.72 | 0.00 | 0.00 |
| R-HSA-1430728 | Metabolism | 0.11 | -0.05 | -0.27 | 0.74 | 0.08 | 0.11 |
| R-HSA-1643685 | Disease | 0.04 | -0.03 | -0.23 | 0.74 | 0.20 | 0.22 |
| R-HSA-211859 | Biological oxidations | 0.08 | -0.12 | -0.17 | 0.74 | 0.32 | 0.33 |
| R-HSA-211945 | Phase I - Functionalization of compounds | -0.15 | -0.22 | -0.19 | 0.73 | 0.26 | 0.26 |
| R-HSA-217271 | FMO oxidises nucleophiles | 0.67 | 0.02 | -0.45 | 0.02 | 0.00 | 0.00 |
| R-HSA-5579029 | Metabolic disorders of biological oxidation enzymes | -0.07 | 0.20 | -0.15 | 0.73 | 0.34 | 0.31 |
| R-HSA-5668914 | Diseases of metabolism | -0.06 | 0.03 | -0.33 | 0.74 | 0.02 | 0.01 |
| R-HSA-389661 | Glyoxylate metabolism and glycine degradation | 0.13 | -0.20 | -0.28 | 0.71 | 0.06 | 0.07 |
| R-HSA-71291 | Metabolism of amino acids and derivatives | 0.12 | 0.07 | -0.10 | 0.74 | 0.37 | 0.39 |
| R-HSA-194315 | Signaling by Rho GTPases | -0.06 | -0.07 | 0.05 | 0.74 | 0.38 | 0.40 |
| R-HSA-73863 | RNA Polymerase I Transcription Termination | 0.93 | 1.00 | -0.17 | 0.00 | 0.00 | 0.00 |
| R-HSA-73864 | RNA Polymerase I Transcription | -0.02 | -0.26 | 0.02 | 0.72 | 0.38 | 0.37 |
| R-HSA-74160 | Gene expression (Transcription) | 0.02 | -0.09 | -0.30 | 0.74 | 0.04 | 0.04 |
| R-HSA-8980692 | RHOA GTPase cycle | -0.10 | 0.00 | -0.49 | 0.74 | 0.00 | 0.00 |
| R-HSA-9012999 | RHO GTPase cycle | -0.10 | -0.04 | 0.13 | 0.74 | 0.35 | 0.37 |
| R-HSA-9013026 | RHOB GTPase cycle | 0.19 | 0.14 | -0.58 | 0.73 | 0.00 | 0.00 |
| R-HSA-9013106 | RHOC GTPase cycle | 0.10 | -0.04 | -0.52 | 0.74 | 0.00 | 0.00 |
| R-HSA-9716542 | Signaling by Rho GTPases, Miro GTPases and RHOBTB3 | -0.06 | -0.07 | 0.04 | 0.74 | 0.38 | 0.40 |
| R-HSA-109582 | Hemostasis | 0.05 | 0.10 | -0.24 | 0.74 | 0.18 | 0.16 |
| R-HSA-114508 | Effects of PIP2 hydrolysis | -0.85 | 0.53 | -0.26 | 0.00 | 0.00 | 0.00 |
| R-HSA-372790 | Signaling by GPCR | 0.15 | 0.12 | -0.09 | 0.74 | 0.37 | 0.39 |
| R-HSA-388396 | GPCR downstream signalling | 0.15 | 0.12 | -0.09 | 0.74 | 0.37 | 0.38 |
| R-HSA-416476 | G alpha (q) signalling events | 0.15 | 0.23 | -0.03 | 0.73 | 0.38 | 0.38 |
| R-HSA-76002 | Platelet activation, signaling and aggregation | 0.05 | 0.12 | -0.19 | 0.74 | 0.29 | 0.27 |
| R-HSA-5663202 | Diseases of signal transduction by growth factor receptors and second messengers | 0.02 | -0.15 | -0.30 | 0.74 | 0.05 | 0.05 |
| R-HSA-6802952 | Signaling by BRAF and RAF1 fusions | -0.21 | -0.20 | -0.38 | 0.72 | 0.00 | 0.00 |
| R-HSA-6802957 | Oncogenic MAPK signaling | -0.20 | -0.21 | -0.38 | 0.72 | 0.00 | 0.00 |
| R-HSA-381038 | XBP1(S) activates chaperone genes | -0.23 | 0.00 | -0.38 | 0.73 | 0.00 | 0.00 |
| R-HSA-381070 | IRE1alpha activates chaperones | -0.32 | 0.07 | -0.37 | 0.66 | 0.00 | 0.00 |
| R-HSA-381119 | Unfolded Protein Response (UPR) | -0.09 | 0.10 | -0.29 | 0.74 | 0.06 | 0.05 |
| R-HSA-381426 | Regulation of Insulin-like Growth Factor (IGF) transport and uptake by Insulin-like Growth Factor Binding Proteins (IGFBPs) | 0.01 | -0.17 | -0.17 | 0.74 | 0.32 | 0.31 |
| R-HSA-392499 | Metabolism of proteins | 0.02 | 0.01 | -0.26 | 0.74 | 0.12 | 0.13 |
| R-HSA-597592 | Post-translational protein modification | 0.03 | -0.07 | -0.34 | 0.74 | 0.01 | 0.01 |
| R-HSA-8957275 | Post-translational protein phosphorylation | -0.02 | -0.22 | -0.22 | 0.73 | 0.22 | 0.20 |
| R-HSA-70171 | Glycolysis | 0.29 | -0.08 | -0.44 | 0.68 | 0.00 | 0.00 |
| R-HSA-70263 | Gluconeogenesis | 0.06 | 0.00 | -0.26 | 0.74 | 0.12 | 0.13 |
| R-HSA-70326 | Glucose metabolism | 0.20 | -0.03 | -0.42 | 0.73 | 0.00 | 0.00 |
| R-HSA-71387 | Metabolism of carbohydrates | 0.13 | -0.13 | -0.33 | 0.73 | 0.01 | 0.02 |
| R-HSA-1483206 | Glycerophospholipid biosynthesis | 0.14 | -0.03 | -0.27 | 0.74 | 0.08 | 0.11 |
| R-HSA-1483257 | Phospholipid metabolism | 0.11 | -0.03 | -0.36 | 0.74 | 0.00 | 0.00 |
| R-HSA-556833 | Metabolism of lipids | 0.18 | 0.01 | -0.33 | 0.74 | 0.01 | 0.02 |
| R-HSA-6798695 | Neutrophil degranulation | 0.09 | -0.02 | -0.26 | 0.74 | 0.12 | 0.13 |
| R-HSA-1428517 | The citric acid (TCA) cycle and respiratory electron transport | -0.14 | -0.13 | -0.28 | 0.74 | 0.07 | 0.08 |
| R-HSA-204174 | Regulation of pyruvate dehydrogenase (PDH) complex | 0.30 | 0.09 | -0.54 | 0.71 | 0.00 | 0.00 |
| R-HSA-5362517 | Signaling by Retinoic Acid | 0.24 | 0.15 | -0.23 | 0.72 | 0.11 | 0.14 |
| R-HSA-70268 | Pyruvate metabolism | 0.17 | 0.00 | -0.44 | 0.74 | 0.00 | 0.00 |
| R-HSA-71406 | Pyruvate metabolism and Citric Acid (TCA) cycle | 0.31 | -0.03 | -0.48 | 0.68 | 0.00 | 0.00 |
| R-HSA-9006931 | Signaling by Nuclear Receptors | 0.04 | 0.03 | -0.18 | 0.74 | 0.31 | 0.32 |
| R-HSA-390522 | Striated Muscle Contraction | -0.35 | -0.20 | -0.33 | 0.66 | 0.00 | 0.01 |
| R-HSA-397014 | Muscle contraction | -0.20 | -0.17 | -0.25 | 0.73 | 0.10 | 0.14 |
| R-HSA-445355 | Smooth Muscle Contraction | -0.12 | -0.20 | -0.21 | 0.73 | 0.24 | 0.24 |
| R-HSA-163200 | Respiratory electron transport, ATP synthesis by chemiosmotic coupling, and heat production by uncoupling proteins. | -0.36 | -0.19 | -0.20 | 0.66 | 0.11 | 0.26 |
| R-HSA-611105 | Respiratory electron transport | -0.36 | -0.21 | -0.20 | 0.65 | 0.10 | 0.24 |
| R-HSA-6799198 | Complex I biogenesis | -0.56 | -0.24 | -0.24 | 0.24 | 0.00 | 0.15 |
| R-HSA-1614517 | Sulfide oxidation to sulfate | 0.44 | 0.68 | -0.10 | 0.01 | 0.14 | 0.00 |
| R-HSA-1614558 | Degradation of cysteine and homocysteine | 0.00 | 0.05 | -0.37 | 0.74 | 0.00 | 0.00 |
| R-HSA-1614635 | Sulfur amino acid metabolism | 0.01 | 0.11 | -0.26 | 0.74 | 0.12 | 0.09 |
| R-HSA-202733 | Cell surface interactions at the vascular wall | -0.25 | 0.18 | -0.31 | 0.66 | 0.02 | 0.01 |
| R-HSA-210991 | Basigin interactions | -0.48 | -0.56 | -0.13 | 0.06 | 0.09 | 0.05 |
| R-HSA-382551 | Transport of small molecules | 0.00 | -0.05 | -0.16 | 0.74 | 0.33 | 0.35 |
| R-HSA-5576891 | Cardiac conduction | -0.16 | -0.11 | -0.23 | 0.74 | 0.16 | 0.20 |
| R-HSA-5578775 | Ion homeostasis | -0.13 | -0.07 | -0.44 | 0.74 | 0.00 | 0.00 |
| R-HSA-5663205 | Infectious disease | 0.11 | 0.03 | -0.16 | 0.74 | 0.32 | 0.35 |
| R-HSA-936837 | Ion transport by P-type ATPases | -0.49 | -0.08 | -0.35 | 0.43 | 0.00 | 0.01 |
| R-HSA-9679191 | Potential therapeutics for SARS | 0.16 | -0.02 | -0.33 | 0.74 | 0.01 | 0.02 |
| R-HSA-9679506 | SARS-CoV Infections | 0.00 | -0.19 | -0.27 | 0.74 | 0.11 | 0.10 |
| R-HSA-983712 | Ion channel transport | -0.31 | -0.05 | -0.24 | 0.70 | 0.08 | 0.20 |
| R-HSA-1280218 | Adaptive Immune System | 0.01 | 0.02 | -0.35 | 0.74 | 0.01 | 0.01 |
| R-HSA-199991 | Membrane Trafficking | 0.06 | -0.07 | -0.38 | 0.74 | 0.00 | 0.00 |
| R-HSA-2132295 | MHC class II antigen presentation | 0.09 | 0.16 | -0.27 | 0.74 | 0.08 | 0.05 |
| R-HSA-5653656 | Vesicle-mediated transport | 0.06 | -0.04 | -0.38 | 0.74 | 0.00 | 0.00 |
| R-HSA-8854214 | TBC/RABGAPs | 0.17 | -0.32 | -0.40 | 0.60 | 0.00 | 0.00 |
| R-HSA-8873719 | RAB geranylgeranylation | -0.14 | 0.07 | -0.32 | 0.74 | 0.02 | 0.02 |
| R-HSA-8876198 | RAB GEFs exchange GTP for GDP on RABs | 0.16 | 0.07 | -0.48 | 0.74 | 0.00 | 0.00 |
| R-HSA-9007101 | Rab regulation of trafficking | 0.10 | -0.11 | -0.46 | 0.74 | 0.00 | 0.00 |
| R-HSA-9013148 | CDC42 GTPase cycle | 0.07 | 0.17 | -0.44 | 0.74 | 0.00 | 0.00 |
| R-HSA-9013149 | RAC1 GTPase cycle | 0.17 | 0.13 | -0.44 | 0.74 | 0.00 | 0.00 |
| R-HSA-9013404 | RAC2 GTPase cycle | 0.05 | -0.01 | -0.44 | 0.74 | 0.00 | 0.00 |
| R-HSA-9013405 | RHOD GTPase cycle | -0.26 | -0.10 | -0.51 | 0.72 | 0.00 | 0.00 |
| R-HSA-9013406 | RHOQ GTPase cycle | -0.04 | 0.21 | -0.44 | 0.73 | 0.00 | 0.00 |
| R-HSA-9013407 | RHOH GTPase cycle | 0.21 | -0.01 | -0.39 | 0.73 | 0.00 | 0.00 |
| R-HSA-9013408 | RHOG GTPase cycle | 0.04 | 0.09 | -0.43 | 0.74 | 0.00 | 0.00 |
| R-HSA-9013409 | RHOJ GTPase cycle | 0.09 | 0.12 | -0.41 | 0.74 | 0.00 | 0.00 |
| R-HSA-9013423 | RAC3 GTPase cycle | 0.20 | 0.13 | -0.41 | 0.73 | 0.00 | 0.00 |
| R-HSA-9035034 | RHOF GTPase cycle | -0.10 | -0.11 | -0.55 | 0.74 | 0.00 | 0.00 |
| R-HSA-9635486 | Infection with Mycobacterium tuberculosis | 0.19 | -0.05 | -0.32 | 0.73 | 0.01 | 0.02 |
| R-HSA-9636383 | Prevention of phagosomal-lysosomal fusion | 0.12 | 0.73 | -0.14 | 0.01 | 0.34 | 0.00 |
| R-HSA-9636569 | Suppression of autophagy | 1.00 | 0.98 | -0.60 | 0.00 | 0.00 | 0.00 |
| R-HSA-9637687 | Suppression of phagosomal maturation | 0.21 | 0.31 | -0.29 | 0.69 | 0.03 | 0.01 |
| R-HSA-9637690 | Response of Mtb to phagocytosis | 0.28 | -0.10 | -0.24 | 0.68 | 0.08 | 0.19 |
| R-HSA-1632852 | Macroautophagy | 0.05 | 0.01 | -0.29 | 0.74 | 0.05 | 0.05 |
| R-HSA-163560 | Triglyceride catabolism | 0.09 | -0.06 | -0.13 | 0.74 | 0.35 | 0.38 |
| R-HSA-6811440 | Retrograde transport at the Trans-Golgi-Network | -0.18 | -0.40 | -0.49 | 0.62 | 0.00 | 0.00 |
| R-HSA-6811442 | Intra-Golgi and retrograde Golgi-to-ER traffic | 0.11 | 0.03 | -0.37 | 0.74 | 0.00 | 0.00 |
| R-HSA-8979227 | Triglyceride metabolism | 0.09 | -0.06 | -0.03 | 0.74 | 0.38 | 0.40 |
| R-HSA-9612973 | Autophagy | 0.05 | 0.01 | -0.29 | 0.74 | 0.06 | 0.06 |
| R-HSA-9613354 | Lipophagy | -0.10 | -0.21 | -0.59 | 0.73 | 0.00 | 0.00 |
| R-HSA-9613829 | Chaperone Mediated Autophagy | 0.08 | -0.05 | -0.12 | 0.74 | 0.36 | 0.38 |
| R-HSA-9615710 | Late endosomal microautophagy | 0.20 | 0.29 | -0.16 | 0.70 | 0.28 | 0.21 |
| R-HSA-9663891 | Selective autophagy | 0.04 | 0.07 | -0.29 | 0.74 | 0.07 | 0.06 |
| R-HSA-1237044 | Erythrocytes take up carbon dioxide and release oxygen | 0.17 | -0.08 | 0.31 | 0.73 | 0.02 | 0.02 |
| R-HSA-1247673 | Erythrocytes take up oxygen and release carbon dioxide | 0.17 | -0.08 | 0.55 | 0.73 | 0.00 | 0.00 |
| R-HSA-1475029 | Reversible hydration of carbon dioxide | 0.62 | 0.29 | 0.31 | 0.07 | 0.00 | 0.02 |
| R-HSA-1480926 | O2/CO2 exchange in erythrocytes | 0.17 | -0.08 | 0.31 | 0.73 | 0.02 | 0.02 |
| R-HSA-199992 | trans-Golgi Network Vesicle Budding | -0.19 | 0.05 | -0.34 | 0.73 | 0.01 | 0.01 |
| R-HSA-432722 | Golgi Associated Vesicle Biogenesis | -0.09 | 0.04 | -0.39 | 0.74 | 0.00 | 0.00 |
| R-HSA-1266738 | Developmental Biology | 0.04 | 0.10 | -0.15 | 0.74 | 0.34 | 0.34 |
| R-HSA-156827 | L13a-mediated translational silencing of Ceruloplasmin expression | 0.03 | 0.34 | 0.12 | 0.68 | 0.37 | 0.29 |
| R-HSA-156842 | Eukaryotic Translation Elongation | -0.05 | 0.25 | 0.18 | 0.71 | 0.30 | 0.26 |
| R-HSA-156902 | Peptide chain elongation | -0.05 | 0.33 | 0.18 | 0.66 | 0.31 | 0.21 |
| R-HSA-168255 | Influenza Infection | -0.01 | 0.20 | -0.04 | 0.73 | 0.38 | 0.39 |
| R-HSA-168273 | Influenza Viral RNA Transcription and Replication | 0.00 | 0.29 | 0.00 | 0.71 | 0.38 | 0.36 |
| R-HSA-1799339 | SRP-dependent cotranslational protein targeting to membrane | -0.05 | 0.30 | 0.12 | 0.69 | 0.36 | 0.32 |
| R-HSA-192823 | Viral mRNA Translation | -0.05 | 0.35 | 0.17 | 0.65 | 0.32 | 0.22 |
| R-HSA-2408522 | Selenoamino acid metabolism | 0.01 | 0.30 | 0.08 | 0.70 | 0.38 | 0.35 |
| R-HSA-2408557 | Selenocysteine synthesis | -0.05 | 0.35 | 0.16 | 0.64 | 0.34 | 0.23 |
| R-HSA-376176 | Signaling by ROBO receptors | 0.07 | 0.09 | -0.04 | 0.74 | 0.38 | 0.40 |
| R-HSA-422475 | Axon guidance | 0.05 | 0.02 | -0.21 | 0.74 | 0.24 | 0.25 |
| R-HSA-6791226 | Major pathway of rRNA processing in the nucleolus and cytosol | -0.04 | 0.32 | 0.02 | 0.68 | 0.38 | 0.35 |
| R-HSA-72312 | rRNA processing | -0.04 | 0.33 | 0.00 | 0.67 | 0.38 | 0.34 |
| R-HSA-72613 | Eukaryotic Translation Initiation | 0.03 | 0.34 | 0.09 | 0.68 | 0.38 | 0.31 |
| R-HSA-72689 | Formation of a pool of free 40S subunits | 0.01 | 0.36 | 0.14 | 0.66 | 0.35 | 0.25 |
| R-HSA-72706 | GTP hydrolysis and joining of the 60S ribosomal subunit | 0.01 | 0.34 | 0.11 | 0.68 | 0.37 | 0.30 |
| R-HSA-72737 | Cap-dependent Translation Initiation | 0.03 | 0.34 | 0.09 | 0.68 | 0.38 | 0.31 |
| R-HSA-72764 | Eukaryotic Translation Termination | -0.04 | 0.32 | 0.16 | 0.68 | 0.33 | 0.26 |
| R-HSA-72766 | Translation | 0.08 | 0.20 | -0.07 | 0.74 | 0.38 | 0.38 |
| R-HSA-8868773 | rRNA processing in the nucleus and cytosol | -0.04 | 0.32 | 0.02 | 0.68 | 0.38 | 0.35 |
| R-HSA-8953854 | Metabolism of RNA | 0.01 | 0.13 | 0.18 | 0.74 | 0.31 | 0.32 |
| R-HSA-9010553 | Regulation of expression of SLITs and ROBOs | 0.04 | 0.14 | 0.05 | 0.74 | 0.38 | 0.40 |
| R-HSA-927802 | Nonsense-Mediated Decay (NMD) | -0.05 | 0.30 | 0.11 | 0.69 | 0.37 | 0.33 |
| R-HSA-9633012 | Response of EIF2AK4 (GCN2) to amino acid deficiency | -0.02 | 0.35 | 0.14 | 0.66 | 0.36 | 0.26 |
| R-HSA-9675108 | Nervous system development | 0.04 | 0.03 | -0.22 | 0.74 | 0.23 | 0.24 |
| R-HSA-9711097 | Cellular response to starvation | -0.03 | 0.26 | 0.00 | 0.71 | 0.38 | 0.38 |
| R-HSA-975956 | Nonsense Mediated Decay (NMD) independent of the Exon Junction Complex (EJC) | -0.06 | 0.32 | 0.15 | 0.67 | 0.34 | 0.27 |
| R-HSA-975957 | Nonsense Mediated Decay (NMD) enhanced by the Exon Junction Complex (EJC) | -0.05 | 0.30 | 0.11 | 0.69 | 0.37 | 0.33 |
| R-HSA-2142753 | Arachidonic acid metabolism | 0.14 | -0.06 | -0.17 | 0.74 | 0.30 | 0.34 |
| R-HSA-2162123 | Synthesis of Prostaglandins (PG) and Thromboxanes (TX) | 0.35 | -0.12 | -0.19 | 0.59 | 0.10 | 0.29 |
| R-HSA-8978868 | Fatty acid metabolism | 0.23 | 0.03 | -0.33 | 0.73 | 0.00 | 0.01 |
| R-HSA-112314 | Neurotransmitter receptors and postsynaptic signal transmission | 0.22 | 0.01 | -0.13 | 0.73 | 0.31 | 0.37 |
| R-HSA-112315 | Transmission across Chemical Synapses | 0.14 | 0.01 | -0.13 | 0.74 | 0.34 | 0.38 |
| R-HSA-112316 | Neuronal System | 0.12 | 0.06 | -0.09 | 0.74 | 0.37 | 0.39 |
| R-HSA-114608 | Platelet degranulation | 0.04 | 0.02 | -0.13 | 0.74 | 0.36 | 0.38 |
| R-HSA-1445148 | Translocation of SLC2A4 (GLUT4) to the plasma membrane | -0.03 | -0.19 | -0.41 | 0.74 | 0.00 | 0.00 |
| R-HSA-157858 | Gap junction trafficking and regulation | -0.13 | 0.16 | -0.09 | 0.73 | 0.37 | 0.38 |
| R-HSA-1640170 | Cell Cycle | 0.01 | -0.04 | -0.29 | 0.74 | 0.06 | 0.06 |
| R-HSA-1852241 | Organelle biogenesis and maintenance | -0.09 | -0.03 | -0.43 | 0.74 | 0.00 | 0.00 |
| R-HSA-190828 | Gap junction trafficking | -0.13 | 0.10 | -0.05 | 0.74 | 0.38 | 0.40 |
| R-HSA-190840 | Microtubule-dependent trafficking of connexons from Golgi to the plasma membrane | 0.11 | 0.12 | -0.06 | 0.74 | 0.38 | 0.40 |
| R-HSA-190861 | Gap junction assembly | 0.11 | 0.12 | 0.01 | 0.74 | 0.38 | 0.40 |
| R-HSA-190872 | Transport of connexons to the plasma membrane | 0.11 | 0.12 | -0.02 | 0.74 | 0.38 | 0.40 |
| R-HSA-195258 | RHO GTPase Effectors | -0.03 | -0.11 | -0.21 | 0.74 | 0.24 | 0.25 |
| R-HSA-199977 | ER to Golgi Anterograde Transport | 0.01 | 0.15 | -0.33 | 0.74 | 0.01 | 0.01 |
| R-HSA-2467813 | Separation of Sister Chromatids | 0.08 | 0.05 | -0.25 | 0.74 | 0.15 | 0.15 |
| R-HSA-2500257 | Resolution of Sister Chromatid Cohesion | 0.02 | 0.23 | -0.30 | 0.73 | 0.04 | 0.01 |
| R-HSA-2555396 | Mitotic Metaphase and Anaphase | 0.03 | 0.02 | -0.26 | 0.74 | 0.13 | 0.13 |
| R-HSA-2565942 | Regulation of PLK1 Activity at G2/M Transition | -0.05 | -0.03 | -0.38 | 0.74 | 0.00 | 0.00 |
| R-HSA-2995410 | Nuclear Envelope (NE) Reassembly | 0.05 | 0.02 | -0.29 | 0.74 | 0.05 | 0.05 |
| R-HSA-3371497 | HSP90 chaperone cycle for steroid hormone receptors (SHR) in the presence of ligand | -0.02 | 0.10 | -0.19 | 0.74 | 0.29 | 0.27 |
| R-HSA-373760 | L1CAM interactions | 0.06 | 0.06 | -0.30 | 0.74 | 0.04 | 0.04 |
| R-HSA-380259 | Loss of Nlp from mitotic centrosomes | 0.03 | -0.06 | -0.40 | 0.74 | 0.00 | 0.00 |
| R-HSA-380270 | Recruitment of mitotic centrosome proteins and complexes | 0.03 | -0.06 | -0.42 | 0.74 | 0.00 | 0.00 |
| R-HSA-380284 | Loss of proteins required for interphase microtubule organization from the centrosome | 0.03 | -0.06 | -0.40 | 0.74 | 0.00 | 0.00 |
| R-HSA-380287 | Centrosome maturation | 0.03 | -0.06 | -0.42 | 0.74 | 0.00 | 0.00 |
| R-HSA-380320 | Recruitment of NuMA to mitotic centrosomes | 0.14 | 0.00 | -0.38 | 0.74 | 0.00 | 0.00 |
| R-HSA-389957 | Prefoldin mediated transfer of substrate to CCT/TriC | -0.06 | 0.04 | -0.07 | 0.74 | 0.38 | 0.40 |
| R-HSA-389958 | Cooperation of Prefoldin and TriC/CCT in actin and tubulin folding | -0.04 | 0.02 | -0.06 | 0.74 | 0.38 | 0.40 |
| R-HSA-389960 | Formation of tubulin folding intermediates by CCT/TriC | 0.02 | 0.08 | -0.08 | 0.74 | 0.38 | 0.40 |
| R-HSA-389977 | Post-chaperonin tubulin folding pathway | 0.03 | 0.00 | -0.09 | 0.74 | 0.38 | 0.40 |
| R-HSA-390466 | Chaperonin-mediated protein folding | 0.04 | 0.18 | -0.15 | 0.74 | 0.35 | 0.32 |
| R-HSA-391251 | Protein folding | -0.01 | 0.14 | -0.16 | 0.74 | 0.34 | 0.32 |
| R-HSA-437239 | Recycling pathway of L1 | 0.20 | 0.11 | -0.26 | 0.73 | 0.08 | 0.11 |
| R-HSA-438064 | Post NMDA receptor activation events | 0.09 | -0.21 | -0.26 | 0.72 | 0.12 | 0.12 |
| R-HSA-442755 | Activation of NMDA receptors and postsynaptic events | 0.09 | -0.21 | -0.28 | 0.72 | 0.08 | 0.08 |
| R-HSA-446203 | Asparagine N-linked glycosylation | -0.01 | 0.00 | -0.31 | 0.74 | 0.03 | 0.03 |
| R-HSA-453274 | Mitotic G2-G2/M phases | 0.07 | -0.05 | -0.31 | 0.74 | 0.03 | 0.03 |
| R-HSA-5358351 | Signaling by Hedgehog | 0.17 | -0.05 | -0.29 | 0.74 | 0.04 | 0.06 |
| R-HSA-5610787 | Hedgehog 'off' state | 0.15 | -0.03 | -0.26 | 0.74 | 0.09 | 0.12 |
| R-HSA-5617833 | Cilium Assembly | 0.06 | 0.00 | -0.43 | 0.74 | 0.00 | 0.00 |
| R-HSA-5620912 | Anchoring of the basal body to the plasma membrane | 0.03 | -0.06 | -0.44 | 0.74 | 0.00 | 0.00 |
| R-HSA-5620924 | Intraflagellar transport | 0.01 | 0.12 | -0.32 | 0.74 | 0.03 | 0.02 |
| R-HSA-5626467 | RHO GTPases activate IQGAPs | 0.00 | -0.01 | -0.15 | 0.74 | 0.34 | 0.36 |
| R-HSA-5663220 | RHO GTPases Activate Formins | 0.07 | 0.19 | -0.29 | 0.74 | 0.06 | 0.03 |
| R-HSA-6807878 | COPI-mediated anterograde transport | 0.05 | 0.19 | -0.31 | 0.74 | 0.04 | 0.01 |
| R-HSA-6811434 | COPI-dependent Golgi-to-ER retrograde traffic | 0.11 | -0.10 | -0.30 | 0.74 | 0.04 | 0.05 |
| R-HSA-6811436 | COPI-independent Golgi-to-ER retrograde traffic | 0.18 | 0.21 | -0.30 | 0.73 | 0.03 | 0.01 |
| R-HSA-68877 | Mitotic Prometaphase | 0.10 | 0.08 | -0.37 | 0.74 | 0.00 | 0.00 |
| R-HSA-68882 | Mitotic Anaphase | 0.03 | 0.02 | -0.26 | 0.74 | 0.13 | 0.13 |
| R-HSA-68886 | M Phase | 0.05 | -0.03 | -0.26 | 0.74 | 0.12 | 0.13 |
| R-HSA-69275 | G2/M Transition | 0.07 | -0.05 | -0.32 | 0.74 | 0.03 | 0.03 |
| R-HSA-69278 | Cell Cycle, Mitotic | 0.04 | -0.03 | -0.29 | 0.74 | 0.07 | 0.07 |
| R-HSA-76005 | Response to elevated platelet cytosolic Ca2+ | 0.03 | 0.03 | -0.14 | 0.74 | 0.35 | 0.37 |
| R-HSA-8852276 | The role of GTSE1 in G2/M progression after G2 checkpoint | 0.10 | -0.03 | -0.12 | 0.74 | 0.36 | 0.39 |
| R-HSA-8854518 | AURKA Activation by TPX2 | 0.03 | -0.06 | -0.41 | 0.74 | 0.00 | 0.00 |
| R-HSA-8856688 | Golgi-to-ER retrograde transport | 0.12 | 0.05 | -0.33 | 0.74 | 0.01 | 0.01 |
| R-HSA-8955332 | Carboxyterminal post-translational modifications of tubulin | 0.33 | 0.04 | -0.20 | 0.68 | 0.10 | 0.26 |
| R-HSA-948021 | Transport to the Golgi and subsequent modification | 0.02 | 0.16 | -0.32 | 0.74 | 0.02 | 0.01 |
| R-HSA-9609646 | HCMV Infection | 0.13 | 0.18 | -0.15 | 0.74 | 0.33 | 0.31 |
| R-HSA-9609690 | HCMV Early Events | 0.05 | 0.21 | -0.13 | 0.73 | 0.36 | 0.33 |
| R-HSA-9609736 | Assembly and cell surface presentation of NMDA receptors | 0.13 | -0.03 | -0.17 | 0.74 | 0.31 | 0.34 |
| R-HSA-9619483 | Activation of AMPK downstream of NMDARs | 0.11 | -0.03 | -0.32 | 0.74 | 0.02 | 0.02 |
| R-HSA-9646399 | Aggrephagy | 0.05 | 0.11 | -0.14 | 0.74 | 0.36 | 0.36 |
| R-HSA-9648025 | EML4 and NUDC in mitotic spindle formation | 0.04 | 0.26 | -0.30 | 0.72 | 0.04 | 0.01 |
| R-HSA-9668328 | Sealing of the nuclear envelope (NE) by ESCRT-III | 0.17 | 0.11 | -0.15 | 0.74 | 0.31 | 0.34 |
| R-HSA-983189 | Kinesins | 0.18 | 0.00 | -0.24 | 0.74 | 0.12 | 0.18 |
| R-HSA-983231 | Factors involved in megakaryocyte development and platelet production | 0.11 | 0.02 | -0.25 | 0.74 | 0.12 | 0.15 |
| R-HSA-109581 | Apoptosis | 0.06 | -0.07 | -0.22 | 0.74 | 0.21 | 0.23 |
| R-HSA-1168372 | Downstream signaling events of B Cell Receptor (BCR) | 0.06 | -0.03 | -0.23 | 0.74 | 0.20 | 0.22 |
| R-HSA-1169091 | Activation of NF-kappaB in B cells | 0.11 | -0.04 | -0.20 | 0.74 | 0.25 | 0.29 |
| R-HSA-1234174 | Cellular response to hypoxia | 0.07 | -0.09 | -0.22 | 0.74 | 0.22 | 0.24 |
| R-HSA-1234176 | Oxygen-dependent proline hydroxylation of Hypoxia-inducible Factor Alpha | 0.10 | -0.08 | -0.19 | 0.74 | 0.27 | 0.30 |
| R-HSA-1236974 | ER-Phagosome pathway | 0.08 | -0.04 | -0.22 | 0.74 | 0.21 | 0.24 |
| R-HSA-1236975 | Antigen processing-Cross presentation | 0.07 | -0.04 | -0.24 | 0.74 | 0.18 | 0.20 |
| R-HSA-1236978 | Cross-presentation of soluble exogenous antigens (endosomes) | 0.12 | -0.05 | -0.13 | 0.74 | 0.35 | 0.38 |
| R-HSA-1257604 | PIP3 activates AKT signaling | 0.15 | -0.04 | -0.29 | 0.74 | 0.05 | 0.07 |
| R-HSA-157118 | Signaling by NOTCH | 0.01 | -0.06 | -0.18 | 0.74 | 0.30 | 0.32 |
| R-HSA-162906 | HIV Infection | 0.13 | -0.04 | -0.27 | 0.74 | 0.08 | 0.11 |
| R-HSA-162909 | Host Interactions of HIV factors | 0.12 | -0.03 | -0.28 | 0.74 | 0.07 | 0.09 |
| R-HSA-169911 | Regulation of Apoptosis | 0.09 | -0.05 | -0.15 | 0.74 | 0.34 | 0.36 |
| R-HSA-174084 | Autodegradation of Cdh1 by Cdh1:APC/C | 0.11 | -0.04 | -0.10 | 0.74 | 0.37 | 0.39 |
| R-HSA-174113 | SCF-beta-TrCP mediated degradation of Emi1 | 0.11 | -0.04 | -0.10 | 0.74 | 0.37 | 0.39 |
| R-HSA-174143 | APC/C-mediated degradation of cell cycle proteins | 0.11 | -0.04 | -0.12 | 0.74 | 0.36 | 0.39 |
| R-HSA-174154 | APC/C:Cdc20 mediated degradation of Securin | 0.11 | -0.04 | -0.10 | 0.74 | 0.37 | 0.39 |
| R-HSA-174178 | APC/C:Cdh1 mediated degradation of Cdc20 and other APC/C:Cdh1 targeted proteins in late mitosis/early G1 | 0.11 | -0.04 | -0.10 | 0.74 | 0.37 | 0.39 |
| R-HSA-174184 | Cdc20:Phospho-APC/C mediated degradation of Cyclin A | 0.11 | -0.04 | -0.12 | 0.74 | 0.36 | 0.38 |
| R-HSA-176408 | Regulation of APC/C activators between G1/S and early anaphase | 0.11 | -0.04 | -0.11 | 0.74 | 0.37 | 0.39 |
| R-HSA-176409 | APC/C:Cdc20 mediated degradation of mitotic proteins | 0.11 | -0.04 | -0.11 | 0.74 | 0.37 | 0.39 |
| R-HSA-176814 | Activation of APC/C and APC/C:Cdc20 mediated degradation of mitotic proteins | 0.11 | -0.04 | -0.12 | 0.74 | 0.36 | 0.39 |
| R-HSA-179419 | APC:Cdc20 mediated degradation of cell cycle proteins prior to satisfation of the cell cycle checkpoint | 0.11 | -0.04 | -0.13 | 0.74 | 0.35 | 0.38 |
| R-HSA-180534 | Vpu mediated degradation of CD4 | 0.11 | -0.04 | -0.13 | 0.74 | 0.35 | 0.38 |
| R-HSA-180585 | Vif-mediated degradation of APOBEC3G | 0.10 | -0.08 | -0.15 | 0.74 | 0.34 | 0.36 |
| R-HSA-187577 | SCF(Skp2)-mediated degradation of p27/p21 | 0.11 | -0.04 | -0.10 | 0.74 | 0.37 | 0.39 |
| R-HSA-195253 | Degradation of beta-catenin by the destruction complex | 0.09 | -0.06 | -0.14 | 0.74 | 0.35 | 0.37 |
| R-HSA-195721 | Signaling by WNT | 0.01 | -0.03 | -0.16 | 0.74 | 0.33 | 0.35 |
| R-HSA-201681 | TCF dependent signaling in response to WNT | 0.00 | -0.13 | -0.13 | 0.74 | 0.36 | 0.37 |
| R-HSA-202403 | TCR signaling | 0.10 | -0.06 | -0.26 | 0.74 | 0.12 | 0.14 |
| R-HSA-202424 | Downstream TCR signaling | 0.11 | -0.04 | -0.24 | 0.74 | 0.16 | 0.19 |
| R-HSA-211733 | Regulation of activated PAK-2p34 by proteasome mediated degradation | 0.10 | -0.06 | -0.13 | 0.74 | 0.36 | 0.38 |
| R-HSA-212436 | Generic Transcription Pathway | 0.01 | -0.14 | -0.29 | 0.74 | 0.06 | 0.06 |
| R-HSA-2454202 | Fc epsilon receptor (FCERI) signaling | 0.09 | -0.07 | -0.29 | 0.74 | 0.06 | 0.07 |
| R-HSA-2871837 | FCERI mediated NF-kB activation | 0.11 | -0.04 | -0.19 | 0.74 | 0.28 | 0.31 |
| R-HSA-349425 | Autodegradation of the E3 ubiquitin ligase COP1 | 0.11 | -0.04 | -0.15 | 0.74 | 0.33 | 0.36 |
| R-HSA-350562 | Regulation of ornithine decarboxylase (ODC) | 0.11 | -0.08 | -0.12 | 0.74 | 0.36 | 0.38 |
| R-HSA-351202 | Metabolism of polyamines | 0.14 | -0.10 | -0.12 | 0.74 | 0.35 | 0.38 |
| R-HSA-382556 | ABC-family proteins mediated transport | 0.13 | -0.04 | -0.12 | 0.74 | 0.36 | 0.39 |
| R-HSA-3858494 | Beta-catenin independent WNT signaling | 0.07 | 0.01 | -0.21 | 0.74 | 0.24 | 0.26 |
| R-HSA-4086400 | PCP/CE pathway | 0.09 | -0.03 | -0.21 | 0.74 | 0.22 | 0.25 |
| R-HSA-446652 | Interleukin-1 family signaling | 0.14 | -0.03 | -0.20 | 0.74 | 0.24 | 0.29 |
| R-HSA-450408 | AUF1 (hnRNP D0) binds and destabilizes mRNA | 0.09 | -0.05 | -0.10 | 0.74 | 0.37 | 0.39 |
| R-HSA-450531 | Regulation of mRNA stability by proteins that bind AU-rich elements | 0.05 | -0.11 | -0.20 | 0.74 | 0.27 | 0.28 |
| R-HSA-453276 | Regulation of mitotic cell cycle | 0.11 | -0.04 | -0.12 | 0.74 | 0.36 | 0.39 |
| R-HSA-453279 | Mitotic G1 phase and G1/S transition | 0.10 | -0.07 | -0.26 | 0.74 | 0.12 | 0.15 |
| R-HSA-4608870 | Asymmetric localization of PCP proteins | 0.11 | -0.04 | -0.19 | 0.74 | 0.28 | 0.31 |
| R-HSA-4641257 | Degradation of AXIN | 0.11 | -0.04 | -0.16 | 0.74 | 0.32 | 0.35 |
| R-HSA-4641258 | Degradation of DVL | 0.11 | -0.04 | -0.18 | 0.74 | 0.29 | 0.32 |
| R-HSA-5357801 | Programmed Cell Death | 0.07 | -0.07 | -0.24 | 0.74 | 0.15 | 0.17 |
| R-HSA-5358346 | Hedgehog ligand biogenesis | 0.12 | -0.10 | -0.12 | 0.74 | 0.36 | 0.38 |
| R-HSA-5362768 | Hh mutants are degraded by ERAD | 0.11 | -0.07 | -0.14 | 0.74 | 0.34 | 0.37 |
| R-HSA-5387390 | Hh mutants abrogate ligand secretion | 0.11 | -0.07 | -0.12 | 0.74 | 0.36 | 0.38 |
| R-HSA-5607761 | Dectin-1 mediated noncanonical NF-kB signaling | 0.08 | -0.04 | -0.16 | 0.74 | 0.33 | 0.35 |
| R-HSA-5607764 | CLEC7A (Dectin-1) signaling | 0.06 | -0.03 | -0.27 | 0.74 | 0.11 | 0.12 |
| R-HSA-5610780 | Degradation of GLI1 by the proteasome | 0.13 | -0.08 | -0.18 | 0.74 | 0.29 | 0.33 |
| R-HSA-5610783 | Degradation of GLI2 by the proteasome | 0.13 | -0.08 | -0.12 | 0.74 | 0.36 | 0.39 |
| R-HSA-5610785 | GLI3 is processed to GLI3R by the proteasome | 0.13 | -0.08 | -0.12 | 0.74 | 0.36 | 0.38 |
| R-HSA-5619084 | ABC transporter disorders | 0.11 | -0.07 | -0.19 | 0.74 | 0.27 | 0.31 |
| R-HSA-5619115 | Disorders of transmembrane transporters | 0.10 | -0.04 | -0.19 | 0.74 | 0.27 | 0.30 |
| R-HSA-5621481 | C-type lectin receptors (CLRs) | 0.04 | -0.06 | -0.26 | 0.74 | 0.12 | 0.13 |
| R-HSA-5632684 | Hedgehog 'on' state | 0.12 | -0.02 | -0.24 | 0.74 | 0.16 | 0.19 |
| R-HSA-5658442 | Regulation of RAS by GAPs | 0.08 | -0.06 | -0.22 | 0.74 | 0.20 | 0.23 |
| R-HSA-5668541 | TNFR2 non-canonical NF-kB pathway | 0.08 | -0.04 | -0.17 | 0.74 | 0.32 | 0.34 |
| R-HSA-5673001 | RAF/MAP kinase cascade | 0.01 | -0.10 | -0.25 | 0.74 | 0.15 | 0.16 |
| R-HSA-5676590 | NIK-->noncanonical NF-kB signaling | 0.08 | -0.04 | -0.16 | 0.74 | 0.33 | 0.35 |
| R-HSA-5678895 | Defective CFTR causes cystic fibrosis | 0.11 | -0.07 | -0.18 | 0.74 | 0.29 | 0.32 |
| R-HSA-5683057 | MAPK family signaling cascades | 0.00 | -0.10 | -0.26 | 0.74 | 0.11 | 0.12 |
| R-HSA-5684996 | MAPK1/MAPK3 signaling | 0.01 | -0.10 | -0.26 | 0.74 | 0.13 | 0.13 |
| R-HSA-5687128 | MAPK6/MAPK4 signaling | 0.08 | -0.06 | -0.20 | 0.74 | 0.26 | 0.29 |
| R-HSA-5688426 | Deubiquitination | -0.02 | -0.10 | -0.28 | 0.74 | 0.08 | 0.08 |
| R-HSA-5689603 | UCH proteinases | 0.06 | -0.08 | -0.13 | 0.74 | 0.36 | 0.38 |
| R-HSA-5689880 | Ub-specific processing proteases | 0.01 | -0.08 | -0.25 | 0.74 | 0.14 | 0.15 |
| R-HSA-6807070 | PTEN Regulation | 0.15 | -0.06 | -0.28 | 0.74 | 0.05 | 0.08 |
| R-HSA-68827 | CDT1 association with the CDC6:ORC:origin complex | 0.11 | -0.04 | -0.15 | 0.74 | 0.33 | 0.36 |
| R-HSA-68867 | Assembly of the pre-replicative complex | 0.11 | -0.04 | -0.18 | 0.74 | 0.30 | 0.33 |
| R-HSA-68949 | Orc1 removal from chromatin | 0.11 | -0.04 | -0.15 | 0.74 | 0.34 | 0.37 |
| R-HSA-69002 | DNA Replication Pre-Initiation | 0.11 | -0.04 | -0.22 | 0.74 | 0.20 | 0.23 |
| R-HSA-69017 | CDK-mediated phosphorylation and removal of Cdc6 | 0.11 | -0.04 | -0.12 | 0.74 | 0.36 | 0.38 |
| R-HSA-69052 | Switching of origins to a post-replicative state | 0.11 | -0.04 | -0.15 | 0.74 | 0.33 | 0.36 |
| R-HSA-69202 | Cyclin E associated events during G1/S transition | 0.11 | -0.04 | -0.21 | 0.74 | 0.23 | 0.27 |
| R-HSA-69206 | G1/S Transition | 0.10 | -0.07 | -0.25 | 0.74 | 0.12 | 0.15 |
| R-HSA-69239 | Synthesis of DNA | 0.11 | -0.04 | -0.17 | 0.74 | 0.30 | 0.33 |
| R-HSA-69242 | S Phase | 0.11 | -0.04 | -0.26 | 0.74 | 0.12 | 0.14 |
| R-HSA-69306 | DNA Replication | 0.11 | -0.04 | -0.18 | 0.74 | 0.29 | 0.32 |
| R-HSA-69481 | G2/M Checkpoints | 0.04 | -0.09 | -0.19 | 0.74 | 0.28 | 0.30 |
| R-HSA-69541 | Stabilization of p53 | 0.11 | -0.04 | -0.14 | 0.74 | 0.34 | 0.37 |
| R-HSA-69563 | p53-Dependent G1 DNA Damage Response | 0.11 | -0.04 | -0.16 | 0.74 | 0.32 | 0.35 |
| R-HSA-69580 | p53-Dependent G1/S DNA damage checkpoint | 0.11 | -0.04 | -0.16 | 0.74 | 0.32 | 0.35 |
| R-HSA-69601 | Ubiquitin Mediated Degradation of Phosphorylated Cdc25A | 0.11 | -0.04 | -0.14 | 0.74 | 0.34 | 0.37 |
| R-HSA-69610 | p53-Independent DNA Damage Response | 0.11 | -0.04 | -0.14 | 0.74 | 0.34 | 0.37 |
| R-HSA-69613 | p53-Independent G1/S DNA damage checkpoint | 0.11 | -0.04 | -0.14 | 0.74 | 0.34 | 0.37 |
| R-HSA-69615 | G1/S DNA Damage Checkpoints | 0.11 | -0.04 | -0.18 | 0.74 | 0.29 | 0.32 |
| R-HSA-69620 | Cell Cycle Checkpoints | 0.01 | -0.01 | -0.23 | 0.74 | 0.19 | 0.20 |
| R-HSA-69656 | Cyclin A:Cdk2-associated events at S phase entry | 0.11 | -0.04 | -0.20 | 0.74 | 0.24 | 0.28 |
| R-HSA-73857 | RNA Polymerase II Transcription | 0.02 | -0.10 | -0.30 | 0.74 | 0.05 | 0.05 |
| R-HSA-75815 | Ubiquitin-dependent degradation of Cyclin D | 0.11 | -0.04 | -0.14 | 0.74 | 0.35 | 0.37 |
| R-HSA-8854050 | FBXL7 down-regulates AURKA during mitotic entry and in early mitosis | 0.11 | -0.04 | -0.13 | 0.74 | 0.35 | 0.38 |
| R-HSA-8878159 | Transcriptional regulation by RUNX3 | 0.04 | -0.05 | -0.24 | 0.74 | 0.16 | 0.18 |
| R-HSA-8878166 | Transcriptional regulation by RUNX2 | 0.14 | 0.04 | -0.22 | 0.74 | 0.20 | 0.23 |
| R-HSA-8878171 | Transcriptional regulation by RUNX1 | 0.01 | -0.02 | -0.19 | 0.74 | 0.30 | 0.31 |
| R-HSA-8932339 | ROS sensing by NFE2L2 | 0.11 | -0.04 | -0.15 | 0.74 | 0.34 | 0.37 |
| R-HSA-8939236 | RUNX1 regulates transcription of genes involved in differentiation of HSCs | 0.03 | -0.05 | 0.02 | 0.74 | 0.38 | 0.40 |
| R-HSA-8939902 | Regulation of RUNX2 expression and activity | 0.13 | 0.03 | -0.17 | 0.74 | 0.30 | 0.33 |
| R-HSA-8941858 | Regulation of RUNX3 expression and activity | 0.08 | -0.02 | -0.14 | 0.74 | 0.34 | 0.37 |
| R-HSA-8948751 | Regulation of PTEN stability and activity | 0.12 | -0.02 | -0.21 | 0.74 | 0.23 | 0.27 |
| R-HSA-8951664 | Neddylation | 0.09 | -0.16 | -0.37 | 0.73 | 0.00 | 0.00 |
| R-HSA-9006925 | Intracellular signaling by second messengers | 0.16 | -0.09 | -0.32 | 0.73 | 0.02 | 0.03 |
| R-HSA-9013694 | Signaling by NOTCH4 | 0.10 | -0.07 | -0.22 | 0.74 | 0.22 | 0.25 |
| R-HSA-9020702 | Interleukin-1 signaling | 0.14 | -0.03 | -0.26 | 0.74 | 0.09 | 0.12 |
| R-HSA-9604323 | Negative regulation of NOTCH4 signaling | 0.10 | -0.07 | -0.14 | 0.74 | 0.34 | 0.37 |
| R-HSA-9707564 | Cytoprotection by HMOX1 | 0.07 | -0.08 | -0.19 | 0.74 | 0.28 | 0.30 |
| R-HSA-9707587 | Regulation of HMOX1 expression and activity | 0.11 | -0.03 | -0.13 | 0.74 | 0.35 | 0.38 |
| R-HSA-9711123 | Cellular response to chemical stress | 0.11 | -0.11 | -0.17 | 0.74 | 0.30 | 0.33 |
| R-HSA-983168 | Antigen processing: Ubiquitination & Proteasome degradation | 0.09 | -0.09 | -0.40 | 0.74 | 0.00 | 0.00 |
| R-HSA-983169 | Class I MHC mediated antigen processing & presentation | 0.04 | -0.07 | -0.39 | 0.74 | 0.00 | 0.00 |
| R-HSA-983705 | Signaling by the B Cell Receptor (BCR) | 0.06 | -0.06 | -0.30 | 0.74 | 0.04 | 0.05 |
| R-HSA-5625740 | RHO GTPases activate PKNs | -0.15 | -0.24 | 0.05 | 0.73 | 0.38 | 0.38 |
| R-HSA-5625900 | RHO GTPases activate CIT | -0.04 | 0.07 | -0.39 | 0.74 | 0.00 | 0.00 |
| R-HSA-5627117 | RHO GTPases Activate ROCKs | -0.01 | 0.09 | -0.46 | 0.74 | 0.00 | 0.00 |
| R-HSA-5627123 | RHO GTPases activate PAKs | -0.05 | 0.05 | -0.36 | 0.74 | 0.00 | 0.00 |
| R-HSA-1500931 | Cell-Cell communication | -0.37 | -0.12 | -0.24 | 0.65 | 0.05 | 0.19 |
| R-HSA-418990 | Adherens junctions interactions | -0.77 | -0.23 | -0.18 | 0.00 | 0.00 | 0.28 |
| R-HSA-421270 | Cell-cell junction organization | -0.77 | -0.23 | -0.07 | 0.00 | 0.00 | 0.38 |
| R-HSA-446728 | Cell junction organization | -0.32 | -0.01 | -0.17 | 0.68 | 0.19 | 0.33 |
| R-HSA-1592230 | Mitochondrial biogenesis | -0.21 | -0.05 | -0.41 | 0.73 | 0.00 | 0.00 |
| R-HSA-163210 | Formation of ATP by chemiosmotic coupling | -0.32 | -0.08 | -0.17 | 0.69 | 0.19 | 0.34 |
| R-HSA-8949613 | Cristae formation | -0.43 | 0.00 | -0.24 | 0.54 | 0.02 | 0.18 |
| R-HSA-3700989 | Transcriptional Regulation by TP53 | -0.04 | -0.21 | -0.29 | 0.73 | 0.06 | 0.05 |
| R-HSA-5628897 | TP53 Regulates Metabolic Genes | -0.04 | -0.27 | -0.29 | 0.72 | 0.07 | 0.04 |
| R-HSA-174824 | Plasma lipoprotein assembly, remodeling, and clearance | 0.05 | -0.19 | -0.19 | 0.73 | 0.29 | 0.28 |
| R-HSA-196854 | Metabolism of vitamins and cofactors | 0.10 | -0.07 | -0.34 | 0.74 | 0.01 | 0.01 |
| R-HSA-2187338 | Visual phototransduction | 0.10 | 0.14 | 0.00 | 0.74 | 0.38 | 0.40 |
| R-HSA-6806667 | Metabolism of fat-soluble vitamins | -0.04 | -0.06 | -0.14 | 0.74 | 0.36 | 0.37 |
| R-HSA-8963888 | Chylomicron assembly | -0.18 | -0.40 | -0.09 | 0.62 | 0.36 | 0.25 |
| R-HSA-8963889 | Assembly of active LPL and LIPC lipase complexes | -0.96 | 0.22 | -0.18 | 0.00 | 0.00 | 0.23 |
| R-HSA-8963898 | Plasma lipoprotein assembly | -0.02 | -0.46 | -0.15 | 0.51 | 0.35 | 0.13 |
| R-HSA-8963899 | Plasma lipoprotein remodeling | -0.23 | -0.22 | -0.09 | 0.72 | 0.35 | 0.38 |
| R-HSA-8963901 | Chylomicron remodeling | -0.29 | -0.28 | 0.07 | 0.67 | 0.32 | 0.35 |
| R-HSA-9709957 | Sensory Perception | 0.11 | 0.18 | 0.28 | 0.74 | 0.07 | 0.07 |
| R-HSA-975634 | Retinoid metabolism and transport | -0.04 | -0.06 | -0.14 | 0.74 | 0.35 | 0.37 |
| R-HSA-977225 | Amyloid fiber formation | -0.35 | -0.12 | 0.08 | 0.67 | 0.27 | 0.39 |
| R-HSA-159236 | Transport of Mature mRNA derived from an Intron-Containing Transcript | 0.29 | 0.53 | -0.37 | 0.30 | 0.00 | 0.00 |
| R-HSA-72163 | mRNA Splicing - Major Pathway | 0.07 | 0.35 | 0.52 | 0.68 | 0.00 | 0.00 |
| R-HSA-72165 | mRNA Splicing - Minor Pathway | 0.02 | 0.19 | -0.20 | 0.74 | 0.27 | 0.21 |
| R-HSA-72172 | mRNA Splicing | 0.07 | 0.35 | 0.51 | 0.68 | 0.00 | 0.00 |
| R-HSA-72187 | mRNA 3'-end processing | 0.19 | 0.51 | -0.28 | 0.40 | 0.05 | 0.00 |
| R-HSA-72202 | Transport of Mature Transcript to Cytoplasm | 0.29 | 0.53 | -0.37 | 0.30 | 0.00 | 0.00 |
| R-HSA-72203 | Processing of Capped Intron-Containing Pre-mRNA | 0.09 | 0.38 | 0.43 | 0.65 | 0.00 | 0.00 |
| R-HSA-73856 | RNA Polymerase II Transcription Termination | 0.26 | 0.55 | -0.22 | 0.27 | 0.12 | 0.00 |
| R-HSA-71288 | Creatine metabolism | -0.12 | -0.08 | -0.18 | 0.74 | 0.30 | 0.32 |
| R-HSA-9696264 | RND3 GTPase cycle | 0.15 | -0.19 | -0.53 | 0.71 | 0.00 | 0.00 |
| R-HSA-9696273 | RND1 GTPase cycle | 0.12 | -0.09 | -0.39 | 0.74 | 0.00 | 0.00 |
| R-HSA-162658 | Golgi Cisternae Pericentriolar Stack Reorganization | 0.12 | 0.15 | -0.41 | 0.74 | 0.00 | 0.00 |
| R-HSA-68875 | Mitotic Prophase | -0.02 | 0.02 | -0.18 | 0.74 | 0.31 | 0.33 |
| R-HSA-1912420 | Pre-NOTCH Processing in Golgi | -0.59 | -0.28 | -0.48 | 0.14 | 0.00 | 0.00 |
| R-HSA-1912422 | Pre-NOTCH Expression and Processing | -0.61 | -0.33 | -0.10 | 0.08 | 0.02 | 0.32 |
| R-HSA-418346 | Platelet homeostasis | 0.09 | 0.14 | -0.22 | 0.74 | 0.21 | 0.18 |
| R-HSA-418359 | Reduction of cytosolic Ca++ levels | -0.46 | 0.03 | -0.37 | 0.45 | 0.00 | 0.00 |
| R-HSA-418360 | Platelet calcium homeostasis | -0.46 | 0.03 | -0.33 | 0.45 | 0.00 | 0.01 |
| R-HSA-8856825 | Cargo recognition for clathrin-mediated endocytosis | -0.12 | -0.23 | -0.33 | 0.73 | 0.01 | 0.01 |
| R-HSA-8856828 | Clathrin-mediated endocytosis | 0.05 | -0.17 | -0.32 | 0.74 | 0.03 | 0.02 |
| R-HSA-70688 | Proline catabolism | -0.82 | -0.73 | -0.03 | 0.00 | 0.00 | 0.00 |
| R-HSA-112310 | Neurotransmitter release cycle | -0.31 | 0.14 | -0.13 | 0.63 | 0.26 | 0.35 |
| R-HSA-210500 | Glutamate Neurotransmitter Release Cycle | -0.91 | -0.02 | -0.27 | 0.00 | 0.00 | 0.11 |
| R-HSA-8964539 | Glutamate and glutamine metabolism | -0.21 | -0.49 | -0.45 | 0.46 | 0.00 | 0.00 |
| R-HSA-1474228 | Degradation of the extracellular matrix | -0.20 | 0.11 | -0.20 | 0.72 | 0.23 | 0.26 |
| R-HSA-1474244 | Extracellular matrix organization | -0.10 | 0.11 | -0.30 | 0.74 | 0.04 | 0.03 |
| R-HSA-216083 | Integrin cell surface interactions | -0.16 | 0.16 | -0.47 | 0.72 | 0.00 | 0.00 |
| R-HSA-425366 | Transport of bile salts and organic acids, metal ions and amine compounds | 0.12 | -0.15 | -0.04 | 0.73 | 0.38 | 0.40 |
| R-HSA-425407 | SLC-mediated transmembrane transport | 0.16 | 0.09 | -0.09 | 0.74 | 0.37 | 0.39 |
| R-HSA-433692 | Proton-coupled monocarboxylate transport | -0.85 | -0.33 | -0.34 | 0.00 | 0.00 | 0.00 |
| R-HSA-5619070 | Defective SLC16A1 causes symptomatic deficiency in lactate transport (SDLT) | -0.85 | -0.33 | -0.43 | 0.00 | 0.00 | 0.00 |
| R-HSA-5619102 | SLC transporter disorders | 0.03 | 0.18 | -0.19 | 0.74 | 0.28 | 0.23 |
| R-HSA-379716 | Cytosolic tRNA aminoacylation | 0.27 | 0.22 | -0.10 | 0.70 | 0.32 | 0.35 |
| R-HSA-379724 | tRNA Aminoacylation | 0.29 | 0.04 | -0.27 | 0.71 | 0.04 | 0.11 |
| R-HSA-141424 | Amplification of signal from the kinetochores | -0.10 | 0.25 | -0.32 | 0.70 | 0.02 | 0.00 |
| R-HSA-141444 | Amplification of signal from unattached kinetochores via a MAD2 inhibitory signal | -0.10 | 0.25 | -0.32 | 0.70 | 0.02 | 0.00 |
| R-HSA-69618 | Mitotic Spindle Checkpoint | -0.10 | 0.25 | -0.29 | 0.70 | 0.06 | 0.02 |
| R-HSA-196780 | Biotin transport and metabolism | 0.51 | 0.36 | -0.50 | 0.26 | 0.00 | 0.00 |
| R-HSA-196849 | Metabolism of water-soluble vitamins and cofactors | 0.13 | 0.00 | -0.44 | 0.74 | 0.00 | 0.00 |
| R-HSA-3296482 | Defects in vitamin and cofactor metabolism | 0.31 | 0.10 | -0.46 | 0.70 | 0.00 | 0.00 |
| R-HSA-3323169 | Defects in biotin (Btn) metabolism | 0.51 | 0.36 | -0.58 | 0.26 | 0.00 | 0.00 |
| R-HSA-3371599 | Defective HLCS causes multiple carboxylase deficiency | 0.51 | 0.36 | -0.60 | 0.26 | 0.00 | 0.00 |
| R-HSA-71032 | Propionyl-CoA catabolism | 0.46 | 0.00 | -0.59 | 0.47 | 0.00 | 0.00 |
| R-HSA-77289 | Mitochondrial Fatty Acid Beta-Oxidation | 0.30 | -0.06 | -0.38 | 0.68 | 0.00 | 0.00 |
| R-HSA-1483166 | Synthesis of PA | 0.36 | 0.10 | -0.29 | 0.67 | 0.01 | 0.05 |
| R-HSA-9659379 | Sensory processing of sound | 0.08 | 0.23 | -0.23 | 0.73 | 0.19 | 0.10 |
| R-HSA-9662360 | Sensory processing of sound by inner hair cells of the cochlea | 0.07 | 0.25 | -0.24 | 0.73 | 0.16 | 0.06 |
| R-HSA-9662361 | Sensory processing of sound by outer hair cells of the cochlea | 0.07 | 0.22 | -0.22 | 0.73 | 0.22 | 0.14 |
| R-HSA-77286 | mitochondrial fatty acid beta-oxidation of saturated fatty acids | 0.42 | -0.12 | -0.31 | 0.46 | 0.00 | 0.04 |
| R-HSA-77310 | Beta oxidation of lauroyl-CoA to decanoyl-CoA-CoA | 0.53 | -0.25 | -0.39 | 0.03 | 0.00 | 0.00 |
| R-HSA-77346 | Beta oxidation of decanoyl-CoA to octanoyl-CoA-CoA | 0.54 | -0.01 | -0.22 | 0.21 | 0.00 | 0.25 |
| R-HSA-77348 | Beta oxidation of octanoyl-CoA to hexanoyl-CoA | 0.63 | 0.05 | -0.24 | 0.06 | 0.00 | 0.18 |
| R-HSA-77350 | Beta oxidation of hexanoyl-CoA to butanoyl-CoA | 0.65 | -0.23 | -0.29 | 0.00 | 0.00 | 0.05 |
| R-HSA-77352 | Beta oxidation of butanoyl-CoA to acetyl-CoA | 0.70 | -0.22 | -0.31 | 0.00 | 0.00 | 0.02 |
| R-HSA-3371453 | Regulation of HSF1-mediated heat shock response | -0.05 | 0.03 | -0.39 | 0.74 | 0.00 | 0.00 |
| R-HSA-3371556 | Cellular response to heat stress | -0.08 | -0.28 | -0.39 | 0.72 | 0.00 | 0.00 |
| R-HSA-9013420 | RHOU GTPase cycle | -0.15 | 0.02 | -0.49 | 0.74 | 0.00 | 0.00 |
| R-HSA-9013424 | RHOV GTPase cycle | -0.17 | 0.01 | -0.44 | 0.74 | 0.00 | 0.00 |
| R-HSA-9696270 | RND2 GTPase cycle | 0.03 | 0.05 | -0.40 | 0.74 | 0.00 | 0.00 |
| R-HSA-6794362 | Protein-protein interactions at synapses | -0.03 | 0.30 | -0.11 | 0.70 | 0.37 | 0.29 |
| R-HSA-8849932 | Synaptic adhesion-like molecules | 0.04 | 0.05 | -0.08 | 0.74 | 0.38 | 0.40 |
| R-HSA-111885 | Opioid Signalling | 0.32 | 0.17 | -0.31 | 0.69 | 0.01 | 0.02 |
| R-HSA-1296041 | Activation of G protein gated Potassium channels | 0.43 | 0.73 | 0.02 | 0.00 | 0.22 | 0.00 |
| R-HSA-1296059 | G protein gated Potassium channels | 0.43 | 0.73 | 0.02 | 0.00 | 0.22 | 0.00 |
| R-HSA-1296065 | Inwardly rectifying K+ channels | 0.43 | 0.73 | -0.03 | 0.00 | 0.22 | 0.00 |
| R-HSA-1296071 | Potassium Channels | 0.43 | 0.73 | 0.05 | 0.00 | 0.21 | 0.00 |
| R-HSA-163359 | Glucagon signaling in metabolic regulation | 0.33 | 0.19 | -0.12 | 0.68 | 0.25 | 0.35 |
| R-HSA-163685 | Integration of energy metabolism | 0.19 | 0.13 | -0.27 | 0.73 | 0.06 | 0.07 |
| R-HSA-202040 | G-protein activation | 0.48 | 0.79 | 0.03 | 0.00 | 0.15 | 0.00 |
| R-HSA-2485179 | Activation of the phototransduction cascade | 0.48 | 0.79 | 0.44 | 0.00 | 0.00 | 0.00 |
| R-HSA-2514856 | The phototransduction cascade | 0.45 | 0.67 | 0.13 | 0.01 | 0.12 | 0.01 |
| R-HSA-2514859 | Inactivation, recovery and regulation of the phototransduction cascade | 0.45 | 0.67 | 0.12 | 0.01 | 0.13 | 0.01 |
| R-HSA-2980736 | Peptide hormone metabolism | 0.04 | -0.02 | 0.04 | 0.74 | 0.38 | 0.40 |
| R-HSA-373080 | Class B/2 (Secretin family receptors) | 0.49 | 0.77 | 0.08 | 0.00 | 0.12 | 0.00 |
| R-HSA-381676 | Glucagon-like Peptide-1 (GLP1) regulates insulin secretion | 0.17 | 0.14 | -0.17 | 0.74 | 0.29 | 0.31 |
| R-HSA-381753 | Olfactory Signaling Pathway | 0.51 | 0.52 | 0.52 | 0.06 | 0.00 | 0.00 |
| R-HSA-381771 | Synthesis, secretion, and inactivation of Glucagon-like Peptide-1 (GLP-1) | 0.03 | 0.34 | 0.28 | 0.68 | 0.07 | 0.03 |
| R-HSA-392170 | ADP signalling through P2Y purinoceptor 12 | 0.52 | 0.79 | 0.00 | 0.00 | 0.11 | 0.00 |
| R-HSA-392451 | G beta:gamma signalling through PI3Kgamma | 0.31 | 0.49 | 0.00 | 0.38 | 0.34 | 0.17 |
| R-HSA-392518 | Signal amplification | 0.51 | 0.59 | -0.12 | 0.02 | 0.06 | 0.01 |
| R-HSA-392851 | Prostacyclin signalling through prostacyclin receptor | 0.49 | 0.77 | 0.06 | 0.00 | 0.13 | 0.00 |
| R-HSA-397795 | G-protein beta:gamma signalling | 0.28 | 0.47 | -0.08 | 0.45 | 0.33 | 0.13 |
| R-HSA-400042 | Adrenaline,noradrenaline inhibits insulin secretion | 0.47 | 0.80 | -0.10 | 0.00 | 0.11 | 0.00 |
| R-HSA-400508 | Incretin synthesis, secretion, and inactivation | 0.03 | 0.34 | 0.28 | 0.68 | 0.08 | 0.03 |
| R-HSA-4086398 | Ca2+ pathway | -0.04 | 0.19 | -0.21 | 0.73 | 0.24 | 0.17 |
| R-HSA-416482 | G alpha (12/13) signalling events | 0.20 | 0.54 | -0.35 | 0.32 | 0.00 | 0.00 |
| R-HSA-418217 | G beta:gamma signalling through PLC beta | 0.43 | 0.73 | 0.03 | 0.00 | 0.22 | 0.00 |
| R-HSA-418555 | G alpha (s) signalling events | 0.44 | 0.70 | -0.01 | 0.01 | 0.21 | 0.01 |
| R-HSA-418592 | ADP signalling through P2Y purinoceptor 1 | 0.44 | 0.48 | -0.04 | 0.21 | 0.19 | 0.15 |
| R-HSA-418594 | G alpha (i) signalling events | 0.31 | 0.16 | -0.10 | 0.70 | 0.29 | 0.38 |
| R-HSA-418597 | G alpha (z) signalling events | 0.55 | 0.81 | -0.13 | 0.00 | 0.03 | 0.00 |
| R-HSA-420092 | Glucagon-type ligand receptors | 0.49 | 0.77 | 0.19 | 0.00 | 0.03 | 0.00 |
| R-HSA-422356 | Regulation of insulin secretion | 0.26 | 0.24 | -0.21 | 0.70 | 0.14 | 0.14 |
| R-HSA-428930 | Thromboxane signalling through TP receptor | 0.48 | 0.79 | -0.01 | 0.00 | 0.15 | 0.00 |
| R-HSA-432040 | Vasopressin regulates renal water homeostasis via Aquaporins | 0.23 | 0.09 | -0.11 | 0.73 | 0.33 | 0.38 |
| R-HSA-445717 | Aquaporin-mediated transport | 0.23 | 0.09 | -0.09 | 0.73 | 0.35 | 0.39 |
| R-HSA-451326 | Activation of kainate receptors upon glutamate binding | 0.43 | 0.73 | -0.02 | 0.00 | 0.22 | 0.00 |
| R-HSA-456926 | Thrombin signalling through proteinase activated receptors (PARs) | 0.54 | 0.48 | -0.04 | 0.07 | 0.08 | 0.15 |
| R-HSA-500657 | Presynaptic function of Kainate receptors | 0.43 | 0.73 | 0.01 | 0.00 | 0.22 | 0.00 |
| R-HSA-500792 | GPCR ligand binding | 0.35 | 0.43 | 0.08 | 0.46 | 0.28 | 0.23 |
| R-HSA-6814122 | Cooperation of PDCL (PhLP1) and TRiC/CCT in G-protein beta folding | 0.07 | 0.23 | -0.03 | 0.73 | 0.38 | 0.38 |
| R-HSA-8939211 | ESR-mediated signaling | -0.09 | -0.07 | -0.16 | 0.74 | 0.33 | 0.35 |
| R-HSA-8964315 | G beta:gamma signalling through BTK | 0.43 | 0.73 | 0.08 | 0.00 | 0.20 | 0.00 |
| R-HSA-8964616 | G beta:gamma signalling through CDC42 | 0.37 | 0.66 | 0.03 | 0.03 | 0.29 | 0.01 |
| R-HSA-9009391 | Extra-nuclear estrogen signaling | 0.12 | 0.02 | -0.19 | 0.74 | 0.27 | 0.31 |
| R-HSA-9658195 | Leishmania infection | 0.28 | -0.01 | -0.16 | 0.71 | 0.23 | 0.35 |
| R-HSA-9660821 | ADORA2B mediated anti-inflammatory cytokines production | 0.39 | 0.31 | 0.02 | 0.56 | 0.27 | 0.36 |
| R-HSA-9662851 | Anti-inflammatory response favouring Leishmania parasite infection | 0.40 | 0.19 | -0.06 | 0.61 | 0.23 | 0.38 |
| R-HSA-9664433 | Leishmania parasite growth and survival | 0.40 | 0.19 | -0.06 | 0.61 | 0.23 | 0.38 |
| R-HSA-977443 | GABA receptor activation | 0.52 | 0.79 | -0.03 | 0.00 | 0.10 | 0.00 |
| R-HSA-977444 | GABA B receptor activation | 0.52 | 0.79 | -0.10 | 0.00 | 0.07 | 0.00 |
| R-HSA-991365 | Activation of GABAB receptors | 0.52 | 0.79 | -0.10 | 0.00 | 0.07 | 0.00 |
| R-HSA-997272 | Inhibition of voltage gated Ca2+ channels via Gbeta/gamma subunits | 0.43 | 0.73 | 0.02 | 0.00 | 0.22 | 0.00 |
| R-HSA-190704 | Oligomerization of connexins into connexons | -0.90 | -0.15 | 0.21 | 0.00 | 0.00 | 0.22 |
| R-HSA-190827 | Transport of connexins along the secretory pathway | -0.90 | -0.15 | 0.21 | 0.00 | 0.00 | 0.22 |
| R-HSA-190873 | Gap junction degradation | -0.62 | 0.03 | -0.26 | 0.04 | 0.00 | 0.13 |
| R-HSA-191650 | Regulation of gap junction activity | -0.54 | 0.30 | -0.81 | 0.01 | 0.00 | 0.00 |
| R-HSA-196025 | Formation of annular gap junctions | -0.62 | 0.03 | -0.23 | 0.04 | 0.00 | 0.21 |
| R-HSA-71384 | Ethanol oxidation | 0.06 | -0.44 | -0.36 | 0.49 | 0.00 | 0.00 |
| R-HSA-1268020 | Mitochondrial protein import | -0.19 | 0.09 | -0.22 | 0.73 | 0.18 | 0.21 |
| R-HSA-71403 | Citric acid cycle (TCA cycle) | 0.42 | -0.04 | -0.53 | 0.54 | 0.00 | 0.00 |
| R-HSA-9609507 | Protein localization | -0.11 | 0.11 | -0.28 | 0.74 | 0.07 | 0.06 |
| R-HSA-75105 | Fatty acyl-CoA biosynthesis | 0.44 | 0.45 | -0.41 | 0.26 | 0.00 | 0.00 |
| R-HSA-2029480 | Fcgamma receptor (FCGR) dependent phagocytosis | 0.31 | -0.24 | -0.40 | 0.52 | 0.00 | 0.00 |
| R-HSA-2029482 | Regulation of actin dynamics for phagocytic cup formation | 0.28 | -0.23 | -0.35 | 0.58 | 0.00 | 0.01 |
| R-HSA-2682334 | EPH-Ephrin signaling | 0.16 | 0.01 | -0.31 | 0.74 | 0.02 | 0.03 |
| R-HSA-3928662 | EPHB-mediated forward signaling | 0.41 | -0.02 | -0.31 | 0.57 | 0.00 | 0.04 |
| R-HSA-5663213 | RHO GTPases Activate WASPs and WAVEs | 0.38 | -0.20 | -0.26 | 0.41 | 0.02 | 0.11 |
| R-HSA-9664407 | Parasite infection | 0.29 | -0.25 | -0.35 | 0.54 | 0.00 | 0.00 |
| R-HSA-9664417 | Leishmania phagocytosis | 0.29 | -0.25 | -0.35 | 0.54 | 0.00 | 0.00 |
| R-HSA-9664422 | FCGR3A-mediated phagocytosis | 0.29 | -0.25 | -0.35 | 0.54 | 0.00 | 0.00 |
| R-HSA-72649 | Translation initiation complex formation | 0.16 | 0.46 | 0.06 | 0.52 | 0.38 | 0.20 |
| R-HSA-72662 | Activation of the mRNA upon binding of the cap-binding complex and eIFs, and subsequent binding to 43S | 0.16 | 0.46 | 0.05 | 0.52 | 0.38 | 0.20 |
| R-HSA-72695 | Formation of the ternary complex, and subsequently, the 43S complex | 0.18 | 0.51 | 0.08 | 0.41 | 0.37 | 0.13 |
| R-HSA-72702 | Ribosomal scanning and start codon recognition | 0.14 | 0.46 | 0.05 | 0.53 | 0.38 | 0.20 |
| R-HSA-447115 | Interleukin-12 family signaling | 0.20 | 0.10 | -0.11 | 0.73 | 0.34 | 0.38 |
| R-HSA-8950505 | Gene and protein expression by JAK-STAT signaling after Interleukin-12 stimulation | 0.20 | 0.19 | 0.01 | 0.73 | 0.37 | 0.39 |
| R-HSA-9020591 | Interleukin-12 signaling | 0.21 | 0.14 | -0.05 | 0.73 | 0.37 | 0.39 |
| R-HSA-373752 | Netrin-1 signaling | -0.06 | -0.36 | -0.32 | 0.67 | 0.02 | 0.01 |
| R-HSA-169893 | Prolonged ERK activation events | -0.10 | -0.25 | -0.38 | 0.73 | 0.00 | 0.00 |
| R-HSA-170984 | ARMS-mediated activation | -0.50 | 0.04 | -0.45 | 0.32 | 0.00 | 0.00 |
| R-HSA-186763 | Downstream signal transduction | -0.20 | -0.36 | -0.65 | 0.66 | 0.00 | 0.00 |
| R-HSA-186797 | Signaling by PDGF | -0.09 | -0.13 | -0.62 | 0.74 | 0.00 | 0.00 |
| R-HSA-354192 | Integrin signaling | -0.34 | -0.15 | -0.28 | 0.68 | 0.02 | 0.09 |
| R-HSA-372708 | p130Cas linkage to MAPK signaling for integrins | -0.32 | -0.06 | -0.26 | 0.69 | 0.04 | 0.12 |
| R-HSA-512988 | Interleukin-3, Interleukin-5 and GM-CSF signaling | -0.30 | -0.54 | -0.44 | 0.27 | 0.00 | 0.00 |
| R-HSA-6806834 | Signaling by MET | -0.39 | 0.14 | -0.50 | 0.49 | 0.00 | 0.00 |
| R-HSA-76009 | Platelet Aggregation (Plug Formation) | -0.27 | 0.08 | -0.28 | 0.70 | 0.03 | 0.06 |
| R-HSA-8848021 | Signaling by PTK6 | -0.36 | -0.01 | -0.25 | 0.65 | 0.04 | 0.15 |
| R-HSA-8849471 | PTK6 Regulates RHO GTPases, RAS GTPase and MAP kinases | -0.31 | -0.29 | -0.39 | 0.65 | 0.00 | 0.00 |
| R-HSA-8875555 | MET activates RAP1 and RAC1 | -0.42 | -0.19 | -0.64 | 0.58 | 0.00 | 0.00 |
| R-HSA-8875656 | MET receptor recycling | -0.56 | -0.75 | -0.62 | 0.00 | 0.00 | 0.00 |
| R-HSA-8875878 | MET promotes cell motility | -0.39 | 0.17 | -0.53 | 0.45 | 0.00 | 0.00 |
| R-HSA-9006927 | Signaling by Non-Receptor Tyrosine Kinases | -0.36 | -0.01 | -0.25 | 0.65 | 0.04 | 0.15 |
| R-HSA-912631 | Regulation of signaling by CBL | -0.61 | -0.44 | -0.44 | 0.03 | 0.00 | 0.00 |
| R-HSA-1989781 | PPARA activates gene expression | 0.37 | 0.09 | -0.35 | 0.65 | 0.00 | 0.00 |
| R-HSA-200425 | Carnitine metabolism | -0.08 | -0.25 | -0.56 | 0.73 | 0.00 | 0.00 |
| R-HSA-400206 | Regulation of lipid metabolism by PPARalpha | 0.37 | 0.09 | -0.36 | 0.65 | 0.00 | 0.00 |
| R-HSA-880009 | Interconversion of 2-oxoglutarate and 2-hydroxyglutarate | 0.42 | -0.35 | -0.66 | 0.06 | 0.00 | 0.00 |
| R-HSA-196843 | Vitamin B2 (riboflavin) metabolism | -0.62 | -0.03 | -0.21 | 0.07 | 0.00 | 0.25 |
| R-HSA-199220 | Vitamin B5 (pantothenate) metabolism | -0.22 | 0.63 | -0.33 | 0.00 | 0.01 | 0.00 |
| R-HSA-2046104 | alpha-linolenic (omega3) and linoleic (omega6) acid metabolism | 0.42 | 0.35 | -0.30 | 0.46 | 0.00 | 0.00 |
| R-HSA-2046105 | Linoleic acid (LA) metabolism | 0.98 | 0.58 | -0.27 | 0.00 | 0.00 | 0.00 |
| R-HSA-2046106 | alpha-linolenic acid (ALA) metabolism | 0.42 | 0.35 | -0.30 | 0.46 | 0.00 | 0.00 |
| R-HSA-75876 | Synthesis of very long-chain fatty acyl-CoAs | 0.46 | 0.30 | -0.35 | 0.44 | 0.00 | 0.00 |
| R-HSA-1227986 | Signaling by ERBB2 | -0.18 | -0.30 | -0.42 | 0.70 | 0.00 | 0.00 |
| R-HSA-6785631 | ERBB2 Regulates Cell Motility | 0.41 | -0.35 | -0.26 | 0.07 | 0.01 | 0.05 |
| R-HSA-1660661 | Sphingolipid de novo biosynthesis | -0.40 | -0.01 | -0.30 | 0.60 | 0.00 | 0.05 |
| R-HSA-192105 | Synthesis of bile acids and bile salts | 0.07 | -0.08 | -0.33 | 0.74 | 0.02 | 0.02 |
| R-HSA-194068 | Bile acid and bile salt metabolism | 0.07 | -0.04 | -0.27 | 0.74 | 0.09 | 0.10 |
| R-HSA-428157 | Sphingolipid metabolism | -0.26 | 0.14 | -0.30 | 0.68 | 0.02 | 0.02 |
| R-HSA-8957322 | Metabolism of steroids | 0.18 | -0.02 | -0.32 | 0.74 | 0.01 | 0.02 |
| R-HSA-1566948 | Elastic fibre formation | -0.01 | 0.23 | -0.45 | 0.73 | 0.00 | 0.00 |
| R-HSA-2129379 | Molecules associated with elastic fibres | -0.01 | 0.23 | -0.43 | 0.73 | 0.00 | 0.00 |
| R-HSA-69273 | Cyclin A/B1/B2 associated events during G2/M transition | -0.12 | -0.24 | -0.30 | 0.73 | 0.04 | 0.03 |
| R-HSA-111465 | Apoptotic cleavage of cellular proteins | 0.02 | -0.01 | -0.32 | 0.74 | 0.03 | 0.03 |
| R-HSA-75153 | Apoptotic execution phase | -0.01 | 0.07 | -0.33 | 0.74 | 0.02 | 0.01 |
| R-HSA-9648002 | RAS processing | -0.07 | 0.29 | -0.09 | 0.69 | 0.38 | 0.32 |
| R-HSA-888590 | GABA synthesis, release, reuptake and degradation | 0.84 | 0.39 | 0.01 | 0.00 | 0.00 | 0.29 |
| R-HSA-916853 | Degradation of GABA | 0.84 | 0.39 | -0.98 | 0.00 | 0.00 | 0.00 |
| R-HSA-156580 | Phase II - Conjugation of compounds | 0.31 | -0.01 | -0.12 | 0.69 | 0.26 | 0.38 |
| R-HSA-156581 | Methylation | 0.24 | -0.02 | -0.25 | 0.72 | 0.08 | 0.17 |
| R-HSA-1369062 | ABC transporters in lipid homeostasis | 0.44 | -0.21 | -0.27 | 0.24 | 0.00 | 0.08 |
| R-HSA-9603798 | Class I peroxisomal membrane protein import | 0.12 | 0.02 | -0.46 | 0.74 | 0.00 | 0.00 |
| R-HSA-170834 | Signaling by TGF-beta Receptor Complex | -0.10 | 0.01 | -0.34 | 0.74 | 0.01 | 0.01 |
| R-HSA-2173793 | Transcriptional activity of SMAD2/SMAD3:SMAD4 heterotrimer | -0.27 | 0.03 | -0.34 | 0.71 | 0.00 | 0.01 |
| R-HSA-2173795 | Downregulation of SMAD2/3:SMAD4 transcriptional activity | -0.24 | 0.24 | -0.34 | 0.62 | 0.00 | 0.00 |
| R-HSA-8852135 | Protein ubiquitination | -0.23 | -0.04 | -0.29 | 0.73 | 0.04 | 0.07 |
| R-HSA-8866652 | Synthesis of active ubiquitin: roles of E1 and E2 enzymes | -0.13 | 0.16 | -0.26 | 0.73 | 0.11 | 0.08 |
| R-HSA-9006936 | Signaling by TGFB family members | -0.10 | 0.01 | -0.33 | 0.74 | 0.02 | 0.02 |
| R-HSA-9033241 | Peroxisomal protein import | 0.00 | 0.10 | -0.37 | 0.74 | 0.00 | 0.00 |
| R-HSA-1482798 | Acyl chain remodeling of CL | 0.56 | -0.25 | -0.60 | 0.01 | 0.00 | 0.00 |
| R-HSA-77285 | Beta oxidation of myristoyl-CoA to lauroyl-CoA | 0.35 | -0.43 | -0.56 | 0.06 | 0.00 | 0.00 |
| R-HSA-77288 | mitochondrial fatty acid beta-oxidation of unsaturated fatty acids | 0.36 | -0.08 | -0.40 | 0.60 | 0.00 | 0.00 |
| R-HSA-77305 | Beta oxidation of palmitoyl-CoA to myristoyl-CoA | 0.39 | 0.00 | -0.30 | 0.60 | 0.00 | 0.05 |
| R-HSA-192905 | vRNP Assembly | 0.35 | -0.74 | -0.46 | 0.00 | 0.00 | 0.00 |
| R-HSA-6794361 | Neurexins and neuroligins | -0.08 | 0.49 | -0.08 | 0.32 | 0.38 | 0.10 |
| R-HSA-15869 | Metabolism of nucleotides | 0.19 | -0.09 | -0.24 | 0.73 | 0.13 | 0.20 |
| R-HSA-499943 | Interconversion of nucleotide di- and triphosphates | 0.08 | -0.24 | -0.09 | 0.71 | 0.38 | 0.37 |
| R-HSA-70635 | Urea cycle | 0.33 | 0.42 | -0.08 | 0.50 | 0.29 | 0.19 |
| R-HSA-70895 | Branched-chain amino acid catabolism | 0.28 | -0.02 | -0.48 | 0.71 | 0.00 | 0.00 |
| R-HSA-1169408 | ISG15 antiviral mechanism | 0.11 | -0.09 | -0.55 | 0.74 | 0.00 | 0.00 |
| R-HSA-1169410 | Antiviral mechanism by IFN-stimulated genes | 0.10 | -0.21 | -0.53 | 0.72 | 0.00 | 0.00 |
| R-HSA-168276 | NS1 Mediated Effects on Host Pathways | 0.48 | -0.13 | -0.54 | 0.26 | 0.00 | 0.00 |
| R-HSA-913531 | Interferon Signaling | 0.09 | -0.13 | -0.43 | 0.74 | 0.00 | 0.00 |
| R-HSA-193648 | NRAGE signals death through JNK | -0.04 | 0.69 | -0.52 | 0.01 | 0.00 | 0.00 |
| R-HSA-193704 | p75 NTR receptor-mediated signalling | -0.04 | 0.17 | -0.44 | 0.74 | 0.00 | 0.00 |
| R-HSA-204998 | Cell death signalling via NRAGE, NRIF and NADE | -0.04 | 0.26 | -0.48 | 0.71 | 0.00 | 0.00 |
| R-HSA-73887 | Death Receptor Signalling | 0.06 | 0.18 | -0.43 | 0.74 | 0.00 | 0.00 |
| R-HSA-140342 | Apoptosis induced DNA fragmentation | -0.22 | 0.45 | -0.32 | 0.21 | 0.01 | 0.00 |
| R-HSA-2559584 | Formation of Senescence-Associated Heterochromatin Foci (SAHF) | -0.36 | 0.69 | -0.26 | 0.00 | 0.03 | 0.00 |
| R-HSA-2559586 | DNA Damage/Telomere Stress Induced Senescence | -0.46 | 0.38 | -0.09 | 0.02 | 0.15 | 0.22 |
| R-HSA-1630316 | Glycosaminoglycan metabolism | 0.09 | 0.22 | -0.32 | 0.73 | 0.02 | 0.01 |
| R-HSA-1638074 | Keratan sulfate/keratin metabolism | 0.18 | 0.00 | -0.46 | 0.74 | 0.00 | 0.00 |
| R-HSA-2022854 | Keratan sulfate biosynthesis | 0.39 | 0.14 | -0.43 | 0.63 | 0.00 | 0.00 |
| R-HSA-2022857 | Keratan sulfate degradation | 0.18 | 0.00 | -0.56 | 0.74 | 0.00 | 0.00 |
| R-HSA-3000178 | ECM proteoglycans | -0.17 | 0.23 | -0.40 | 0.68 | 0.00 | 0.00 |
| R-HSA-3560782 | Diseases associated with glycosaminoglycan metabolism | 0.21 | 0.23 | -0.39 | 0.72 | 0.00 | 0.00 |
| R-HSA-3656225 | Defective CHST6 causes MCDC1 | 0.39 | 0.14 | -0.55 | 0.63 | 0.00 | 0.00 |
| R-HSA-3656243 | Defective ST3GAL3 causes MCT12 and EIEE15 | 0.39 | 0.14 | -0.58 | 0.63 | 0.00 | 0.00 |
| R-HSA-3656244 | Defective B4GALT1 causes B4GALT1-CDG (CDG-2d) | 0.39 | 0.14 | -0.53 | 0.63 | 0.00 | 0.00 |
| R-HSA-3781865 | Diseases of glycosylation | 0.04 | 0.04 | -0.39 | 0.74 | 0.00 | 0.00 |
| R-HSA-109606 | Intrinsic Pathway for Apoptosis | 0.09 | -0.30 | -0.23 | 0.67 | 0.19 | 0.13 |
| R-HSA-111447 | Activation of BAD and translocation to mitochondria | -0.10 | -0.72 | -0.31 | 0.01 | 0.03 | 0.00 |
| R-HSA-114452 | Activation of BH3-only proteins | -0.11 | -0.37 | -0.29 | 0.66 | 0.06 | 0.02 |
| R-HSA-69473 | G2/M DNA damage checkpoint | -0.22 | -0.26 | -0.20 | 0.70 | 0.21 | 0.22 |
| R-HSA-75035 | Chk1/Chk2(Cds1) mediated inactivation of Cyclin B:Cdk1 complex | -0.10 | -0.72 | -0.24 | 0.01 | 0.18 | 0.00 |
| R-HSA-9614085 | FOXO-mediated transcription | -0.08 | -0.75 | -0.28 | 0.00 | 0.07 | 0.00 |
| R-HSA-9614399 | Regulation of localization of FOXO transcription factors | -0.16 | -0.66 | -0.29 | 0.06 | 0.05 | 0.00 |
| R-HSA-111457 | Release of apoptotic factors from the mitochondria | 0.28 | -0.22 | -0.20 | 0.60 | 0.15 | 0.25 |
| R-HSA-111458 | Formation of apoptosome | 0.41 | -0.27 | -0.29 | 0.19 | 0.00 | 0.04 |
| R-HSA-111459 | Activation of caspases through apoptosome-mediated cleavage | 0.91 | 0.09 | -0.27 | 0.00 | 0.00 | 0.08 |
| R-HSA-111461 | Cytochrome c-mediated apoptotic response | 0.41 | -0.27 | -0.33 | 0.19 | 0.00 | 0.01 |
| R-HSA-111463 | SMAC (DIABLO) binds to IAPs | 0.32 | -0.17 | -0.32 | 0.60 | 0.00 | 0.03 |
| R-HSA-111464 | SMAC(DIABLO)-mediated dissociation of IAP:caspase complexes | 0.32 | -0.17 | -0.32 | 0.60 | 0.00 | 0.03 |
| R-HSA-111469 | SMAC, XIAP-regulated apoptotic response | 0.32 | -0.17 | -0.32 | 0.60 | 0.00 | 0.03 |
| R-HSA-111471 | Apoptotic factor-mediated response | 0.28 | -0.29 | -0.30 | 0.49 | 0.01 | 0.03 |
| R-HSA-2151201 | Transcriptional activation of mitochondrial biogenesis | 0.33 | 0.01 | -0.48 | 0.68 | 0.00 | 0.00 |
| R-HSA-3299685 | Detoxification of Reactive Oxygen Species | 0.35 | -0.19 | -0.08 | 0.51 | 0.27 | 0.39 |
| R-HSA-5218859 | Regulated Necrosis | 0.22 | 0.00 | -0.25 | 0.73 | 0.08 | 0.15 |
| R-HSA-5620971 | Pyroptosis | 0.52 | 0.07 | -0.18 | 0.33 | 0.02 | 0.31 |
| R-HSA-9627069 | Regulation of the apoptosome activity | 0.41 | -0.27 | -0.29 | 0.19 | 0.00 | 0.04 |
| R-HSA-8964540 | Alanine metabolism | -0.49 | -0.17 | -0.46 | 0.45 | 0.00 | 0.00 |
| R-HSA-114516 | Disinhibition of SNARE formation | -0.60 | 0.90 | -0.32 | 0.00 | 0.00 | 0.00 |
| R-HSA-449836 | Other interleukin signaling | -0.05 | 0.64 | -0.26 | 0.03 | 0.12 | 0.00 |
| R-HSA-165159 | MTOR signalling | -0.03 | -0.17 | -0.46 | 0.74 | 0.00 | 0.00 |
| R-HSA-166208 | mTORC1-mediated signalling | -0.05 | -0.03 | -0.36 | 0.74 | 0.00 | 0.00 |
| R-HSA-380972 | Energy dependent regulation of mTOR by LKB1-AMPK | -0.03 | -0.25 | -0.48 | 0.72 | 0.00 | 0.00 |
| R-HSA-8943724 | Regulation of PTEN gene transcription | 0.28 | -0.26 | -0.31 | 0.53 | 0.01 | 0.02 |
| R-HSA-9639288 | Amino acids regulate mTORC1 | -0.08 | -0.08 | -0.31 | 0.74 | 0.03 | 0.04 |
| R-HSA-204005 | COPII-mediated vesicle transport | -0.06 | -0.01 | -0.37 | 0.74 | 0.00 | 0.00 |
| R-HSA-9700206 | Signaling by ALK in cancer | 0.08 | -0.25 | -0.35 | 0.71 | 0.01 | 0.00 |
| R-HSA-9725370 | Signaling by ALK fusions and activated point mutants | 0.08 | -0.25 | -0.35 | 0.71 | 0.01 | 0.00 |
| R-HSA-6805567 | Keratinization | 0.17 | 0.55 | 0.24 | 0.31 | 0.14 | 0.01 |
| R-HSA-6809371 | Formation of the cornified envelope | 0.17 | 0.55 | 0.17 | 0.31 | 0.29 | 0.04 |
| R-HSA-8862803 | Deregulated CDK5 triggers multiple neurodegenerative pathways in Alzheimer's disease models | -0.30 | -0.42 | -0.21 | 0.52 | 0.14 | 0.09 |
| R-HSA-8863678 | Neurodegenerative Diseases | -0.30 | -0.42 | -0.21 | 0.52 | 0.14 | 0.09 |
| R-HSA-9645723 | Diseases of programmed cell death | -0.27 | -0.40 | -0.01 | 0.59 | 0.36 | 0.28 |
| R-HSA-71240 | Tryptophan catabolism | 0.44 | -0.16 | -0.16 | 0.33 | 0.08 | 0.34 |
| R-HSA-159227 | Transport of the SLBP independent Mature mRNA | 0.55 | 0.44 | -0.51 | 0.10 | 0.00 | 0.00 |
| R-HSA-159230 | Transport of the SLBP Dependant Mature mRNA | 0.55 | 0.44 | -0.51 | 0.10 | 0.00 | 0.00 |
| R-HSA-159231 | Transport of Mature mRNA Derived from an Intronless Transcript | 0.55 | 0.44 | -0.49 | 0.10 | 0.00 | 0.00 |
| R-HSA-159234 | Transport of Mature mRNAs Derived from Intronless Transcripts | 0.55 | 0.44 | -0.49 | 0.10 | 0.00 | 0.00 |
| R-HSA-162587 | HIV Life Cycle | -0.03 | 0.02 | -0.31 | 0.74 | 0.04 | 0.03 |
| R-HSA-162599 | Late Phase of HIV Life Cycle | -0.03 | 0.02 | -0.31 | 0.74 | 0.04 | 0.04 |
| R-HSA-165054 | Rev-mediated nuclear export of HIV RNA | -0.02 | 0.28 | -0.51 | 0.71 | 0.00 | 0.00 |
| R-HSA-168271 | Transport of Ribonucleoproteins into the Host Nucleus | 0.61 | 0.06 | -0.57 | 0.09 | 0.00 | 0.00 |
| R-HSA-168274 | Export of Viral Ribonucleoproteins from Nucleus | -0.04 | 0.21 | -0.55 | 0.73 | 0.00 | 0.00 |
| R-HSA-168325 | Viral Messenger RNA Synthesis | 0.84 | 0.64 | -0.44 | 0.00 | 0.00 | 0.00 |
| R-HSA-168333 | NEP/NS2 Interacts with the Cellular Export Machinery | -0.04 | 0.21 | -0.55 | 0.73 | 0.00 | 0.00 |
| R-HSA-170822 | Regulation of Glucokinase by Glucokinase Regulatory Protein | 0.84 | 0.64 | -0.56 | 0.00 | 0.00 | 0.00 |
| R-HSA-176033 | Interactions of Vpr with host cellular proteins | 0.68 | 0.56 | -0.48 | 0.00 | 0.00 | 0.00 |
| R-HSA-177243 | Interactions of Rev with host cellular proteins | 0.08 | 0.08 | -0.48 | 0.74 | 0.00 | 0.00 |
| R-HSA-180746 | Nuclear import of Rev protein | 0.61 | 0.06 | -0.50 | 0.09 | 0.00 | 0.00 |
| R-HSA-180910 | Vpr-mediated nuclear import of PICs | 0.84 | 0.64 | -0.51 | 0.00 | 0.00 | 0.00 |
| R-HSA-191859 | snRNP Assembly | 0.40 | 0.16 | -0.36 | 0.62 | 0.00 | 0.00 |
| R-HSA-194441 | Metabolism of non-coding RNA | 0.40 | 0.16 | -0.36 | 0.62 | 0.00 | 0.00 |
| R-HSA-211000 | Gene Silencing by RNA | -0.26 | -0.30 | -0.11 | 0.67 | 0.32 | 0.33 |
| R-HSA-2980766 | Nuclear Envelope Breakdown | 0.07 | 0.75 | -0.47 | 0.00 | 0.00 | 0.00 |
| R-HSA-2990846 | SUMOylation | 0.61 | 0.52 | -0.37 | 0.01 | 0.00 | 0.00 |
| R-HSA-3108214 | SUMOylation of DNA damage response and repair proteins | 0.84 | 0.64 | -0.46 | 0.00 | 0.00 | 0.00 |
| R-HSA-3108232 | SUMO E3 ligases SUMOylate target proteins | 0.73 | 0.49 | -0.37 | 0.00 | 0.00 | 0.00 |
| R-HSA-3232142 | SUMOylation of ubiquitinylation proteins | 0.84 | 0.64 | -0.46 | 0.00 | 0.00 | 0.00 |
| R-HSA-3301854 | Nuclear Pore Complex (NPC) Disassembly | 0.61 | 0.77 | -0.49 | 0.00 | 0.00 | 0.00 |
| R-HSA-4085377 | SUMOylation of SUMOylation proteins | 0.84 | 0.64 | -0.47 | 0.00 | 0.00 | 0.00 |
| R-HSA-4551638 | SUMOylation of chromatin organization proteins | 0.84 | 0.64 | -0.33 | 0.00 | 0.00 | 0.00 |
| R-HSA-4570464 | SUMOylation of RNA binding proteins | 0.69 | 0.65 | -0.43 | 0.00 | 0.00 | 0.00 |
| R-HSA-4615885 | SUMOylation of DNA replication proteins | 0.84 | 0.64 | -0.38 | 0.00 | 0.00 | 0.00 |
| R-HSA-5578749 | Transcriptional regulation by small RNAs | -0.26 | -0.13 | -0.09 | 0.72 | 0.34 | 0.39 |
| R-HSA-5619107 | Defective TPR may confer susceptibility towards thyroid papillary carcinoma (TPC) | 0.84 | 0.64 | -0.56 | 0.00 | 0.00 | 0.00 |
| R-HSA-6784531 | tRNA processing in the nucleus | -0.34 | -0.43 | -0.34 | 0.47 | 0.00 | 0.00 |
| R-HSA-72306 | tRNA processing | -0.09 | -0.15 | -0.33 | 0.74 | 0.01 | 0.01 |
| R-HSA-9610379 | HCMV Late Events | 0.07 | 0.07 | -0.10 | 0.74 | 0.37 | 0.39 |
| R-HSA-2161522 | Abacavir transport and metabolism | 0.92 | 1.04 | -0.30 | 0.00 | 0.00 | 0.00 |
| R-HSA-2161541 | Abacavir metabolism | 0.92 | 1.04 | -0.43 | 0.00 | 0.00 | 0.00 |
| R-HSA-74259 | Purine catabolism | 0.29 | -0.49 | -0.34 | 0.05 | 0.00 | 0.00 |
| R-HSA-8956319 | Nucleobase catabolism | 0.15 | -0.21 | -0.31 | 0.70 | 0.02 | 0.03 |
| R-HSA-166658 | Complement cascade | 0.18 | 0.33 | -0.12 | 0.69 | 0.34 | 0.24 |
| R-HSA-198933 | Immunoregulatory interactions between a Lymphoid and a non-Lymphoid cell | -0.31 | 0.57 | -0.17 | 0.00 | 0.20 | 0.00 |
| R-HSA-977606 | Regulation of Complement cascade | 0.12 | 0.29 | -0.11 | 0.71 | 0.36 | 0.30 |
| R-HSA-6803157 | Antimicrobial peptides | -0.18 | 0.06 | 0.24 | 0.73 | 0.13 | 0.19 |
| R-HSA-5368286 | Mitochondrial translation initiation | -0.24 | 0.45 | -0.18 | 0.16 | 0.23 | 0.03 |
| R-HSA-5368287 | Mitochondrial translation | -0.15 | 0.29 | -0.21 | 0.65 | 0.23 | 0.11 |
| R-HSA-5389840 | Mitochondrial translation elongation | -0.15 | 0.29 | -0.18 | 0.65 | 0.29 | 0.17 |
| R-HSA-5419276 | Mitochondrial translation termination | -0.24 | 0.45 | -0.19 | 0.16 | 0.22 | 0.03 |
| R-HSA-532668 | N-glycan trimming in the ER and Calnexin/Calreticulin cycle | -0.14 | -0.42 | -0.35 | 0.59 | 0.00 | 0.00 |
| R-HSA-901042 | Calnexin/calreticulin cycle | -0.15 | -0.32 | -0.38 | 0.69 | 0.00 | 0.00 |
| R-HSA-9678108 | SARS-CoV-1 Infection | -0.55 | -0.56 | -0.22 | 0.02 | 0.01 | 0.01 |
| R-HSA-9683686 | Maturation of spike protein | -0.88 | -0.86 | -0.38 | 0.00 | 0.00 | 0.00 |
| R-HSA-9683701 | Translation of Structural Proteins | -0.84 | -0.42 | -0.25 | 0.00 | 0.00 | 0.04 |
| R-HSA-9694516 | SARS-CoV-2 Infection | -0.34 | -0.53 | -0.21 | 0.24 | 0.11 | 0.02 |
| R-HSA-9694548 | Maturation of spike protein | -0.40 | -0.67 | -0.28 | 0.02 | 0.01 | 0.00 |
| R-HSA-9694635 | Translation of Structural Proteins | -0.47 | -0.44 | -0.21 | 0.23 | 0.03 | 0.07 |
| R-HSA-1234158 | Regulation of gene expression by Hypoxia-inducible Factor | -0.82 | -0.48 | -0.44 | 0.00 | 0.00 | 0.00 |
| R-HSA-383280 | Nuclear Receptor transcription pathway | 0.74 | 0.68 | -0.30 | 0.00 | 0.00 | 0.00 |
| R-HSA-429914 | Deadenylation-dependent mRNA decay | -0.13 | 0.22 | -0.29 | 0.71 | 0.06 | 0.02 |
| R-HSA-429947 | Deadenylation of mRNA | -0.03 | 0.05 | -0.43 | 0.74 | 0.00 | 0.00 |
| R-HSA-4085001 | Sialic acid metabolism | 0.13 | -0.46 | -0.11 | 0.34 | 0.36 | 0.17 |
| R-HSA-446193 | Biosynthesis of the N-glycan precursor (dolichol lipid-linked oligosaccharide, LLO) and transfer to a nascent protein | -0.04 | -0.27 | -0.25 | 0.72 | 0.15 | 0.11 |
| R-HSA-446219 | Synthesis of substrates in N-glycan biosythesis | -0.04 | -0.27 | -0.21 | 0.72 | 0.25 | 0.20 |
| R-HSA-156590 | Glutathione conjugation | 0.41 | 0.12 | -0.24 | 0.61 | 0.02 | 0.14 |
| R-HSA-196836 | Vitamin C (ascorbate) metabolism | 0.32 | 0.25 | -0.55 | 0.67 | 0.00 | 0.00 |
| R-HSA-140837 | Intrinsic Pathway of Fibrin Clot Formation | 0.31 | 0.11 | -0.12 | 0.70 | 0.27 | 0.37 |
| R-HSA-140877 | Formation of Fibrin Clot (Clotting Cascade) | 0.26 | 0.11 | -0.14 | 0.72 | 0.27 | 0.35 |
| R-HSA-9651496 | Defects of contact activation system (CAS) and kallikrein/kinin system (KKS) | 0.61 | 0.17 | -0.08 | 0.11 | 0.02 | 0.38 |
| R-HSA-9657689 | Defective SERPING1 causes hereditary angioedema | 0.68 | -0.06 | 0.08 | 0.01 | 0.00 | 0.40 |
| R-HSA-9671793 | Diseases of hemostasis | 0.61 | 0.17 | -0.08 | 0.11 | 0.02 | 0.38 |
| R-HSA-373755 | Semaphorin interactions | 0.04 | -0.12 | -0.37 | 0.74 | 0.00 | 0.00 |
| R-HSA-399954 | Sema3A PAK dependent Axon repulsion | 0.25 | -0.19 | -0.25 | 0.65 | 0.07 | 0.14 |
| R-HSA-70221 | Glycogen breakdown (glycogenolysis) | 0.08 | -0.64 | -0.56 | 0.02 | 0.00 | 0.00 |
| R-HSA-8982491 | Glycogen metabolism | -0.08 | -0.57 | -0.54 | 0.24 | 0.00 | 0.00 |
| R-HSA-9636667 | Manipulation of host energy metabolism | 0.68 | -0.67 | -0.38 | 0.00 | 0.00 | 0.00 |
| R-HSA-1660662 | Glycosphingolipid metabolism | -0.12 | 0.28 | -0.30 | 0.67 | 0.04 | 0.01 |
| R-HSA-399956 | CRMPs in Sema3A signaling | 0.03 | -0.29 | -0.21 | 0.70 | 0.26 | 0.19 |
| R-HSA-1483249 | Inositol phosphate metabolism | 0.67 | 0.17 | -0.38 | 0.03 | 0.00 | 0.00 |
| R-HSA-1855183 | Synthesis of IP2, IP, and Ins in the cytosol | 0.78 | 0.31 | -0.46 | 0.00 | 0.00 | 0.00 |
| R-HSA-3322077 | Glycogen synthesis | -0.24 | -0.43 | -0.56 | 0.56 | 0.00 | 0.00 |
| R-HSA-70370 | Galactose catabolism | 0.33 | -0.51 | -0.36 | 0.01 | 0.00 | 0.00 |
| R-HSA-3928665 | EPH-ephrin mediated repulsion of cells | 0.03 | 0.10 | -0.32 | 0.74 | 0.03 | 0.02 |
| R-HSA-432720 | Lysosome Vesicle Biogenesis | -0.28 | -0.06 | -0.23 | 0.71 | 0.10 | 0.21 |
| R-HSA-5099900 | WNT5A-dependent internalization of FZD4 | -0.02 | 0.02 | -0.27 | 0.74 | 0.10 | 0.09 |
| R-HSA-5140745 | WNT5A-dependent internalization of FZD2, FZD5 and ROR2 | -0.02 | 0.02 | -0.27 | 0.74 | 0.09 | 0.09 |
| R-HSA-8949215 | Mitochondrial calcium ion transport | -0.20 | -0.05 | -0.42 | 0.73 | 0.00 | 0.00 |
| R-HSA-8949664 | Processing of SMDT1 | -0.26 | -0.04 | -0.38 | 0.72 | 0.00 | 0.00 |
| R-HSA-9012852 | Signaling by NOTCH3 | -0.25 | 0.26 | -0.34 | 0.59 | 0.00 | 0.00 |
| R-HSA-9013507 | NOTCH3 Activation and Transmission of Signal to the Nucleus | -0.88 | 0.12 | -0.36 | 0.00 | 0.00 | 0.00 |
| R-HSA-9017802 | Noncanonical activation of NOTCH3 | -1.00 | -0.22 | -0.61 | 0.00 | 0.00 | 0.00 |
| R-HSA-375165 | NCAM signaling for neurite out-growth | -0.18 | -0.12 | -0.44 | 0.74 | 0.00 | 0.00 |
| R-HSA-445095 | Interaction between L1 and Ankyrins | -0.32 | 0.29 | -0.35 | 0.38 | 0.00 | 0.00 |
| R-HSA-8939243 | RUNX1 interacts with co-factors whose precise effect on RUNX1 targets is not known | -0.60 | -0.09 | -0.46 | 0.11 | 0.00 | 0.00 |
| R-HSA-75205 | Dissolution of Fibrin Clot | 0.34 | -0.03 | -0.23 | 0.65 | 0.06 | 0.21 |
| R-HSA-3229121 | Glycogen storage diseases | -0.65 | -0.48 | -0.45 | 0.01 | 0.00 | 0.00 |
| R-HSA-3878781 | Glycogen storage disease type IV (GBE1) | -0.93 | -0.90 | -0.81 | 0.00 | 0.00 | 0.00 |
| R-HSA-5663084 | Diseases of carbohydrate metabolism | -0.46 | -0.21 | -0.39 | 0.50 | 0.00 | 0.00 |
| R-HSA-189445 | Metabolism of porphyrins | -0.05 | -0.27 | -0.16 | 0.72 | 0.33 | 0.28 |
| R-HSA-189483 | Heme degradation | 0.12 | -0.25 | -0.12 | 0.70 | 0.36 | 0.35 |
| R-HSA-5633007 | Regulation of TP53 Activity | -0.07 | -0.05 | -0.36 | 0.74 | 0.00 | 0.00 |
| R-HSA-6804759 | Regulation of TP53 Activity through Association with Co-factors | -0.76 | 0.40 | -0.20 | 0.00 | 0.00 | 0.05 |
| R-HSA-381042 | PERK regulates gene expression | -0.15 | 0.50 | -0.18 | 0.19 | 0.28 | 0.02 |
| R-HSA-72731 | Recycling of eIF2:GDP | 0.49 | 0.42 | -0.23 | 0.22 | 0.01 | 0.01 |
| R-HSA-9648895 | Response of EIF2AK1 (HRI) to heme deficiency | 0.49 | 0.42 | -0.09 | 0.22 | 0.10 | 0.18 |
| R-HSA-212165 | Epigenetic regulation of gene expression | -0.20 | -0.28 | -0.11 | 0.70 | 0.35 | 0.34 |
| R-HSA-427389 | ERCC6 (CSB) and EHMT2 (G9a) positively regulate rRNA expression | -0.30 | -0.39 | 0.15 | 0.57 | 0.23 | 0.14 |
| R-HSA-5250913 | Positive epigenetic regulation of rRNA expression | -0.20 | -0.28 | 0.01 | 0.70 | 0.37 | 0.37 |
| R-HSA-73772 | RNA Polymerase I Promoter Escape | -0.63 | -0.39 | 0.12 | 0.03 | 0.01 | 0.18 |
| R-HSA-73854 | RNA Polymerase I Promoter Clearance | -0.21 | -0.52 | 0.02 | 0.38 | 0.37 | 0.12 |
| R-HSA-8953750 | Transcriptional Regulation by E2F6 | 0.68 | -0.37 | -0.36 | 0.00 | 0.00 | 0.00 |
| R-HSA-1442490 | Collagen degradation | -0.11 | 0.19 | -0.31 | 0.72 | 0.03 | 0.01 |
| R-HSA-1474290 | Collagen formation | -0.05 | 0.07 | -0.36 | 0.74 | 0.00 | 0.00 |
| R-HSA-1650814 | Collagen biosynthesis and modifying enzymes | 0.00 | 0.15 | -0.44 | 0.74 | 0.00 | 0.00 |
| R-HSA-2022090 | Assembly of collagen fibrils and other multimeric structures | -0.18 | 0.06 | -0.40 | 0.73 | 0.00 | 0.00 |
| R-HSA-2173782 | Binding and Uptake of Ligands by Scavenger Receptors | 0.10 | 0.14 | -0.25 | 0.74 | 0.14 | 0.12 |
| R-HSA-2214320 | Anchoring fibril formation | -0.45 | 0.50 | -0.55 | 0.00 | 0.00 | 0.00 |
| R-HSA-2243919 | Crosslinking of collagen fibrils | -0.45 | 0.50 | -0.58 | 0.00 | 0.00 | 0.00 |
| R-HSA-3000157 | Laminin interactions | -0.23 | 0.15 | -0.54 | 0.70 | 0.00 | 0.00 |
| R-HSA-3000171 | Non-integrin membrane-ECM interactions | -0.34 | 0.28 | -0.47 | 0.37 | 0.00 | 0.00 |
| R-HSA-3000480 | Scavenging by Class A Receptors | -0.05 | 0.11 | -0.29 | 0.74 | 0.07 | 0.05 |
| R-HSA-419037 | NCAM1 interactions | -0.17 | -0.08 | -0.40 | 0.74 | 0.00 | 0.00 |
| R-HSA-8948216 | Collagen chain trimerization | -0.13 | 0.16 | -0.49 | 0.73 | 0.00 | 0.00 |
| R-HSA-170968 | Frs2-mediated activation | 0.00 | -0.17 | -0.31 | 0.74 | 0.03 | 0.03 |
| R-HSA-9006335 | Signaling by Erythropoietin | -0.59 | -0.71 | -0.49 | 0.00 | 0.00 | 0.00 |
| R-HSA-9027284 | Erythropoietin activates RAS | -0.59 | -0.71 | -0.41 | 0.00 | 0.00 | 0.00 |
| R-HSA-1655829 | Regulation of cholesterol biosynthesis by SREBP (SREBF) | 0.39 | 0.14 | -0.48 | 0.63 | 0.00 | 0.00 |
| R-HSA-191273 | Cholesterol biosynthesis | 0.88 | 0.33 | -0.42 | 0.00 | 0.00 | 0.00 |
| R-HSA-2426168 | Activation of gene expression by SREBF (SREBP) | 0.57 | 0.39 | -0.47 | 0.11 | 0.00 | 0.00 |
| R-HSA-71336 | Pentose phosphate pathway | 0.18 | -0.16 | -0.16 | 0.71 | 0.29 | 0.33 |
| R-HSA-74217 | Purine salvage | 0.36 | -0.12 | -0.30 | 0.58 | 0.00 | 0.04 |
| R-HSA-8956321 | Nucleotide salvage | 0.36 | -0.12 | -0.28 | 0.58 | 0.01 | 0.08 |
| R-HSA-446343 | Localization of the PINCH-ILK-PARVIN complex to focal adhesions | 0.37 | 0.55 | -0.60 | 0.17 | 0.00 | 0.00 |
| R-HSA-446353 | Cell-extracellular matrix interactions | -0.11 | 0.00 | -0.49 | 0.74 | 0.00 | 0.00 |
| R-HSA-425410 | Metal ion SLC transporters | 0.79 | 0.10 | -0.33 | 0.00 | 0.00 | 0.01 |
| R-HSA-5619049 | Defective SLC40A1 causes hemochromatosis 4 (HFE4) (macrophages) | 0.79 | 0.10 | -0.35 | 0.00 | 0.00 | 0.00 |
| R-HSA-5619060 | Defective CP causes aceruloplasminemia (ACERULOP) | 0.79 | 0.10 | -0.35 | 0.00 | 0.00 | 0.00 |
| R-HSA-917937 | Iron uptake and transport | -0.18 | 0.05 | -0.24 | 0.73 | 0.13 | 0.17 |
| R-HSA-110314 | Recognition of DNA damage by PCNA-containing replication complex | 0.00 | 0.43 | -0.36 | 0.54 | 0.00 | 0.00 |
| R-HSA-5696394 | DNA Damage Recognition in GG-NER | -0.08 | -0.40 | -0.38 | 0.63 | 0.00 | 0.00 |
| R-HSA-5696395 | Formation of Incision Complex in GG-NER | -0.10 | 0.16 | -0.37 | 0.73 | 0.00 | 0.00 |
| R-HSA-5696398 | Nucleotide Excision Repair | -0.05 | -0.24 | -0.33 | 0.73 | 0.02 | 0.01 |
| R-HSA-5696399 | Global Genome Nucleotide Excision Repair (GG-NER) | -0.09 | -0.32 | -0.36 | 0.70 | 0.00 | 0.00 |
| R-HSA-5696400 | Dual Incision in GG-NER | 0.00 | 0.43 | -0.37 | 0.54 | 0.00 | 0.00 |
| R-HSA-6781823 | Formation of TC-NER Pre-Incision Complex | 0.02 | -0.23 | -0.34 | 0.73 | 0.01 | 0.01 |
| R-HSA-6781827 | Transcription-Coupled Nucleotide Excision Repair (TC-NER) | 0.02 | -0.23 | -0.34 | 0.73 | 0.01 | 0.01 |
| R-HSA-6782135 | Dual incision in TC-NER | 0.17 | 0.60 | -0.31 | 0.15 | 0.02 | 0.00 |
| R-HSA-6782210 | Gap-filling DNA repair synthesis and ligation in TC-NER | 0.17 | 0.60 | -0.30 | 0.15 | 0.03 | 0.00 |
| R-HSA-73893 | DNA Damage Bypass | 0.17 | 0.15 | -0.42 | 0.74 | 0.00 | 0.00 |
| R-HSA-73894 | DNA Repair | -0.02 | -0.16 | -0.32 | 0.74 | 0.02 | 0.02 |
| R-HSA-3238698 | WNT ligand biogenesis and trafficking | 0.00 | 0.49 | -0.12 | 0.41 | 0.37 | 0.07 |
| R-HSA-428890 | Role of ABL in ROBO-SLIT signaling | 0.13 | 0.23 | -0.63 | 0.73 | 0.00 | 0.00 |
| R-HSA-162791 | Attachment of GPI anchor to uPAR | -0.67 | -0.47 | -0.40 | 0.01 | 0.00 | 0.00 |
| R-HSA-163125 | Post-translational modification: synthesis of GPI-anchored proteins | 0.13 | 0.22 | -0.16 | 0.73 | 0.31 | 0.26 |
| R-HSA-190236 | Signaling by FGFR | 0.02 | -0.13 | -0.23 | 0.74 | 0.20 | 0.21 |
| R-HSA-5654738 | Signaling by FGFR2 | 0.02 | -0.13 | -0.22 | 0.74 | 0.22 | 0.23 |
| R-HSA-6803529 | FGFR2 alternative splicing | 0.20 | 0.13 | -0.25 | 0.73 | 0.11 | 0.13 |
| R-HSA-193639 | p75NTR signals via NF-kB | 0.08 | 0.35 | -0.32 | 0.67 | 0.02 | 0.00 |
| R-HSA-205043 | NRIF signals cell death from the nucleus | 0.08 | 0.35 | -0.37 | 0.67 | 0.00 | 0.00 |
| R-HSA-209543 | p75NTR recruits signalling complexes | 0.08 | 0.35 | -0.28 | 0.67 | 0.07 | 0.01 |
| R-HSA-209560 | NF-kB is activated and signals survival | 0.08 | 0.35 | -0.25 | 0.67 | 0.14 | 0.02 |
| R-HSA-5205647 | Mitophagy | -0.09 | 0.20 | -0.30 | 0.72 | 0.04 | 0.01 |
| R-HSA-5205685 | PINK1-PRKN Mediated Mitophagy | -0.09 | 0.20 | -0.30 | 0.72 | 0.04 | 0.02 |
| R-HSA-9664873 | Pexophagy | 0.08 | 0.35 | -0.42 | 0.67 | 0.00 | 0.00 |
| R-HSA-2980767 | Activation of NIMA Kinases NEK9, NEK6, NEK7 | 0.37 | 0.90 | -0.34 | 0.00 | 0.00 | 0.00 |
| R-HSA-1227990 | Signaling by ERBB2 in Cancer | -0.58 | -0.79 | -0.41 | 0.00 | 0.00 | 0.00 |
| R-HSA-1236382 | Constitutive Signaling by Ligand-Responsive EGFR Cancer Variants | -0.62 | -0.54 | -0.45 | 0.01 | 0.00 | 0.00 |
| R-HSA-1643713 | Signaling by EGFR in Cancer | -0.62 | -0.54 | -0.34 | 0.01 | 0.00 | 0.00 |
| R-HSA-5213460 | RIPK1-mediated regulated necrosis | 0.01 | -0.05 | -0.32 | 0.74 | 0.02 | 0.02 |
| R-HSA-5637810 | Constitutive Signaling by EGFRvIII | -0.58 | -0.79 | -0.56 | 0.00 | 0.00 | 0.00 |
| R-HSA-5637812 | Signaling by EGFRvIII in Cancer | -0.58 | -0.79 | -0.56 | 0.00 | 0.00 | 0.00 |
| R-HSA-5637815 | Signaling by Ligand-Responsive EGFR Variants in Cancer | -0.62 | -0.54 | -0.45 | 0.01 | 0.00 | 0.00 |
| R-HSA-5675482 | Regulation of necroptotic cell death | 0.01 | -0.05 | -0.32 | 0.74 | 0.02 | 0.02 |
| R-HSA-8863795 | Downregulation of ERBB2 signaling | -0.25 | -0.14 | -0.27 | 0.72 | 0.05 | 0.10 |
| R-HSA-9013418 | RHOBTB2 GTPase cycle | -0.23 | -0.14 | 0.89 | 0.73 | 0.00 | 0.00 |
| R-HSA-9634285 | Constitutive Signaling by Overexpressed ERBB2 | -0.58 | -0.79 | -0.60 | 0.00 | 0.00 | 0.00 |
| R-HSA-9652282 | Drug-mediated inhibition of ERBB2 signaling | -0.57 | -0.93 | -0.54 | 0.00 | 0.00 | 0.00 |
| R-HSA-9664565 | Signaling by ERBB2 KD Mutants | -0.58 | -0.79 | -0.39 | 0.00 | 0.00 | 0.00 |
| R-HSA-9665230 | Drug resistance in ERBB2 KD mutants | -0.57 | -0.93 | -0.54 | 0.00 | 0.00 | 0.00 |
| R-HSA-9665233 | Resistance of ERBB2 KD mutants to trastuzumab | -0.57 | -0.93 | -0.54 | 0.00 | 0.00 | 0.00 |
| R-HSA-9665244 | Resistance of ERBB2 KD mutants to sapitinib | -0.57 | -0.93 | -0.54 | 0.00 | 0.00 | 0.00 |
| R-HSA-9665245 | Resistance of ERBB2 KD mutants to tesevatinib | -0.57 | -0.93 | -0.54 | 0.00 | 0.00 | 0.00 |
| R-HSA-9665246 | Resistance of ERBB2 KD mutants to neratinib | -0.57 | -0.93 | -0.54 | 0.00 | 0.00 | 0.00 |
| R-HSA-9665247 | Resistance of ERBB2 KD mutants to osimertinib | -0.57 | -0.93 | -0.54 | 0.00 | 0.00 | 0.00 |
| R-HSA-9665249 | Resistance of ERBB2 KD mutants to afatinib | -0.57 | -0.93 | -0.54 | 0.00 | 0.00 | 0.00 |
| R-HSA-9665250 | Resistance of ERBB2 KD mutants to AEE788 | -0.57 | -0.93 | -0.54 | 0.00 | 0.00 | 0.00 |
| R-HSA-9665251 | Resistance of ERBB2 KD mutants to lapatinib | -0.57 | -0.93 | -0.54 | 0.00 | 0.00 | 0.00 |
| R-HSA-9665348 | Signaling by ERBB2 ECD mutants | -0.58 | -0.79 | -0.61 | 0.00 | 0.00 | 0.00 |
| R-HSA-9665686 | Signaling by ERBB2 TMD/JMD mutants | -0.58 | -0.79 | -0.32 | 0.00 | 0.00 | 0.00 |
| R-HSA-9665737 | Drug resistance in ERBB2 TMD/JMD mutants | -0.57 | -0.93 | -0.54 | 0.00 | 0.00 | 0.00 |
| R-HSA-9706574 | RHOBTB GTPase Cycle | -0.10 | -0.25 | 0.84 | 0.72 | 0.00 | 0.00 |
| R-HSA-2142789 | Ubiquinol biosynthesis | 0.22 | -0.30 | -0.49 | 0.56 | 0.00 | 0.00 |
| R-HSA-8978934 | Metabolism of cofactors | 0.15 | -0.30 | -0.22 | 0.64 | 0.18 | 0.15 |
| R-HSA-180024 | DARPP-32 events | 0.01 | -0.25 | -0.36 | 0.72 | 0.00 | 0.00 |
| R-HSA-2025928 | Calcineurin activates NFAT | -0.78 | 0.83 | -0.29 | 0.00 | 0.00 | 0.00 |
| R-HSA-2871809 | FCERI mediated Ca+2 mobilization | -0.15 | -0.09 | -0.38 | 0.74 | 0.00 | 0.00 |
| R-HSA-5607763 | CLEC7A (Dectin-1) induces NFAT activation | -0.01 | 0.25 | -0.42 | 0.72 | 0.00 | 0.00 |
| R-HSA-8849175 | Threonine catabolism | -0.16 | -0.19 | -0.25 | 0.73 | 0.13 | 0.14 |
| R-HSA-983170 | Antigen Presentation: Folding, assembly and peptide loading of class I MHC | 0.06 | 0.04 | -0.47 | 0.74 | 0.00 | 0.00 |
| R-HSA-264870 | Caspase-mediated cleavage of cytoskeletal proteins | 0.09 | -0.11 | -0.53 | 0.74 | 0.00 | 0.00 |
| R-HSA-6785807 | Interleukin-4 and Interleukin-13 signaling | 0.11 | 0.07 | -0.27 | 0.74 | 0.08 | 0.08 |
| R-HSA-9013422 | RHOBTB1 GTPase cycle | 0.09 | -0.36 | 0.89 | 0.60 | 0.00 | 0.00 |
| R-HSA-2168880 | Scavenging of heme from plasma | 0.24 | 0.17 | -0.05 | 0.72 | 0.36 | 0.39 |
| R-HSA-917729 | Endosomal Sorting Complex Required For Transport (ESCRT) | 0.25 | -0.15 | -0.25 | 0.68 | 0.08 | 0.16 |
| R-HSA-9615017 | FOXO-mediated transcription of oxidative stress, metabolic and neuronal genes | -0.25 | -0.69 | -0.33 | 0.02 | 0.01 | 0.00 |
| R-HSA-168275 | Entry of Influenza Virion into Host Cell via Endocytosis | -0.57 | -0.08 | -0.08 | 0.19 | 0.04 | 0.40 |
| R-HSA-177504 | Retrograde neurotrophin signalling | 0.12 | 0.30 | -0.15 | 0.71 | 0.32 | 0.22 |
| R-HSA-8866427 | VLDLR internalisation and degradation | 0.17 | 0.14 | -0.19 | 0.74 | 0.25 | 0.26 |
| R-HSA-8964038 | LDL clearance | 0.13 | 0.13 | -0.16 | 0.74 | 0.31 | 0.32 |
| R-HSA-8964043 | Plasma lipoprotein clearance | 0.13 | -0.02 | -0.16 | 0.74 | 0.31 | 0.35 |
| R-HSA-112311 | Neurotransmitter clearance | 0.16 | 0.00 | -0.12 | 0.74 | 0.35 | 0.38 |
| R-HSA-379397 | Enzymatic degradation of dopamine by COMT | 0.16 | 0.00 | -0.26 | 0.74 | 0.09 | 0.12 |
| R-HSA-379398 | Enzymatic degradation of Dopamine by monoamine oxidase | 0.16 | 0.00 | -0.23 | 0.74 | 0.15 | 0.20 |
| R-HSA-379401 | Dopamine clearance from the synaptic cleft | 0.16 | 0.00 | -0.08 | 0.74 | 0.37 | 0.40 |
| R-HSA-177929 | Signaling by EGFR | -0.45 | -0.12 | -0.43 | 0.53 | 0.00 | 0.00 |
| R-HSA-182971 | EGFR downregulation | -0.36 | 0.03 | -0.32 | 0.64 | 0.00 | 0.02 |
| R-HSA-6807004 | Negative regulation of MET activity | -0.60 | -0.15 | -0.47 | 0.13 | 0.00 | 0.00 |
| R-HSA-8875360 | InlB-mediated entry of Listeria monocytogenes into host cell | -0.60 | -0.15 | -0.42 | 0.13 | 0.00 | 0.00 |
| R-HSA-8876384 | Listeria monocytogenes entry into host cells | -0.70 | -0.28 | -0.42 | 0.01 | 0.00 | 0.00 |
| R-HSA-983695 | Antigen activates B Cell Receptor (BCR) leading to generation of second messengers | 0.16 | -0.54 | -0.46 | 0.09 | 0.00 | 0.00 |
| R-HSA-525793 | Myogenesis | -0.40 | -0.60 | -0.36 | 0.06 | 0.00 | 0.00 |
| R-HSA-2173796 | SMAD2/SMAD3:SMAD4 heterotrimer regulates transcription | -0.58 | -0.18 | -0.34 | 0.19 | 0.00 | 0.01 |
| R-HSA-168277 | Influenza Virus Induced Apoptosis | 1.00 | 0.83 | 0.08 | 0.00 | 0.00 | 0.00 |
| R-HSA-180897 | Vpr-mediated induction of apoptosis by mitochondrial outer membrane permeabilization | 0.60 | 0.52 | -0.07 | 0.01 | 0.02 | 0.08 |
| R-HSA-425397 | Transport of vitamins, nucleosides, and related molecules | 0.43 | 0.37 | -0.15 | 0.42 | 0.10 | 0.14 |
| R-HSA-83936 | Transport of nucleosides and free purine and pyrimidine bases across the plasma membrane | 0.60 | 0.52 | 0.00 | 0.01 | 0.03 | 0.12 |
| R-HSA-381340 | Transcriptional regulation of white adipocyte differentiation | 0.34 | 0.33 | -0.38 | 0.60 | 0.00 | 0.00 |
| R-HSA-9024446 | NR1H2 and NR1H3-mediated signaling | 0.02 | 0.14 | -0.34 | 0.74 | 0.01 | 0.00 |
| R-HSA-9031528 | NR1H2 & NR1H3 regulate gene expression linked to triglyceride lipolysis in adipose | -1.01 | 0.05 | -0.26 | 0.00 | 0.00 | 0.13 |
| R-HSA-2142688 | Synthesis of 5-eicosatetraenoic acids | 0.34 | -0.22 | -0.30 | 0.49 | 0.01 | 0.04 |
| R-HSA-2142712 | Synthesis of 12-eicosatetraenoic acid derivatives | 0.71 | 0.12 | -0.11 | 0.01 | 0.00 | 0.37 |
| R-HSA-2142770 | Synthesis of 15-eicosatetraenoic acid derivatives | 0.71 | 0.12 | -0.39 | 0.01 | 0.00 | 0.00 |
| R-HSA-2672351 | Stimuli-sensing channels | -0.03 | 0.21 | -0.17 | 0.73 | 0.32 | 0.25 |
| R-HSA-6798163 | Choline catabolism | 0.33 | -0.01 | -0.33 | 0.67 | 0.00 | 0.02 |
| R-HSA-71064 | Lysine catabolism | 0.17 | -0.47 | -0.46 | 0.22 | 0.00 | 0.00 |
| R-HSA-8981607 | Intracellular oxygen transport | -0.73 | -0.25 | 0.07 | 0.01 | 0.00 | 0.36 |
| R-HSA-430039 | mRNA decay by 5' to 3' exoribonuclease | -0.29 | 0.48 | -0.26 | 0.05 | 0.05 | 0.00 |
| R-HSA-373076 | Class A/1 (Rhodopsin-like receptors) | 0.23 | 0.14 | 0.07 | 0.73 | 0.36 | 0.39 |
| R-HSA-375276 | Peptide ligand-binding receptors | 0.23 | 0.14 | 0.11 | 0.73 | 0.34 | 0.38 |
| R-HSA-444473 | Formyl peptide receptors bind formyl peptides and many other ligands | 0.44 | 0.35 | -0.30 | 0.43 | 0.00 | 0.00 |
| R-HSA-196807 | Nicotinate metabolism | 0.18 | -0.08 | -0.41 | 0.73 | 0.00 | 0.00 |
| R-HSA-197264 | Nicotinamide salvaging | 0.11 | -0.40 | -0.42 | 0.52 | 0.00 | 0.00 |
| R-HSA-3928663 | EPHA-mediated growth cone collapse | 0.01 | 0.02 | -0.38 | 0.74 | 0.00 | 0.00 |
| R-HSA-400685 | Sema4D in semaphorin signaling | 0.10 | 0.05 | -0.46 | 0.74 | 0.00 | 0.00 |
| R-HSA-416572 | Sema4D induced cell migration and growth-cone collapse | 0.01 | 0.02 | -0.51 | 0.74 | 0.00 | 0.00 |
| R-HSA-9662834 | CD163 mediating an anti-inflammatory response | 0.52 | -0.47 | -0.40 | 0.00 | 0.00 | 0.00 |
| R-HSA-71737 | Pyrophosphate hydrolysis | 0.51 | -0.34 | -0.36 | 0.01 | 0.00 | 0.00 |
| R-HSA-8963684 | Tyrosine catabolism | 0.28 | 0.35 | 0.02 | 0.63 | 0.35 | 0.33 |
| R-HSA-8963691 | Phenylalanine and tyrosine metabolism | 0.49 | 0.13 | -0.08 | 0.44 | 0.11 | 0.39 |
| R-HSA-1980143 | Signaling by NOTCH1 | 0.32 | 0.55 | -0.34 | 0.22 | 0.00 | 0.00 |
| R-HSA-2122947 | NOTCH1 Intracellular Domain Regulates Transcription | 0.13 | 0.39 | -0.32 | 0.63 | 0.02 | 0.00 |
| R-HSA-2644602 | Signaling by NOTCH1 PEST Domain Mutants in Cancer | 0.13 | 0.39 | -0.33 | 0.63 | 0.01 | 0.00 |
| R-HSA-2644603 | Signaling by NOTCH1 in Cancer | 0.13 | 0.39 | -0.33 | 0.63 | 0.01 | 0.00 |
| R-HSA-2644606 | Constitutive Signaling by NOTCH1 PEST Domain Mutants | 0.13 | 0.39 | -0.33 | 0.63 | 0.01 | 0.00 |
| R-HSA-2894858 | Signaling by NOTCH1 HD+PEST Domain Mutants in Cancer | 0.13 | 0.39 | -0.33 | 0.63 | 0.01 | 0.00 |
| R-HSA-2894862 | Constitutive Signaling by NOTCH1 HD+PEST Domain Mutants | 0.13 | 0.39 | -0.33 | 0.63 | 0.01 | 0.00 |
| R-HSA-3214815 | HDACs deacetylate histones | -0.05 | -0.03 | 0.03 | 0.74 | 0.38 | 0.40 |
| R-HSA-3247509 | Chromatin modifying enzymes | -0.06 | -0.09 | -0.30 | 0.74 | 0.04 | 0.05 |
| R-HSA-350054 | Notch-HLH transcription pathway | 1.01 | 0.32 | -0.38 | 0.00 | 0.00 | 0.00 |
| R-HSA-3899300 | SUMOylation of transcription cofactors | 0.76 | 0.34 | -0.42 | 0.00 | 0.00 | 0.00 |
| R-HSA-4839726 | Chromatin organization | -0.06 | -0.09 | -0.30 | 0.74 | 0.04 | 0.05 |
| R-HSA-8986944 | Transcriptional Regulation by MECP2 | 0.36 | 0.02 | -0.24 | 0.65 | 0.03 | 0.17 |
| R-HSA-9005891 | Loss of function of MECP2 in Rett syndrome | 0.97 | -0.18 | -0.32 | 0.00 | 0.00 | 0.02 |
| R-HSA-9005895 | Pervasive developmental disorders | 0.97 | -0.18 | -0.32 | 0.00 | 0.00 | 0.02 |
| R-HSA-9022537 | Loss of MECP2 binding ability to the NCoR/SMRT complex | 1.01 | 0.32 | -0.42 | 0.00 | 0.00 | 0.00 |
| R-HSA-9022692 | Regulation of MECP2 expression and activity | 0.69 | -0.34 | -0.39 | 0.00 | 0.00 | 0.00 |
| R-HSA-9029569 | NR1H3 & NR1H2 regulate gene expression linked to cholesterol transport and efflux | 0.39 | 0.06 | -0.36 | 0.63 | 0.00 | 0.00 |
| R-HSA-9623433 | NR1H2 & NR1H3 regulate gene expression to control bile acid homeostasis | 1.01 | 0.32 | -0.36 | 0.00 | 0.00 | 0.00 |
| R-HSA-9675151 | Disorders of Developmental Biology | 0.97 | -0.18 | -0.32 | 0.00 | 0.00 | 0.02 |
| R-HSA-9697154 | Disorders of Nervous System Development | 0.97 | -0.18 | -0.32 | 0.00 | 0.00 | 0.02 |
| R-HSA-381033 | ATF6 (ATF6-alpha) activates chaperones | 0.05 | 0.00 | -0.16 | 0.74 | 0.34 | 0.35 |
| R-HSA-381183 | ATF6 (ATF6-alpha) activates chaperone genes | 0.05 | 0.00 | -0.09 | 0.74 | 0.38 | 0.39 |
| R-HSA-446107 | Type I hemidesmosome assembly | -0.09 | 0.33 | -0.21 | 0.65 | 0.23 | 0.07 |
| R-HSA-193634 | Axonal growth inhibition (RHOA activation) | -0.03 | 0.02 | 0.02 | 0.74 | 0.38 | 0.40 |
| R-HSA-193697 | p75NTR regulates axonogenesis | -0.03 | 0.02 | 0.00 | 0.74 | 0.38 | 0.40 |
| R-HSA-114604 | GPVI-mediated activation cascade | -0.13 | 0.42 | -0.33 | 0.44 | 0.01 | 0.00 |
| R-HSA-3000170 | Syndecan interactions | -0.47 | 0.45 | -0.49 | 0.00 | 0.00 | 0.00 |
| R-HSA-430116 | GP1b-IX-V activation signalling | -0.33 | -0.01 | -0.42 | 0.68 | 0.00 | 0.00 |
| R-HSA-75892 | Platelet Adhesion to exposed collagen | -0.46 | 0.82 | -0.31 | 0.00 | 0.00 | 0.00 |
| R-HSA-8874081 | MET activates PTK2 signaling | -0.37 | 0.37 | -0.50 | 0.11 | 0.00 | 0.00 |
| R-HSA-3906995 | Diseases associated with O-glycosylation of proteins | -0.04 | 0.68 | -0.37 | 0.01 | 0.00 | 0.00 |
| R-HSA-5083628 | Defective POMGNT1 causes MDDGA3, MDDGB3 and MDDGC3 | -0.29 | 0.68 | -0.62 | 0.00 | 0.00 | 0.00 |
| R-HSA-5083629 | Defective POMT2 causes MDDGA2, MDDGB2 and MDDGC2 | -0.29 | 0.68 | -0.41 | 0.00 | 0.00 | 0.00 |
| R-HSA-5083633 | Defective POMT1 causes MDDGA1, MDDGB1 and MDDGC1 | -0.29 | 0.68 | -0.41 | 0.00 | 0.00 | 0.00 |
| R-HSA-5173105 | O-linked glycosylation | -0.04 | 0.68 | -0.33 | 0.01 | 0.02 | 0.00 |
| R-HSA-9619665 | EGR2 and SOX10-mediated initiation of Schwann cell myelination | -0.45 | 0.26 | -0.34 | 0.13 | 0.00 | 0.00 |
| R-HSA-9615933 | Postmitotic nuclear pore complex (NPC) reformation | 0.37 | -0.26 | -0.45 | 0.33 | 0.00 | 0.00 |
| R-HSA-2032785 | YAP1- and WWTR1 (TAZ)-stimulated gene expression | -0.61 | -0.76 | -0.43 | 0.00 | 0.00 | 0.00 |
| R-HSA-5578768 | Physiological factors | -0.61 | -0.76 | -0.46 | 0.00 | 0.00 | 0.00 |
| R-HSA-110362 | POLB-Dependent Long Patch Base Excision Repair | 0.50 | 0.26 | -0.19 | 0.39 | 0.02 | 0.17 |
| R-HSA-110373 | Resolution of AP sites via the multiple-nucleotide patch replacement pathway | 0.50 | 0.26 | -0.25 | 0.39 | 0.00 | 0.05 |
| R-HSA-73884 | Base Excision Repair | -0.07 | 0.01 | -0.09 | 0.74 | 0.38 | 0.40 |
| R-HSA-73933 | Resolution of Abasic Sites (AP sites) | 0.50 | 0.26 | -0.26 | 0.39 | 0.00 | 0.03 |
| R-HSA-1474151 | Tetrahydrobiopterin (BH4) synthesis, recycling, salvage and regulation | -0.06 | -0.43 | -0.09 | 0.57 | 0.38 | 0.22 |
| R-HSA-168928 | DDX58/IFIH1-mediated induction of interferon-alpha/beta | -0.17 | -0.20 | -0.41 | 0.73 | 0.00 | 0.00 |
| R-HSA-202131 | Metabolism of nitric oxide: NOS3 activation and regulation | 0.12 | 0.14 | -0.23 | 0.74 | 0.18 | 0.16 |
| R-HSA-203615 | eNOS activation | 0.15 | -0.02 | -0.22 | 0.74 | 0.18 | 0.23 |
| R-HSA-3000484 | Scavenging by Class F Receptors | 0.26 | 0.00 | -0.15 | 0.72 | 0.26 | 0.36 |
| R-HSA-3371511 | HSF1 activation | -0.05 | -0.72 | -0.27 | 0.01 | 0.11 | 0.00 |
| R-HSA-3371568 | Attenuation phase | -0.38 | -0.55 | -0.14 | 0.16 | 0.18 | 0.06 |
| R-HSA-3371571 | HSF1-dependent transactivation | -0.21 | -0.69 | -0.28 | 0.02 | 0.06 | 0.00 |
| R-HSA-5218920 | VEGFR2 mediated vascular permeability | -0.33 | -0.32 | -0.36 | 0.62 | 0.00 | 0.00 |
| R-HSA-5336415 | Uptake and function of diphtheria toxin | -0.09 | -0.26 | -0.24 | 0.72 | 0.17 | 0.13 |
| R-HSA-5339562 | Uptake and actions of bacterial toxins | 0.01 | -0.23 | -0.16 | 0.73 | 0.34 | 0.31 |
| R-HSA-5601884 | PIWI-interacting RNA (piRNA) biogenesis | -0.26 | -0.97 | -0.09 | 0.00 | 0.34 | 0.00 |
| R-HSA-9018519 | Estrogen-dependent gene expression | -0.62 | -0.30 | -0.15 | 0.08 | 0.01 | 0.28 |
| R-HSA-112040 | G-protein mediated events | 0.43 | 0.00 | -0.48 | 0.53 | 0.00 | 0.00 |
| R-HSA-112043 | PLC beta mediated events | 0.37 | -0.20 | -0.50 | 0.45 | 0.00 | 0.00 |
| R-HSA-399997 | Acetylcholine regulates insulin secretion | 0.17 | 0.60 | -0.31 | 0.16 | 0.02 | 0.00 |
| R-HSA-400451 | Free fatty acids regulate insulin secretion | 0.67 | 0.76 | -0.30 | 0.00 | 0.00 | 0.00 |
| R-HSA-434316 | Fatty Acids bound to GPR40 (FFAR1) regulate insulin secretion | 0.69 | 1.00 | -0.31 | 0.00 | 0.00 | 0.00 |
| R-HSA-163282 | Mitochondrial transcription initiation | -0.34 | 0.41 | -0.46 | 0.09 | 0.00 | 0.00 |
| R-HSA-75944 | Transcription from mitochondrial promoters | -0.34 | 0.41 | -0.47 | 0.09 | 0.00 | 0.00 |
| R-HSA-5689901 | Metalloprotease DUBs | -0.47 | 0.06 | -0.08 | 0.38 | 0.15 | 0.40 |
| R-HSA-5693532 | DNA Double-Strand Break Repair | -0.35 | 0.40 | -0.28 | 0.10 | 0.02 | 0.00 |
| R-HSA-5693538 | Homology Directed Repair | -0.54 | 0.61 | -0.22 | 0.00 | 0.01 | 0.00 |
| R-HSA-5693565 | Recruitment and ATM-mediated phosphorylation of repair and signaling proteins at DNA double strand breaks | -0.35 | 0.40 | -0.15 | 0.10 | 0.21 | 0.13 |
| R-HSA-5693567 | HDR through Homologous Recombination (HRR) or Single Strand Annealing (SSA) | -0.54 | 0.61 | -0.21 | 0.00 | 0.01 | 0.00 |
| R-HSA-5693571 | Nonhomologous End-Joining (NHEJ) | -0.47 | 0.65 | -0.16 | 0.00 | 0.07 | 0.00 |
| R-HSA-5693606 | DNA Double Strand Break Response | -0.35 | 0.40 | -0.16 | 0.10 | 0.19 | 0.10 |
| R-HSA-5693607 | Processing of DNA double-strand break ends | -0.54 | 0.61 | -0.19 | 0.00 | 0.02 | 0.00 |
| R-HSA-140834 | Extrinsic Pathway of Fibrin Clot Formation | 0.57 | 0.17 | -0.35 | 0.22 | 0.00 | 0.00 |
| R-HSA-159740 | Gamma-carboxylation of protein precursors | 0.46 | 0.32 | -0.28 | 0.42 | 0.00 | 0.01 |
| R-HSA-159763 | Transport of gamma-carboxylated protein precursors from the endoplasmic reticulum to the Golgi apparatus | 0.46 | 0.32 | -0.20 | 0.42 | 0.03 | 0.10 |
| R-HSA-159782 | Removal of aminoterminal propeptides from gamma-carboxylated proteins | 0.46 | 0.32 | -0.24 | 0.42 | 0.01 | 0.04 |
| R-HSA-159854 | Gamma-carboxylation, transport, and amino-terminal cleavage of proteins | 0.46 | 0.32 | -0.31 | 0.42 | 0.00 | 0.00 |
| R-HSA-163841 | Gamma carboxylation, hypusine formation and arylsulfatase activation | 0.43 | 0.17 | -0.36 | 0.56 | 0.00 | 0.00 |
| R-HSA-9662001 | Defective factor VIII causes hemophilia A | 0.59 | 0.24 | -0.21 | 0.16 | 0.00 | 0.14 |
| R-HSA-9668250 | Defective factor IX causes hemophilia B | 0.57 | 0.17 | -0.04 | 0.22 | 0.05 | 0.39 |
| R-HSA-9672383 | Defective factor IX causes thrombophilia | 0.57 | 0.17 | -0.14 | 0.22 | 0.02 | 0.34 |
| R-HSA-9672396 | Defective cofactor function of FVIIIa variant | 0.57 | 0.17 | -0.14 | 0.22 | 0.02 | 0.34 |
| R-HSA-9673202 | Defective F9 variant does not activate FX | 0.57 | 0.17 | -0.14 | 0.22 | 0.02 | 0.34 |
| R-HSA-9673218 | Defective F9 secretion | 0.88 | 0.20 | 0.10 | 0.00 | 0.00 | 0.38 |
| R-HSA-9673221 | Defective F9 activation | 0.88 | 0.20 | 0.29 | 0.00 | 0.00 | 0.06 |
| R-HSA-9673240 | Defective gamma-carboxylation of F9 | 0.88 | 0.20 | -0.41 | 0.00 | 0.00 | 0.00 |
| R-HSA-1482788 | Acyl chain remodelling of PC | -0.50 | -0.09 | -0.04 | 0.41 | 0.13 | 0.40 |
| R-HSA-1482839 | Acyl chain remodelling of PE | -0.50 | -0.09 | -0.10 | 0.41 | 0.10 | 0.39 |
| R-HSA-4043911 | Defective PMM2 causes PMM2-CDG (CDG-1a) | -0.77 | -0.99 | -0.56 | 0.00 | 0.00 | 0.00 |
| R-HSA-446205 | Synthesis of GDP-mannose | 0.09 | -0.31 | -0.29 | 0.67 | 0.05 | 0.03 |
| R-HSA-5609975 | Diseases associated with glycosylation precursor biosynthesis | -0.18 | -0.47 | -0.37 | 0.50 | 0.00 | 0.00 |
| R-HSA-977347 | Serine biosynthesis | 0.83 | 0.11 | -0.44 | 0.00 | 0.00 | 0.00 |
| R-HSA-9617629 | Regulation of FOXO transcriptional activity by acetylation | -0.29 | -0.95 | -0.55 | 0.00 | 0.00 | 0.00 |
| R-HSA-1236977 | Endosomal/Vacuolar pathway | 0.17 | -0.07 | -0.41 | 0.73 | 0.00 | 0.00 |
| R-HSA-2172127 | DAP12 interactions | -0.09 | -0.04 | -0.40 | 0.74 | 0.00 | 0.00 |
| R-HSA-2424491 | DAP12 signaling | -0.09 | -0.04 | -0.43 | 0.74 | 0.00 | 0.00 |
| R-HSA-877300 | Interferon gamma signaling | 0.29 | 0.30 | -0.36 | 0.66 | 0.00 | 0.00 |
| R-HSA-909733 | Interferon alpha/beta signaling | 0.54 | 0.07 | -0.40 | 0.29 | 0.00 | 0.00 |
| R-HSA-390450 | Folding of actin by CCT/TriC | -0.12 | 0.04 | -0.13 | 0.74 | 0.35 | 0.38 |
| R-HSA-390471 | Association of TriC/CCT with target proteins during biosynthesis | -0.15 | 0.12 | -0.35 | 0.73 | 0.01 | 0.00 |
| R-HSA-5620920 | Cargo trafficking to the periciliary membrane | -0.03 | -0.02 | -0.45 | 0.74 | 0.00 | 0.00 |
| R-HSA-5620922 | BBSome-mediated cargo-targeting to cilium | 0.06 | 0.09 | -0.41 | 0.74 | 0.00 | 0.00 |
| R-HSA-112399 | IRS-mediated signalling | -0.59 | -0.66 | -0.17 | 0.00 | 0.01 | 0.00 |
| R-HSA-112412 | SOS-mediated signalling | -0.59 | -0.66 | -0.59 | 0.00 | 0.00 | 0.00 |
| R-HSA-1169092 | Activation of RAS in B cells | -0.72 | -0.56 | -0.49 | 0.00 | 0.00 | 0.00 |
| R-HSA-1226099 | Signaling by FGFR in disease | 0.01 | -0.16 | -0.30 | 0.74 | 0.05 | 0.05 |
| R-HSA-1236394 | Signaling by ERBB4 | -0.42 | -0.42 | -0.30 | 0.36 | 0.00 | 0.01 |
| R-HSA-1250196 | SHC1 events in ERBB2 signaling | -0.59 | -0.66 | -0.35 | 0.00 | 0.00 | 0.00 |
| R-HSA-1250347 | SHC1 events in ERBB4 signaling | -0.59 | -0.66 | -0.21 | 0.00 | 0.00 | 0.00 |
| R-HSA-1433557 | Signaling by SCF-KIT | 0.17 | -0.02 | -0.29 | 0.74 | 0.04 | 0.07 |
| R-HSA-179812 | GRB2 events in EGFR signaling | -0.59 | -0.66 | -0.27 | 0.00 | 0.00 | 0.00 |
| R-HSA-180336 | SHC1 events in EGFR signaling | -0.59 | -0.66 | -0.30 | 0.00 | 0.00 | 0.00 |
| R-HSA-1963640 | GRB2 events in ERBB2 signaling | -0.59 | -0.66 | -0.22 | 0.00 | 0.00 | 0.00 |
| R-HSA-210993 | Tie2 Signaling | -0.59 | -0.66 | -0.66 | 0.00 | 0.00 | 0.00 |
| R-HSA-2179392 | EGFR Transactivation by Gastrin | -0.59 | -0.66 | -0.34 | 0.00 | 0.00 | 0.00 |
| R-HSA-2404192 | Signaling by Type 1 Insulin-like Growth Factor 1 Receptor (IGF1R) | -0.59 | -0.66 | -0.19 | 0.00 | 0.01 | 0.00 |
| R-HSA-2428924 | IGF1R signaling cascade | -0.59 | -0.66 | -0.17 | 0.00 | 0.01 | 0.00 |
| R-HSA-2428928 | IRS-related events triggered by IGF1R | -0.59 | -0.66 | -0.17 | 0.00 | 0.01 | 0.01 |
| R-HSA-2428933 | SHC-related events triggered by IGF1R | -0.59 | -0.66 | -0.56 | 0.00 | 0.00 | 0.00 |
| R-HSA-2871796 | FCERI mediated MAPK activation | -0.01 | -0.35 | -0.41 | 0.67 | 0.00 | 0.00 |
| R-HSA-442742 | CREB1 phosphorylation through NMDA receptor-mediated activation of RAS signaling | -0.06 | -0.52 | -0.25 | 0.36 | 0.14 | 0.01 |
| R-HSA-442982 | Ras activation upon Ca2+ influx through NMDA receptor | -0.48 | -0.58 | -0.22 | 0.04 | 0.02 | 0.01 |
| R-HSA-5218921 | VEGFR2 mediated cell proliferation | 0.02 | -0.44 | -0.35 | 0.52 | 0.01 | 0.00 |
| R-HSA-5621575 | CD209 (DC-SIGN) signaling | -0.10 | -0.31 | -0.43 | 0.71 | 0.00 | 0.00 |
| R-HSA-5654687 | Downstream signaling of activated FGFR1 | -0.59 | -0.66 | -0.20 | 0.00 | 0.01 | 0.00 |
| R-HSA-5654688 | SHC-mediated cascade:FGFR1 | -0.59 | -0.66 | -0.04 | 0.00 | 0.04 | 0.01 |
| R-HSA-5654693 | FRS-mediated FGFR1 signaling | -0.59 | -0.66 | -0.07 | 0.00 | 0.04 | 0.01 |
| R-HSA-5654696 | Downstream signaling of activated FGFR2 | -0.59 | -0.66 | -0.21 | 0.00 | 0.00 | 0.00 |
| R-HSA-5654699 | SHC-mediated cascade:FGFR2 | -0.59 | -0.66 | -0.06 | 0.00 | 0.04 | 0.01 |
| R-HSA-5654700 | FRS-mediated FGFR2 signaling | -0.59 | -0.66 | -0.09 | 0.00 | 0.03 | 0.01 |
| R-HSA-5654704 | SHC-mediated cascade:FGFR3 | -0.59 | -0.66 | -0.15 | 0.00 | 0.02 | 0.01 |
| R-HSA-5654706 | FRS-mediated FGFR3 signaling | -0.59 | -0.66 | -0.18 | 0.00 | 0.01 | 0.00 |
| R-HSA-5654708 | Downstream signaling of activated FGFR3 | -0.59 | -0.66 | -0.29 | 0.00 | 0.00 | 0.00 |
| R-HSA-5654712 | FRS-mediated FGFR4 signaling | -0.59 | -0.66 | -0.07 | 0.00 | 0.04 | 0.01 |
| R-HSA-5654716 | Downstream signaling of activated FGFR4 | -0.59 | -0.66 | -0.20 | 0.00 | 0.00 | 0.00 |
| R-HSA-5654719 | SHC-mediated cascade:FGFR4 | -0.59 | -0.66 | -0.04 | 0.00 | 0.04 | 0.01 |
| R-HSA-5654736 | Signaling by FGFR1 | -0.07 | -0.25 | -0.26 | 0.73 | 0.12 | 0.09 |
| R-HSA-5654741 | Signaling by FGFR3 | -0.07 | -0.25 | -0.30 | 0.73 | 0.05 | 0.03 |
| R-HSA-5654743 | Signaling by FGFR4 | -0.07 | -0.25 | -0.24 | 0.73 | 0.17 | 0.14 |
| R-HSA-5655253 | Signaling by FGFR2 in disease | -0.59 | -0.66 | -0.18 | 0.00 | 0.01 | 0.00 |
| R-HSA-5655291 | Signaling by FGFR4 in disease | -0.59 | -0.66 | -0.49 | 0.00 | 0.00 | 0.00 |
| R-HSA-5655302 | Signaling by FGFR1 in disease | 0.01 | -0.16 | -0.37 | 0.74 | 0.00 | 0.00 |
| R-HSA-5655332 | Signaling by FGFR3 in disease | -0.59 | -0.66 | -0.22 | 0.00 | 0.00 | 0.00 |
| R-HSA-5673000 | RAF activation | -0.08 | -0.27 | -0.39 | 0.72 | 0.00 | 0.00 |
| R-HSA-5674135 | MAP2K and MAPK activation | -0.07 | -0.12 | -0.27 | 0.74 | 0.10 | 0.10 |
| R-HSA-5675221 | Negative regulation of MAPK pathway | -0.03 | -0.26 | -0.38 | 0.72 | 0.00 | 0.00 |
| R-HSA-6802946 | Signaling by moderate kinase activity BRAF mutants | -0.11 | -0.19 | -0.28 | 0.74 | 0.07 | 0.07 |
| R-HSA-6802948 | Signaling by high-kinase activity BRAF mutants | -0.09 | -0.16 | -0.30 | 0.74 | 0.05 | 0.05 |
| R-HSA-6802949 | Signaling by RAS mutants | -0.11 | -0.19 | -0.28 | 0.74 | 0.07 | 0.07 |
| R-HSA-6802953 | RAS signaling downstream of NF1 loss-of-function variants | -0.72 | -0.56 | -0.49 | 0.00 | 0.00 | 0.00 |
| R-HSA-6802955 | Paradoxical activation of RAF signaling by kinase inactive BRAF | -0.11 | -0.19 | -0.28 | 0.74 | 0.07 | 0.07 |
| R-HSA-74751 | Insulin receptor signalling cascade | -0.01 | -0.54 | -0.17 | 0.26 | 0.33 | 0.04 |
| R-HSA-74752 | Signaling by Insulin receptor | -0.17 | -0.35 | -0.18 | 0.67 | 0.27 | 0.18 |
| R-HSA-881907 | Gastrin-CREB signalling pathway via PKC and MAPK | -0.01 | -0.54 | -0.23 | 0.26 | 0.21 | 0.01 |
| R-HSA-8851805 | MET activates RAS signaling | -0.59 | -0.66 | -0.55 | 0.00 | 0.00 | 0.00 |
| R-HSA-8853334 | Signaling by FGFR3 fusions in cancer | -0.59 | -0.66 | -0.63 | 0.00 | 0.00 | 0.00 |
| R-HSA-8853338 | Signaling by FGFR3 point mutants in cancer | -0.59 | -0.66 | -0.22 | 0.00 | 0.00 | 0.00 |
| R-HSA-8951936 | RUNX3 regulates p14-ARF | -0.71 | 0.15 | -0.43 | 0.00 | 0.00 | 0.00 |
| R-HSA-9006115 | Signaling by NTRK2 (TRKB) | -0.29 | -0.25 | -0.49 | 0.68 | 0.00 | 0.00 |
| R-HSA-9026519 | Activated NTRK2 signals through RAS | -0.59 | -0.66 | -0.47 | 0.00 | 0.00 | 0.00 |
| R-HSA-9028731 | Activated NTRK2 signals through FRS2 and FRS3 | -0.59 | -0.66 | -0.47 | 0.00 | 0.00 | 0.00 |
| R-HSA-9034015 | Signaling by NTRK3 (TRKC) | -0.59 | -0.66 | -0.50 | 0.00 | 0.00 | 0.00 |
| R-HSA-9034864 | Activated NTRK3 signals through RAS | -0.59 | -0.66 | -0.48 | 0.00 | 0.00 | 0.00 |
| R-HSA-9607240 | FLT3 Signaling | -0.32 | -0.22 | -0.34 | 0.68 | 0.00 | 0.01 |
| R-HSA-9634635 | Estrogen-stimulated signaling through PRKCZ | 0.12 | -0.19 | -0.21 | 0.72 | 0.22 | 0.22 |
| R-HSA-9649913 | RAS GTPase cycle mutants | -0.72 | -0.56 | -0.44 | 0.00 | 0.00 | 0.00 |
| R-HSA-9649948 | Signaling downstream of RAS mutants | -0.11 | -0.19 | -0.28 | 0.74 | 0.07 | 0.07 |
| R-HSA-9656223 | Signaling by RAF1 mutants | -0.08 | -0.19 | -0.31 | 0.74 | 0.04 | 0.03 |
| R-HSA-9669938 | Signaling by KIT in disease | 0.01 | -0.16 | -0.62 | 0.74 | 0.00 | 0.00 |
| R-HSA-9670439 | Signaling by phosphorylated juxtamembrane, extracellular and kinase domain KIT mutants | 0.01 | -0.16 | -0.62 | 0.74 | 0.00 | 0.00 |
| R-HSA-9671555 | Signaling by PDGFR in disease | 0.14 | -0.23 | -0.65 | 0.70 | 0.00 | 0.00 |
| R-HSA-9673767 | Signaling by PDGFRA transmembrane, juxtamembrane and kinase domain mutants | 0.01 | -0.16 | -0.71 | 0.74 | 0.00 | 0.00 |
| R-HSA-9673770 | Signaling by PDGFRA extracellular domain mutants | 0.01 | -0.16 | -0.71 | 0.74 | 0.00 | 0.00 |
| R-HSA-9674555 | Signaling by CSF3 (G-CSF) | -0.11 | -0.22 | -0.45 | 0.73 | 0.00 | 0.00 |
| R-HSA-9682385 | FLT3 signaling in disease | -0.59 | -0.05 | -0.34 | 0.13 | 0.00 | 0.01 |
| R-HSA-9703465 | Signaling by FLT3 fusion proteins | -0.53 | -0.22 | -0.55 | 0.31 | 0.00 | 0.00 |
| R-HSA-9703648 | Signaling by FLT3 ITD and TKD mutants | -0.59 | -0.66 | -0.36 | 0.00 | 0.00 | 0.00 |
| R-HSA-196757 | Metabolism of folate and pterines | 0.18 | -0.56 | -0.51 | 0.05 | 0.00 | 0.00 |
| R-HSA-9018676 | Biosynthesis of D-series resolvins | 0.38 | -0.14 | 0.00 | 0.50 | 0.27 | 0.40 |
| R-HSA-9018677 | Biosynthesis of DHA-derived SPMs | 0.21 | -0.08 | -0.19 | 0.72 | 0.22 | 0.30 |
| R-HSA-9018678 | Biosynthesis of specialized proresolving mediators (SPMs) | 0.19 | -0.04 | -0.19 | 0.73 | 0.23 | 0.30 |
| R-HSA-9018679 | Biosynthesis of EPA-derived SPMs | 0.38 | -0.14 | -0.18 | 0.50 | 0.09 | 0.30 |
| R-HSA-9018896 | Biosynthesis of E-series 18(S)-resolvins | 0.38 | -0.14 | -0.14 | 0.50 | 0.16 | 0.36 |
| R-HSA-9020265 | Biosynthesis of aspirin-triggered D-series resolvins | 0.38 | -0.14 | -0.09 | 0.50 | 0.23 | 0.39 |
| R-HSA-9023661 | Biosynthesis of E-series 18(R)-resolvins | 0.38 | -0.14 | -0.24 | 0.50 | 0.03 | 0.18 |
| R-HSA-156588 | Glucuronidation | 0.16 | -0.10 | 0.01 | 0.73 | 0.38 | 0.40 |
| R-HSA-173599 | Formation of the active cofactor, UDP-glucuronate | 0.21 | 0.16 | -0.73 | 0.73 | 0.00 | 0.00 |
| R-HSA-975576 | N-glycan antennae elongation in the medial/trans-Golgi | 0.37 | 0.78 | -0.20 | 0.00 | 0.09 | 0.00 |
| R-HSA-975578 | Reactions specific to the complex N-glycan synthesis pathway | 0.37 | 0.78 | -0.33 | 0.00 | 0.00 | 0.00 |
| R-HSA-3214847 | HATs acetylate histones | -0.13 | -0.09 | -0.22 | 0.74 | 0.21 | 0.24 |
| R-HSA-110328 | Recognition and association of DNA glycosylase with site containing an affected pyrimidine | -0.63 | -0.24 | 0.02 | 0.07 | 0.02 | 0.38 |
| R-HSA-110329 | Cleavage of the damaged pyrimidine | -0.63 | -0.24 | 0.02 | 0.07 | 0.02 | 0.38 |
| R-HSA-110330 | Recognition and association of DNA glycosylase with site containing an affected purine | -0.63 | -0.24 | 0.08 | 0.07 | 0.01 | 0.36 |
| R-HSA-110331 | Cleavage of the damaged purine | -0.63 | -0.24 | 0.08 | 0.07 | 0.01 | 0.36 |
| R-HSA-1221632 | Meiotic synapsis | -0.63 | 0.18 | -0.11 | 0.00 | 0.01 | 0.36 |
| R-HSA-1474165 | Reproduction | -0.48 | 0.17 | -0.04 | 0.21 | 0.16 | 0.39 |
| R-HSA-1500620 | Meiosis | -0.63 | 0.01 | -0.05 | 0.04 | 0.02 | 0.40 |
| R-HSA-157579 | Telomere Maintenance | -0.47 | -0.28 | -0.13 | 0.44 | 0.10 | 0.32 |
| R-HSA-171306 | Packaging Of Telomere Ends | -0.63 | -0.24 | 0.11 | 0.07 | 0.01 | 0.33 |
| R-HSA-1912408 | Pre-NOTCH Transcription and Translation | -0.63 | -0.39 | -0.02 | 0.03 | 0.02 | 0.29 |
| R-HSA-201722 | Formation of the beta-catenin:TCF transactivating complex | -0.64 | -0.47 | 0.05 | 0.01 | 0.01 | 0.16 |
| R-HSA-212300 | PRC2 methylates histones and DNA | -0.30 | -0.39 | 0.13 | 0.57 | 0.26 | 0.16 |
| R-HSA-2299718 | Condensation of Prophase Chromosomes | -0.31 | -0.52 | 0.16 | 0.30 | 0.22 | 0.02 |
| R-HSA-2559582 | Senescence-Associated Secretory Phenotype (SASP) | -0.17 | -0.20 | 0.05 | 0.73 | 0.38 | 0.39 |
| R-HSA-427359 | SIRT1 negatively regulates rRNA expression | -0.63 | -0.39 | 0.25 | 0.03 | 0.00 | 0.01 |
| R-HSA-427413 | NoRC negatively regulates rRNA expression | -0.63 | -0.39 | 0.03 | 0.03 | 0.02 | 0.27 |
| R-HSA-5250924 | B-WICH complex positively regulates rRNA expression | -0.37 | -0.27 | 0.09 | 0.61 | 0.25 | 0.34 |
| R-HSA-5250941 | Negative epigenetic regulation of rRNA expression | -0.63 | -0.39 | 0.02 | 0.03 | 0.02 | 0.28 |
| R-HSA-5334118 | DNA methylation | -0.63 | -0.39 | 0.27 | 0.03 | 0.00 | 0.01 |
| R-HSA-5617472 | Activation of anterior HOX genes in hindbrain development during early embryogenesis | -0.30 | -0.39 | -0.07 | 0.57 | 0.32 | 0.28 |
| R-HSA-5619507 | Activation of HOX genes during differentiation | -0.30 | -0.39 | -0.07 | 0.57 | 0.32 | 0.28 |
| R-HSA-5625886 | Activated PKN1 stimulates transcription of AR (androgen receptor) regulated genes KLK2 and KLK3 | -0.63 | -0.39 | 0.31 | 0.03 | 0.00 | 0.00 |
| R-HSA-606279 | Deposition of new CENPA-containing nucleosomes at the centromere | -0.25 | -0.37 | 0.05 | 0.62 | 0.36 | 0.28 |
| R-HSA-73728 | RNA Polymerase I Promoter Opening | -0.43 | -0.55 | 0.33 | 0.10 | 0.00 | 0.00 |
| R-HSA-73886 | Chromosome Maintenance | -0.28 | -0.29 | -0.14 | 0.67 | 0.28 | 0.30 |
| R-HSA-73927 | Depurination | -0.63 | -0.24 | 0.08 | 0.07 | 0.01 | 0.36 |
| R-HSA-73928 | Depyrimidination | -0.63 | -0.24 | 0.02 | 0.07 | 0.02 | 0.38 |
| R-HSA-73929 | Base-Excision Repair, AP Site Formation | -0.63 | -0.24 | 0.01 | 0.07 | 0.02 | 0.38 |
| R-HSA-774815 | Nucleosome assembly | -0.25 | -0.37 | 0.05 | 0.62 | 0.36 | 0.28 |
| R-HSA-8866654 | E3 ubiquitin ligases ubiquitinate target proteins | -0.41 | -0.12 | -0.27 | 0.61 | 0.01 | 0.10 |
| R-HSA-8936459 | RUNX1 regulates genes involved in megakaryocyte differentiation and platelet function | -0.38 | 0.07 | 0.00 | 0.59 | 0.28 | 0.40 |
| R-HSA-912446 | Meiotic recombination | -0.63 | -0.39 | 0.15 | 0.03 | 0.00 | 0.12 |
| R-HSA-9616222 | Transcriptional regulation of granulopoiesis | -0.45 | -0.04 | 0.06 | 0.51 | 0.17 | 0.40 |
| R-HSA-9670095 | Inhibition of DNA recombination at telomere | -0.63 | -0.39 | 0.00 | 0.03 | 0.02 | 0.29 |
| R-HSA-9710421 | Defective pyroptosis | -0.21 | -0.38 | 0.12 | 0.63 | 0.33 | 0.19 |
| R-HSA-174403 | Glutathione synthesis and recycling | 0.40 | 0.30 | -0.26 | 0.55 | 0.01 | 0.03 |
| R-HSA-5578998 | Defective OPLAH causes OPLAHD | 0.69 | -0.48 | -0.37 | 0.00 | 0.00 | 0.00 |
| R-HSA-2022377 | Metabolism of Angiotensinogen to Angiotensins | 0.15 | -0.04 | 0.17 | 0.74 | 0.30 | 0.33 |
| R-HSA-166663 | Initial triggering of complement | 0.29 | 0.37 | -0.21 | 0.60 | 0.13 | 0.05 |
| R-HSA-166786 | Creation of C4 and C2 activators | 0.20 | 0.08 | -0.20 | 0.73 | 0.20 | 0.26 |
| R-HSA-163680 | AMPK inhibits chREBP transcriptional activation activity | 0.12 | 0.05 | -0.45 | 0.74 | 0.00 | 0.00 |
| R-HSA-2453864 | Retinoid cycle disease events | 0.01 | -0.20 | -0.04 | 0.73 | 0.38 | 0.39 |
| R-HSA-2453902 | The canonical retinoid cycle in rods (twilight vision) | 0.01 | -0.20 | 0.03 | 0.73 | 0.38 | 0.39 |
| R-HSA-2474795 | Diseases associated with visual transduction | 0.01 | -0.20 | -0.04 | 0.73 | 0.38 | 0.39 |
| R-HSA-9675143 | Diseases of the neuronal system | 0.01 | -0.20 | -0.04 | 0.73 | 0.38 | 0.39 |
| R-HSA-196783 | Coenzyme A biosynthesis | -0.70 | 0.91 | -0.58 | 0.00 | 0.00 | 0.00 |
| R-HSA-373753 | Nephrin family interactions | -0.48 | -0.22 | -0.51 | 0.46 | 0.00 | 0.00 |
| R-HSA-3214841 | PKMTs methylate histone lysines | 0.07 | -0.24 | -0.21 | 0.72 | 0.24 | 0.21 |
| R-HSA-3214858 | RMTs methylate histone arginines | -0.02 | -0.61 | 0.03 | 0.10 | 0.38 | 0.03 |
| R-HSA-6804758 | Regulation of TP53 Activity through Acetylation | 0.56 | 0.26 | -0.45 | 0.21 | 0.00 | 0.00 |
| R-HSA-73762 | RNA Polymerase I Transcription Initiation | 0.68 | -0.37 | -0.26 | 0.00 | 0.00 | 0.04 |
| R-HSA-425381 | Bicarbonate transporters | 0.03 | -0.21 | -0.29 | 0.73 | 0.06 | 0.05 |
| R-HSA-425393 | Transport of inorganic cations/anions and amino acids/oligopeptides | 0.00 | 0.06 | -0.08 | 0.74 | 0.38 | 0.39 |
| R-HSA-6790901 | rRNA modification in the nucleus and cytosol | -0.05 | 0.65 | -0.15 | 0.02 | 0.34 | 0.00 |
| R-HSA-9725371 | Nuclear events stimulated by ALK signaling in cancer | 0.03 | -0.16 | -0.28 | 0.74 | 0.09 | 0.09 |
| R-HSA-1834949 | Cytosolic sensors of pathogen-associated DNA | -0.81 | -0.17 | -0.44 | 0.00 | 0.00 | 0.00 |
| R-HSA-73780 | RNA Polymerase III Chain Elongation | -0.80 | -0.29 | -0.23 | 0.00 | 0.00 | 0.13 |
| R-HSA-73980 | RNA Polymerase III Transcription Termination | -0.42 | -0.33 | -0.31 | 0.50 | 0.00 | 0.01 |
| R-HSA-74158 | RNA Polymerase III Transcription | -0.42 | -0.33 | -0.34 | 0.50 | 0.00 | 0.00 |
| R-HSA-749476 | RNA Polymerase III Abortive And Retractive Initiation | -0.42 | -0.33 | -0.34 | 0.50 | 0.00 | 0.00 |
| R-HSA-76046 | RNA Polymerase III Transcription Initiation | -0.80 | -0.29 | -0.30 | 0.00 | 0.00 | 0.02 |
| R-HSA-76061 | RNA Polymerase III Transcription Initiation From Type 1 Promoter | -0.80 | -0.29 | -0.29 | 0.00 | 0.00 | 0.03 |
| R-HSA-76066 | RNA Polymerase III Transcription Initiation From Type 2 Promoter | -0.80 | -0.29 | -0.32 | 0.00 | 0.00 | 0.01 |
| R-HSA-76071 | RNA Polymerase III Transcription Initiation From Type 3 Promoter | -0.80 | -0.29 | -0.26 | 0.00 | 0.00 | 0.07 |
| R-HSA-164378 | PKA activation in glucagon signalling | 0.26 | -0.18 | -0.50 | 0.66 | 0.00 | 0.00 |
| R-HSA-1638091 | Heparan sulfate/heparin (HS-GAG) metabolism | 0.03 | 0.17 | -0.30 | 0.74 | 0.05 | 0.02 |
| R-HSA-1793185 | Chondroitin sulfate/dermatan sulfate metabolism | 0.12 | 0.27 | -0.31 | 0.72 | 0.02 | 0.00 |
| R-HSA-1971475 | A tetrasaccharide linker sequence is required for GAG synthesis | 0.12 | 0.27 | -0.32 | 0.72 | 0.02 | 0.00 |
| R-HSA-2022870 | Chondroitin sulfate biosynthesis | 0.17 | 0.28 | -0.22 | 0.71 | 0.19 | 0.10 |
| R-HSA-2022923 | Dermatan sulfate biosynthesis | 0.17 | 0.28 | -0.45 | 0.71 | 0.00 | 0.00 |
| R-HSA-2024101 | CS/DS degradation | 0.17 | 0.28 | -0.39 | 0.71 | 0.00 | 0.00 |
| R-HSA-3560783 | Defective B4GALT7 causes EDS, progeroid type | 0.12 | 0.27 | -0.39 | 0.72 | 0.00 | 0.00 |
| R-HSA-3560801 | Defective B3GAT3 causes JDSSDHD | 0.12 | 0.27 | -0.38 | 0.72 | 0.00 | 0.00 |
| R-HSA-3595172 | Defective CHST3 causes SEDCJD | 0.17 | 0.28 | -0.36 | 0.71 | 0.00 | 0.00 |
| R-HSA-3595174 | Defective CHST14 causes EDS, musculocontractural type | 0.17 | 0.28 | -0.40 | 0.71 | 0.00 | 0.00 |
| R-HSA-3595177 | Defective CHSY1 causes TPBS | 0.17 | 0.28 | -0.38 | 0.71 | 0.00 | 0.00 |
| R-HSA-4420332 | Defective B3GALT6 causes EDSP2 and SEMDJL1 | 0.12 | 0.27 | -0.39 | 0.72 | 0.00 | 0.00 |
| R-HSA-193368 | Synthesis of bile acids and bile salts via 7alpha-hydroxycholesterol | -0.31 | 0.11 | -0.29 | 0.65 | 0.02 | 0.04 |
| R-HSA-389887 | Beta-oxidation of pristanoyl-CoA | 0.89 | 0.80 | -0.55 | 0.00 | 0.00 | 0.00 |
| R-HSA-390918 | Peroxisomal lipid metabolism | 0.18 | 0.40 | -0.42 | 0.62 | 0.00 | 0.00 |
| R-HSA-9033500 | TYSND1 cleaves peroxisomal proteins | -0.14 | 0.13 | -0.53 | 0.73 | 0.00 | 0.00 |
| R-HSA-3769402 | Deactivation of the beta-catenin transactivating complex | -0.71 | -0.27 | -0.31 | 0.01 | 0.00 | 0.02 |
| R-HSA-392517 | Rap1 signalling | -0.02 | -0.30 | -0.46 | 0.71 | 0.00 | 0.00 |
| R-HSA-450604 | KSRP (KHSRP) binds and destabilizes mRNA | -0.19 | -0.41 | -0.27 | 0.60 | 0.07 | 0.02 |
| R-HSA-9013700 | NOTCH4 Activation and Transmission of Signal to the Nucleus | -0.26 | -0.77 | -0.46 | 0.00 | 0.00 | 0.00 |
| R-HSA-879415 | Advanced glycosylation endproduct receptor signaling | 0.38 | 0.04 | -0.32 | 0.64 | 0.00 | 0.02 |
| R-HSA-168268 | Virus Assembly and Release | -0.03 | -0.31 | -0.32 | 0.70 | 0.03 | 0.01 |
| R-HSA-168316 | Assembly of Viral Components at the Budding Site | -0.03 | -0.31 | -0.32 | 0.70 | 0.03 | 0.01 |
| R-HSA-1237112 | Methionine salvage pathway | 0.16 | 0.16 | -0.14 | 0.74 | 0.33 | 0.33 |
| R-HSA-8963693 | Aspartate and asparagine metabolism | -0.30 | 0.15 | -0.44 | 0.64 | 0.00 | 0.00 |
| R-HSA-209563 | Axonal growth stimulation | 0.34 | -0.23 | -0.23 | 0.46 | 0.05 | 0.17 |
| R-HSA-8964208 | Phenylalanine metabolism | 0.63 | -0.01 | -0.17 | 0.04 | 0.00 | 0.34 |
| R-HSA-73621 | Pyrimidine catabolism | 0.04 | 0.23 | -0.12 | 0.73 | 0.37 | 0.33 |
| R-HSA-5357905 | Regulation of TNFR1 signaling | 0.04 | 0.29 | -0.45 | 0.71 | 0.00 | 0.00 |
| R-HSA-5357956 | TNFR1-induced NFkappaB signaling pathway | 0.04 | 0.29 | -0.47 | 0.71 | 0.00 | 0.00 |
| R-HSA-5626978 | TNFR1-mediated ceramide production | 0.55 | 0.68 | -0.22 | 0.00 | 0.00 | 0.00 |
| R-HSA-75893 | TNF signaling | 0.04 | 0.29 | -0.46 | 0.71 | 0.00 | 0.00 |
| R-HSA-380994 | ATF4 activates genes in response to endoplasmic reticulum stress | -0.57 | 0.57 | -0.16 | 0.00 | 0.01 | 0.01 |
| R-HSA-2173788 | Downregulation of TGF-beta receptor signaling | 0.14 | 0.19 | -0.38 | 0.73 | 0.00 | 0.00 |
| R-HSA-2173789 | TGF-beta receptor signaling activates SMADs | -0.06 | 0.11 | -0.37 | 0.74 | 0.00 | 0.00 |
| R-HSA-450520 | HuR (ELAVL1) binds and stabilizes mRNA | -0.12 | -0.56 | -0.45 | 0.26 | 0.00 | 0.00 |
| R-HSA-9634638 | Estrogen-dependent nuclear events downstream of ESR-membrane signaling | 0.06 | -0.36 | -0.31 | 0.63 | 0.03 | 0.01 |
| R-HSA-9707616 | Heme signaling | -0.28 | 0.03 | -0.53 | 0.70 | 0.00 | 0.00 |
| R-HSA-351906 | Apoptotic cleavage of cell adhesion proteins | 0.06 | 0.03 | -0.42 | 0.74 | 0.00 | 0.00 |
| R-HSA-74182 | Ketone body metabolism | 0.02 | -0.03 | -0.45 | 0.74 | 0.00 | 0.00 |
| R-HSA-77108 | Utilization of Ketone Bodies | -0.10 | -0.29 | -0.42 | 0.71 | 0.00 | 0.00 |
| R-HSA-77111 | Synthesis of Ketone Bodies | -0.10 | 0.07 | -0.53 | 0.74 | 0.00 | 0.00 |
| R-HSA-168643 | Nucleotide-binding domain, leucine rich repeat containing receptor (NLR) signaling pathways | -0.18 | -0.37 | -0.41 | 0.65 | 0.00 | 0.00 |
| R-HSA-5676934 | Protein repair | 0.03 | -0.48 | -0.46 | 0.41 | 0.00 | 0.00 |
| R-HSA-622312 | Inflammasomes | -0.03 | -0.24 | -0.29 | 0.73 | 0.06 | 0.05 |
| R-HSA-844456 | The NLRP3 inflammasome | -0.03 | -0.24 | -0.39 | 0.73 | 0.00 | 0.00 |
| R-HSA-9660826 | Purinergic signaling in leishmaniasis infection | -0.08 | 0.03 | -0.30 | 0.74 | 0.04 | 0.04 |
| R-HSA-9664424 | Cell recruitment (pro-inflammatory response) | -0.08 | 0.03 | -0.30 | 0.74 | 0.04 | 0.04 |
| R-HSA-936440 | Negative regulators of DDX58/IFIH1 signaling | -0.17 | -0.01 | -0.50 | 0.74 | 0.00 | 0.00 |
| R-HSA-1362409 | Mitochondrial iron-sulfur cluster biogenesis | -0.52 | 0.13 | -0.29 | 0.14 | 0.00 | 0.03 |
| R-HSA-110056 | MAPK3 (ERK1) activation | 0.23 | -0.60 | -0.55 | 0.01 | 0.00 | 0.00 |
| R-HSA-112409 | RAF-independent MAPK1/3 activation | 0.18 | -0.44 | -0.41 | 0.32 | 0.00 | 0.00 |
| R-HSA-445144 | Signal transduction by L1 | 0.51 | -0.26 | -0.40 | 0.04 | 0.00 | 0.00 |
| R-HSA-5210891 | Uptake and function of anthrax toxins | 0.31 | -0.17 | -0.37 | 0.61 | 0.00 | 0.00 |
| R-HSA-5674499 | Negative feedback regulation of MAPK pathway | 0.48 | -0.34 | -0.39 | 0.02 | 0.00 | 0.00 |
| R-HSA-5684264 | MAP3K8 (TPL2)-dependent MAPK1/3 activation | -0.22 | 0.15 | -0.30 | 0.70 | 0.03 | 0.03 |
| R-HSA-9652169 | Signaling by MAP2K mutants | 0.48 | -0.34 | -0.27 | 0.02 | 0.00 | 0.05 |
| R-HSA-6782315 | tRNA modification in the nucleus and cytosol | 0.62 | -0.04 | -0.32 | 0.04 | 0.00 | 0.02 |
| R-HSA-6804756 | Regulation of TP53 Activity through Phosphorylation | -0.06 | -0.16 | -0.35 | 0.74 | 0.01 | 0.01 |
| R-HSA-164938 | Nef-mediates down modulation of cell surface receptors by recruiting them to clathrin adapters | 0.20 | -0.04 | -0.31 | 0.73 | 0.02 | 0.04 |
| R-HSA-164952 | The role of Nef in HIV-1 replication and disease pathogenesis | 0.15 | -0.02 | -0.30 | 0.74 | 0.04 | 0.05 |
| R-HSA-167590 | Nef Mediated CD4 Down-regulation | 0.35 | -0.02 | -0.28 | 0.65 | 0.01 | 0.09 |
| R-HSA-182218 | Nef Mediated CD8 Down-regulation | 0.66 | 0.30 | -0.08 | 0.03 | 0.01 | 0.32 |
| R-HSA-399719 | Trafficking of AMPA receptors | 0.38 | -0.15 | -0.19 | 0.51 | 0.10 | 0.30 |
| R-HSA-399721 | Glutamate binding, activation of AMPA receptors and synaptic plasticity | 0.38 | -0.15 | -0.19 | 0.51 | 0.10 | 0.30 |
| R-HSA-416993 | Trafficking of GluR2-containing AMPA receptors | 0.44 | -0.02 | -0.15 | 0.51 | 0.09 | 0.36 |
| R-HSA-71262 | Carnitine synthesis | 0.63 | 0.82 | -0.35 | 0.00 | 0.00 | 0.00 |
| R-HSA-388841 | Costimulation by the CD28 family | 0.03 | -0.09 | -0.38 | 0.74 | 0.00 | 0.00 |
| R-HSA-389356 | CD28 co-stimulation | -0.08 | -0.07 | -0.34 | 0.74 | 0.01 | 0.01 |
| R-HSA-389359 | CD28 dependent Vav1 pathway | -0.08 | -0.07 | -0.25 | 0.74 | 0.15 | 0.17 |
| R-HSA-418885 | DCC mediated attractive signaling | -0.06 | 0.22 | -0.42 | 0.73 | 0.00 | 0.00 |
| R-HSA-428543 | Inactivation of CDC42 and RAC1 | 0.21 | 0.47 | -0.34 | 0.49 | 0.00 | 0.00 |
| R-HSA-5625970 | RHO GTPases activate KTN1 | -0.15 | -0.01 | -0.24 | 0.74 | 0.16 | 0.20 |
| R-HSA-5689877 | Josephin domain DUBs | -0.18 | -0.35 | -0.20 | 0.67 | 0.24 | 0.17 |
| R-HSA-9609523 | Insertion of tail-anchored proteins into the endoplasmic reticulum membrane | -0.17 | 0.21 | -0.06 | 0.70 | 0.37 | 0.38 |
| R-HSA-168638 | NOD1/2 Signaling Pathway | -0.32 | -0.51 | -0.49 | 0.33 | 0.00 | 0.00 |
| R-HSA-2151209 | Activation of PPARGC1A (PGC-1alpha) by phosphorylation | -0.10 | -0.52 | -0.75 | 0.37 | 0.00 | 0.00 |
| R-HSA-376172 | DSCAM interactions | 0.03 | -0.47 | -0.11 | 0.42 | 0.37 | 0.15 |
| R-HSA-964975 | Vitamins B6 activation to pyridoxal phosphate | 0.16 | -0.63 | -0.49 | 0.01 | 0.00 | 0.00 |
| R-HSA-211897 | Cytochrome P450 - arranged by substrate type | -0.39 | -0.07 | -0.13 | 0.63 | 0.19 | 0.38 |
| R-HSA-211935 | Fatty acids | -0.41 | -0.45 | -0.10 | 0.33 | 0.20 | 0.19 |
| R-HSA-211958 | Miscellaneous substrates | -0.41 | -0.45 | -0.14 | 0.33 | 0.15 | 0.15 |
| R-HSA-211979 | Eicosanoids | -0.56 | -0.32 | 0.05 | 0.18 | 0.06 | 0.33 |
| R-HSA-2142691 | Synthesis of Leukotrienes (LT) and Eoxins (EX) | -0.16 | 0.17 | -0.09 | 0.72 | 0.37 | 0.38 |
| R-HSA-2122948 | Activated NOTCH1 Transmits Signal to the Nucleus | -0.02 | 0.66 | -0.33 | 0.02 | 0.02 | 0.00 |
| R-HSA-5635838 | Activation of SMO | 0.71 | 0.86 | -0.20 | 0.00 | 0.00 | 0.00 |
| R-HSA-140875 | Common Pathway of Fibrin Clot Formation | 0.23 | 0.13 | -0.24 | 0.73 | 0.10 | 0.14 |
| R-HSA-354194 | GRB2:SOS provides linkage to MAPK signaling for Integrins | -0.29 | -0.07 | -0.34 | 0.71 | 0.00 | 0.01 |
| R-HSA-5260271 | Diseases of Immune System | -0.10 | -0.12 | -0.27 | 0.74 | 0.09 | 0.10 |
| R-HSA-5602358 | Diseases associated with the TLR signaling cascade | -0.10 | -0.12 | -0.27 | 0.74 | 0.09 | 0.10 |
| R-HSA-5602498 | MyD88 deficiency (TLR2/4) | -0.10 | -0.12 | -0.28 | 0.74 | 0.08 | 0.09 |
| R-HSA-5603041 | IRAK4 deficiency (TLR2/4) | -0.10 | -0.12 | -0.27 | 0.74 | 0.09 | 0.10 |
| R-HSA-5686938 | Regulation of TLR by endogenous ligand | -0.10 | -0.16 | -0.10 | 0.74 | 0.37 | 0.38 |
| R-HSA-8877330 | RUNX1 and FOXP3 control the development of regulatory T lymphocytes (Tregs) | -0.69 | 0.86 | -0.25 | 0.00 | 0.00 | 0.00 |
| R-HSA-8931987 | RUNX1 regulates estrogen receptor mediated transcription | -0.69 | 0.86 | -0.53 | 0.00 | 0.00 | 0.00 |
| R-HSA-8934593 | Regulation of RUNX1 Expression and Activity | -0.69 | 0.86 | -0.64 | 0.00 | 0.00 | 0.00 |
| R-HSA-8935964 | RUNX1 regulates expression of components of tight junctions | -0.43 | 0.80 | -0.49 | 0.00 | 0.00 | 0.00 |
| R-HSA-8939242 | RUNX1 regulates transcription of genes involved in differentiation of keratinocytes | -0.69 | 0.86 | -0.27 | 0.00 | 0.00 | 0.00 |
| R-HSA-8939245 | RUNX1 regulates transcription of genes involved in BCR signaling | -0.69 | 0.86 | -0.41 | 0.00 | 0.00 | 0.00 |
| R-HSA-8939246 | RUNX1 regulates transcription of genes involved in differentiation of myeloid cells | -0.25 | 0.80 | -0.42 | 0.00 | 0.00 | 0.00 |
| R-HSA-8939247 | RUNX1 regulates transcription of genes involved in interleukin signaling | -0.69 | 0.86 | -0.45 | 0.00 | 0.00 | 0.00 |
| R-HSA-8939256 | RUNX1 regulates transcription of genes involved in WNT signaling | -0.69 | 0.86 | -0.33 | 0.00 | 0.00 | 0.00 |
| R-HSA-8940973 | RUNX2 regulates osteoblast differentiation | -0.03 | 0.23 | -0.35 | 0.73 | 0.01 | 0.00 |
| R-HSA-8941284 | RUNX2 regulates chondrocyte maturation | -0.69 | 0.86 | -0.33 | 0.00 | 0.00 | 0.00 |
| R-HSA-8941326 | RUNX2 regulates bone development | -0.03 | 0.23 | -0.34 | 0.73 | 0.01 | 0.00 |
| R-HSA-8941332 | RUNX2 regulates genes involved in cell migration | -0.69 | 0.86 | -0.38 | 0.00 | 0.00 | 0.00 |
| R-HSA-8941333 | RUNX2 regulates genes involved in differentiation of myeloid cells | -0.25 | 0.80 | -0.34 | 0.00 | 0.00 | 0.00 |
| R-HSA-8949275 | RUNX3 Regulates Immune Response and Cell Migration | -0.69 | 0.86 | -0.50 | 0.00 | 0.00 | 0.00 |
| R-HSA-8951911 | RUNX3 regulates RUNX1-mediated transcription | -0.69 | 0.86 | -0.49 | 0.00 | 0.00 | 0.00 |
| R-HSA-379726 | Mitochondrial tRNA aminoacylation | 0.22 | -0.44 | -0.50 | 0.24 | 0.00 | 0.00 |
| R-HSA-438066 | Unblocking of NMDA receptors, glutamate binding and activation | -0.35 | -0.59 | -0.01 | 0.11 | 0.31 | 0.05 |
| R-HSA-9617324 | Negative regulation of NMDA receptor-mediated neuronal transmission | -0.35 | -0.59 | -0.17 | 0.11 | 0.17 | 0.02 |
| R-HSA-9620244 | Long-term potentiation | -0.35 | -0.59 | -0.07 | 0.11 | 0.29 | 0.05 |
| R-HSA-5357609 | Glycogen storage disease type II (GAA) | -0.83 | -0.59 | -0.79 | 0.00 | 0.00 | 0.00 |
| R-HSA-111367 | SLBP independent Processing of Histone Pre-mRNAs | 0.67 | 0.78 | 0.02 | 0.00 | 0.01 | 0.00 |
| R-HSA-75067 | Processing of Capped Intronless Pre-mRNA | 0.67 | 0.78 | -0.22 | 0.00 | 0.00 | 0.00 |
| R-HSA-77588 | SLBP Dependent Processing of Replication-Dependent Histone Pre-mRNAs | 0.67 | 0.78 | -0.01 | 0.00 | 0.01 | 0.00 |
| R-HSA-1482801 | Acyl chain remodelling of PS | -0.50 | -0.09 | -0.07 | 0.41 | 0.11 | 0.40 |
| R-HSA-1566977 | Fibronectin matrix formation | -0.98 | 0.30 | -0.52 | 0.00 | 0.00 | 0.00 |
| R-HSA-9700645 | ALK mutants bind TKIs | -0.24 | 0.01 | -0.40 | 0.72 | 0.00 | 0.00 |
| R-HSA-9637628 | Modulation by Mtb of host immune system | -0.63 | 0.41 | -0.45 | 0.00 | 0.00 | 0.00 |
| R-HSA-5083635 | Defective B3GALTL causes Peters-plus syndrome (PpS) | 0.21 | 0.67 | -0.39 | 0.04 | 0.00 | 0.00 |
| R-HSA-5173214 | O-glycosylation of TSR domain-containing proteins | 0.21 | 0.67 | -0.38 | 0.04 | 0.00 | 0.00 |
| R-HSA-6804760 | Regulation of TP53 Activity through Methylation | -0.88 | 0.39 | -0.31 | 0.00 | 0.00 | 0.00 |
| R-HSA-917977 | Transferrin endocytosis and recycling | -0.16 | -0.14 | -0.25 | 0.74 | 0.13 | 0.16 |
| R-HSA-5609974 | Defective PGM1 causes PGM1-CDG (CDG1t) | -0.68 | -0.88 | -0.97 | 0.00 | 0.00 | 0.00 |
| R-HSA-110312 | Translesion synthesis by REV1 | -0.76 | 0.47 | -0.35 | 0.00 | 0.00 | 0.00 |
| R-HSA-110313 | Translesion synthesis by Y family DNA polymerases bypasses lesions on DNA template | -0.03 | 0.06 | -0.37 | 0.74 | 0.00 | 0.00 |
| R-HSA-110320 | Translesion Synthesis by POLH | -0.03 | 0.06 | -0.32 | 0.74 | 0.03 | 0.02 |
| R-HSA-1253288 | Downregulation of ERBB4 signaling | -0.76 | 0.47 | -0.50 | 0.00 | 0.00 | 0.00 |
| R-HSA-1295596 | Spry regulation of FGF signaling | 0.06 | -0.19 | -0.37 | 0.73 | 0.00 | 0.00 |
| R-HSA-1358803 | Downregulation of ERBB2:ERBB3 signaling | -0.76 | 0.47 | -0.29 | 0.00 | 0.00 | 0.00 |
| R-HSA-162588 | Budding and maturation of HIV virion | -0.12 | 0.15 | -0.22 | 0.73 | 0.20 | 0.17 |
| R-HSA-168927 | TICAM1, RIP1-mediated IKK complex recruitment | -0.47 | 0.02 | -0.35 | 0.42 | 0.00 | 0.01 |
| R-HSA-174048 | APC/C:Cdc20 mediated degradation of Cyclin B | -0.76 | 0.47 | -0.05 | 0.00 | 0.00 | 0.16 |
| R-HSA-174490 | Membrane binding and targetting of GAG proteins | -0.76 | 0.47 | -0.27 | 0.00 | 0.00 | 0.00 |
| R-HSA-174495 | Synthesis And Processing Of GAG, GAGPOL Polyproteins | -0.76 | 0.47 | -0.27 | 0.00 | 0.00 | 0.00 |
| R-HSA-175474 | Assembly Of The HIV Virion | -0.76 | 0.47 | -0.28 | 0.00 | 0.00 | 0.00 |
| R-HSA-179409 | APC-Cdc20 mediated degradation of Nek2A | -0.76 | 0.47 | -0.12 | 0.00 | 0.00 | 0.09 |
| R-HSA-1980145 | Signaling by NOTCH2 | -0.76 | 0.47 | -0.18 | 0.00 | 0.00 | 0.03 |
| R-HSA-2173791 | TGF-beta receptor signaling in EMT (epithelial to mesenchymal transition) | -0.47 | -0.01 | -0.24 | 0.46 | 0.02 | 0.19 |
| R-HSA-2559585 | Oncogene Induced Senescence | 0.12 | -0.13 | -0.25 | 0.73 | 0.12 | 0.15 |
| R-HSA-2691230 | Signaling by NOTCH1 HD Domain Mutants in Cancer | -0.76 | 0.47 | -0.29 | 0.00 | 0.00 | 0.00 |
| R-HSA-2691232 | Constitutive Signaling by NOTCH1 HD Domain Mutants | -0.76 | 0.47 | -0.29 | 0.00 | 0.00 | 0.00 |
| R-HSA-2979096 | NOTCH2 Activation and Transmission of Signal to the Nucleus | -0.76 | 0.47 | -0.34 | 0.00 | 0.00 | 0.00 |
| R-HSA-3134975 | Regulation of innate immune responses to cytosolic DNA | -0.76 | 0.47 | -0.50 | 0.00 | 0.00 | 0.00 |
| R-HSA-3785653 | Myoclonic epilepsy of Lafora | -0.48 | -0.28 | -0.41 | 0.41 | 0.00 | 0.00 |
| R-HSA-400253 | Circadian Clock | -0.13 | -0.05 | -0.47 | 0.74 | 0.00 | 0.00 |
| R-HSA-445989 | TAK1 activates NFkB by phosphorylation and activation of IKKs complex | -0.60 | -0.02 | -0.43 | 0.10 | 0.00 | 0.00 |
| R-HSA-450321 | JNK (c-Jun kinases) phosphorylation and activation mediated by activated human TAK1 | -0.76 | 0.47 | -0.48 | 0.00 | 0.00 | 0.00 |
| R-HSA-4641263 | Regulation of FZD by ubiquitination | -0.76 | 0.47 | -0.21 | 0.00 | 0.00 | 0.01 |
| R-HSA-5654726 | Negative regulation of FGFR1 signaling | 0.06 | -0.19 | -0.13 | 0.73 | 0.36 | 0.36 |
| R-HSA-5654727 | Negative regulation of FGFR2 signaling | 0.06 | -0.19 | -0.12 | 0.73 | 0.37 | 0.37 |
| R-HSA-5654732 | Negative regulation of FGFR3 signaling | 0.06 | -0.19 | -0.17 | 0.73 | 0.31 | 0.31 |
| R-HSA-5654733 | Negative regulation of FGFR4 signaling | 0.06 | -0.19 | -0.10 | 0.73 | 0.37 | 0.38 |
| R-HSA-5655862 | Translesion synthesis by POLK | -0.76 | 0.47 | -0.38 | 0.00 | 0.00 | 0.00 |
| R-HSA-5656121 | Translesion synthesis by POLI | -0.76 | 0.47 | -0.36 | 0.00 | 0.00 | 0.00 |
| R-HSA-5656169 | Termination of translesion DNA synthesis | -0.76 | 0.47 | -0.37 | 0.00 | 0.00 | 0.00 |
| R-HSA-5685942 | HDR through Homologous Recombination (HRR) | -0.76 | 0.47 | -0.34 | 0.00 | 0.00 | 0.00 |
| R-HSA-5689896 | Ovarian tumor domain proteases | -0.27 | -0.30 | -0.38 | 0.67 | 0.00 | 0.00 |
| R-HSA-5696397 | Gap-filling DNA repair synthesis and ligation in GG-NER | -0.76 | 0.47 | -0.27 | 0.00 | 0.00 | 0.00 |
| R-HSA-6783310 | Fanconi Anemia Pathway | -0.76 | 0.47 | -0.38 | 0.00 | 0.00 | 0.00 |
| R-HSA-6804757 | Regulation of TP53 Degradation | -0.23 | -0.12 | -0.32 | 0.73 | 0.01 | 0.02 |
| R-HSA-6806003 | Regulation of TP53 Expression and Degradation | -0.23 | -0.12 | -0.34 | 0.73 | 0.01 | 0.01 |
| R-HSA-69231 | Cyclin D associated events in G1 | -0.19 | -0.12 | -0.25 | 0.73 | 0.12 | 0.16 |
| R-HSA-69236 | G1 Phase | -0.19 | -0.12 | -0.25 | 0.73 | 0.12 | 0.16 |
| R-HSA-8849469 | PTK6 Regulates RTKs and Their Effectors AKT1 and DOK1 | -0.76 | 0.47 | -0.10 | 0.00 | 0.00 | 0.11 |
| R-HSA-8876493 | InlA-mediated entry of Listeria monocytogenes into host cells | -0.82 | -0.04 | -0.29 | 0.00 | 0.00 | 0.07 |
| R-HSA-8948747 | Regulation of PTEN localization | -0.76 | 0.47 | -0.42 | 0.00 | 0.00 | 0.00 |
| R-HSA-901032 | ER Quality Control Compartment (ERQC) | -0.38 | -0.19 | -0.41 | 0.64 | 0.00 | 0.00 |
| R-HSA-9013973 | TICAM1-dependent activation of IRF3/IRF7 | -0.76 | 0.47 | -0.37 | 0.00 | 0.00 | 0.00 |
| R-HSA-9014325 | TICAM1,TRAF6-dependent induction of TAK1 complex | -0.76 | 0.47 | -0.43 | 0.00 | 0.00 | 0.00 |
| R-HSA-936964 | Activation of IRF3/IRF7 mediated by TBK1/IKK epsilon | -0.76 | 0.47 | -0.33 | 0.00 | 0.00 | 0.00 |
| R-HSA-937039 | IRAK1 recruits IKK complex | -0.76 | 0.47 | -0.30 | 0.00 | 0.00 | 0.00 |
| R-HSA-937041 | IKK complex recruitment mediated by RIP1 | -0.47 | 0.02 | -0.29 | 0.42 | 0.00 | 0.05 |
| R-HSA-937042 | IRAK2 mediated activation of TAK1 complex | -0.76 | 0.47 | -0.42 | 0.00 | 0.00 | 0.00 |
| R-HSA-937072 | TRAF6-mediated induction of TAK1 complex within TLR4 complex | -0.76 | 0.47 | -0.32 | 0.00 | 0.00 | 0.00 |
| R-HSA-9645460 | Alpha-protein kinase 1 signaling pathway | -0.76 | 0.47 | -0.51 | 0.00 | 0.00 | 0.00 |
| R-HSA-9683683 | Maturation of protein E | -0.76 | 0.47 | -0.08 | 0.00 | 0.00 | 0.13 |
| R-HSA-9694493 | Maturation of protein E | -0.76 | 0.47 | -0.08 | 0.00 | 0.00 | 0.13 |
| R-HSA-9705462 | Inactivation of CSF3 (G-CSF) signaling | 0.02 | -0.10 | -0.40 | 0.74 | 0.00 | 0.00 |
| R-HSA-9706369 | Negative regulation of FLT3 | -0.05 | 0.22 | -0.15 | 0.72 | 0.34 | 0.29 |
| R-HSA-9706377 | FLT3 signaling by CBL mutants | -0.76 | 0.47 | -0.05 | 0.00 | 0.00 | 0.16 |
| R-HSA-9708530 | Regulation of BACH1 activity | -0.76 | 0.47 | -0.11 | 0.00 | 0.00 | 0.10 |
| R-HSA-975110 | TRAF6 mediated IRF7 activation in TLR7/8 or 9 signaling | -0.76 | 0.47 | -0.22 | 0.00 | 0.00 | 0.01 |
| R-HSA-975144 | IRAK1 recruits IKK complex upon TLR7/8 or 9 stimulation | -0.76 | 0.47 | -0.30 | 0.00 | 0.00 | 0.00 |
| R-HSA-975163 | IRAK2 mediated activation of TAK1 complex upon TLR7/8 or 9 stimulation | -0.76 | 0.47 | -0.39 | 0.00 | 0.00 | 0.00 |
| R-HSA-189451 | Heme biosynthesis | -0.28 | -0.19 | -0.18 | 0.71 | 0.20 | 0.29 |
| R-HSA-5579006 | Defective GSS causes GSS deficiency | 0.58 | 0.82 | -0.31 | 0.00 | 0.00 | 0.00 |
| R-HSA-5633008 | TP53 Regulates Transcription of Cell Death Genes | -0.07 | 0.09 | -0.16 | 0.74 | 0.33 | 0.34 |
| R-HSA-6803205 | TP53 regulates transcription of several additional cell death genes whose specific roles in p53-dependent apoptosis remain uncertain | -0.07 | 0.09 | -0.31 | 0.74 | 0.03 | 0.02 |
| R-HSA-2995383 | Initiation of Nuclear Envelope (NE) Reformation | -0.23 | -0.05 | -0.26 | 0.73 | 0.08 | 0.14 |
| R-HSA-1483191 | Synthesis of PC | 0.17 | 0.03 | -0.24 | 0.74 | 0.14 | 0.18 |
| R-HSA-390247 | Beta-oxidation of very long chain fatty acids | 0.16 | 0.43 | -0.45 | 0.58 | 0.00 | 0.00 |
| R-HSA-111931 | PKA-mediated phosphorylation of CREB | 0.16 | -0.39 | -0.46 | 0.47 | 0.00 | 0.00 |
| R-HSA-111933 | Calmodulin induced events | 0.16 | -0.44 | -0.47 | 0.35 | 0.00 | 0.00 |
| R-HSA-111996 | Ca-dependent events | 0.27 | -0.35 | -0.48 | 0.36 | 0.00 | 0.00 |
| R-HSA-111997 | CaM pathway | 0.16 | -0.44 | -0.47 | 0.35 | 0.00 | 0.00 |
| R-HSA-1489509 | DAG and IP3 signaling | 0.24 | -0.42 | -0.51 | 0.23 | 0.00 | 0.00 |
| R-HSA-163615 | PKA activation | 0.16 | -0.39 | -0.42 | 0.47 | 0.00 | 0.00 |
| R-HSA-442720 | CREB1 phosphorylation through the activation of Adenylate Cyclase | 0.16 | -0.39 | -0.34 | 0.47 | 0.00 | 0.00 |
| R-HSA-9010642 | ROBO receptors bind AKAP5 | 0.31 | -0.64 | -0.67 | 0.00 | 0.00 | 0.00 |
| R-HSA-9664323 | FCGR3A-mediated IL10 synthesis | 0.26 | -0.38 | -0.50 | 0.31 | 0.00 | 0.00 |
| R-HSA-2028269 | Signaling by Hippo | -0.09 | 0.17 | -0.54 | 0.73 | 0.00 | 0.00 |
| R-HSA-8984722 | Interleukin-35 Signalling | 0.14 | -0.11 | -0.52 | 0.73 | 0.00 | 0.00 |
| R-HSA-9020956 | Interleukin-27 signaling | 0.14 | -0.11 | -0.56 | 0.73 | 0.00 | 0.00 |
| R-HSA-352238 | Breakdown of the nuclear lamina | -0.99 | 0.72 | -0.02 | 0.00 | 0.00 | 0.00 |
| R-HSA-4419969 | Depolymerisation of the Nuclear Lamina | -0.99 | 0.72 | -0.22 | 0.00 | 0.00 | 0.00 |
| R-HSA-1236973 | Cross-presentation of particulate exogenous antigens (phagosomes) | 0.65 | 0.51 | -0.30 | 0.01 | 0.00 | 0.00 |
| R-HSA-3000471 | Scavenging by Class B Receptors | 0.34 | 0.18 | -0.01 | 0.67 | 0.31 | 0.39 |
| R-HSA-434313 | Intracellular metabolism of fatty acids regulates insulin secretion | 0.65 | 0.51 | -0.27 | 0.01 | 0.00 | 0.00 |
| R-HSA-173623 | Classical antibody-mediated complement activation | 0.20 | 0.08 | -0.12 | 0.73 | 0.34 | 0.38 |
| R-HSA-5423646 | Aflatoxin activation and detoxification | -0.54 | -0.13 | -0.13 | 0.30 | 0.05 | 0.37 |
| R-HSA-1855204 | Synthesis of IP3 and IP4 in the cytosol | 0.56 | 0.48 | -0.36 | 0.05 | 0.00 | 0.00 |
| R-HSA-166665 | Terminal pathway of complement | -0.06 | 0.06 | -0.03 | 0.74 | 0.38 | 0.40 |
| R-HSA-6809583 | Retinoid metabolism disease events | 0.48 | -0.05 | -0.13 | 0.37 | 0.08 | 0.38 |
| R-HSA-6799990 | Metal sequestration by antimicrobial proteins | -0.37 | 0.01 | -0.20 | 0.64 | 0.11 | 0.29 |
| R-HSA-446210 | Synthesis of UDP-N-acetyl-glucosamine | -0.16 | -0.06 | -0.36 | 0.74 | 0.00 | 0.00 |
| R-HSA-2393930 | Phosphate bond hydrolysis by NUDT proteins | 0.50 | -0.86 | -0.39 | 0.00 | 0.00 | 0.00 |
| R-HSA-9673768 | Signaling by membrane-tethered fusions of PDGFRA or PDGFRB | 0.66 | -0.49 | -0.46 | 0.00 | 0.00 | 0.00 |
| R-HSA-173736 | Alternative complement activation | 0.46 | 0.96 | -0.18 | 0.00 | 0.04 | 0.00 |
| R-HSA-170670 | Adenylate cyclase inhibitory pathway | 0.70 | 0.91 | -0.41 | 0.00 | 0.00 | 0.00 |
| R-HSA-1679131 | Trafficking and processing of endosomal TLR | 0.40 | -0.94 | -0.27 | 0.00 | 0.01 | 0.00 |
| R-HSA-1433559 | Regulation of KIT signaling | 0.10 | -0.39 | -0.48 | 0.55 | 0.00 | 0.00 |
| R-HSA-5668599 | RHO GTPases Activate NADPH Oxidases | 0.07 | -0.26 | -0.22 | 0.71 | 0.23 | 0.19 |
| R-HSA-8853659 | RET signaling | 0.05 | -0.39 | -0.40 | 0.58 | 0.00 | 0.00 |
| R-HSA-210990 | PECAM1 interactions | 0.42 | 0.20 | -0.56 | 0.57 | 0.00 | 0.00 |
| R-HSA-432142 | Platelet sensitization by LDL | 0.25 | -0.23 | -0.37 | 0.62 | 0.00 | 0.00 |
| R-HSA-156584 | Cytosolic sulfonation of small molecules | 0.17 | -0.34 | -0.05 | 0.56 | 0.37 | 0.33 |
| R-HSA-1369007 | Mitochondrial ABC transporters | -0.79 | 0.08 | -0.71 | 0.00 | 0.00 | 0.00 |
| R-HSA-2564830 | Cytosolic iron-sulfur cluster assembly | -0.23 | 0.06 | -0.25 | 0.72 | 0.09 | 0.14 |
| R-HSA-8983711 | OAS antiviral response | 0.04 | -0.90 | -0.44 | 0.00 | 0.00 | 0.00 |
| R-HSA-73817 | Purine ribonucleoside monophosphate biosynthesis | 0.24 | 0.27 | -0.28 | 0.70 | 0.04 | 0.02 |
| R-HSA-8956320 | Nucleobase biosynthesis | 0.24 | 0.27 | -0.25 | 0.70 | 0.08 | 0.05 |
| R-HSA-264876 | Insulin processing | -0.45 | -1.00 | -0.38 | 0.00 | 0.00 | 0.00 |
| R-HSA-3928664 | Ephrin signaling | -0.01 | 0.05 | -0.26 | 0.74 | 0.12 | 0.11 |
| R-HSA-447043 | Neurofascin interactions | -0.44 | 0.59 | -0.81 | 0.00 | 0.00 | 0.00 |
| R-HSA-164940 | Nef mediated downregulation of MHC class I complex cell surface expression | -0.11 | -0.08 | -0.43 | 0.74 | 0.00 | 0.00 |
| R-HSA-1222556 | ROS and RNS production in phagocytes | -0.39 | -0.18 | -0.21 | 0.63 | 0.07 | 0.25 |
| R-HSA-77387 | Insulin receptor recycling | -0.29 | -0.20 | -0.19 | 0.70 | 0.17 | 0.27 |
| R-HSA-2408550 | Metabolism of ingested H2SeO4 and H2SeO3 into H2Se | -0.14 | -0.50 | -0.46 | 0.43 | 0.00 | 0.00 |
| R-HSA-5620916 | VxPx cargo-targeting to cilium | -0.71 | -0.90 | -0.41 | 0.00 | 0.00 | 0.00 |
| R-HSA-193807 | Synthesis of bile acids and bile salts via 27-hydroxycholesterol | -0.31 | 0.11 | -0.32 | 0.65 | 0.00 | 0.01 |
| R-HSA-211976 | Endogenous sterols | -0.37 | 0.31 | -0.30 | 0.22 | 0.01 | 0.00 |
| R-HSA-5579013 | Defective CYP7B1 causes SPG5A and CBAS3 | -0.21 | 0.55 | -0.39 | 0.04 | 0.00 | 0.00 |
| R-HSA-5682113 | Defective ABCA1 causes TGD | 0.03 | -0.16 | -0.07 | 0.74 | 0.38 | 0.39 |
| R-HSA-8963896 | HDL assembly | 0.31 | -0.47 | -0.25 | 0.04 | 0.05 | 0.02 |
| R-HSA-8964011 | HDL clearance | 0.03 | -0.16 | -0.10 | 0.74 | 0.37 | 0.38 |
| R-HSA-8964058 | HDL remodeling | 0.04 | -0.22 | -0.04 | 0.73 | 0.38 | 0.39 |
| R-HSA-163358 | PKA-mediated phosphorylation of key metabolic factors | 0.45 | -0.62 | -0.41 | 0.00 | 0.00 | 0.00 |
| R-HSA-9022535 | Loss of phosphorylation of MECP2 at T308 | 0.93 | -0.68 | -0.14 | 0.00 | 0.00 | 0.00 |
| R-HSA-9634600 | Regulation of glycolysis by fructose 2,6-bisphosphate metabolism | 0.25 | -0.52 | -0.42 | 0.04 | 0.00 | 0.00 |
| R-HSA-9026395 | Biosynthesis of DHA-derived sulfido conjugates | 0.77 | 0.56 | -0.40 | 0.00 | 0.00 | 0.00 |
| R-HSA-9026762 | Biosynthesis of maresin conjugates in tissue regeneration (MCTR) | 0.77 | 0.56 | -0.40 | 0.00 | 0.00 | 0.00 |
| R-HSA-1483255 | PI Metabolism | -0.01 | 0.01 | -0.50 | 0.74 | 0.00 | 0.00 |
| R-HSA-1660499 | Synthesis of PIPs at the plasma membrane | -0.01 | 0.01 | -0.61 | 0.74 | 0.00 | 0.00 |
| R-HSA-171319 | Telomere Extension By Telomerase | -0.22 | -0.10 | -0.32 | 0.73 | 0.01 | 0.02 |
| R-HSA-180786 | Extension of Telomeres | -0.22 | -0.10 | -0.34 | 0.73 | 0.00 | 0.01 |
| R-HSA-5694530 | Cargo concentration in the ER | -0.10 | -0.11 | -0.27 | 0.74 | 0.10 | 0.11 |
| R-HSA-163754 | Insulin effects increased synthesis of Xylulose-5-Phosphate | -0.48 | -0.38 | 0.04 | 0.30 | 0.14 | 0.28 |
| R-HSA-5365859 | RA biosynthesis pathway | 0.24 | 0.02 | -0.07 | 0.72 | 0.35 | 0.40 |
| R-HSA-5652084 | Fructose metabolism | 0.11 | 0.08 | -0.25 | 0.74 | 0.12 | 0.12 |
| R-HSA-70350 | Fructose catabolism | 0.03 | 0.23 | -0.36 | 0.73 | 0.00 | 0.00 |
| R-HSA-201451 | Signaling by BMP | -0.19 | -0.43 | -0.38 | 0.58 | 0.00 | 0.00 |
| R-HSA-198725 | Nuclear Events (kinase and transcription factor activation) | 0.21 | -0.12 | -0.31 | 0.71 | 0.01 | 0.03 |
| R-HSA-198753 | ERK/MAPK targets | 0.26 | -0.32 | -0.43 | 0.46 | 0.00 | 0.00 |
| R-HSA-450282 | MAPK targets/ Nuclear events mediated by MAP kinases | 0.26 | -0.32 | -0.47 | 0.46 | 0.00 | 0.00 |
| R-HSA-450341 | Activation of the AP-1 family of transcription factors | 0.46 | -0.63 | -0.47 | 0.00 | 0.00 | 0.00 |
| R-HSA-1187000 | Fertilization | 0.30 | 0.97 | 0.00 | 0.00 | 0.34 | 0.00 |
| R-HSA-1300645 | Acrosome Reaction and Sperm:Oocyte Membrane Binding | 0.30 | 0.97 | -0.20 | 0.00 | 0.14 | 0.00 |
| R-HSA-202670 | ERKs are inactivated | 0.26 | -0.17 | -0.32 | 0.66 | 0.01 | 0.02 |
| R-HSA-140179 | Amine Oxidase reactions | -0.59 | -0.30 | -0.53 | 0.13 | 0.00 | 0.00 |
| R-HSA-141333 | Biogenic amines are oxidatively deaminated to aldehydes by MAOA and MAOB | -0.59 | -0.30 | -0.51 | 0.13 | 0.00 | 0.00 |
| R-HSA-3814836 | Glycogen storage disease type XV (GYG1) | -0.65 | -0.32 | -0.78 | 0.04 | 0.00 | 0.00 |
| R-HSA-3828062 | Glycogen storage disease type 0 (muscle GYS1) | -0.65 | -0.32 | -0.78 | 0.04 | 0.00 | 0.00 |
| R-HSA-211981 | Xenobiotics | -0.31 | -0.50 | -0.12 | 0.36 | 0.27 | 0.12 |
| R-HSA-211999 | CYP2E1 reactions | -0.31 | -0.50 | -0.11 | 0.36 | 0.29 | 0.13 |
| R-HSA-9018682 | Biosynthesis of maresins | -0.31 | -0.53 | -0.20 | 0.28 | 0.15 | 0.03 |
| R-HSA-9027307 | Biosynthesis of maresin-like SPMs | -0.31 | -0.50 | -0.13 | 0.36 | 0.27 | 0.11 |
| R-HSA-8876725 | Protein methylation | 0.42 | -0.05 | -0.20 | 0.52 | 0.05 | 0.28 |
| R-HSA-9678110 | Attachment and Entry | 0.31 | -0.99 | -0.12 | 0.00 | 0.27 | 0.00 |
| R-HSA-9694614 | Attachment and Entry | 0.31 | -0.99 | -0.20 | 0.00 | 0.13 | 0.00 |
| R-HSA-196071 | Metabolism of steroid hormones | 0.02 | -0.37 | -0.16 | 0.63 | 0.34 | 0.21 |
| R-HSA-196108 | Pregnenolone biosynthesis | 0.55 | -0.77 | -0.23 | 0.00 | 0.00 | 0.00 |
| R-HSA-5652227 | Fructose biosynthesis | 0.22 | -0.16 | -0.12 | 0.69 | 0.32 | 0.37 |
| R-HSA-5357786 | TNFR1-induced proapoptotic signaling | 0.33 | -0.27 | -0.49 | 0.41 | 0.00 | 0.00 |
| R-HSA-399955 | SEMA3A-Plexin repulsion signaling by inhibiting Integrin adhesion | 0.29 | -0.30 | -0.29 | 0.44 | 0.02 | 0.03 |
| R-HSA-416550 | Sema4D mediated inhibition of cell attachment and migration | 0.21 | -0.08 | -0.35 | 0.72 | 0.00 | 0.01 |
| R-HSA-198203 | PI3K/AKT activation | -0.17 | -0.49 | -0.43 | 0.46 | 0.00 | 0.00 |
| R-HSA-5666185 | RHO GTPases Activate Rhotekin and Rhophilins | -0.05 | 0.20 | -0.32 | 0.73 | 0.03 | 0.01 |
| R-HSA-8985586 | SLIT2:ROBO1 increases RHOA activity | -0.17 | -0.49 | -0.43 | 0.46 | 0.00 | 0.00 |
| R-HSA-1592389 | Activation of Matrix Metalloproteinases | 0.43 | 0.31 | 0.17 | 0.50 | 0.09 | 0.24 |
| R-HSA-9690406 | Transcriptional regulation of testis differentiation | 0.29 | 0.42 | -0.17 | 0.53 | 0.21 | 0.07 |
| R-HSA-2029485 | Role of phospholipids in phagocytosis | 0.76 | -0.33 | -0.55 | 0.00 | 0.00 | 0.00 |
| R-HSA-6811555 | PI5P Regulates TP53 Acetylation | 0.50 | 0.57 | -0.48 | 0.04 | 0.00 | 0.00 |
| R-HSA-73843 | 5-Phosphoribose 1-diphosphate biosynthesis | 0.41 | -0.64 | -0.14 | 0.00 | 0.13 | 0.01 |
| R-HSA-5619050 | Defective SLC4A1 causes hereditary spherocytosis type 4 (HSP4), distal renal tubular acidosis (dRTA) and dRTA with hemolytic anemia (dRTA-HA) | -0.57 | -0.13 | 0.11 | 0.22 | 0.03 | 0.38 |
| R-HSA-918233 | TRAF3-dependent IRF activation pathway | -0.61 | -0.12 | -0.32 | 0.10 | 0.00 | 0.02 |
| R-HSA-933541 | TRAF6 mediated IRF7 activation | -0.61 | -0.12 | -0.34 | 0.10 | 0.00 | 0.01 |
| R-HSA-933542 | TRAF6 mediated NF-kB activation | -0.52 | -0.31 | -0.54 | 0.27 | 0.00 | 0.00 |
| R-HSA-933543 | NF-kB activation through FADD/RIP-1 pathway mediated by caspase-8 and -10 | -0.61 | -0.12 | -0.62 | 0.10 | 0.00 | 0.00 |
| R-HSA-391160 | Signal regulatory protein family interactions | -0.50 | -0.71 | -0.21 | 0.00 | 0.02 | 0.00 |
| R-HSA-112308 | Presynaptic depolarization and calcium channel opening | -0.69 | -0.24 | -0.14 | 0.02 | 0.00 | 0.33 |
| R-HSA-8866423 | VLDL assembly | 0.37 | -1.03 | 0.03 | 0.00 | 0.29 | 0.00 |
| R-HSA-9020933 | Interleukin-23 signaling | 0.36 | -0.44 | -0.41 | 0.04 | 0.00 | 0.00 |
| R-HSA-202433 | Generation of second messenger molecules | -0.31 | -0.47 | -0.35 | 0.41 | 0.00 | 0.00 |
| R-HSA-1482883 | Acyl chain remodeling of DAG and TAG | -0.72 | 0.86 | 0.25 | 0.00 | 0.00 | 0.00 |
| R-HSA-426048 | Arachidonate production from DAG | -0.72 | 0.86 | -0.50 | 0.00 | 0.00 | 0.00 |
| R-HSA-1663150 | The activation of arylsulfatases | 0.31 | -0.40 | -0.41 | 0.16 | 0.00 | 0.00 |
| R-HSA-8941413 | Events associated with phagocytolytic activity of PMN cells | -0.88 | -0.07 | 0.42 | 0.00 | 0.00 | 0.00 |
| R-HSA-203641 | NOSTRIN mediated eNOS trafficking | 0.05 | 0.03 | -0.36 | 0.74 | 0.00 | 0.00 |
| R-HSA-4641262 | Disassembly of the destruction complex and recruitment of AXIN to the membrane | -0.02 | -0.37 | -0.17 | 0.65 | 0.33 | 0.19 |
| R-HSA-9617828 | FOXO-mediated transcription of cell cycle genes | 0.22 | -1.00 | -0.22 | 0.00 | 0.14 | 0.00 |
| R-HSA-113501 | Inhibition of replication initiation of damaged DNA by RB1/E2F1 | 0.04 | -0.42 | -0.45 | 0.55 | 0.00 | 0.00 |
| R-HSA-113510 | E2F mediated regulation of DNA replication | 0.04 | -0.42 | -0.38 | 0.55 | 0.00 | 0.00 |
| R-HSA-163767 | PP2A-mediated dephosphorylation of key metabolic factors | 0.04 | -0.42 | -0.30 | 0.55 | 0.04 | 0.01 |
| R-HSA-196299 | Beta-catenin phosphorylation cascade | -0.08 | -0.22 | -0.15 | 0.73 | 0.34 | 0.33 |
| R-HSA-199418 | Negative regulation of the PI3K/AKT network | 0.26 | -0.06 | -0.29 | 0.71 | 0.02 | 0.06 |
| R-HSA-2465910 | MASTL Facilitates Mitotic Progression | 0.04 | -0.42 | -0.03 | 0.55 | 0.38 | 0.26 |
| R-HSA-389513 | CTLA4 inhibitory signaling | 0.19 | -0.10 | -0.50 | 0.73 | 0.00 | 0.00 |
| R-HSA-4791275 | Signaling by WNT in cancer | -0.08 | -0.22 | -0.24 | 0.73 | 0.17 | 0.15 |
| R-HSA-4839735 | Signaling by AXIN mutants | 0.19 | -0.10 | -0.13 | 0.73 | 0.33 | 0.37 |
| R-HSA-4839743 | Signaling by CTNNB1 phospho-site mutants | -0.08 | -0.22 | -0.16 | 0.73 | 0.33 | 0.31 |
| R-HSA-4839744 | Signaling by APC mutants | 0.19 | -0.10 | -0.13 | 0.73 | 0.33 | 0.37 |
| R-HSA-4839748 | Signaling by AMER1 mutants | 0.19 | -0.10 | -0.13 | 0.73 | 0.33 | 0.37 |
| R-HSA-5339716 | Signaling by GSK3beta mutants | -0.08 | -0.22 | -0.16 | 0.73 | 0.33 | 0.31 |
| R-HSA-5358747 | S33 mutants of beta-catenin aren't phosphorylated | -0.08 | -0.22 | -0.16 | 0.73 | 0.33 | 0.31 |
| R-HSA-5358749 | S37 mutants of beta-catenin aren't phosphorylated | -0.08 | -0.22 | -0.16 | 0.73 | 0.33 | 0.31 |
| R-HSA-5358751 | S45 mutants of beta-catenin aren't phosphorylated | -0.08 | -0.22 | -0.16 | 0.73 | 0.33 | 0.31 |
| R-HSA-5358752 | T41 mutants of beta-catenin aren't phosphorylated | -0.08 | -0.22 | -0.16 | 0.73 | 0.33 | 0.31 |
| R-HSA-5467337 | APC truncation mutants have impaired AXIN binding | 0.19 | -0.10 | -0.13 | 0.73 | 0.33 | 0.37 |
| R-HSA-5467340 | AXIN missense mutants destabilize the destruction complex | 0.19 | -0.10 | -0.13 | 0.73 | 0.33 | 0.37 |
| R-HSA-5467348 | Truncations of AMER1 destabilize the destruction complex | 0.19 | -0.10 | -0.13 | 0.73 | 0.33 | 0.37 |
| R-HSA-6811558 | PI5P, PP2A and IER3 Regulate PI3K/AKT Signaling | 0.26 | -0.06 | -0.28 | 0.71 | 0.03 | 0.09 |
| R-HSA-3214842 | HDMs demethylate histones | -0.09 | 0.67 | -0.01 | 0.01 | 0.38 | 0.01 |
| R-HSA-3134973 | LRR FLII-interacting protein 1 (LRRFIP1) activates type I IFN production | -0.89 | -0.55 | -0.66 | 0.00 | 0.00 | 0.00 |
| R-HSA-4411364 | Binding of TCF/LEF:CTNNB1 to target gene promoters | -0.89 | -0.55 | -0.19 | 0.00 | 0.00 | 0.02 |
| R-HSA-8853884 | Transcriptional Regulation by VENTX | -0.89 | -0.55 | -0.27 | 0.00 | 0.00 | 0.00 |
| R-HSA-8951430 | RUNX3 regulates WNT signaling | -0.89 | -0.55 | -0.27 | 0.00 | 0.00 | 0.00 |
| R-HSA-446388 | Regulation of cytoskeletal remodeling and cell spreading by IPP complex components | -0.39 | 0.32 | -0.60 | 0.15 | 0.00 | 0.00 |
| R-HSA-1855167 | Synthesis of pyrophosphates in the cytosol | 0.56 | -0.40 | -0.41 | 0.00 | 0.00 | 0.00 |
| R-HSA-447038 | NrCAM interactions | -0.44 | 0.59 | -0.41 | 0.00 | 0.00 | 0.00 |
| R-HSA-447041 | CHL1 interactions | 0.19 | -0.13 | -0.36 | 0.72 | 0.00 | 0.00 |
| R-HSA-159418 | Recycling of bile acids and salts | 0.08 | 0.08 | -0.18 | 0.74 | 0.30 | 0.30 |
| R-HSA-879518 | Transport of organic anions | 0.08 | 0.08 | -0.19 | 0.74 | 0.29 | 0.30 |
| R-HSA-3065676 | SUMO is conjugated to E1 (UBA2:SAE1) | 0.13 | 0.60 | -0.05 | 0.15 | 0.38 | 0.03 |
| R-HSA-3065678 | SUMO is transferred from E1 to E2 (UBE2I, UBC9) | 0.13 | 0.60 | -0.10 | 0.15 | 0.37 | 0.01 |
| R-HSA-3215018 | Processing and activation of SUMO | 0.13 | 0.60 | -0.12 | 0.15 | 0.36 | 0.01 |
| R-HSA-2142845 | Hyaluronan metabolism | -0.41 | 0.77 | -0.32 | 0.00 | 0.00 | 0.00 |
| R-HSA-2160916 | Hyaluronan uptake and degradation | -0.41 | 0.77 | -0.40 | 0.00 | 0.00 | 0.00 |
| R-HSA-2408508 | Metabolism of ingested SeMet, Sec, MeSec into H2Se | -0.02 | 0.19 | -0.31 | 0.73 | 0.04 | 0.01 |
| R-HSA-5578997 | Defective AHCY causes HMAHCHD | -0.52 | 0.22 | -0.06 | 0.05 | 0.09 | 0.38 |
| R-HSA-5659996 | RPIA deficiency: failed conversion of R5P to RU5P | 0.68 | 0.84 | -0.05 | 0.00 | 0.01 | 0.00 |
| R-HSA-6791461 | RPIA deficiency: failed conversion of RU5P to R5P | 0.68 | 0.84 | -0.05 | 0.00 | 0.01 | 0.00 |
| R-HSA-6791465 | Pentose phosphate pathway disease | 0.08 | 0.24 | -0.02 | 0.73 | 0.38 | 0.38 |
| R-HSA-5223345 | Miscellaneous transport and binding events | -0.19 | -0.82 | -0.39 | 0.00 | 0.00 | 0.00 |
| R-HSA-1483148 | Synthesis of PG | -0.47 | 0.20 | -0.21 | 0.18 | 0.03 | 0.17 |
| R-HSA-1059683 | Interleukin-6 signaling | 0.61 | 0.34 | -0.62 | 0.07 | 0.00 | 0.00 |
| R-HSA-111453 | BH3-only proteins associate with and inactivate anti-apoptotic BCL-2 members | 0.34 | 0.14 | -0.18 | 0.68 | 0.14 | 0.28 |
| R-HSA-1266695 | Interleukin-7 signaling | 0.34 | 0.14 | -0.09 | 0.68 | 0.28 | 0.39 |
| R-HSA-1839117 | Signaling by cytosolic FGFR1 fusion mutants | 0.26 | -0.03 | -0.69 | 0.71 | 0.00 | 0.00 |
| R-HSA-1839124 | FGFR1 mutant receptor activation | 0.26 | -0.03 | -0.35 | 0.71 | 0.00 | 0.01 |
| R-HSA-198745 | Signalling to STAT3 | 0.34 | 0.14 | -0.16 | 0.68 | 0.19 | 0.32 |
| R-HSA-201556 | Signaling by ALK | 0.34 | 0.14 | -0.47 | 0.68 | 0.00 | 0.00 |
| R-HSA-2586552 | Signaling by Leptin | 0.34 | 0.14 | -0.46 | 0.68 | 0.00 | 0.00 |
| R-HSA-2892247 | POU5F1 (OCT4), SOX2, NANOG activate genes related to proliferation | 0.34 | 0.14 | 0.17 | 0.68 | 0.18 | 0.33 |
| R-HSA-451927 | Interleukin-2 family signaling | 0.26 | -0.03 | -0.45 | 0.71 | 0.00 | 0.00 |
| R-HSA-452723 | Transcriptional regulation of pluripotent stem cells | 0.34 | 0.14 | 0.07 | 0.68 | 0.30 | 0.39 |
| R-HSA-6783589 | Interleukin-6 family signaling | 0.61 | 0.34 | -0.41 | 0.07 | 0.00 | 0.00 |
| R-HSA-6783783 | Interleukin-10 signaling | 0.34 | 0.14 | -0.27 | 0.68 | 0.02 | 0.06 |
| R-HSA-8849474 | PTK6 Activates STAT3 | 0.34 | 0.14 | -0.30 | 0.68 | 0.01 | 0.03 |
| R-HSA-8854691 | Interleukin-20 family signaling | 0.61 | 0.34 | -0.30 | 0.07 | 0.00 | 0.00 |
| R-HSA-8875791 | MET activates STAT3 | 0.34 | 0.14 | -0.58 | 0.68 | 0.00 | 0.00 |
| R-HSA-8983432 | Interleukin-15 signaling | -0.06 | -0.31 | -0.69 | 0.70 | 0.00 | 0.00 |
| R-HSA-8985947 | Interleukin-9 signaling | 0.61 | 0.34 | -0.66 | 0.07 | 0.00 | 0.00 |
| R-HSA-9008059 | Interleukin-37 signaling | 0.34 | 0.14 | -0.23 | 0.68 | 0.06 | 0.17 |
| R-HSA-9020958 | Interleukin-21 signaling | 0.61 | 0.34 | -0.44 | 0.07 | 0.00 | 0.00 |
| R-HSA-9701898 | STAT3 nuclear events downstream of ALK signaling | 0.34 | 0.14 | -0.55 | 0.68 | 0.00 | 0.00 |
| R-HSA-982772 | Growth hormone receptor signaling | 0.59 | -0.05 | 0.02 | 0.08 | 0.04 | 0.40 |
| R-HSA-5683826 | Surfactant metabolism | 0.16 | -0.03 | -0.20 | 0.74 | 0.23 | 0.28 |
| R-HSA-8847453 | Synthesis of PIPs in the nucleus | 0.54 | 0.88 | -0.39 | 0.00 | 0.00 | 0.00 |
| R-HSA-111446 | Activation of BIM and translocation to mitochondria | 0.03 | 0.79 | 0.06 | 0.00 | 0.38 | 0.00 |
| R-HSA-2142670 | Synthesis of epoxy (EET) and dihydroxyeicosatrienoic acids (DHET) | -0.19 | 0.02 | -0.21 | 0.73 | 0.20 | 0.25 |
| R-HSA-392154 | Nitric oxide stimulates guanylate cyclase | 0.09 | -0.55 | -0.31 | 0.14 | 0.03 | 0.00 |
| R-HSA-418457 | cGMP effects | 0.09 | -0.55 | -0.35 | 0.14 | 0.00 | 0.00 |
| R-HSA-1251985 | Nuclear signaling by ERBB4 | 0.24 | -0.82 | -0.27 | 0.00 | 0.05 | 0.00 |
| R-HSA-8864260 | Transcriptional regulation by the AP-2 (TFAP2) family of transcription factors | 0.24 | -0.82 | -0.20 | 0.00 | 0.18 | 0.00 |
| R-HSA-8964026 | Chylomicron clearance | 0.24 | -0.82 | -0.18 | 0.00 | 0.22 | 0.00 |
| R-HSA-2022928 | HS-GAG biosynthesis | 0.02 | 0.25 | -0.35 | 0.72 | 0.01 | 0.00 |
| R-HSA-2024096 | HS-GAG degradation | -0.14 | 0.03 | -0.36 | 0.74 | 0.00 | 0.00 |
| R-HSA-3656237 | Defective EXT2 causes exostoses 2 | 0.02 | 0.25 | -0.42 | 0.72 | 0.00 | 0.00 |
| R-HSA-3656253 | Defective EXT1 causes exostoses 1, TRPS2 and CHDS | 0.02 | 0.25 | -0.42 | 0.72 | 0.00 | 0.00 |
| R-HSA-193144 | Estrogen biosynthesis | -0.76 | 0.36 | 0.01 | 0.00 | 0.00 | 0.32 |
| R-HSA-5579030 | Defective CYP19A1 causes AEXS | -0.76 | 0.36 | 0.28 | 0.00 | 0.00 | 0.03 |
| R-HSA-6787639 | GDP-fucose biosynthesis | -0.44 | -0.21 | -0.35 | 0.55 | 0.00 | 0.01 |
| R-HSA-5660668 | CLEC7A/inflammasome pathway | -0.57 | -0.16 | -0.59 | 0.22 | 0.00 | 0.00 |
| R-HSA-844615 | The AIM2 inflammasome | -0.57 | -0.16 | -0.12 | 0.22 | 0.03 | 0.37 |
| R-HSA-425561 | Sodium/Calcium exchangers | -0.03 | 0.33 | -0.13 | 0.68 | 0.36 | 0.24 |
| R-HSA-5619056 | Defective HK1 causes hexokinase deficiency (HK deficiency) | -0.06 | 0.60 | -0.66 | 0.06 | 0.00 | 0.00 |
| R-HSA-72200 | mRNA Editing: C to U Conversion | -0.65 | -1.04 | 0.09 | 0.00 | 0.01 | 0.00 |
| R-HSA-75072 | mRNA Editing | -0.65 | -1.04 | -0.12 | 0.00 | 0.01 | 0.00 |
| R-HSA-75094 | Formation of the Editosome | -0.65 | -1.04 | 0.09 | 0.00 | 0.01 | 0.00 |
| R-HSA-877312 | Regulation of IFNG signaling | 0.89 | 0.53 | -0.26 | 0.00 | 0.00 | 0.00 |
| R-HSA-9013508 | NOTCH3 Intracellular Domain Regulates Transcription | 0.38 | 0.40 | -0.34 | 0.46 | 0.00 | 0.00 |
| R-HSA-912694 | Regulation of IFNA signaling | 0.89 | 0.53 | -0.21 | 0.00 | 0.00 | 0.00 |
| R-HSA-111932 | CaMK IV-mediated phosphorylation of CREB | 0.14 | -0.67 | -0.43 | 0.00 | 0.00 | 0.00 |
| R-HSA-442729 | CREB1 phosphorylation through the activation of CaMKII/CaMKK/CaMKIV cascasde | 0.14 | -0.67 | -0.40 | 0.00 | 0.00 | 0.00 |
| R-HSA-5576892 | Phase 0 - rapid depolarisation | 0.14 | -0.67 | -0.12 | 0.00 | 0.36 | 0.01 |
| R-HSA-194306 | Neurophilin interactions with VEGF and VEGFR | 0.83 | -0.85 | -0.48 | 0.00 | 0.00 | 0.00 |
| R-HSA-2978092 | Abnormal conversion of 2-oxoglutarate to 2-hydroxyglutarate | 0.23 | -0.01 | -0.18 | 0.73 | 0.22 | 0.32 |
| R-HSA-389542 | NADPH regeneration | 0.23 | -0.01 | -0.18 | 0.73 | 0.22 | 0.32 |
| R-HSA-5661270 | Formation of xylulose-5-phosphate | 0.45 | 0.46 | -0.43 | 0.23 | 0.00 | 0.00 |
| R-HSA-450513 | Tristetraprolin (TTP, ZFP36) binds and destabilizes mRNA | -0.13 | -0.42 | -0.30 | 0.60 | 0.04 | 0.01 |
| R-HSA-9636249 | Inhibition of nitric oxide production | 0.38 | -0.51 | -0.77 | 0.00 | 0.00 | 0.00 |
| R-HSA-389599 | Alpha-oxidation of phytanate | -0.51 | -0.10 | -0.23 | 0.39 | 0.01 | 0.21 |
| R-HSA-6811438 | Intra-Golgi traffic | -0.40 | -0.62 | -0.45 | 0.04 | 0.00 | 0.00 |
| R-HSA-9664535 | LTC4-CYSLTR mediated IL4 production | -0.12 | 0.53 | -0.24 | 0.16 | 0.16 | 0.00 |
| R-HSA-9657688 | Defective factor XII causes hereditary angioedema | 0.62 | 0.39 | 0.09 | 0.04 | 0.02 | 0.27 |
| R-HSA-9672391 | Defective F8 cleavage by thrombin | 0.62 | 0.39 | -0.07 | 0.04 | 0.02 | 0.24 |
| R-HSA-9031628 | NGF-stimulated transcription | -0.11 | 1.05 | -0.25 | 0.00 | 0.14 | 0.00 |
| R-HSA-164944 | Nef and signal transduction | -0.01 | 0.05 | -0.28 | 0.74 | 0.09 | 0.08 |
| R-HSA-2219528 | PI3K/AKT Signaling in Cancer | -0.07 | -0.10 | -0.25 | 0.74 | 0.14 | 0.15 |
| R-HSA-2219530 | Constitutive Signaling by Aberrant PI3K in Cancer | -0.07 | -0.10 | -0.23 | 0.74 | 0.20 | 0.22 |
| R-HSA-428540 | Activation of RAC1 | -0.01 | 0.05 | -0.37 | 0.74 | 0.00 | 0.00 |
| R-HSA-9032500 | Activated NTRK2 signals through FYN | 0.30 | 0.57 | -0.59 | 0.19 | 0.00 | 0.00 |
| R-HSA-9032759 | NTRK2 activates RAC1 | 0.30 | 0.57 | -0.49 | 0.19 | 0.00 | 0.00 |
| R-HSA-9032845 | Activated NTRK2 signals through CDK5 | 0.30 | 0.57 | -0.26 | 0.19 | 0.03 | 0.00 |
| R-HSA-9619229 | Activation of RAC1 downstream of NMDARs | 0.30 | 0.57 | -0.17 | 0.19 | 0.20 | 0.01 |
| R-HSA-9664420 | Killing mechanisms | 0.30 | 0.57 | -0.38 | 0.19 | 0.00 | 0.00 |
| R-HSA-9673324 | WNT5:FZD7-mediated leishmania damping | 0.30 | 0.57 | -0.38 | 0.19 | 0.00 | 0.00 |
| R-HSA-166020 | Transfer of LPS from LBP carrier to CD14 | -0.34 | -0.24 | -0.36 | 0.66 | 0.00 | 0.00 |
| R-HSA-418889 | Caspase activation via Dependence Receptors in the absence of ligand | -0.36 | -0.26 | -0.41 | 0.63 | 0.00 | 0.00 |
| R-HSA-5357769 | Caspase activation via extrinsic apoptotic signalling pathway | -0.36 | -0.26 | -0.27 | 0.63 | 0.02 | 0.07 |
| R-HSA-163765 | ChREBP activates metabolic gene expression | 0.45 | 0.38 | -0.64 | 0.36 | 0.00 | 0.00 |
| R-HSA-6785470 | tRNA processing in the mitochondrion | 0.19 | 0.86 | -0.29 | 0.00 | 0.04 | 0.00 |
| R-HSA-6787450 | tRNA modification in the mitochondrion | 0.19 | 0.86 | -0.36 | 0.00 | 0.00 | 0.00 |
| R-HSA-8868766 | rRNA processing in the mitochondrion | 0.19 | 0.86 | -0.38 | 0.00 | 0.00 | 0.00 |
| R-HSA-2206281 | Mucopolysaccharidoses | -0.46 | -0.42 | -0.46 | 0.29 | 0.00 | 0.00 |
| R-HSA-2206308 | MPS IV - Morquio syndrome B | -0.46 | -0.42 | -0.30 | 0.29 | 0.00 | 0.01 |
| R-HSA-4341670 | Defective NEU1 causes sialidosis | -0.46 | -0.42 | -0.10 | 0.29 | 0.14 | 0.23 |
| R-HSA-196791 | Vitamin D (calciferol) metabolism | 0.17 | 0.68 | -0.18 | 0.03 | 0.26 | 0.00 |
| R-HSA-5609976 | Defective GALK1 can cause Galactosemia II (GALCT2) | 0.15 | -0.59 | 0.03 | 0.03 | 0.38 | 0.04 |
| R-HSA-8853383 | Lysosomal oligosaccharide catabolism | 0.15 | -0.75 | 0.17 | 0.00 | 0.30 | 0.00 |
| R-HSA-5578995 | Defective TPMT causes TPMT deficiency | 0.01 | -0.85 | -0.24 | 0.00 | 0.18 | 0.00 |
| R-HSA-189085 | Digestion of dietary carbohydrate | 0.31 | 0.34 | 0.10 | 0.62 | 0.30 | 0.31 |
| R-HSA-8935690 | Digestion | -0.02 | 0.24 | 0.41 | 0.72 | 0.00 | 0.00 |
| R-HSA-8963743 | Digestion and absorption | -0.02 | 0.24 | 0.37 | 0.72 | 0.00 | 0.00 |
| R-HSA-450385 | Butyrate Response Factor 1 (BRF1) binds and destabilizes mRNA | -0.16 | 0.06 | -0.28 | 0.74 | 0.06 | 0.07 |
| R-HSA-9660537 | Signaling by MRAS-complex mutants | 0.17 | -0.25 | -0.50 | 0.67 | 0.00 | 0.00 |
| R-HSA-9726840 | SHOC2 M1731 mutant abolishes MRAS complex function | 0.17 | -0.25 | -0.50 | 0.67 | 0.00 | 0.00 |
| R-HSA-9726842 | Gain-of-function MRAS complexes activate RAF signaling | 0.17 | -0.25 | -0.50 | 0.67 | 0.00 | 0.00 |
| R-HSA-5657560 | Hereditary fructose intolerance | -0.40 | 0.69 | -0.26 | 0.00 | 0.02 | 0.00 |
| R-HSA-139910 | Activation of BMF and translocation to mitochondria | -0.28 | 0.60 | -0.68 | 0.00 | 0.00 | 0.00 |
| R-HSA-205025 | NADE modulates death signalling | -0.30 | -0.77 | -0.33 | 0.00 | 0.00 | 0.00 |
| R-HSA-2142816 | Synthesis of (16-20)-hydroxyeicosatetraenoic acids (HETE) | -0.08 | 0.67 | 0.22 | 0.01 | 0.22 | 0.00 |
| R-HSA-5579000 | Defective CYP1B1 causes Glaucoma | -0.08 | 0.67 | -0.37 | 0.01 | 0.00 | 0.00 |
| R-HSA-1660514 | Synthesis of PIPs at the Golgi membrane | -0.58 | -0.98 | -0.37 | 0.00 | 0.00 | 0.00 |
| R-HSA-181430 | Norepinephrine Neurotransmitter Release Cycle | -0.42 | 0.41 | -0.20 | 0.02 | 0.06 | 0.03 |
| R-HSA-380612 | Metabolism of serotonin | -0.42 | 0.41 | -0.35 | 0.02 | 0.00 | 0.00 |
| R-HSA-380615 | Serotonin clearance from the synaptic cleft | -0.42 | 0.41 | -0.47 | 0.02 | 0.00 | 0.00 |
| R-HSA-5579012 | Defective MAOA causes BRUNS | -0.42 | 0.41 | -0.35 | 0.02 | 0.00 | 0.00 |
| R-HSA-4043916 | Defective MPI causes MPI-CDG (CDG-1b) | 0.95 | 0.38 | -0.59 | 0.00 | 0.00 | 0.00 |
| R-HSA-70921 | Histidine catabolism | 0.49 | 0.16 | -0.19 | 0.45 | 0.03 | 0.25 |
| R-HSA-9018681 | Biosynthesis of protectins | 0.18 | 0.11 | 0.08 | 0.74 | 0.37 | 0.40 |
| R-HSA-196741 | Cobalamin (Cbl, vitamin B12) transport and metabolism | -0.47 | -0.96 | -0.33 | 0.00 | 0.00 | 0.00 |
| R-HSA-3296469 | Defects in cobalamin (B12) metabolism | -0.47 | -0.96 | -0.39 | 0.00 | 0.00 | 0.00 |
| R-HSA-3359475 | Defective MMAA causes methylmalonic aciduria type cblA | -0.47 | -0.96 | -0.71 | 0.00 | 0.00 | 0.00 |
| R-HSA-3359478 | Defective MUT causes methylmalonic aciduria mut type | -0.47 | -0.96 | -0.71 | 0.00 | 0.00 | 0.00 |
| R-HSA-8849468 | PTK6 Regulates Proteins Involved in RNA Processing | -0.85 | 0.46 | -0.05 | 0.00 | 0.00 | 0.17 |
| R-HSA-9635465 | Suppression of apoptosis | 0.09 | -0.13 | -0.01 | 0.74 | 0.38 | 0.40 |
| R-HSA-174577 | Activation of C3 and C5 | -0.28 | -0.75 | -0.38 | 0.00 | 0.00 | 0.00 |
| R-HSA-6791055 | TALDO1 deficiency: failed conversion of SH7P, GA3P to Fru(6)P, E4P | -0.51 | -0.36 | 0.05 | 0.25 | 0.11 | 0.29 |
| R-HSA-6791462 | TALDO1 deficiency: failed conversion of Fru(6)P, E4P to SH7P, GA3P | -0.51 | -0.36 | 0.05 | 0.25 | 0.11 | 0.29 |
| R-HSA-112382 | Formation of RNA Pol II elongation complex | 0.06 | -0.50 | -0.29 | 0.31 | 0.05 | 0.00 |
| R-HSA-167152 | Formation of HIV elongation complex in the absence of HIV Tat | 0.06 | -0.50 | -0.27 | 0.31 | 0.10 | 0.01 |
| R-HSA-167169 | HIV Transcription Elongation | 0.06 | -0.50 | -0.26 | 0.31 | 0.12 | 0.01 |
| R-HSA-167172 | Transcription of the HIV genome | 0.06 | -0.50 | -0.24 | 0.31 | 0.17 | 0.02 |
| R-HSA-167200 | Formation of HIV-1 elongation complex containing HIV-1 Tat | 0.06 | -0.50 | -0.26 | 0.31 | 0.12 | 0.01 |
| R-HSA-167238 | Pausing and recovery of Tat-mediated HIV elongation | 0.06 | -0.50 | -0.21 | 0.31 | 0.24 | 0.03 |
| R-HSA-167243 | Tat-mediated HIV elongation arrest and recovery | 0.06 | -0.50 | -0.21 | 0.31 | 0.24 | 0.03 |
| R-HSA-167246 | Tat-mediated elongation of the HIV-1 transcript | 0.06 | -0.50 | -0.26 | 0.31 | 0.12 | 0.01 |
| R-HSA-167287 | HIV elongation arrest and recovery | 0.06 | -0.50 | -0.23 | 0.31 | 0.20 | 0.02 |
| R-HSA-167290 | Pausing and recovery of HIV elongation | 0.06 | -0.50 | -0.23 | 0.31 | 0.20 | 0.02 |
| R-HSA-674695 | RNA Polymerase II Pre-transcription Events | 0.06 | -0.50 | -0.26 | 0.31 | 0.11 | 0.01 |
| R-HSA-6796648 | TP53 Regulates Transcription of DNA Repair Genes | 0.06 | -0.50 | -0.34 | 0.31 | 0.01 | 0.00 |
| R-HSA-75955 | RNA Polymerase II Transcription Elongation | 0.06 | -0.50 | -0.29 | 0.31 | 0.05 | 0.00 |
| R-HSA-4085023 | Defective GFPT1 causes CMSTA1 | -0.31 | -0.30 | -0.85 | 0.65 | 0.00 | 0.00 |
| R-HSA-211728 | Regulation of PAK-2p34 activity by PS-GAP/RHG10 | -0.31 | -0.47 | -0.48 | 0.41 | 0.00 | 0.00 |
| R-HSA-211736 | Stimulation of the cell death response by PAK-2p34 | -0.31 | -0.47 | -0.87 | 0.41 | 0.00 | 0.00 |
| R-HSA-5624138 | Trafficking of myristoylated proteins to the cilium | 0.01 | 0.06 | -0.60 | 0.74 | 0.00 | 0.00 |
| R-HSA-6783984 | Glycine degradation | 0.45 | -0.78 | -0.05 | 0.00 | 0.17 | 0.00 |
| R-HSA-3772470 | Negative regulation of TCF-dependent signaling by WNT ligand antagonists | 0.11 | 0.23 | -0.27 | 0.73 | 0.09 | 0.04 |
| R-HSA-109704 | PI3K Cascade | -0.45 | -0.76 | -0.14 | 0.00 | 0.11 | 0.00 |
| R-HSA-180292 | GAB1 signalosome | -0.45 | -0.76 | -0.40 | 0.00 | 0.00 | 0.00 |
| R-HSA-1963642 | PI3K events in ERBB2 signaling | -0.45 | -0.76 | -0.32 | 0.00 | 0.00 | 0.00 |
| R-HSA-2730905 | Role of LAT2/NTAL/LAB on calcium mobilization | -0.45 | -0.76 | -0.37 | 0.00 | 0.00 | 0.00 |
| R-HSA-5654689 | PI-3K cascade:FGFR1 | -0.45 | -0.76 | -0.07 | 0.00 | 0.18 | 0.00 |
| R-HSA-5654695 | PI-3K cascade:FGFR2 | -0.45 | -0.76 | -0.09 | 0.00 | 0.17 | 0.00 |
| R-HSA-5654710 | PI-3K cascade:FGFR3 | -0.45 | -0.76 | -0.18 | 0.00 | 0.07 | 0.00 |
| R-HSA-5654720 | PI-3K cascade:FGFR4 | -0.45 | -0.76 | -0.06 | 0.00 | 0.18 | 0.00 |
| R-HSA-74749 | Signal attenuation | 0.22 | -0.54 | -0.34 | 0.04 | 0.00 | 0.00 |
| R-HSA-8851907 | MET activates PI3K/AKT signaling | -0.45 | -0.76 | -0.79 | 0.00 | 0.00 | 0.00 |
| R-HSA-8865999 | MET activates PTPN11 | -0.45 | -0.76 | -0.78 | 0.00 | 0.00 | 0.00 |
| R-HSA-9028335 | Activated NTRK2 signals through PI3K | -0.45 | -0.76 | -0.60 | 0.00 | 0.00 | 0.00 |
| R-HSA-912526 | Interleukin receptor SHC signaling | -0.45 | -0.76 | -0.42 | 0.00 | 0.00 | 0.00 |
| R-HSA-9645135 | STAT5 Activation | -0.45 | -0.76 | -0.46 | 0.00 | 0.00 | 0.00 |
| R-HSA-9702518 | STAT5 activation downstream of FLT3 ITD mutants | -0.45 | -0.76 | -0.22 | 0.00 | 0.03 | 0.00 |
| R-HSA-9022699 | MECP2 regulates neuronal receptors and channels | -0.16 | 0.42 | -0.10 | 0.40 | 0.36 | 0.16 |
| R-HSA-193775 | Synthesis of bile acids and bile salts via 24-hydroxycholesterol | -0.42 | -0.34 | -0.34 | 0.48 | 0.00 | 0.00 |
| R-HSA-211994 | Sterols are 12-hydroxylated by CYP8B1 | -0.42 | -0.34 | -0.33 | 0.48 | 0.00 | 0.01 |
| R-HSA-8937144 | Aryl hydrocarbon receptor signalling | -0.75 | -0.04 | 0.05 | 0.00 | 0.00 | 0.40 |
| R-HSA-111995 | phospho-PLA2 pathway | 0.96 | 0.17 | -0.65 | 0.00 | 0.00 | 0.00 |
| R-HSA-112411 | MAPK1 (ERK2) activation | 0.96 | 0.17 | -0.64 | 0.00 | 0.00 | 0.00 |
| R-HSA-444257 | RSK activation | 0.56 | -0.43 | -0.25 | 0.00 | 0.00 | 0.04 |
| R-HSA-9652817 | Signaling by MAPK mutants | 0.96 | 0.17 | -0.38 | 0.00 | 0.00 | 0.00 |
| R-HSA-193048 | Androgen biosynthesis | -0.24 | -0.29 | -0.18 | 0.69 | 0.24 | 0.24 |
| R-HSA-5578999 | Defective GCLC causes HAGGSD | 0.13 | 0.07 | -0.38 | 0.74 | 0.00 | 0.00 |
| R-HSA-2142700 | Synthesis of Lipoxins (LX) | 0.01 | 0.33 | -0.12 | 0.69 | 0.37 | 0.24 |
| R-HSA-8964572 | Lipid particle organization | -0.22 | -0.77 | -0.22 | 0.00 | 0.17 | 0.00 |
| R-HSA-9029558 | NR1H2 & NR1H3 regulate gene expression linked to lipogenesis | -0.05 | 0.51 | -0.30 | 0.29 | 0.04 | 0.00 |
| R-HSA-5263617 | Metabolism of ingested MeSeO2H into MeSeH | -0.30 | -0.77 | -0.54 | 0.00 | 0.00 | 0.00 |
| R-HSA-9645722 | Defective Intrinsic Pathway for Apoptosis Due to p14ARF Loss of Function | -0.29 | -0.28 | 0.13 | 0.67 | 0.26 | 0.27 |

Table 1: The list of pathways from the integrated analysis of EL, TL proteomics and transcriptomics integrated analysis.

| **Pathway Name** | **Stable Identifier** |
| --- | --- |
| Prefoldin mediated transfer of substrate to CCT/TriC | (R-HSA-389957) |
| Cooperation of Prefoldin and TriC/CCT in actin and tubulin folding | (R-HSA- 389958)) |
| Respiratory electron transport, ATP synthesis by chemiosmotic coupling, and  heat production by uncoupling proteins. | (R-HSA-163200) |
| Respiratory electron transport | (R-HSA-611105) |
| Gene and protein expression by JAK-STAT signaling after Interleukin-12  stimulation | (R-HSA-8950505) |
| Interleukin-12 signaling | (R-HSA-9020591) |
| The citric acid (TCA) cycle and respiratory electron transport | (R-HSA-1428517) |
| Chaperonin-mediated protein folding | (R-HSA-390466) |
| Interleukin-12 family signaling | (R-HSA-447115) |
| Protein folding | (R-HSA-391251) |
| Complex I biogenesis | (R-HSA-6799198) |
| Beta oxidation of octanoyl-CoA to hexanoyl-CoA | (R-HSA-77348) |
| Beta oxidation of lauroyl-CoA to decanoyl-CoA-CoA | (R-HSA-77310) |
| Beta oxidation of hexanoyl-CoA to butanoyl-CoA | (R-HSA-77350) |
| Mitochondrial protein import | (R-HSA-1268020) |
| Beta oxidation of decanoyl-CoA to octanoyl-CoA-CoA | (R-HSA-77346) |
| Depolymerisation of the Nuclear Lamina | (R-HSA-4419969) |
| RHOBTB2 GTPase cycle | (R-HSA-9013418) |
| Initiation of Nuclear Envelope (NE) Reformation | (R-HSA-2995383) |
| Protein localization | (R-HSA-9609507) |
| RHOBTB GTPase Cycle | (R-HSA-9706574) |
| Breakdown of the nuclear lamina | (R-HSA-352238) |
| Drug resistance in ERBB2 KD mutants | (R-HSA-9665230) |
| Drug-mediated inhibition of ERBB2 signaling | (R-HSA-9652282) |
| Resistance of ERBB2 KD mutants to AEE788 | (R-HSA-9665250) |

Table 2: The list of most significantly regulated Reactome pathways identified from the EL proteomics entity input.

| **Pathway Name** | **Stable Identifier** |
| --- | --- |
| Interleukin-12 family signaling | (R-HSA-447115) |
| COPI-mediated anterograde transport | (R-HSA-6807878) |
| L1CAM interactions | (R-HSA-373760) |
| Interleukin-12 signaling | (R-HSA-9020591) |
| Senescence-Associated Secretory Phenotype (SASP) | (R-HSA-2559582) |
| Post NMDA receptor activation events | (R-HSA-438064) |
| Microtubule-dependent trafficking of connexons from Golgi to the plasma  membrane | (R-HSA-190840) |
| Recycling pathway of L1 | (R-HSA-437239) |
| Transport of connexons to the plasma membrane | (R-HSA-190872) |
| Signaling by Interleukins | (R-HSA-449147) |
| ER to Golgi Anterograde Transport | (R-HSA-199977) |
| Post-chaperonin tubulin folding pathway | (R-HSA-389977) |
| Activation of NMDA receptors and postsynaptic events | (R-HSA-442755) |
| Signaling by MAP2K mutants | (R-HSA-9652169) |
| Signaling by ALK fusions and activated point mutants | (R-HSA-9725370) |
| Signaling by ALK in cancer | (R-HSA-9700206) |
| RAF-independent MAPK1/3 activation | (R-HSA-112409) |
| Prefoldin mediated transfer of substrate to CCT/TriC | (R-HSA-389957) |
| Formation of tubulin folding intermediates by CCT/TriC | (R-HSA-389960) |
| Gene and protein expression by JAK-STAT signaling after Interleukin-12  stimulation | (R-HSA-8950505) |
| Negative feedback regulation of MAPK pathway | (R-HSA-5674499) |
| Cellular Senescence | (R-HSA-2559583) |
| Activation of AMPK downstream of NMDARs | (R-HSA-9619483) |
| Formation of the cornified envelope | (R-HSA-6809371) |
| RHO GTPases activate IQGAPs | (R-HSA-5626467) |

Table 3: The list of most significantly regulated Reactome pathways identified from the TL proteomics entity input.

| **Pathway Name** | **Stable Identifier** |
| --- | --- |
| rRNA processing in the nucleus and cytosol | (R-HSA-8868773) |
| Major pathway of rRNA processing in the nucleolus and cytosol | (R-HSA-6791226) |
| rRNA processing | (R-HSA-72312) |
| Interferon Signaling | (R-HSA-913531) |
| Transcriptional regulation of white adipocyte differentiation | (R-HSA-381340) |
| PERK regulates gene expression | (R-HSA-381042) |
| Loss of MECP2 binding ability to 5hmC-DNA | (R-HSA-9022534) |
| Response of EIF2AK1 (HRI) to heme deficiency | (R-HSA-9648895) |
| Response of EIF2AK4 (GCN2) to amino acid deficiency | (R-HSA-9633012) |
| Translocation of SLC2A4 (GLUT4) to the plasma membrane | (R-HSA-1445148) |
| ATF4 activates genes in response to endoplasmic reticulum stress | (R-HSA-380994) |
| Assembly of collagen fibrils and other multimeric structures | (R-HSA-2022090) |
| Cellular response to starvation | (R-HSA-9711097) |
| Regulation of MECP2 expression and activity | (R-HSA-9022692) |
| NGF-stimulated transcription | (R-HSA-9031628) |
| Loss of MECP2 binding ability to 5mC-DNA | (R-HSA-9022538) |
| Collagen chain trimerization | (R-HSA-8948216) |
| ATF6 (ATF6-alpha) activates chaperone genes | (R-HSA-381183) |
| MECP2 regulates transcription of genes involved in GABA signaling | (R-HSA-9022927) |
| Reversible hydration of carbon dioxide | (R-HSA-1475029) |
| ATF6 (ATF6-alpha) activates chaperones | (R-HSA-381033) |
| Unfolded Protein Response (UPR) | (R-HSA-381119) |
| Interferon gamma signaling | (R-HSA-877300) |

Table 4: The list of most significantly regulated Reactome pathways identified from the transcriptomics analysis.
