## Supplementary material for "Compartmentalisation proteomics revealed endolysosomal protein network changes in a goat model of atrial fibrillation": Supplementary File 3.docx

### Supplementary File 3 contains;

### 1. Table 1: The complete list of the most significantly regulated proteins identified uniquely in EL fraction

### 2. Figure 1: The PCA plot showing the outliers of proteomics and transcriptomics biological replicates.

| **No.** | **Gene Name** | **Protein name** |
| --- | --- | --- |
| 1 | PFDN4 | Prefoldin subunit 4 |
| 2 | SMCO4 | Single-pass membrane and coiled-coil domain-containing protein 4 |
| 3 | HECTD3 | E3 ubiquitin-protein ligase HECTD3 |
| 4 | WDR78 | WD_REPEATS_REGION domain-containing protein |
| 5 | RTN3 | Reticulon |
| 6 | HSPB7 | SHSP domain-containing protein |
| 7 | RNF17 | RING finger protein 17 |
| 8 | NAV2 | Calponin-homology (CH) domain-containing protein |
| 9 | PPP1R11 | E3 ubiquitin-protein ligase PPP1R11 |
| 10 | PKIG | cAMP-dependent protein kinase inhibitor |
| 11 | COX6A2 | Cytochrome c oxidase subunit |
| 12 | HSPB2 | SHSP domain-containing protein |
| 13 | ENO2 | 2-phospho-D-glycerate hydro-lyase |
| 14 | ATP5PF | ATP synthase-coupling factor 6, mitochondrial |
| 15 | ACTC1 | Actin, alpha cardiac muscle 1 |
| 16 | POLR3D | DNA-directed RNA polymerase III subunit RPC4 |
| 17 | CLTB | Clathrin light chain |
| 18 | SFPQ | Splicing factor, proline- and glutamine-rich |
| 19 | NDUFB1 | Complex I-MNLL |
| 20 | BDH1 | D-beta-hydroxybutyrate dehydrogenase, mitochondrial |
| 21 | COBL | Protein cordon-bleu |
| 22 | ARL6IP5 | PRA1 family protein |
| 23 | MACF1 | Microtubule-actin cross-linking factor 1, isoforms 1/2/3/5 |
| 24 | MPRIP | Myosin phosphatase Rho-interacting protein |
| 25 | CXADR | Coxsackievirus and adenovirus receptor |
| 26 | PPP1R12C | Protein phosphatase 1 regulatory subunit |
| 27 | NDUFS6 | NADH dehydrogenase [ubiquinone] iron-sulfur protein 6, mitochondrial |
| 28 | LMNA | Prelamin-A/C |
| 29 | DGKD | Diacylglycerol kinase |
| 30 | MAP4 | Microtubule-associated protein |
| 31 | AUTS2 | Autism susceptibility gene 2 protein |
| 32 | NDUFB8 | NADH dehydrogenase [ubiquinone] 1 beta subcomplex subunit 8, mitochondrial |
| 33 | USP9X | Ubiquitinyl hydrolase 1 |
| 34 | RPL24 | TRASH domain-containing protein |
| 35 | UFC1 | Ubiquitin-fold modifier-conjugating enzyme 1 |
| 36 | HHATL | Protein-cysteine N-palmitoyltransferase HHAT-like protein |
| 37 | DYNLRB1 | Dynein light chain roadblock |
| 38 | ATP5PD | ATP synthase subunit d, mitochondrial |
| 39 | NDUFB7 | Complex I-B18 |
| 40 | GANAB | Gal_mutarotas_2 domain-containing protein |
| 41 | LARS1 | Leucyl-tRNA synthetase |
| 42 | VAPA | Vesicle-associated membrane protein-associated protein A |
| 43 | SIRT3 | NAD-dependent protein deacetylase |
| 44 | SMYD2 | N-lysine methyltransferase SMYD2 |
| 45 | ACADSB | Short/branched chain specific acyl-CoA dehydrogenase |
| 46 | GORASP2 | GRASP55_65 domain-containing protein |
| 47 | GLRX | Glutaredoxin domain-containing protein |
| 48 | NDUFB9 | Complex I-B22 |
| 49 | FN1 | Fibronectin |
| 50 | PLG | Plasminogen (Fragment) |
| 51 | RPLP2 | 60S acidic ribosomal protein P2 |
| 52 | TAGLN2 | Transgelin |
| 53 | CDNF | Cerebral dopamine neurotrophic factor |
| 54 | NDUFA6 | NADH dehydrogenase [ubiquinone] 1 alpha subcomplex subunit 6 |
| 55 | EHD1 | EH domain-containing protein 1 |
| 56 | CCT2 | CCT-beta |
| 57 | COPS7A | PCI domain-containing protein |
| 58 | MTX2 | Metaxin-2 |
| 59 | GAA | P-type domain-containing protein |
| 60 | FLNB | Filamin-B |
| 61 | RNH1 | Ribonuclease inhibitor |
| 62 | SLC25A6 | ADP/ATP translocase 3 |
| 63 | ME1 | Malic enzyme |
| 64 | MEMO1 | Mediator of ErbB2-driven cell motility 1 |
| 65 | ETHE1 | Lactamase_B domain-containing protein |
| 66 | NCOR2 | Nuclear receptor corepressor 2 |
| 67 | DST | Dystonin |
| 68 | IPO5 | Importin N-terminal domain-containing protein |
| 69 | NAE1 | NEDD8-activating enzyme E1 regulatory subunit |
| 70 | PLPBP | Pyridoxal phosphate homeostasis protein |
| 71 | SMS | PABS domain-containing protein |
| 72 | RTN4IP1 | PKS_ER domain-containing protein |
| 73 | GRHPR | Glyoxylate reductase/hydroxypyruvate reductase |
| 74 | PDXP | Pyridoxal phosphate phosphatase |
| 75 | PSMB4 | Proteasome subunit beta |
| 76 | TUBB6 | Tubulin beta chain |
| 77 | GPD1 | Glycerol-3-phosphate dehydrogenase [NAD(+)] |
| 78 | FDPS | Farnesyl pyrophosphate synthase |
| 79 | CBX1 | Chromobox protein homolog 1 |
| 80 | PCBD2 | 4a-hydroxytetrahydrobiopterin dehydratase |
| 81 | EIF3E | Eukaryotic translation initiation factor 3 subunit E |
| 82 | PLBD1 | Phospholipase B-like |
| 83 | PLIN3 | Perilipin |
| 84 | VPS25 | ESCRT-II complex subunit VPS25 |
| 85 | EOGT | EGF domain-specific O-linked N-acetylglucosamine transferase |
| 86 | MAPKAPK3 | Non-specific serine/threonine protein kinase |
| 87 | SNRPD3 | Small nuclear ribonucleoprotein Sm D3 |
| 88 | CAVIN1 | Caveolae-associated protein 1 |
| 89 | CORO1C | Coronin |
| 90 | MAT2A | S-adenosylmethionine synthase |
| 91 | cytb | Cytochrome b |
| 92 | ITGAX | VWFA domain-containing protein |
| 93 | PMPCB | Beta-MPP |
| 94 | ASS1 | Argininosuccinate synthase |

Table 1: The most significantly regulated proteins identified uniquely in EL fraction

Star Methods

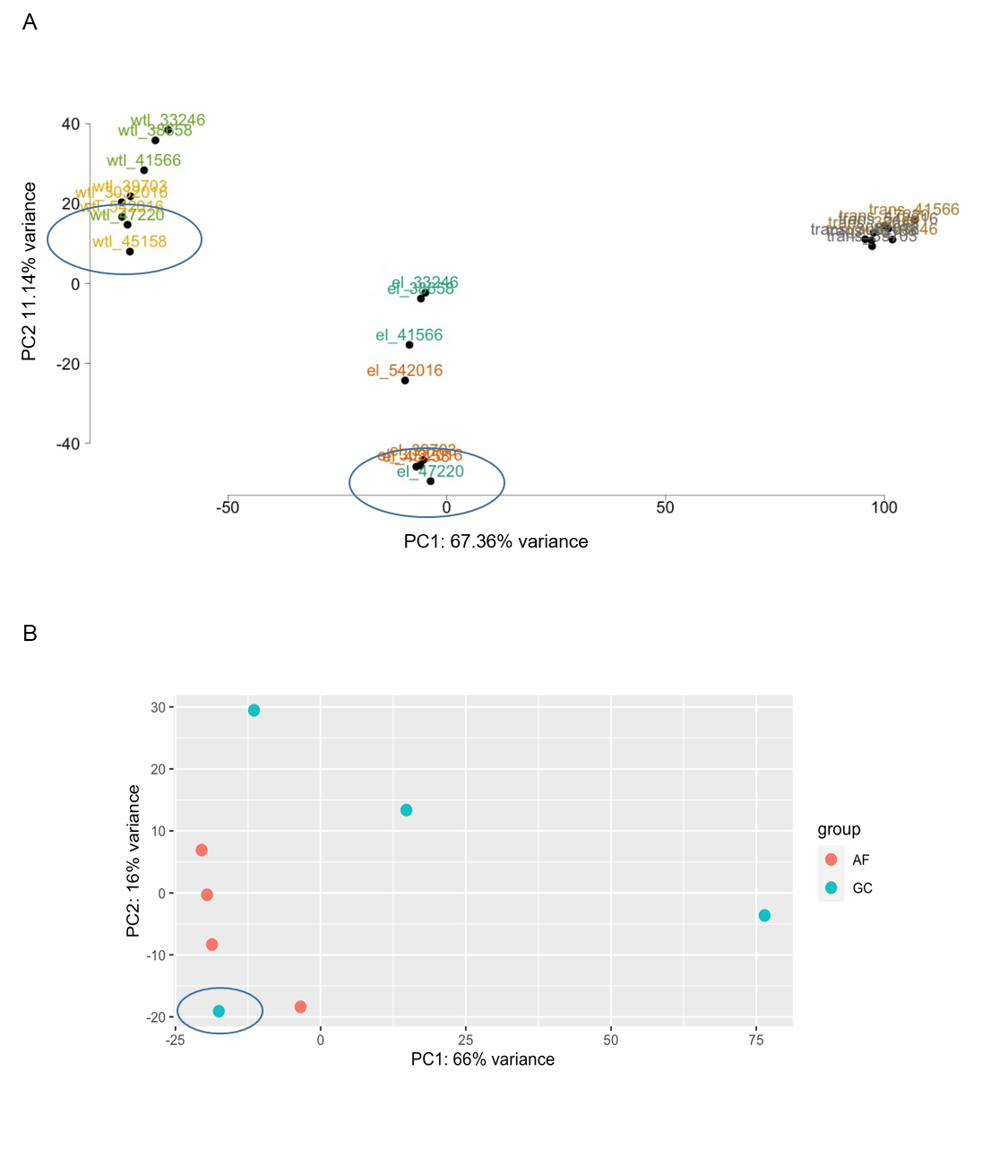

Figure 1: The PCA plots showing the outliers of proteomics and transcriptomics biological replicates. **A.** The shared PCA plot for the proteomics of whole tissue lysate and Endolysosome fraction compared to transcriptomics shows that Sham model 47220 and AF model 45158 are deviated from rest of the proteomics samples. **B.** PCA plot of the transcriptomics shows the deviation of the goat control/ sham model sample no. 33246 (AF = Atrial Fibrillation, GC= Goat Control/ Sham).
