## Supplementary material for "Compartmentalisation proteomics revealed endolysosomal protein network changes in a goat model of atrial fibrillation": Supplementary File 4.docx

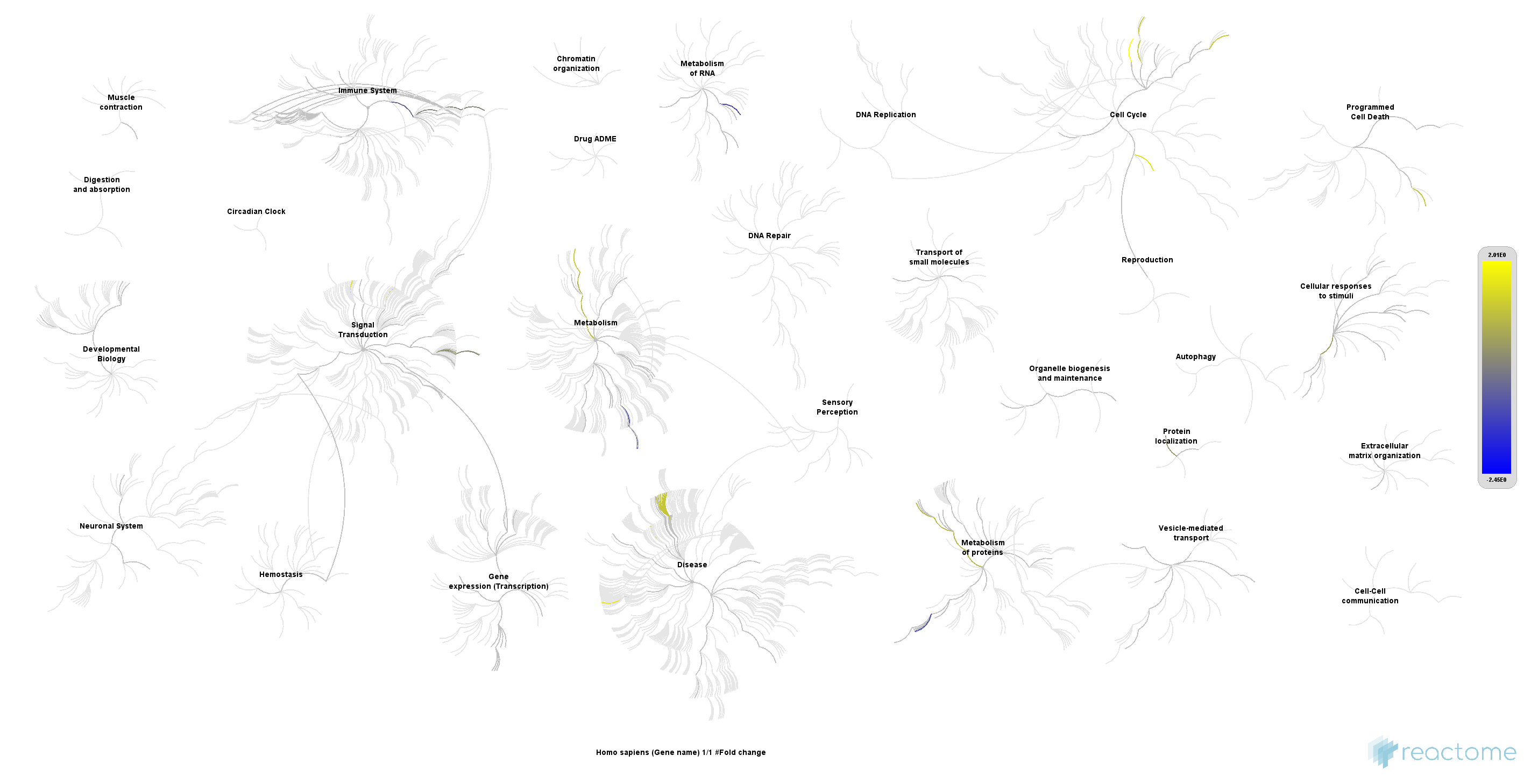
Supplementary File 4 Figure 1: EL most significant proteins displayed in a Reactome genome-wide, hierarchical visualization of pathways in a space filling graph. Highest up and down regulated pathways are depicted in Yellow and blue.
