## Supplementary figures and images for "Compartmentalisation proteomics revealed endolysosomal protein network changes in a goat model of atrial fibrillation"

### Supplementary File 4.tif

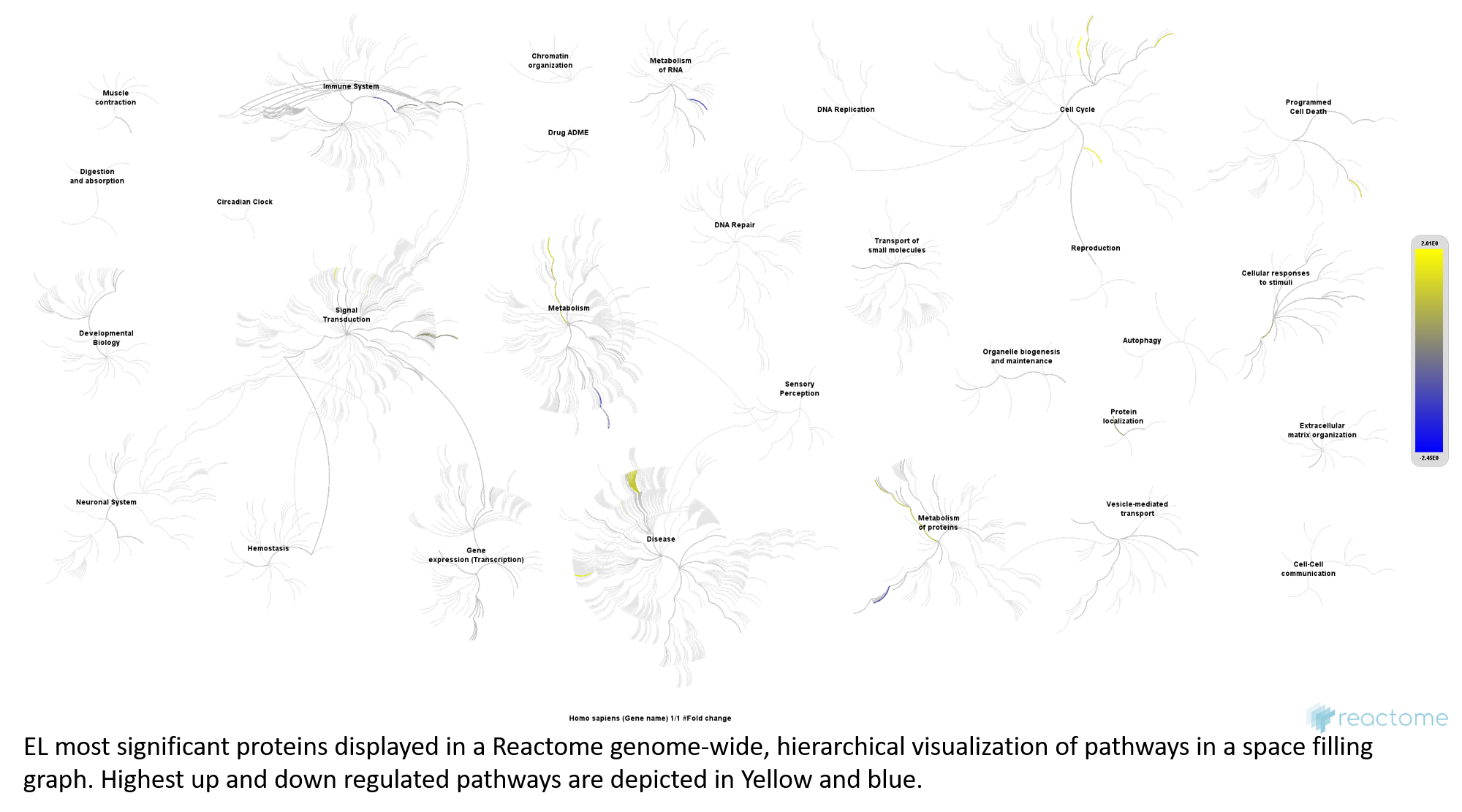

### Supplementary File 5.tif

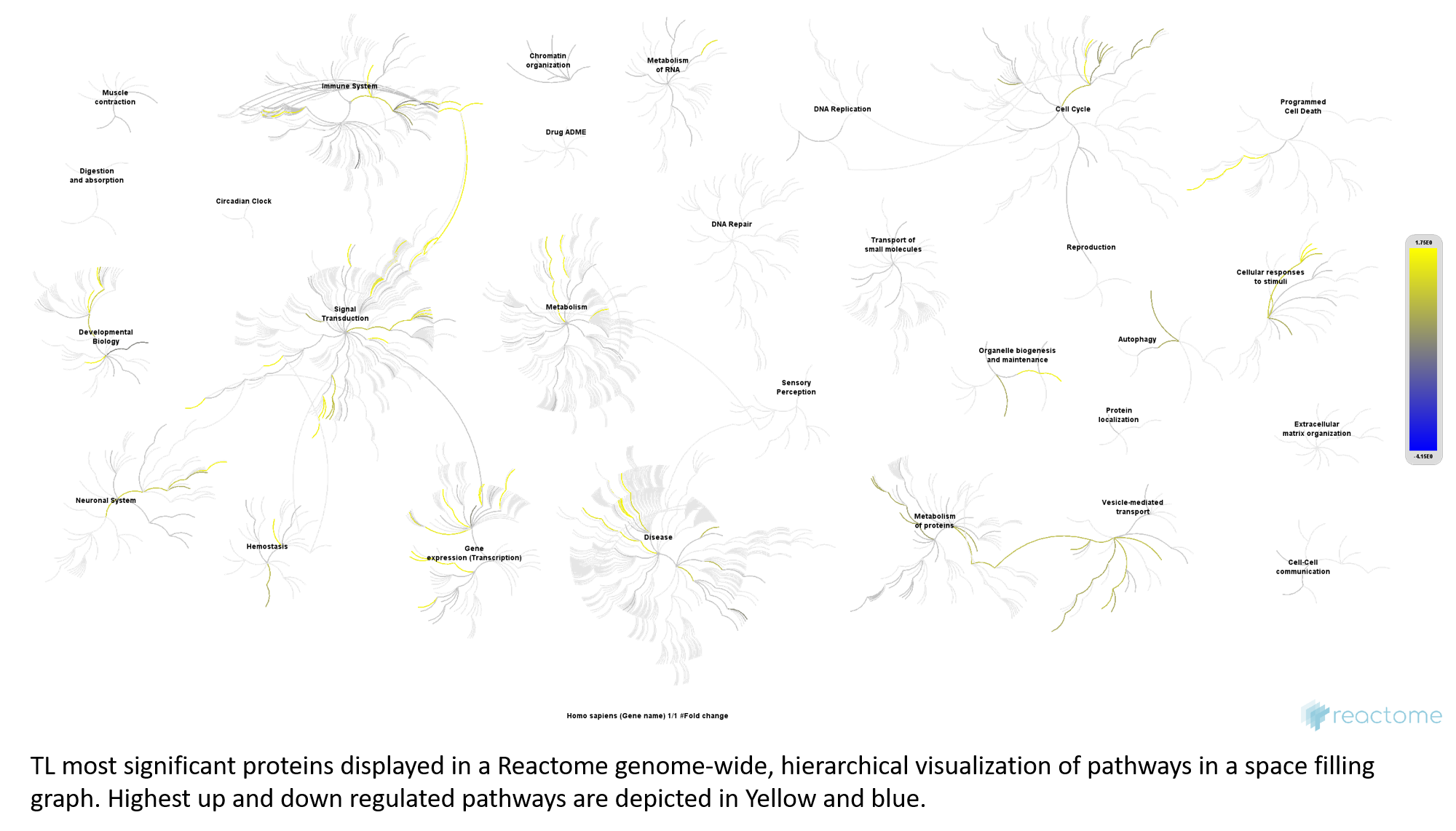

### Supplementary File 6.tif

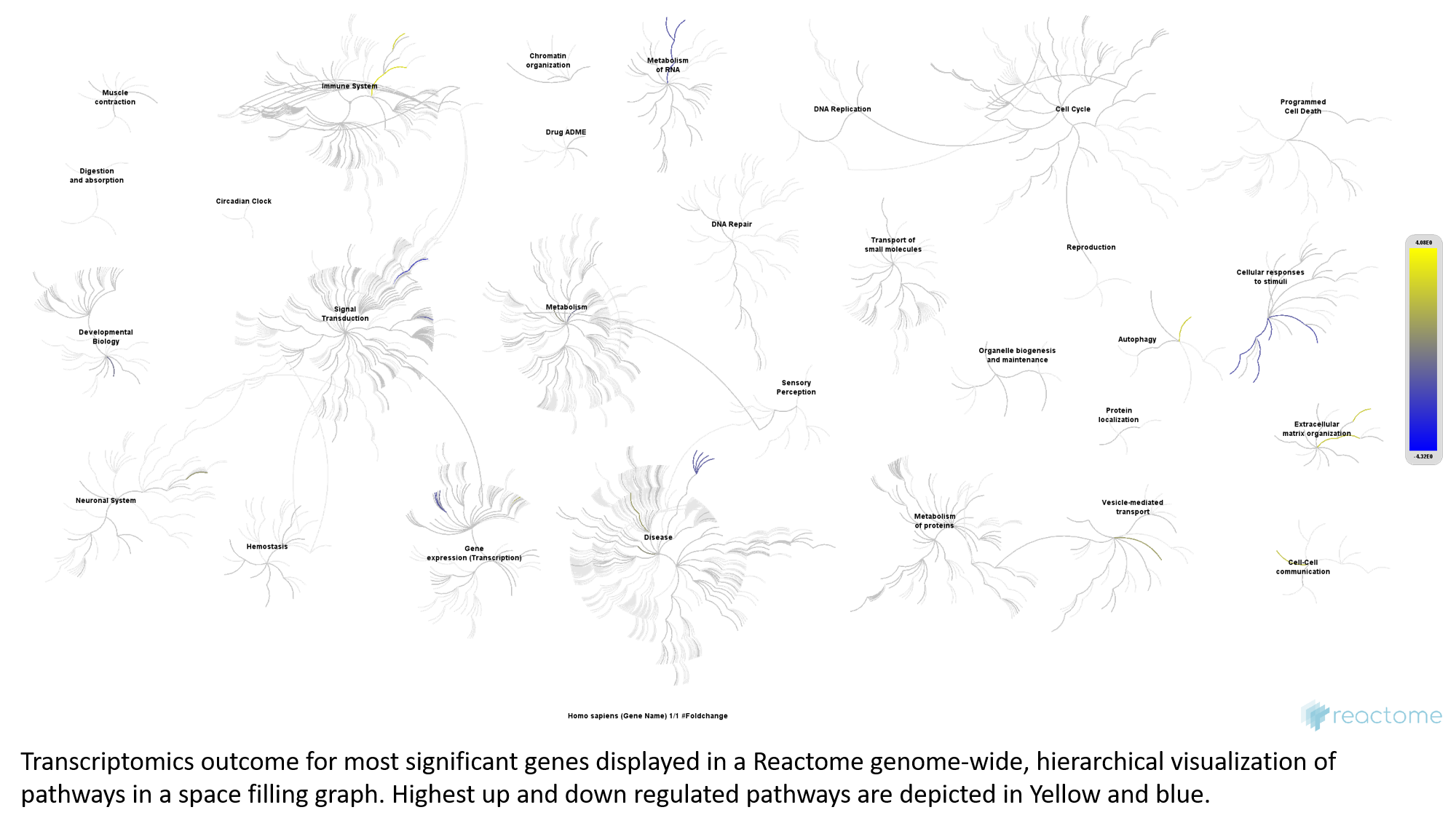
